# Genetic variant pathogenicity prediction trained using disease-specific clinical sequencing datasets

**DOI:** 10.1101/334235

**Authors:** Perry Evans, Chao Wu, Amanda Lindy, Dianalee A. McKnight, Matthew Lebo, Mahdi Sarmady, Ahmad N. Abou Tayoun

## Abstract

Recent advances in DNA sequencing technologies have expanded our understanding of the molecular underpinnings for several genetic disorders, and increased the utilization of genomic tests by clinicians. Given the paucity of evidence to assess each variant, and the difficulty of experimentally evaluating a variant’s clinical significance, many of the thousand variants that can be generated by clinical tests are reported as variants of unknown clinical significance. However, the creation of population-scale variant databases can significantly improve clinical variant interpretation. Specifically, pathogenicity prediction for novel missense variants can now utilize features describing regional variant constraint. Constrained genomic regions are those that have an unusually low variant count in the general population. Several computational methods have been introduced to capture these regions and incorporate them into pathogenicity classifiers, but these methods have yet to be compared on an independent clinical variant dataset. Here we introduce one variant dataset derived from clinical sequencing panels, and use it to compare the ability of different genomic constraint metrics to determine missense variant pathogenicity. This dataset is compiled from 17,071 patients surveyed with clinical genomic sequencing for cardiomyopathy, epilepsy, or RASopathies. We further utilize this dataset to demonstrate the necessity of disease-specific classifiers, and to train PathoPredictor, a disease-specific ensemble classifier of pathogenicity based on regional constraint and variant level features. PathoPredictor achieves an average precision greater than 90% for variants from all 99 tested disease genes while approaching 100% accuracy for some genes. Accumulation of larger clinical variant datasets and their utilization to train existing pathogenicity metrics can significantly enhance their performance in a disease and gene-specific manner.

## Introduction

Comprehensive sequencing has become the cornerstone of genomic medicine and research. However, unlike previous targeted or single gene testing, multigene sequencing can yield thousands of rare variants often requiring manual clinical correlation and interpretation. Unlike synonymous (or silent) and loss of function (mainly nonsense, frameshift, and canonical splice site) variants where the impact on the protein can be relatively easily predicted, novel missense variants are the most challenging to interpret, often leading to inconclusive genomic reports and leaving clinicians and families with daunting uncertainties and anxieties. On the other hand, researchers are currently incapable of studying the impact of every possible missense variant in the ∼20,000 genes of the human genome. Therefore, novel clinical-grade approaches are needed to assist clinicians and researchers in determining the pathogenicity of missense variants.

Machine learning has yielded several pathogenicity prediction tools built with variant features and previously assigned pathogenic and benign labels. Collections of labeled variant labels for classifier training and testing include the Human Gene Mutation Database (HGMD) (Stenson et al. 2009), the Leiden Open Variation Database (Fokkema et al. 2011), and ClinVar (Landrum et al. 2016). Additionally, frequently occurring variants from databases like the Genome Aggregation Database (gnomAD) (Lek et al. 2016) are used as a substitute for benign variants. Variant features can describe single positions, like genomic sequence context and amino acid conservation, or regions that contain the variant, like protein domains and variation constraint.

Two uses of simple region features are seen in the Functional Analysis through Hidden Markov Models (FATHMM) (Shihab et al. 2013) and the variant effect scoring tool (VEST) (Carter et al. 2013). FATHMM and VEST were found to be the most important features for determining pathogenicity in an ensemble of 18 prediction scores called REVEL (Ioannidis et al. 2016). VEST distinguished disease missense variants in HGMD from high frequency (allele frequency >1%) missense variants from the Exome Sequencing Project (ESP) (Auer et al. 2016) using a random forest with 86 features from the SNVBox database (Wong et al. 2011). These features describe amino acid substitutions, regional amino acid composition, conservation scores, local protein structure, and annotations of functional protein sites. FATHMM scored variants by their conservation in homologous sequences, weighted by the tolerance of each variant’s protein family (Pfam) domain or SUPERFAMILY (Gough et al. 2001) to mutations observed in HGMD and the set of functionally neutral UniProt variants (The UniProt Consortium 2017). VEST’s inclusion of functional protein sites, and FATHMM’s Pfam domain tolerance consideration enabled them to capture regional protein features such as domain structure and conservation, but did not capture regional tolerance to genetic variation.

Large variant collections like the Exome Aggregation Consortium (ExAC) dataset (Lek et al. 2016) and gnomAD have enabled metrics that summarize purifying selection within genomic regions. Regions with high purifying selection are constrained and have less population variation than expected. Regions with low purifying selection are unconstrained and have equal or more population variation compared to expectation. With this knowledge, classifiers might flag a variant as pathogenic if it lies within a genomic region that selects against variants (Amr et al. 2017). Here we examine three constraint metrics derived from ExAC or gnomAD. One such constraint metric is the constrained coding region (CCR) percentile, which compared observed variant counts from gnomAD to those predicted by CpG density (Havrilla et al. 2019). A similar feature called missense depletion was constructed for the MPC (missense badness, PolyPhen-2, and constraint) pathogenicity classifier of *de novo* missense variants (Samocha et al. 2017). MPC’s missense depletion feature was measured as the fraction of expected ExAC variation that was observed in exons. Only ExAC variants with minor allele frequencies below 0.1% were considered. The expected rate of rare missense variants was based on a model that utilized both gene and sequence context specific mutation rates (Samocha et al. 2014). An additional pathogenicity feature introduced by MPC was missense badness, which accounted for an amino acid substitution’s increase in deleteriousness when it occurs in a missense-constrained region. The third constraint metric is the missense tolerance ratio (MTR), which was calculated in 31 codon windows using missense and synonymous variant frequencies from ExAC and gnomAD (Traynelis et al. 2017). MTR is the ratio of the observed missense variant fraction to the missense variant fraction calculated from all possible variants in the window when all nucleotide changes are equally likely. A variant in a low MTR region is expected to have a high chance of being pathogenic.

In this manuscript, we evaluate the effectiveness of region based pathogenicity predictors in a clinical setting. We introduce three patient variant training datasets gathered from clinical sequencing panels for cardiomyopathy, epilepsy, and RASopathies. These datasets cover 17,071 patients. All variants have been manually classified by two main clinical laboratories, whose members significantly contributed to the development of the American College of Medical Genetics and Genomics/Association for Molecular Pathology (ACMG/AMP) sequence variant interpretation guidelines (Richards et al. 2015). We use each dataset to compare CCR, FATHMM, missense badness, missense depletion, MTR, and VEST, and to train disease-specific predictors. These clinical variant sets have the advantage of being consistently reviewed in a clinically sound manner, and originate from focused disease studies. This allows us to explore the hypothesis that disease-specific classifiers, first introduced for smaller gene sets (Homburger et al. 2016; Tavtigian et al. 2008), are better than general genome wide classifiers. We also introduce PathoPredictor, a disease-specific pathogenicity score trained with clinical sequencing panel variants to combine the pathogenicity scores compared here.

## Results

### Variants and genes studied

We focus on patient variants from three disease panels: cardiomyopathy, epilepsy, and RASopathies (Figure 1). We also investigate the subset of epilepsy dominant genes: *CDKL5, KCNQ2, KCNQ3, PCDH19, SCN1A, SCN1B, SCN2A, SCN8A, SLC2A1, SPTAN1, STXBP1*, and *TSC1*. These genes account for a large number of epilepsy pathogenic variants and, since they follow a dominant inheritance pattern, might have distinct characteristics impacting variant prediction relative to all other epilepsy genes (see below). For each disease variant set, we compare the performance of CCR, FATHMM, missense badness, missense depletion, MTR, and VEST using panel and ClinVar variants with pathogenicity labels. We also build PathoPredictor, an ensemble classifier of pathogenicity, and test it with variants from ClinVar not found in our disease panel variant sets. To ensure the reliability of ClinVar variant pathogenicity labels, we examine only unambiguously pathogenic or benign variants, and split ClinVar into two variant groups: all ClinVar variants and those that have been reviewed. Note that few of these variant collections have an equal amount of pathogenic and benign variants, with a drastic imbalance for cardiomyopathy panel variants.

**Figure 1.**
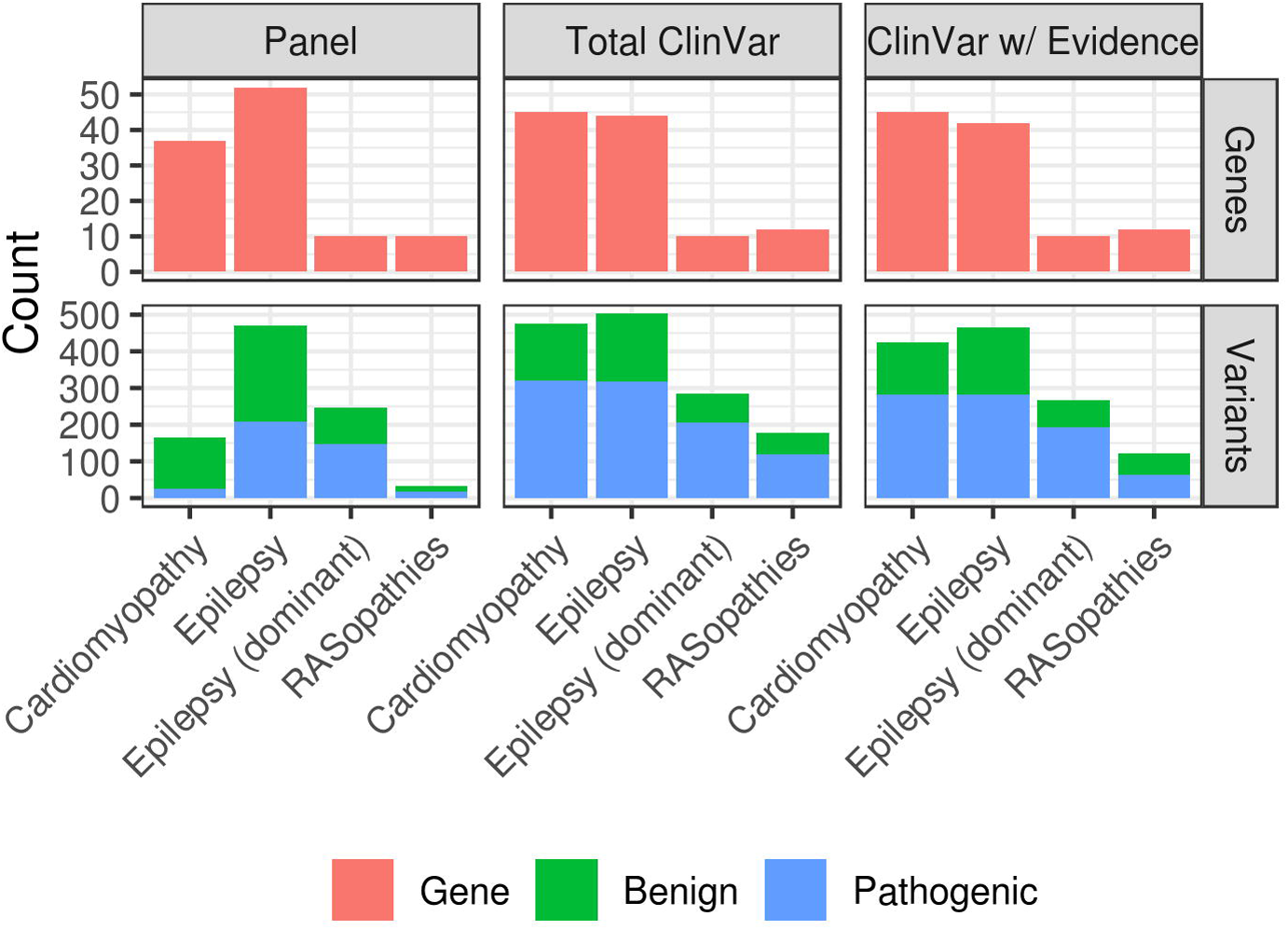
Study datasets. Missense variant and gene counts are shown by disease panel and ClinVar variant set. We only use ClinVar variants from panel genes, and consider either any ClinVar variant (Total ClinVar), or ClinVar variants that have been reviewed (ClinVar w/ Evidence). ClinVar variants are restricted to those with no conflicting pathogenicity assignments, and any genomic position from the panel data is removed from the ClinVar variant sets.

### Disease-specific classifier evaluation

We use disease panel and ClinVar datasets to compare pathogenicity classifiers, and to train and test PathoPredictor (Figure 2). For each disease, we used panel and ClinVar variants to build precision-recall curves using pathogenicity scores from CCR, FATHMM, missense badness, missense depletion, MTR, and VEST. These curves were summarized using average precision. To evaluate PathoPredictor, we examined each disease panel gene, trained a model using disease panel variants not found in the selected gene, and tested the model using panel and ClinVar variants from the gene. By training PathoPredictor with disease-specific variants, we collect variants that belong to genes that are more likely to share a common biological pathway, and might have similar tolerance to variants.

**Figure 2.**
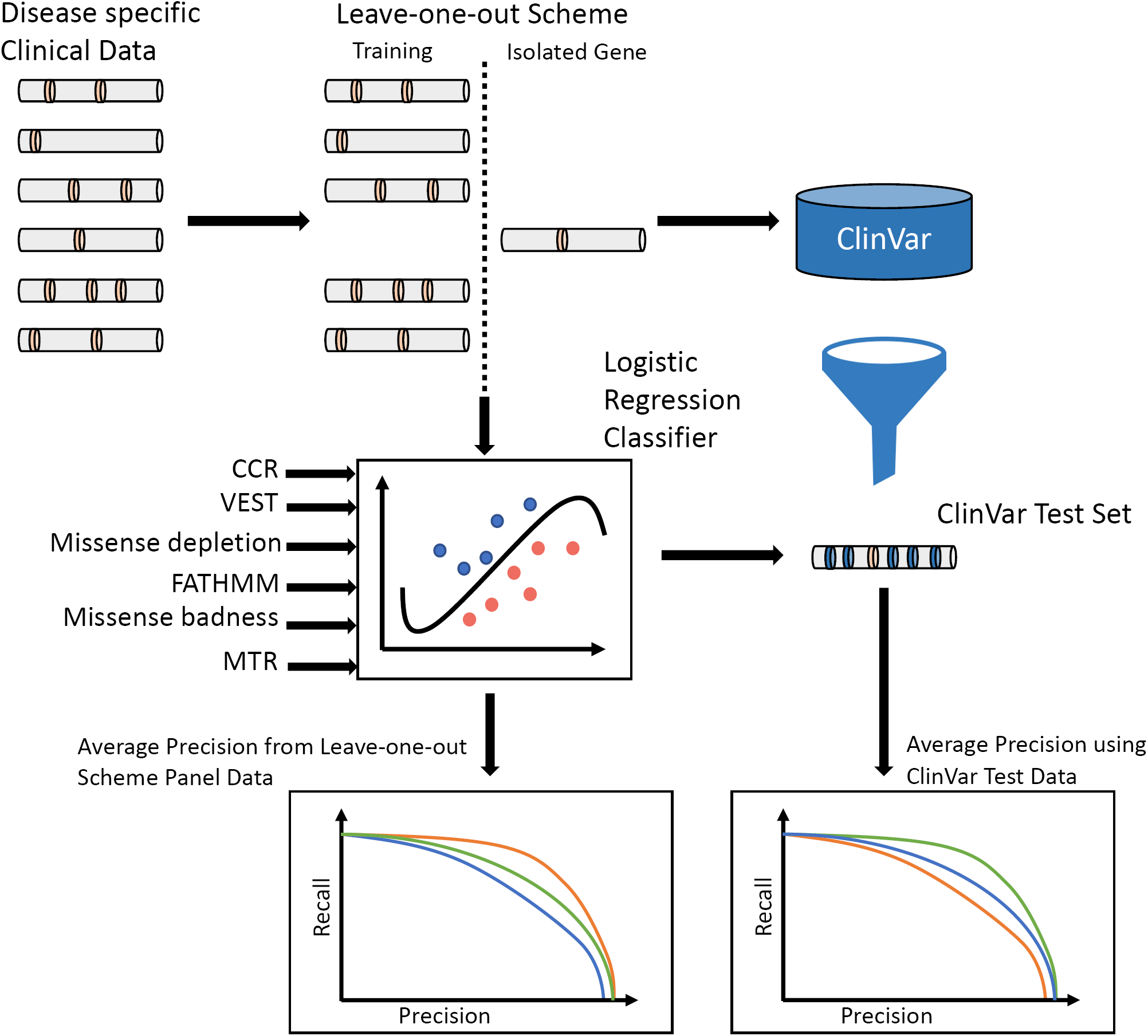
Method description. Our goal is to build disease-specific classifiers of missense variant pathogenicity using variants from clinical panels. For all genes in a disease panel, we trained a model using variants from all other genes except the gene in question, and tested the model using variants from that gene of interest. We then used ClinVar variants from the gene of interest as an independent test set. Test results were summarized as average precision scores.

PathoPredictor predicts the pathogenicity of disease panel variants with an average precision higher than that obtained with any single feature (Figure 3). The average precision of PathoPredictor is greater than 90% for all disease panels. CCR has the highest average precision among the single features. The majority of the time, PathoPredictor’s performance is significantly better than that of any single feature in the 24 comparisons made across six features and four panel variant sets (18 of the total 24 comparisons in all four panels, p < 0.05). PathoPredictor was not significantly better than CCR for the dominant epilepsy genes, where it is expected that regional constraint is most critical (see above). The remaining five exceptions were in the RASopathies and cardiomyopathy panels, which both had the lowest variant count (Figure 1). However, when evaluating with more variants (below), a significant advantage of PathoPredictor over all single features was observed (Figure 4, p < 0.03).

**Figure 3.**
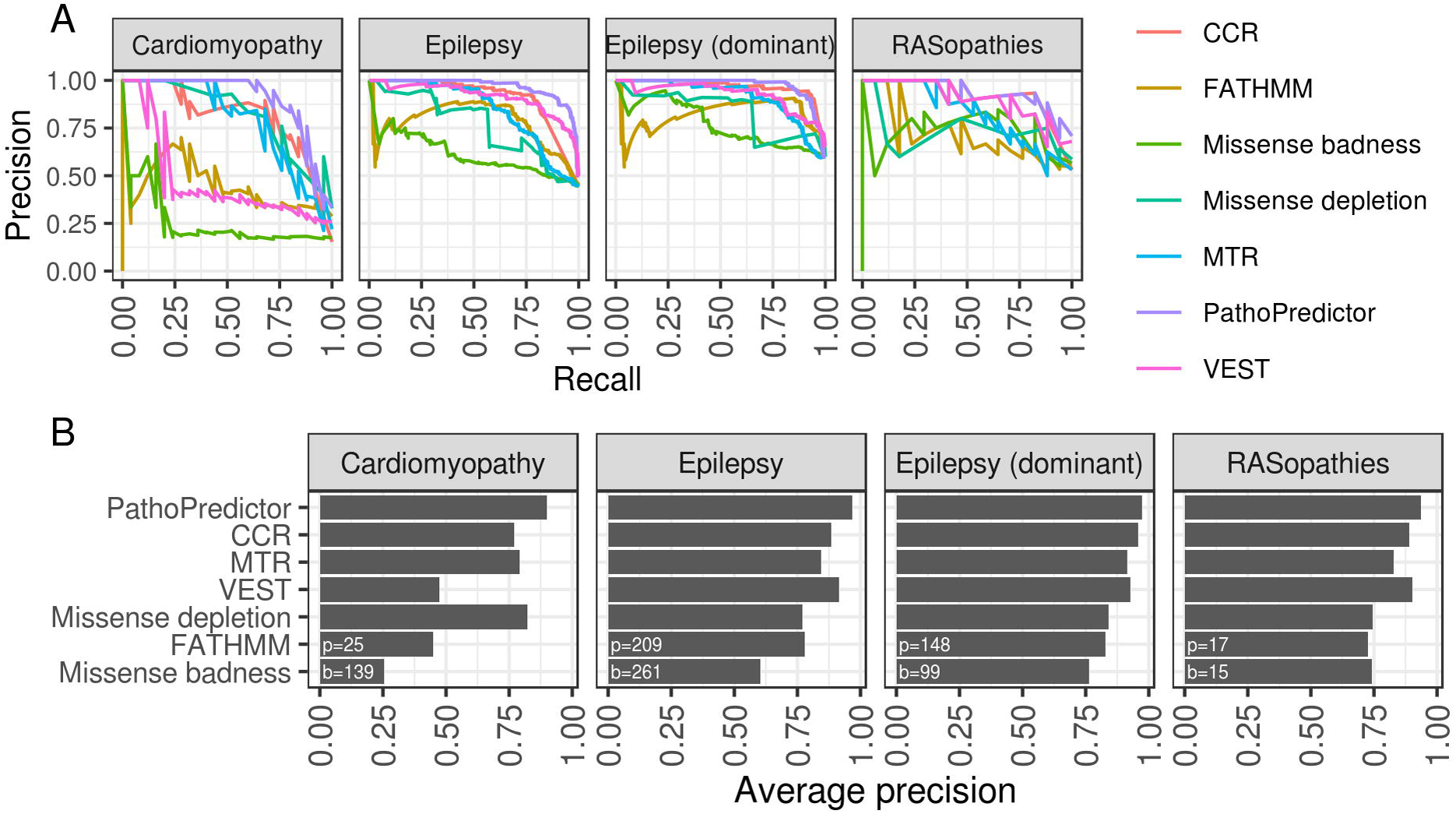
Disease-specific classifier performance using disease panel cross-validation. For each disease panel, we used a hold-one-gene-out approach to evaluate a logistic regression model’s ability to predict pathogenicity. For all genes in a disease panel, we trained PathoPredictor using variants from all other genes, and tested the model using variants from the gene of interest. Using the held-out gene variant prediction scores, we A) computed a precision-recall curve and B) summarized the curve as the average precision. We then computed a precision-recall curve for each individual feature using untransformed scores. The numbers of pathogenic (p) and benign (b) variants investigated are shown at the bottom left of each panel in B). For all epilepsy variants, PathoPredictor performs significantly better than any single feature (p < 10^-4^), and PathoPredictor only failed to be significantly better in 6 of the 24 total feature comparisons (CCR, VEST, and missense depletion for RASopathies, CCR for dominant epilepsy genes, and missense depletion and MTR for cardiomyopathy).

**Figure 4.**
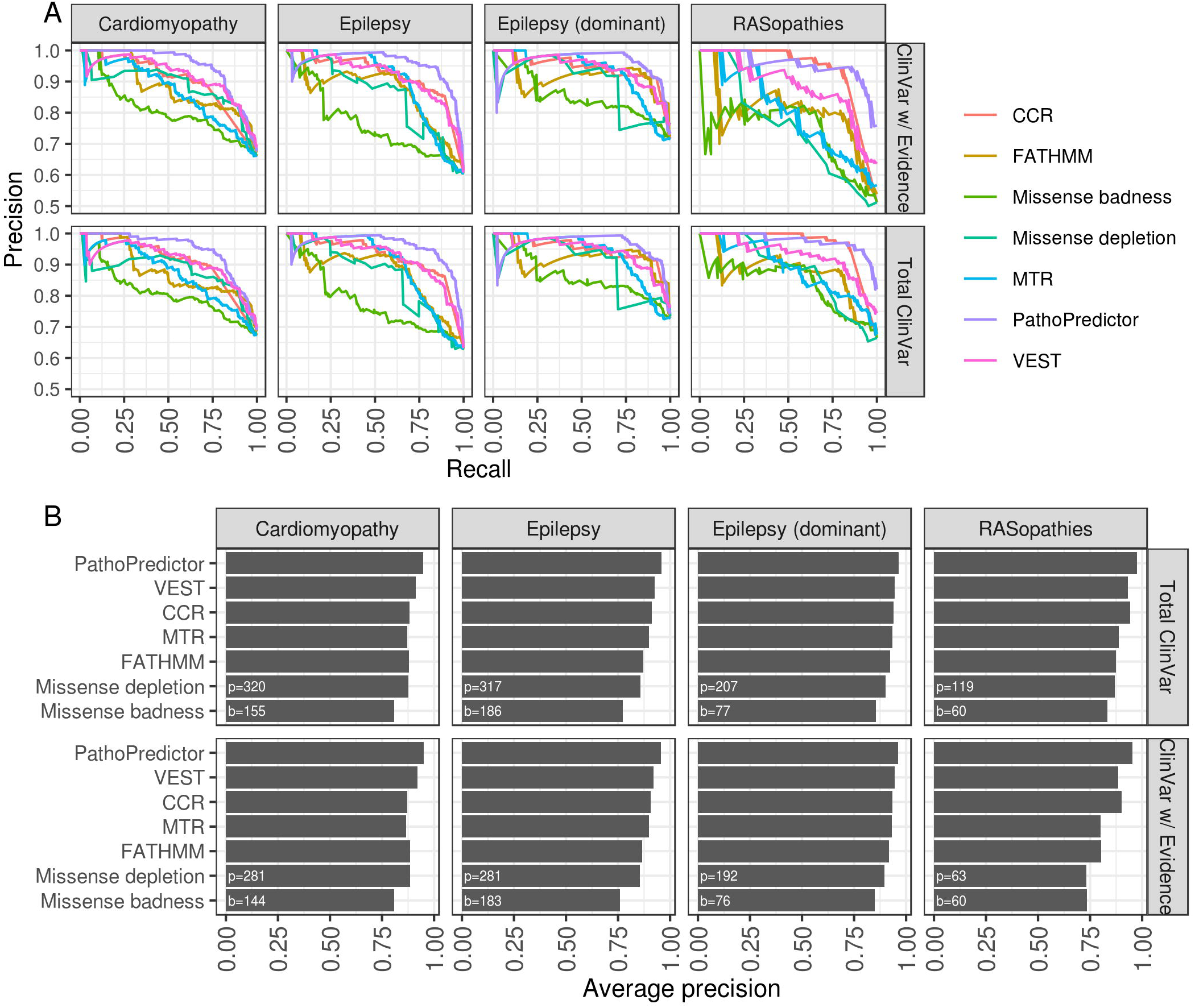
Disease-specific classifier performance using disease panel data for training and ClinVar data for testing. For each disease panel, we applied the hold-one-gene-out models from Figure 3 to ClinVar variants from the held-out gene to obtain pathogenicity prediction scores. We compared PathoPredictor to each feature using A) a precision-recall curves and B) average precisions summarizing each curve. We used either all ClinVar variants (Total ClinVar), or ClinVar variants with a review status that included at least one submitter or an expert panel (Clinvar w/ Evidence). The numbers of pathogenic (p) and benign (b) variants investigated are shown at the bottom left of each panel in B). PathoPredictor performs significantly better than any single feature when examining all ClinVar variants (p < 0.03).

PathoPredictor’s performance was next evaluated using a larger independent variant set from ClinVar (Figure 4). For each disease panel, we find that PathoPredictor performs significantly better than any single feature when examining all ClinVar variants (p < 0.03). PathoPredictor has similar performance when using all of ClinVar and the reviewed subset of ClinVar, achieving an average precision greater than 95% for all variant sets. The poor average precision obtained when using VEST, FATHMM, and missense badness to predict cardiomyopathy panel variants was not replicated using ClinVar variants in cardiomyopathy panel genes. This discrepancy can be attributed to the lower number of cardiomyopathy panel variants, especially pathogenic variants (Figure 1). PathoPredictor showed consistent results for RASopathy ClinVar and panel variants, however, given the larger number of ClinVar variants, the improved performance of PathoPredictor was now statistically significant in ClinVar (p < 0.004).

As a further test of PathoPredictor, we trained PathoPredictor with ClinVar variants, and evaluated each classifier with disease panel variants (Figure 5). For training we used either total ClinVar variants, or those that had been reviewed, and restricted ClinVar training variants to genes from the disease panel used for evaluation. PathoPredictor achieved an average precision of at least 90% for all evaluations. PathoPredictor performed better than each of its six features (p < 0.05), except for missense depletion for cardiomyopathy panel variants, CCR, missense badness, and VEST for RASopathy panel variants (most likely due to limited cardiomyopathy and RASopathy panel variants), and CCR for dominant epilepsy panel variants. These findings are consistent with the disease panel hold-one-gene-out approach in Figure 3.

**Figure 5.**
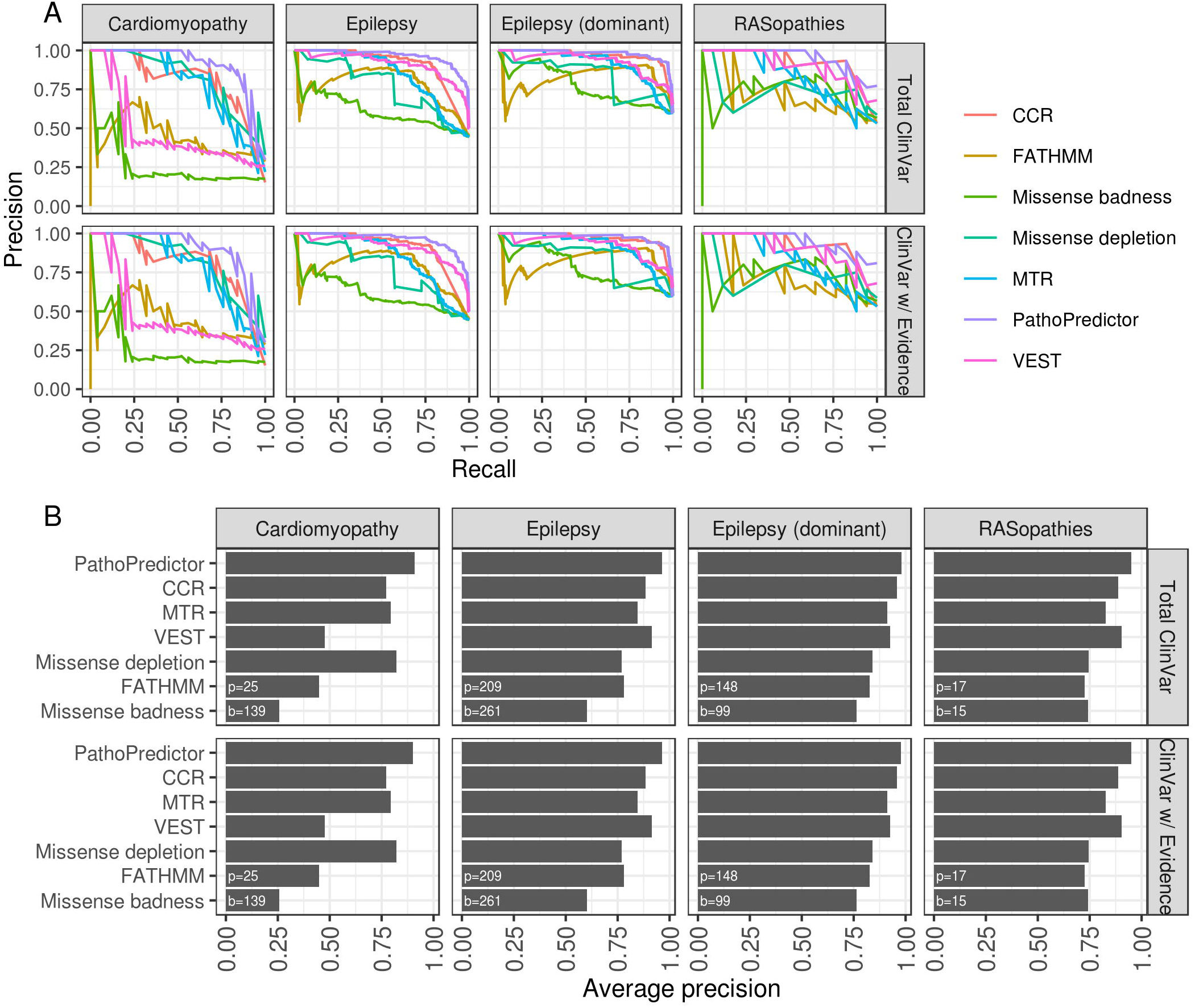
Disease-specific classifier performance using ClinVar data for training and disease panel data for testing. For each disease panel, we collected ClinVar variants in panel genes, using either all ClinVar variants (Total ClinVar), or reviewed ClinVar variants (ClinVar w/ Evidence). PathoPredictor training and evaluation for each disease panel proceeded with a hold-one-gene approach. Disease panel variants from the gene of interest were used for evaluation, and ClinVar variants from all remaining disease panel genes were used for training. Using the held-out gene variant prediction scores, we A) computed a precision-recall curve and B) summarized the curve with its average precision. We then computed a precision-recall curve for each individual feature using untransformed scores. The numbers of pathogenic (p) and benign (b) variants investigated are shown at the bottom left of each panel in B). PathoPredictor performed better than each of its six features (p < 0.05), except for missense depletion for cardiomyopathy panel variants, CCR, missense badness, and VEST for RASopathy panel variants, and CCR for dominant epilepsy panel variants.

Assessing the gene-wise performance of PathoPredictor is challenging because most genes have a small variant sample size. However, some genes with high variant count were found to best demonstrate the utility of PathoPredictor (Figure 6A). When using the hold-one-gene-out approach for training and evaluation on disease panel data, PathoPredictor had an accuracy of 95% for the 27 pathogenic and 10 benign variants in *KCNQ2*. When training on panel variants, and validating with ClinVar variants, PathoPredictor had a 96% accuracy for 41 pathogenic and 6 benign ClinVar variants in *KCNQ2*. High accuracies were also observed for *RAF1, SCN2A, SCN5A*, and *STXBP1*.

**Figure 6.**
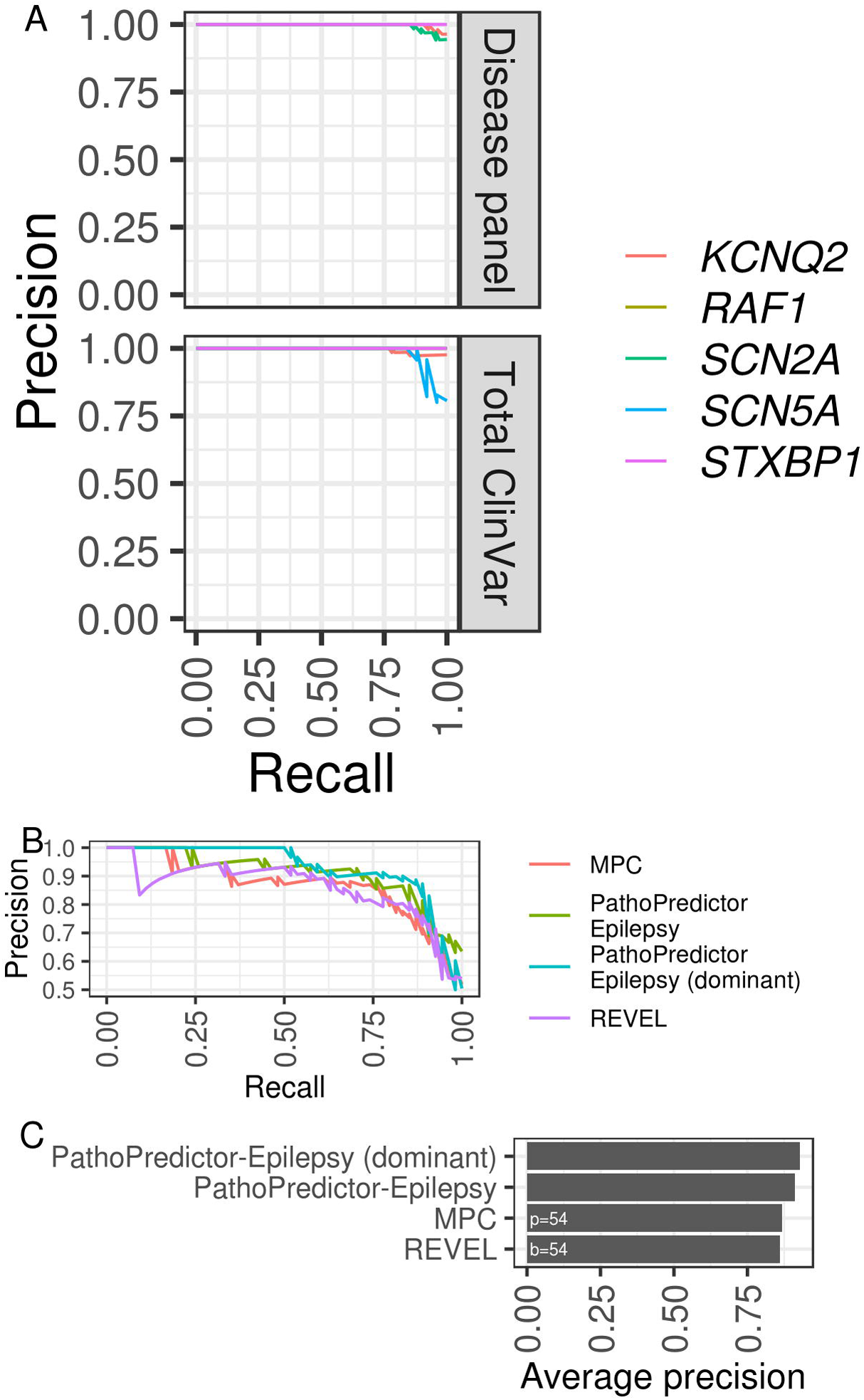
PathoPredictor performance. A) Precision-recall curves are shown for select genes evaluated during cross-validation with the disease panel dataset and tested with ClinVar variants. The curve for *RAF1* closely follows, and is obscured by that of *SCN2A*. For KCNQ2, PathoPredictor has an accuracy of 95% for panel variants and 96% for ClinVar variants. B) PathoPredictor epilepsy-specific classifiers were compared to REVEL and MPC. *De novo* missense variants in epilepsy panel genes were used as pathogenic variants. Epilepsy panel gene missense variants from unaffected siblings of autism spectrum disorder patients were used as benign variants. PathoPredictor was trained as in Figure 4, but only utilizing the full and dominant epilepsy datasets. Variants were filtered using the same methods applied to ClinVar variants, and additional filters were applied to remove training data for MPC. C) We summarized each scoring metric’s precision-recall curve as the average precision. Both PathoPredictor classifiers achieved a greater average precision than REVEL (p < 0.05), and the dominant epilepsy classifier performed better than MPC (p < 0.05). P: pathogenic; b: benign.

### Comparing PathoPredictor with MPC and REVEL

REVEL is a state-of-art ensemble classifier of pathogenicity. Built using 18 prediction scores, it has more features than PathoPredictor, but does not contain recent genomic constraint features like CCR and missense depletion. MPC is a recent classifier of pathogenicity that was trained to combine genomic constraint features with PolyPhen-2 scores using missense pathogenic ClinVar variants and a benign variant set constructed using missense variants with 1% or higher ExAC frequencies. We utilized the set of *de novo* variants to compare PathoPredictor, MPC, and REVEL (Figure 6B, see Methods). We focused on our PathoPredictor epilepsy classifiers, as they were expected to be most relevant to the neurodevelopmental and autism disorder variants from the validation *de novo* dataset. We found that the dominant epilepsy trained PathoPredictor achieved more than 94% average precision which was significantly higher than that of REVEL or MPC (p < 0.05, Figure 6C). Both PathoPredictor classifiers achieved a greater average precision than REVEL (p < 0.05, Figure 6C).

## Discussion

We have shown that the efficacy of variant pathogenicity prediction varies by disease, whereby each disease dictates a unique combination of classifier features. We have also presented PathoPredictor, a new missense variant pathogenicity predictor trained with variants from clinical sequencing results to produce pathogenicity scores from disease-specific combinations of regional constraint and variant features. PathoPredictor achieves an average precision greater than its components: CCR, FATHMM, missense depletion, missense badness, MTR, Pfam domain status, and VEST. FATHMM, Pfam domain status, and VEST capture regional constraint by using domains and protein families, while CCR, missense depletion, missense badness, and MTR locate genomic regions with less natural population variants than expected by null models of variation.

The evaluation of PathoPredictor and other variant classification tools is limited by available data describing pathogenic and benign variants. Ideally, this data would come from unbiased functional, mechanistic, tissue-based studies. Since these dataset do not exist in large quantities, we have chosen to utilize ClinVar and established variant interpretation protocols to determine a ground truth for variant pathogenicity. The performance of ParhoPredictor is dependent on the quality of these annotations, and additional functional studies are needed to construct better databases for the training and evaluation of pathogenicity classifiers.

CCR was determined to be the most useful feature for classification, replicating results from the CCR manuscript (Havrilla et al. 2019). Consistent with a recent survey of pathogenicity predictor performance using ClinVar variants (Ghosh, Oak, and Plon 2017), we find that VEST outperforms FATHMM for ClinVar variants. Missense depletion and badness were consistently the worst performing classification scores. The differing performance of these tools by disease panel demonstrates the utility of constructing PathoPredictor as a disease-specific combination of tools.

To construct PathoPredictor, we introduced a unique variant dataset derived from clinical panel sequencing results for cardiomyopathy, epilepsy, and RASopathy patients. A benefit of this dataset compared to ClinVar is that the variants are labeled and obtained in a more homogeneous way, which helps remove data biases. Furthermore, the variant labels followed clinical interpretation standards similar to the ACMG/AMP guidelines, making the dataset more similar to real world clinical use cases. The clinically classified (pathogenic and benign) variants incorporated several pieces of evidence, mainly segregation, variant effect, functional, and allele frequency data, with limited reliance on computer predictions (or none) thus ensuring that there are no biases towards any one prediction tool. To avoid any further biases in our training and test datasets, we removed all variants previously used to train any of the component features. However, this significantly reduced the number of variants to optimize and evaluate PathoPreditor. Further testing and optimization, with larger clinically curated variant datasets, is required to confirm PathoPredictor’s superior performance, and its utility in a clinical setting. An additional limitation of PathoPredictor, and other pathogenicity scores mentioned here, is that they are trained and evaluated using missense variants, ignoring synonymous variants that may impact splicing.

We demonstrated the utility of PathoPredictor using missense variants from ClinVar and a variant set of *de novo* variants previously used to compare REVEL and CCR. PathoPredictor performs significantly better than its constituent features when evaluated with ClinVar. However, a recent study of ClinVar variants concluded that while ClinVar has improved over time, it contains incorrect pathogenic labels for some endocrine tumor syndrome variant labels (Toledo and NGS in PPGL (NGSnPPGL) Study Group 2018), as an example. While this ClinVar problem could affect our results, we also find that PathoPredictor has a significantly higher average precision than REVEL when testing with the *de novo* variants, which is consistent with CCR’s improvement over REVEL using this same dataset (Havrilla et al. 2019).

In conclusion, we recommend using PathoPredictor scores to predict missense variant pathogenicity for cardiomyopathy, epilepsy, and RASopathies. Predictions for all possible missense variants for disease panel genes are located in Supplemental Table S2.

## Methods

### Classifier

We used Python’s scikit-learn machine learning library to train a logistic regression model to predict the pathogenicity of missense variants from clinical panels. Variants were classified by two well known clinical laboratories, GeneDx (Gaithersburg, MD) and the Laboratory for Molecular Medicine (Harvard Medical School, MA), using variant interpretation protocols that are well within the most recent 2015 ACMG/AMP guidelines. Pathogenic and likely pathogenic variants were assigned values of one, while benign and likely benign variants were assigned values of zero. During training, we used L2 regularization with a regularization strength of one. Our model included six features corresponding to measures of pathogenicity, one Pfam domain indicator, and all pairwise combinations of features. All model terms were standardized by removing the mean and scaling by the standard deviation with scikit-learn.

### Pathogenicity scores as classifier features

We used seven features in our classifier: CCR, FATHMM, missense depletion, missense badness, MTR, VEST, and Pfam protein domains. FATHMM and VEST can provide multiple scores for one variant, depending on isoforms. VEST scores were taken as the minimum VEST v3.0 score provided by dbNSFP v2.9 (Liu, Jian, and Boerwinkle 2013). FATHMM scores were taken as the negative minimum FATHMM v2.3 score provided by dbNSFP v2.9. FATHMM scores were negated so that their interpretation would match the other features. CCR scores were taken as the CCR percentile (ccr_pct) from the CCR BED file v1.20171112 (Havrilla et al. 2019). Missense depletion and badness scores were taken from the constraint MPC VCF file v2 as obs_exp and mis_badness, respectively (Samocha et al. 2017). Missense depletion was negated so that its interpretation would match the other features. The MTR manuscript authors provided chromosome-specific tab delimited text files containing MTR scores and associated metrics for single genomic positions. We extracted MTR scores, negated them so that their interpretation would match the other features, and constructed BED files where each line corresponded to a region of consecutive positions with identical MTR scores. A BED file of Pfam domain locations was downloaded from the UCSC Genome Browser. We assigned each variant a Pfam score of one if its position fell within a domain, and zero otherwise. Note that we omit this simple Pfam domain feature from figures comparing feature classification performance because we do not expect it to perform well by itself. Feature values were assigned to variants as described below in the variant annotation and filtering pipeline.

For each disease dataset we used Python’s scikit-learn library to standardize each feature by removing its mean and scaling by its standard deviation. Disease panel, ClinVar, and *de novo* variants for the same disease were processed together so that their features would be on the same scale.

### Variant sets

We utilized three missense variant sources in this study: disease panels and ClinVar for model training and validation (using unique variant sets), and neurodevelopmental patients for comparing PathPredicor to MPC and REVEL.

GeneDx provided clinical sequencing panel results for epilepsy, while the Laboratory for Molecular Medicine provided their clinically curated data for cardiomyopathy and RASopathies. The number of patients investigated differed by gene. The maximum number of patients observed was 5466 for cardiomyopathy, 8583 for epilepsy, and 3022 for RASopathies. No gene was shared between the three datasets. Variants were provided in Human Genome Variation Society (HGVS) c. notation (Dunnen et al. 2016), and were converted to VCF files of hg19 based variants using Mutalyzer (Wildeman et al. 2008) and custom scripts. To construct variant sets for our classifier, we discarded variants of uncertain significance. We formed a benign set of variants using “benign” and “likely benign” variants. Similarly, our pathogenic variant set consisted of “pathogenic” and “likely pathogenic” variants. These labeled variants were used for training disease-specific classifiers (see below). Diseases panel variants and labels were deposited into Supplemental Table S1.

Variants from ClinVar were chosen as a validation set. We restricted ClinVar genes to those found in the disease panels, and removed any ClinVar genomic position found in the disease panels, producing an independent variant set. The hg19 ClinVar VCF file was downloaded on February 25, 2018, and limited to unambiguously pathogenic or likely pathogenic and benign or likely benign variants with no conflicts according to CLINSIG (Landrum et al. 2016). We considered ClinVar variants with any review status as one test set, and consulted CLNREVSTAT (Landrum et al. 2016) to produce a second ClinVar test variant set restricted to reviewed variants.

As in the CCR paper, we compared PathoPredictor, REVEL, and MPC using *de novo* missense variants from 5,620 neurodevelopmental disorder patients and 2,078 unaffected siblings of autism spectrum disorder patients (Samocha et al. 2017; Havrilla et al. 2019). *De novo* variants from patients were considered pathogenic, and *de novo* variants from unaffected siblings were considered benign. HGVS formatted variants were uploaded to VariantValidator (Freeman et al. 2018), and a VCF file was constructed from the results. This file was normalized with vt (Tan, Abecasis, and Kang 2015). To avoid evaluating with any tool’s training data, we removed disease panel variants, ClinVar variants, and benign variants present in more than 1% of ExAC (MPC’s benign training data).

### Disease-specific classifier evaluation

We compared estimated pathogenicity probabilities produced by our trained models with each pathogenicity score used as a model feature via precision-recall curves and average precision, as implemented in scikit-learn. Precision-recall curves and average precision are useful here due to the possibility of imbalances between pathogenic and benign variant counts. Average precision is an approximation of the area under the method’s precision-recall curve. We ran three experiments with disease panel and ClinVar variants to evaluate the performance of PathoPredictor. First, we used cross validation with disease panel variants. Second, we trained a model with disease panel variants, and validated it with ClinVar variants. Third, we trained a model with ClinVar variants, and validated it with disease panel variants. Comparisons were conducted in a leave one gene out manner. We iterated over all genes in a disease, holding out the gene of interest, and training a model using variants from all remaining genes. This model was applied to variants from the gene of interest, ensuring that a given gene was never used for training and validation. ClinVar variant datasets were restricted to variants not found in the disease panel results. Precision-recall curves and average precision scores were made for each pathogenicity score by aggregating the results from each gene evaluation. The DeLong test as implemented in R’s pROC package (Robin et al. 2012) was used to compare areas under receiver operating characteristics curves produced by predictors. We used this test to gauge the significance of differences between classifiers.

### Comparing PathoPredictor with MPC and REVEL

We applied our epilepsy variant trained PathoPredictor (using all or dominant epilepsy genes) to *de novo* missense variants. For the evaluation of PathoPredictor, MPC, and REVEL, we used 54 pathogenic missense variants located in the epilepsy panel genes. Limiting the benign missense dataset to epilepsy genes produced only 6 benign variants for evaluation, so we randomly selected 48 variants from the full benign missense dataset of 969 variants not located in epilepsy genes so that the pathogenic and benign evaluation sets would have the same size. PathoPredictor, MPC, and REVEL scores were compared using precision-recall curves, average precision, and the DeLong test.

### Variant annotation and filtering pipeline

Our pipeline began with VCF files containing 6,382 *de novo*, 345,849 ClinVar and 7,840 disease panel variants labeled as benign, pathogenic, or VUS. Snpeff v4.3.1T (Cingolani et al. 2012) was used to determine variant effects in GRCh37.75. SnpSift v4.3 (Cingolani et al. 2012) was used to annotate variants with allele frequencies from the ESP, FATHMM scores, and VEST scores from dbNSFP. We annotated variants with values from BED (CCR and Pfam) and VCF (missense badness and depletion and calculated ESP frequencies from ESP6500SI-V2) files using vcfanno v0.2.8 (Pedersen, Layer, and Quinlan 2016). The ESP frequencies are needed next when remove the training data used for VEST.

We next removed variants that had been used to train FATHMM (∼49,500 disease variants and ∼37,000 putatively neutral variants) or VEST (∼45,000 disease variants and ∼45,000 putatively neutral variants), ensuring that these features would not have an advantage when comparing pathogenicity scores, and that our validation datasets would not overlap with any variants used for training. Both FATHMM and VEST were trained with damaging mutations from HGMD, but they differed in their choice of neutral missense variant set. FATHMM was trained with neutral variants from UniProt (The UniProt Consortium 2017) and VEST was trained with missense variants from ESP achieving an population frequency of 1% or higher.

We then removed variants found in the set of 154,257 DM (damaging mutation) in HGMD Professional 2016.1. To address frequent ESP variants, we took the variant frequency as the maximum of dbNSFP fields ESP6500_EA_AF, ESP6500_AA_AF, and the total ESP allele frequency determined using vcfanno. We discarded variants (68 for ClinVar, and 59 disease panel) when this maximum value reached at least 0.01. To remove neutral UniProt variants, we used “Polymorphism” annotations to build a list of neutral codons relative to hg38. Polymorphism annotations were downloaded from www.uniprot.org/docs/humsavar.txt, and joined with hg38 codon coordinates from UniProt. Both were downloaded on April 5th, 2018. We used liftOver (Hinrichs et al. 2006) to convert these to hg19, and removed any variant found in any of 921,722 neutral codons.

Final variant sets were taken as missense variants with CCR and missense depletion, and MTR scores to avoid missing data issues. After applying all the above filters, 666 disease panel, 1,159 ClinVar, and 108 *de novo* missense variants were used in this study.

### Software availability

Source code for this manuscript is available at https://github.com/samesense/pathopredictor, and included as Supplemental Code. A docker image for running PathoPredictor is available at https://hub.docker.com/r/samesense/pathopredictor/. Diseases panel variants and labels were deposited into Supplemental Table S1. Predictions for all possible missense variants for disease panel genes are located in Supplemental Table S2.

## Acknowledgments

We thank all members of GeneDx and the Laboratory for Molecular Medicine (LMM) who manually curated and classified the disease variants used in our study.

