## Supplemental material for "Genetic variant pathogenicity prediction trained using disease-specific clinical sequencing datasets"

**Supplemental Materials**

**Table S1**. Missense disease panel variants. All missense clinical sequencing variants from cardiomyopathy, epilepsy, and rasopathies are included with hg19 coordinates. Each variant has an associated disease and pathogenicity (V for VUS, B for benign, and P for pathogenic).

**Table S2**. Disease panel gene predictions. Every possible missense variant for disease panel genes is listed with predictions from each disease panel classifier. The pathopredictor_class column holds the predicted pathogenicity (1 for pathogenic, and 0 for benign). The pathopredictor_score column holds the prediction score yielded by the model. This score ranges from 0 (most benign) to 1 (most pathogenic).

**Supplemental Code**. Source code for this manuscript is included as a Snakemake workflow and associated scripts. This code is maintained at <https://github.com/samesense/pathopredictor>. A docker image for running PathoPredictor is available at <https://hub.docker.com/r/samesense/pathopredictor/>.
