## Supplementary material for "Genetic variant pathogenicity prediction trained using disease-specific clinical sequencing datasets": Source Code: ahmad_grant_2017.md

### Epilepsy evaluation

Here are the missense variants in genes.
![missense_counts](plots/gene_missense_counts.png)

Here's the MPC distribution.
![mpc_dist](plots/mpc_hist.png)

Here's the leave one out method.

Here's the ROC curve.

![clinvar mpc missense roc](plots/missense_fg_roc.png)

### Clinvar evaluation

Clinvar was downloaded February 2016. It has 10806 variants in 9234 positions. Genes were limited to those in the XXX in the epilepsy panel, and Positions were limited to those where all variants have the same clinical classification. 2350 positions have 2517 pathogenic or likely pathogenic variants, and 1997 positions have 2050 benign or likely benign variants. We focus on missense variants because those are the most difficult to classify. There are 1023 pathogenic missense positions with 1059 variants, and 518 benign missense positions with 523 variants.

934 missense pathogenic positions have 966 variants with MPC scores. MPC scores have not been computed for all transcripts yet, so some genes have no scores. The genes with the most missing clinvar variants are FOXG1 (24), MECP2 (16), and TSC2 (30). 371 benign missense positions have 374 variants with MPC scores. First, we classify these positions using MPC. Next we classify these variants using MPC, MPC+path-fraction, or path-fraction. This demonstrates that a combined MPC+path-fraction works better than either alone. When a variant is outside of a Pfam domain, it is assigned a dummy domain that it flanked by two Pfam domains, or the start/end of a gene and a Pfam domain.

![clinvar mpc missense roc](plots/missense_clinvar_roc_feature_union.png)

Note that clinvar has not been limited by populaiton frequency.
