## Supplementary material for "Genetic variant pathogenicity prediction trained using disease-specific clinical sequencing datasets": Source Code: conf.py

```
# -*- coding: utf-8 -*-
#
# mahdi_epi documentation build configuration file, created by
# sphinx-quickstart.
#
# This file is execfile()d with the current directory set to its containing dir.
#
# Note that not all possible configuration values are present in this
# autogenerated file.
#
# All configuration values have a default; values that are commented out
# serve to show the default.

import os
import sys

# If extensions (or modules to document with autodoc) are in another directory,
# add these directories to sys.path here. If the directory is relative to the
# documentation root, use os.path.abspath to make it absolute, like shown here.
# sys.path.insert(0, os.path.abspath('.'))

# -- General configuration -----------------------------------------------------

# If your documentation needs a minimal Sphinx version, state it here.
# needs_sphinx = '1.0'

# Add any Sphinx extension module names here, as strings. They can be extensions
# coming with Sphinx (named 'sphinx.ext.*') or your custom ones.
extensions = []

# Add any paths that contain templates here, relative to this directory.
templates_path = ['_templates']

# The suffix of source filenames.
source_suffix = '.rst'

# The encoding of source files.
# source_encoding = 'utf-8-sig'

# The master toctree document.
master_doc = 'index'

# General information about the project.
project = u'mahdi_epi'

# The version info for the project you're documenting, acts as replacement for
# |version| and |release|, also used in various other places throughout the
# built documents.
#
# The short X.Y version.
version = '0.1'
# The full version, including alpha/beta/rc tags.
release = '0.1'

# The language for content autogenerated by Sphinx. Refer to documentation
# for a list of supported languages.
# language = None

# There are two options for replacing |today|: either, you set today to some
# non-false value, then it is used:
# today = ''
# Else, today_fmt is used as the format for a strftime call.
# today_fmt = '%B %d, %Y'

# List of patterns, relative to source directory, that match files and
# directories to ignore when looking for source files.
exclude_patterns = ['_build']

# The reST default role (used for this markup: `text`) to use for all documents.
# default_role = None

# If true, '()' will be appended to :func: etc. cross-reference text.
# add_function_parentheses = True

# If true, the current module name will be prepended to all description
# unit titles (such as .. function::).
# add_module_names = True

# If true, sectionauthor and moduleauthor directives will be shown in the
# output. They are ignored by default.
# show_authors = False

# The name of the Pygments (syntax highlighting) style to use.
pygments_style = 'sphinx'

# A list of ignored prefixes for module index sorting.
# modindex_common_prefix = []


# -- Options for HTML output ---------------------------------------------------

# The theme to use for HTML and HTML Help pages.  See the documentation for
# a list of builtin themes.
html_theme = 'default'

# Theme options are theme-specific and customize the look and feel of a theme
# further.  For a list of options available for each theme, see the
# documentation.
# html_theme_options = {}

# Add any paths that contain custom themes here, relative to this directory.
# html_theme_path = []

# The name for this set of Sphinx documents.  If None, it defaults to
# "<project> v<release> documentation".
# html_title = None

# A shorter title for the navigation bar.  Default is the same as html_title.
# html_short_title = None

# The name of an image file (relative to this directory) to place at the top
# of the sidebar.
# html_logo = None

# The name of an image file (within the static path) to use as favicon of the
# docs.  This file should be a Windows icon file (.ico) being 16x16 or 32x32
# pixels large.
# html_favicon = None

# Add any paths that contain custom static files (such as style sheets) here,
# relative to this directory. They are copied after the builtin static files,
# so a file named "default.css" will overwrite the builtin "default.css".
html_static_path = ['_static']

# If not '', a 'Last updated on:' timestamp is inserted at every page bottom,
# using the given strftime format.
# html_last_updated_fmt = '%b %d, %Y'

# If true, SmartyPants will be used to convert quotes and dashes to
# typographically correct entities.
# html_use_smartypants = True

# Custom sidebar templates, maps document names to template names.
# html_sidebars = {}

# Additional templates that should be rendered to pages, maps page names to
# template names.
# html_additional_pages = {}

# If false, no module index is generated.
# html_domain_indices = True

# If false, no index is generated.
# html_use_index = True

# If true, the index is split into individual pages for each letter.
# html_split_index = False

# If true, links to the reST sources are added to the pages.
# html_show_sourcelink = True

# If true, "Created using Sphinx" is shown in the HTML footer. Default is True.
# html_show_sphinx = True

# If true, "(C) Copyright ..." is shown in the HTML footer. Default is True.
# html_show_copyright = True

# If true, an OpenSearch description file will be output, and all pages will
# contain a <link> tag referring to it.  The value of this option must be the
# base URL from which the finished HTML is served.
# html_use_opensearch = ''

# This is the file name suffix for HTML files (e.g. ".xhtml").
# html_file_suffix = None

# Output file base name for HTML help builder.
htmlhelp_basename = 'mahdi_epidoc'


# -- Options for LaTeX output --------------------------------------------------

latex_elements = {
    # The paper size ('letterpaper' or 'a4paper').
    # 'papersize': 'letterpaper',

    # The font size ('10pt', '11pt' or '12pt').
    # 'pointsize': '10pt',

    # Additional stuff for the LaTeX preamble.
    # 'preamble': '',
}

# Grouping the document tree into LaTeX files. List of tuples
# (source start file, target name, title, author, documentclass [howto/manual]).
latex_documents = [
    ('index',
     'mahdi_epi.tex',
     u'mahdi_epi Documentation',
     u"samesense", 'manual'),
]

# The name of an image file (relative to this directory) to place at the top of
# the title page.
# latex_logo = None

# For "manual" documents, if this is true, then toplevel headings are parts,
# not chapters.
# latex_use_parts = False

# If true, show page references after internal links.
# latex_show_pagerefs = False

# If true, show URL addresses after external links.
# latex_show_urls = False

# Documents to append as an appendix to all manuals.
# latex_appendices = []

# If false, no module index is generated.
# latex_domain_indices = True


# -- Options for manual page output --------------------------------------------

# One entry per manual page. List of tuples
# (source start file, name, description, authors, manual section).
man_pages = [
    ('index', 'mahdi_epi', u'mahdi_epi Documentation',
     [u"samesense"], 1)
]

# If true, show URL addresses after external links.
# man_show_urls = False


# -- Options for Texinfo output ------------------------------------------------

# Grouping the document tree into Texinfo files. List of tuples
# (source start file, target name, title, author,
#  dir menu entry, description, category)
texinfo_documents = [
    ('index', 'mahdi_epi', u'mahdi_epi Documentation',
     u"samesense", 'mahdi_epi',
     'Variant effect prediction using domains', 'Miscellaneous'),
]

# Documents to append as an appendix to all manuals.
# texinfo_appendices = []

# If false, no module index is generated.
# texinfo_domain_indices = True

# How to display URL addresses: 'footnote', 'no', or 'inline'.
# texinfo_show_urls = 'footnote'
```
