## Supplementary material for "Genetic variant pathogenicity prediction trained using disease-specific clinical sequencing datasets": Source Code: how_to_predict.md

Predicting w/ PathoPredictor
==============================

Use docker container [samesense/pathopredictor](https://hub.docker.com/r/samesense/pathopredictor/) to make predictions. The container includes all software, but annotation datasets need to be downloaded. You need to specify a data directory mount when using docker. Your input and output files will live here.

```
docker run -e USER=$USER -e USERID=$UI --user $(id -u) -it -v /absolute/local/data/:/opt/pathopredictor/data/ samesense/pathopredictor /bin/bash
```

Put your vcf file(s) in `/absolute/local/data/interim/user_preds/`. Using the `interim/user_preds/` directory is required, but only use the first part of the path to data when envoking docker, like this `-v /absolute/local/data/`.

Edit `/opt/pathopredictor/configs/sm_predict_ex_config.json` so that it contains the names (w/ no vcf suffix) of your vcf files.

While in the docker container, navigate to `/opt/pathopredictor/src/rules/`. The prediction pipeline requires 32G of ram. The prediction pipeline is run with Snakemake, after updating the container's path:

```
export PATH=/opt/conda/envs/pathopredictor/bin/:$PATH
/opt/conda/envs/pathopredictor/bin/snakemake --configfile ../../configs/sm_predict_ex_config.json -s sf_predict.py all_predictions
```

To see the pipeline rules that will be run:

```
/opt/conda/envs/pathopredictor/bin/snakemake --configfile ../../configs/sm_predict_ex_config.json -s sf_predict.py -n all_predictions
```
