## Supplementary material for "Genetic variant pathogenicity prediction trained using disease-specific clinical sequencing datasets": Source Code: ISMB18.md

The abstract should be a brief and concise summary of the background/motivation for the study,
method and result(s), as well as conclusion/statement of significance.
The additional 1-page pdf can be used to provide additional details,
as well as figures and graphics that support the main points of the abstract.
