## Supplementary material for "Genetic variant pathogenicity prediction trained using disease-specific clinical sequencing datasets": Source Code: goals.md

### 110418
* standardize features

### 100418
* roc evaluation for panel and clinvar - DONE. Still looks good.
* write predictor methods
* standardize or reverse negative scores so fathmm does not do bad
