## Supplementary material for "Genetic variant pathogenicity prediction trained using disease-specific clinical sequencing datasets": Source Code: log.md

### 2019_04_09

#### variant count stats
```
$ cd /mnt/isilon/dbhi_bfx/perry/projects/sarmadi/mahdi_epi/data/interim/full
# subtract one from each count for header
$ cat ndenovo.dat | cut -f 1,2,3,5 | sc | wc -l
$ cat clinvar.dat | cut -f 1,2,3,4 | sc | wc -l
$ cat panel.dat | cut -f 7,22,23,5 | sc | wc -l
```

### results: within panel
```
# pval auc cmp
$ cd /mnt/isilon/dbhi_bfx/perry/projects/sarmadi/mahdi_epi/data/interim/auc_cmp
$ cat clinvar.panel.ccr-vest-fathmm-missense_badness-missense_depletion-mtr-is_domain | sort -k3gr
# worst avg pre
$ cd /mnt/isilon/dbhi_bfx/perry/projects/sarmadi/mahdi_epi/data/interim/pred_panel_eval
$  cat ccr-vest-fathmm-missense_badness-missense_depletion-mtr-is_domain | grep Path | cut -f 2
```

### results: eval clinvar
```
# pvals
$ cd /mnt/isilon/dbhi_bfx/perry/projects/sarmadi/mahdi_epi/data/interim/auc_cmp
$ sort -k3gr clinvar.clinvar.ccr-vest-fathmm-missense_badness-missense_depletion-mtr-is_domain
# avg pre
$ cd /mnt/isilon/dbhi_bfx/perry/projects/sarmadi/mahdi_epi/data/interim/EVAL_clinvar/pred_clinvar_eval
$ grep Path clinvar_*.ccr-vest-fathmm-missense_badness-missense_depletion-mtr-is_domain | sort -k2gr
```

#### results: clinvar train; test panel
```
$ cd /mnt/isilon/dbhi_bfx/perry/projects/sarmadi/mahdi_epi/data/interim/train_clinvar_test_panel_singleTrue/EVAL/pred_panel_eval
$ grep Path ccr-vest-fathmm-missense_badness-missense_depletion-mtr-is_domain 
$ cd /mnt/isilon/dbhi_bfx/perry/projects/sarmadi/mahdi_epi/data/interim/train_clinvar_test_panel_singleFalse/EVAL/pred_panel_eval
# same grep
```

#### fig5 clinvar train; test panel
```
$ cd /mnt/isilon/dbhi_bfx/perry/projects/sarmadi/mahdi_epi/data/interim/auc_cmp_train_clinvar_test_panel
# some evidence in clinvar; sort by pval
$ sort -k3gr singleTrue.ccr-vest-fathmm-missense_badness-missense_depletion-mtr-is_domain
$ sort -k3gr singleFalse.ccr-vest-fathmm-missense_badness-missense_depletion-mtr-is_domain
```

#### fig6a single gene eval and results
```
$ cd /mnt/isilon/dbhi_bfx/perry/projects/sarmadi/mahdi_epi/data/interim/single_gene_stats
# KCNQ2 accuracy for within panel and clinvar
$ grep KCNQ2 panel.ccr-vest-fathmm-missense_badness-missense_depletion-mtr-is_domain
$ grep KCNQ2 clinvar.ccr-vest-fathmm-missense_badness-missense_depletion-mtr-is_domain
```

#### fig6c pp vs mpc/revel
```
$ cd /mnt/isilon/dbhi_bfx/perry/projects/sarmadi/mahdi_epi/data/interim/auc_cmp
$ grep mpc ndenovo.clinvar.ccr-vest-fathmm-missense_badness-missense_depletion-mtr-is_domain
$ grep revel ndenovo.clinvar.ccr-vest-fathmm-missense_badness-missense_depletion-mtr-is_domain
```

### revel vs pp results
```
$ cd /mnt/isilon/dbhi_bfx/perry/projects/sarmadi/mahdi_epi/data/interim/EVAL_ndenovo/pred_clinvar_eval
$ grep Path clinvar_tot.ccr-vest-fathmm-missense_badness-missense_depletion-mtr-is_domain
```

### 2019_04_08
* source activate PP4
* sm -s sf.py -j60 --use-singularity --singularity-args "-B /mnt/isilon/:/mnt/isilon" upload_all

### 2018_11_14
* dropped training vars that were at same position, but w/ diff alt alleles

### 2018_11_09
* duplicate clinvars. 115256528

### 2018_10_26
* update methods for mtr
* mtr score parsing was fixed. It performs well now.

### 2018_10_12
* ndenovo test w/ all denovo shows epi limited is a bit better than mpc/revel
* use path from epi genes w/ all ndenovo looks good for pp
* sample = number of ndenovo benign is better
* pvals do not completely reflect this

### 2018_10_10
* ndenovo has few benign

### 2018_10_08
* add mtr to feature ls
* add mpc to final eval

## 2018_06_21
* Rejected from plos machine learning edition
* Trying genome research. Gotta mk png, and smaller images to imbed in gdocs. Pngs done.
* fix ref format
* make cover letter match reqs

### 2018_06_01
* Finish prediction pipeline
* where to put vcfanno data?

### 2018_05_30
* submitted

### 2018_05_29
* If I add all panel genes for epi, how many more? 68 epi genes initially. 52 for panel evaluation
* I've updated all the figures. Next update paper numbers and results. Yes! Automate numbers.

### 2018_05_23
* finish count table
* test auc. I already have it. It looks good! use proc test.roc
* cmp revel. wtf _merge?

### 2018_05_22
* fix top genes
* count vars
* eval revel denovo

### 2018_05_21
* cmp revel. revel is better
* pvals for scores
* top gene pr curves
* var counts as pipeline moves

### 2018_05_18
* single gene fig, method, result
* background
* discussion
* fig1 and fig importance
* I don't see how clinvar single ran w/ the right data. need to account for clinvar set in main prediction data
* I can't seem to parse gnomad this morining.
* Clinvar single vs total was not a problem, but the script should have been clearer. It's under control now.
* No training data after removing vars. Try not using vest cutoff. Fathmm too. That's a lot of work. There are some left after the vest and fathmm filters.
* Also must redo clinvar restrictions for these guys. Do this only if vest and fathmm are removed.
* only epi has enough training vars
* gene specific stuff looks too bad to publish, so it's not going in

### 2018_05_17
* Mehods done except single gene
* single gene: single_gene_collapse_.003_.1.pdf. Need to filter gnomad first. Running. parse_gnomad like clinvar and panel. Don't forget to filter vars after w/ limit_eval_general. remove panel vars from extra gnomad vars, and remove extra gnomad vars from clinvar test
* what about the count figure? How is it made if global eval limits my genes? I should update fig1. It uses results, so it should be fine.

### 2018_05_10
* First tackle background
* Then methods
