## Supplementary material for "Genetic variant pathogenicity prediction trained using disease-specific clinical sequencing datasets": Source Code: make.bat

```
@ECHO OFF

REM Command file for Sphinx documentation

if "%SPHINXBUILD%" == "" (
	set SPHINXBUILD=sphinx-build
)
set BUILDDIR=_build
set ALLSPHINXOPTS=-d %BUILDDIR%/doctrees %SPHINXOPTS% .
set I18NSPHINXOPTS=%SPHINXOPTS% .
if NOT "%PAPER%" == "" (
	set ALLSPHINXOPTS=-D latex_paper_size=%PAPER% %ALLSPHINXOPTS%
	set I18NSPHINXOPTS=-D latex_paper_size=%PAPER% %I18NSPHINXOPTS%
)

if "%1" == "" goto help

if "%1" == "help" (
	:help
	echo.Please use `make ^<target^>` where ^<target^> is one of
	echo.  html       to make standalone HTML files
	echo.  dirhtml    to make HTML files named index.html in directories
	echo.  singlehtml to make a single large HTML file
	echo.  pickle     to make pickle files
	echo.  json       to make JSON files
	echo.  htmlhelp   to make HTML files and a HTML help project
	echo.  qthelp     to make HTML files and a qthelp project
	echo.  devhelp    to make HTML files and a Devhelp project
	echo.  epub       to make an epub
	echo.  latex      to make LaTeX files, you can set PAPER=a4 or PAPER=letter
	echo.  text       to make text files
	echo.  man        to make manual pages
	echo.  texinfo    to make Texinfo files
	echo.  gettext    to make PO message catalogs
	echo.  changes    to make an overview over all changed/added/deprecated items
	echo.  linkcheck  to check all external links for integrity
	echo.  doctest    to run all doctests embedded in the documentation if enabled
	goto end
)

if "%1" == "clean" (
	for /d %%i in (%BUILDDIR%\*) do rmdir /q /s %%i
	del /q /s %BUILDDIR%\*
	goto end
)

if "%1" == "html" (
	%SPHINXBUILD% -b html %ALLSPHINXOPTS% %BUILDDIR%/html
	if errorlevel 1 exit /b 1
	echo.
	echo.Build finished. The HTML pages are in %BUILDDIR%/html.
	goto end
)

if "%1" == "dirhtml" (
	%SPHINXBUILD% -b dirhtml %ALLSPHINXOPTS% %BUILDDIR%/dirhtml
	if errorlevel 1 exit /b 1
	echo.
	echo.Build finished. The HTML pages are in %BUILDDIR%/dirhtml.
	goto end
)

if "%1" == "singlehtml" (
	%SPHINXBUILD% -b singlehtml %ALLSPHINXOPTS% %BUILDDIR%/singlehtml
	if errorlevel 1 exit /b 1
	echo.
	echo.Build finished. The HTML pages are in %BUILDDIR%/singlehtml.
	goto end
)

if "%1" == "pickle" (
	%SPHINXBUILD% -b pickle %ALLSPHINXOPTS% %BUILDDIR%/pickle
	if errorlevel 1 exit /b 1
	echo.
	echo.Build finished; now you can process the pickle files.
	goto end
)

if "%1" == "json" (
	%SPHINXBUILD% -b json %ALLSPHINXOPTS% %BUILDDIR%/json
	if errorlevel 1 exit /b 1
	echo.
	echo.Build finished; now you can process the JSON files.
	goto end
)

if "%1" == "htmlhelp" (
	%SPHINXBUILD% -b htmlhelp %ALLSPHINXOPTS% %BUILDDIR%/htmlhelp
	if errorlevel 1 exit /b 1
	echo.
	echo.Build finished; now you can run HTML Help Workshop with the ^
.hhp project file in %BUILDDIR%/htmlhelp.
	goto end
)

if "%1" == "qthelp" (
	%SPHINXBUILD% -b qthelp %ALLSPHINXOPTS% %BUILDDIR%/qthelp
	if errorlevel 1 exit /b 1
	echo.
	echo.Build finished; now you can run "qcollectiongenerator" with the ^
.qhcp project file in %BUILDDIR%/qthelp, like this:
	echo.^> qcollectiongenerator %BUILDDIR%\qthelp\mahdi_epi.qhcp
	echo.To view the help file:
	echo.^> assistant -collectionFile %BUILDDIR%\qthelp\mahdi_epi.ghc
	goto end
)

if "%1" == "devhelp" (
	%SPHINXBUILD% -b devhelp %ALLSPHINXOPTS% %BUILDDIR%/devhelp
	if errorlevel 1 exit /b 1
	echo.
	echo.Build finished.
	goto end
)

if "%1" == "epub" (
	%SPHINXBUILD% -b epub %ALLSPHINXOPTS% %BUILDDIR%/epub
	if errorlevel 1 exit /b 1
	echo.
	echo.Build finished. The epub file is in %BUILDDIR%/epub.
	goto end
)

if "%1" == "latex" (
	%SPHINXBUILD% -b latex %ALLSPHINXOPTS% %BUILDDIR%/latex
	if errorlevel 1 exit /b 1
	echo.
	echo.Build finished; the LaTeX files are in %BUILDDIR%/latex.
	goto end
)

if "%1" == "text" (
	%SPHINXBUILD% -b text %ALLSPHINXOPTS% %BUILDDIR%/text
	if errorlevel 1 exit /b 1
	echo.
	echo.Build finished. The text files are in %BUILDDIR%/text.
	goto end
)

if "%1" == "man" (
	%SPHINXBUILD% -b man %ALLSPHINXOPTS% %BUILDDIR%/man
	if errorlevel 1 exit /b 1
	echo.
	echo.Build finished. The manual pages are in %BUILDDIR%/man.
	goto end
)

if "%1" == "texinfo" (
	%SPHINXBUILD% -b texinfo %ALLSPHINXOPTS% %BUILDDIR%/texinfo
	if errorlevel 1 exit /b 1
	echo.
	echo.Build finished. The Texinfo files are in %BUILDDIR%/texinfo.
	goto end
)

if "%1" == "gettext" (
	%SPHINXBUILD% -b gettext %I18NSPHINXOPTS% %BUILDDIR%/locale
	if errorlevel 1 exit /b 1
	echo.
	echo.Build finished. The message catalogs are in %BUILDDIR%/locale.
	goto end
)

if "%1" == "changes" (
	%SPHINXBUILD% -b changes %ALLSPHINXOPTS% %BUILDDIR%/changes
	if errorlevel 1 exit /b 1
	echo.
	echo.The overview file is in %BUILDDIR%/changes.
	goto end
)

if "%1" == "linkcheck" (
	%SPHINXBUILD% -b linkcheck %ALLSPHINXOPTS% %BUILDDIR%/linkcheck
	if errorlevel 1 exit /b 1
	echo.
	echo.Link check complete; look for any errors in the above output ^
or in %BUILDDIR%/linkcheck/output.txt.
	goto end
)

if "%1" == "doctest" (
	%SPHINXBUILD% -b doctest %ALLSPHINXOPTS% %BUILDDIR%/doctest
	if errorlevel 1 exit /b 1
	echo.
	echo.Testing of doctests in the sources finished, look at the ^
results in %BUILDDIR%/doctest/output.txt.
	goto end
)

:end
```
