## Supplementary material for "Genetic variant pathogenicity prediction trained using disease-specific clinical sequencing datasets": Source Code: meeting_050917.md

### 050917

### Updates
* Performance of just pathogenic fraction
* Clinvar performance

### Pathogenic fraction performance
![roc](plots/roc.png)

### Clinvar performance
* 806 clinvar variants that whose positions are always benign/pathogenic
    * 652 pathogenic
    * 154 benign
* Say pos 1:1234 A>T is path, and A>C is benign. These variants are ignored for now
* AUC evaluation
    * mpc auc 0.854802406183
    * tree auc 0.876005895945
    * tree-no-mpc auc 0.841307465541
![clinvar roc](plots/clinvar_roc.png)

### Plans
* HGMD pathogenic vs clinvar benign?
* Add mtr score evaluation
* Remove clinvar and hgmd variants from training data
* Evaluate mpc on same subset for path/benign labeled data
