## Supplementary material for "Genetic variant pathogenicity prediction trained using disease-specific clinical sequencing datasets": Source Code: meeting_080817.md

## 080817

### Methods
* use mutalyzer to convert c. to genome coords (had trouble w/ indels, the hg19 coords in xlsx are wrong)
* mk vcf file
* annotate with snpeff, dbnsfp
* annotate 1kg, exac_coverage, evs, kaviar, exac, pfam, hgmd, clinvar, cadd, etc with vxfanno
* ready for gemini db
* have exac in gemini db

### variant positions within pfam domains
* 13 BENIGN
* 180 LIKLEY_PATHOGENIC
* 205 LIKELY_BENIGN
* 272 PATHOGENIC
* 1031 VUS

### variant positions with no pfam domain
* 31 BENIGN
* 88 LIKLEY_PATHOGENIC
* 193 PATHOGENIC
* 394 LIKELY_BENIGN
* 1491 VUS

### pfam domains with more than one variant position
* 526 pfam domains hit by at least one variant
* 127 pfam domains with one variant (cannot be evaluated with neighbor variants)

### dist of var position counts in pfam domains
* ex: one domain with 22 variants positions
* 1 22
* 1 30
* 1 53
* 2 13
* 2 14
* 2 15
* 2 16
* 2 18
* 7 12
* 8 10
* 14 7
* 17 8
* 18 9
* 33 5
* 33 6
* 66 4
* 77 3
* 113 2
* 125 1

### dist of pahogenic var position counts in pfam domains
* ex: one domain with 22 pathogenic variant positions
* 1 12 (Ion_trans:482, SCN2A)
* 1 22 (Ion_trans:491, SCN1A)
* 2 10 (KCNQ_channel:11, KCNQ2)
* 2 8
* 2 9
* 5 6
* 8 5
* 13 4
* 24 3
* 28 2
* 94 1

### pfam domains with the most hits (all clinical classifications)
* 16 Ion_trans:553
* 16 Neur_chan_memb:3
* 18 KCNQ_channel:11
* 18 Neur_chan_memb:129
* 22 IRK:0
* 30 Ion_trans:491
* 53 Neur_chan_memb:71

### var types inside pfam domains
* 4 splice_region_variant
* 54 stop_gained
* 280 synonymous_variant
* 1363 missense_variant

### plan
* get read genome coords?
* fix mutalyzer indels
* compare synon vars to exac to find correction factor
* limit to rare vars? and find domains with significant burden when compared to exac
* use burden domain feature + other features to predict benign/likely benign vs pathogenic
