## Supplementary material for "Genetic variant pathogenicity prediction trained using disease-specific clinical sequencing datasets": Source Code: meeting_081517.md

## 081517

### Methods
* use mutalyzer to convert c. to genome coords (had trouble w/ indels)
* mk vcf file
* annotate with snpeff, dbnsfp
* annotate 1kg, exac_coverage, evs, kaviar, exac, pfam, hgmd, clinvar, cadd, etc with vcfanno
* limit to rare vars: less than 1% in 1kg total populations
* test for domain enrichment for missense and nonsense
    * gather all rare positions of interest for disease and exac
    * count the number of variant alleles for these positions
    * when a position is not present, assume it is wt for the max number of alleles observed at a position
    * use fisher's exact test for each domain (include no-domain for now)
    * use bh to correct p-values

### Correlation: exac vs disease
* [silent variants have no correlation](http://franklin.research.chop.edu:8102/notebooks/epi_linked/notebooks/merge_count_data.ipynb)
* [missense variants have a cor of 0.37](http://franklin.research.chop.edu:8102/notebooks/epi_linked/notebooks/merge_count_data_missense.ipynb)

### Nonsense pfam enrichment (qval < .05)
* TSC1	Hamartin
* PCDH19	Cadherin
* KCNQ2	Ion_trans, Ion_trans_2
* CDKL5	Pkinase_Tyr, Pkinase
* SCN1A	Ion_trans

### Missense pfam enrichment (qval < .01)
* KCNQ2	Ion_trans_2, Ion_trans, KCNQ_channel 
* CHRNB2	Neur_chan_memb
* STXBP1	Sec1
* SCN1A	Ion_trans, Sugar_tf, MFS_1
* GABRG2	Neur_chan_memb
* MECP2	MBD
* SCN8A	Ion_trans
* ALDH7A1	Aldedh
* SPTAN1	Spectrin
* SCN1B	V-set, ig
* CDKL5	Pkinase
* GABRA1	Neur_chan_memb
* CNTNAP2	Laminin_G_2
* CTSD	Asp
* FOXG1	Fork_head
* SLC2A1	Sugar_tr, MFS_1
* SCARB2	CD36

### plan
* fix mutalyzer indels
* include frameshift burden
    * lof
    * missese + inframe
* are enriched doamains more likely to have pathogenic vars?    
* use burden domain feature + other features to predict benign/likely benign vs pathogenic
