## Supplementary material for "Genetic variant pathogenicity prediction trained using disease-specific clinical sequencing datasets": Source Code: meeting_082217.md

### 082217

### Updates
* Fixed mutalyzer indels except for complex indels (ins+del)
* burden w/ pathogenic burden evaluation:
    * enrichmed: qval < .01
    * not enriched: qval > .2
    * ignored 'none' domains
    * lof, missense + inframe, total
    * lof is not enriched in pathogenic. Are there some gene/domains I can spot check? relaxing qval and domain does not help either missense nor lof
* can't dl mtr scores, will use missense-badness instead

### plan
* annotate w/ missense-badness
* try to predict using domain status and missese badness vs mis-sense badness alone
* why does lof not work? why so many vars in these few lof domains? B/c it is none? Yes, perhaps. Can ignore this.
* annotate exac freqs for init data
* email for mtr scores
* use vus as denominator as well
* make [paper plot](https://mail.google.com/mail/u/0/#inbox/15e0ad768f930135) too
* counts of sig domains for each type
* count of vars in those domains
