## Supplementary material for "Genetic variant pathogenicity prediction trained using disease-specific clinical sequencing datasets": Source Code: meeting_082917.md

### 290817

### Updates
* annotated mpc scores. [Disease enriched domains have higher MPC scores](plots/mpc.by_pfam_enrichment.png)
* [scored pathogenic/benign w/ MPC](http://franklin.research.chop.edu:8102/notebooks/epi_linked/notebooks/predict-mpc.ipynb#)

### [No-enrichment / leave one out cross validation roc](plots/roc.png)
* ~100 domains have at least 2 benign/pathogenic vars
* Hold out one labeled var from each domain (ignore none domain)
* 91 vars for testing (limited to vars w/ mpc>0 and in some domain w/ at least 2 labeled vars)
* [Small decision tree based on missense constraint and pathogenic domain fraction](plots/mtr_tree.x.pdf)
* [Mostly using pathogenic data](plots/eval_data.png)
* Beat missense constriant scores    

### Evaluate pathogenic burden enrichment with respect to MPC scores
* [Pathogenic fraction (path+benign) for MPC>0](plots/rare.path_frac_wo_vus.pfam.mpcLow_0.mpcHigh_100.png)
    * 90% of variants in disease enriched regions are pathogenic (good)
    * 60% of variants in exac enriched regions are benign (good)
    * 40% of variatns in non-enriched regions are benign (bad)
* [Pathogenic fraction (path+bening) for high MPC>1.4](plots/rare.path_frac_wo_vus.pfam.mpcLow_1.4.mpcHigh_100.png)
    * For burden domains to improve MPC, exac enriched, or non-enriched, domains need to rule out high MPC variants that are not pathogenic.
    * Exac/non-enriched domains need to be enriched in benign variants for migh MPC, but
    * 15% of variants in exac regions are benign for high MPC, so these regions cannot be used to rule out migh MPC that should be benign
* [Pathogenic fraction (path+bening) for low MPC<1.4 variants](plots/rare.path_frac_wo_vus.pfam.mpcLow_0.mpcHigh_1.4.png)
    * For burden domains to iproved MPC, disease-enriched domains need to recover low MPC variatns that should be pathogenic.
    * Disease enriched domains need to be mostly pathogenic for low MPC variants
    * ~40% of enriched domains are pathogenic for low MPC, so these cannot be used to recover low MPC that should be pathogenic
* Conclusion
    * Need to train domains on pathogenic/benign
    * Or come up w/ better enriched domains

### Plans
* Need other dataset
* Examine problematic domains?
* eval clinvar and hgmd
* show fraction performance alone and its distribution
