## Supplementary material for "Genetic variant pathogenicity prediction trained using disease-specific clinical sequencing datasets": Source Code: meeting_102417.md

### 241017

### Updates
* Domain pathogenic freq: sum of path variant frequencies
* Entropy
* Avg freq
* Domain benign freq
* Random forest classifier for feature importance

### Feature importance
* path_frac_t, 1. feature 3 (0.447124)
* benign_ent, 2. feature 10 (0.149873)
* path_ent, 3. feature 7 (0.118855)
* mpc, 4. feature 0 (0.099523)
* benign_freq, 5. feature 8 (0.034581)
* in_none_pfam, 6. feature 4 (0.031161)
* path_avg, 7. feature 6 (0.028193)
* path_benign_freq_r, 8. feature 13 (0.025850)
* benign_avg, 9. feature 9 (0.024398)
* path_freq, 10. feature 5 (0.019332)
* size_t, 11. feature 1 (0.010453)
* af_1kg_all, 12. feature 11 (0.007878)
* mtr, 13. feature 12 (0.002779)
* path_na_t, 14. feature 2 (0.000000)

### Clinvar select gene evaluation
* 'SCN1A','SCN2A','KCNQ2', 'KCNQ3', 'CDKL5', 'PCDH19', 'SCN1B', 'SCN8A', 'SLC2A1', 'SPTAN1', 'STXBP1', 'TSC1'
* http://franklin.research.chop.edu:8101/notebooks/epi_linked/notebooks/predict-for-missense-clinvar-union_features-regression-freq-rf.ipynb
