## Supplementary material for "Genetic variant pathogenicity prediction trained using disease-specific clinical sequencing datasets": Source Code: meeting_260917.md

## 260917

### Updates
* Annotated new data. Very hard b/c of some very old c. missing from VEP and mutalyzer. Made new conversion tool.
* Show mpc hist
![mpc hist](plots/mpc_hist_paper.png)
* Show new roc with very good mpc [notebook](http://franklin.research.chop.edu:8102/notebooks/epi_linked/notebooks/predict-for-missense-lab2-union_features.ipynb)
* panel 1: 70 genes
* panel 2: 45 genes (path dom/rec) (21 common)
    * 102 missense variants
    * 37 path
    * 65 benign
* why not the same genes on panels?
