## Supplementary material for "Genetic variant pathogenicity prediction trained using disease-specific clinical sequencing datasets": Source Code: notes.md

missense badness scores: ftp://ftp.broadinstitute.org/pub/ExAC_release/release1/regional_missense_constraint/README_fordist_mpc_values

plot varinat counts by gene
```
ggplot(data=d) + geom_bar(aes(y=size,x=gene), stat="identity") + facet_grid(.~y) + coord_flip() + theme_bw() + xlab('') + ylab('Variant count')
``````
