## Supplementary material for "Genetic variant pathogenicity prediction trained using disease-specific clinical sequencing datasets": Source Code: results.md

## Results

### Clinvar counts
* [total missense clinvar](https://github.com/samesense/mahdi_epi/blob/master/notebooks/clinvar-report.ipynb)

### [Predict panel variant status](results_predict_panel.md)

### [Predict clinvar variant status](results_predict_clinvar.md)
