## Supplementary material for "Genetic variant pathogenicity prediction trained using disease-specific clinical sequencing datasets": Source Code: results_predict_clinvar.md

### Goal
We are training one MPC cutoff using labeled missense variants from clinical panels, and then predicting that status of missense variants from ClinVar and Denovo-db. Panel training is compared to a hold one gene out training strategy. ClinVar variants have been split by annotation confidence: expert, multiple evidence, single evidence, and all. Each level is determined in a this or better manner. Denovo-db has no benign variants, so it cannot be used for training.

### Panels
    * Cardiomyopathy
    * Noonan
    * UC epilepsy
    * GeneDx epilepsy
    * Focused epilepsy gene panel:
        * SCN1A, SCN2A, KCNQ2, KCNQ3, CDKL5,
        * PCDH19, SCN1B, SCN8A, SLC2A1,
        * SPTAN1, STXBP1, TSC
    * Combined epilepsy
    * Combined focused epilepsy
    * All combined
    
### Questions
* Does training with a panel from above do better than MPC>=2 when testing on all of ClinVar? Yes, all panels are better.
* Does leave one out clinvar do better than the panels? Yes, much better.
* Does training with a panel from above do better than MPC>=2 when testing on all of denovo-db? Yes.
* Does the prediction of ClinVar hold-one-out improve relative to MPC>2 when increasing the quality of ClinVar annotations? Yes, but only for the expert annotations.
* Does the prediction of ClinVar w/ panel training improve relative to MPC>2 when increasing the quality of ClinVar annotations? No, it seems that performance gets worse when increasing ClinVar quality.
* Does training with a panel form above for single gene model do better than a glboal model for Clinvar?
* Does training with a panel form above for single gene model do better than a glboal model for denovo-db?
