## Supplementary material for "Genetic variant pathogenicity prediction trained using disease-specific clinical sequencing datasets": Source Code: results_predict_panel.md

### Predict panel variant status

#### Panels
* Cardiomyopathy
* Noonan
* UC epilepsy
* GeneDx epilepsy
* Focused epilepsy gene panel:
    * SCN1A, SCN2A, KCNQ2, KCNQ3, CDKL5,
    * PCDH19, SCN1B, SCN8A, SLC2A1,
    * SPTAN1, STXBP1, TSC

#### Questions
* Does training w/ clinvar do better than MPC>=2?
* Does training with the panel do better than MPC>=2?
* Does training with panel genes from clinvar do better than all of clinvar? (I'm still evaluating all of the panel, not just panl genes in clinvar. This could be why the results are not as good as total clinvar.)
* Does training with the panel do better than training with limited clinvar (only genes on panel)?
* Do gene specific models do better than global models?

#### Global cutoff methods
* MPC>2 is pathogenic (baseline MPC cutoff)
* hold out test gene from panel and train
* train using all of clinvar minus testing variants
* train using clinvar panel genes minus testing variants

#### Single gene cutoff
* When genes have at least 5 pathogenic and 5 benign variants, evaluate the gene
* Use all methods from the global evaluation, but evaluation is limited by above statement
* Train using clinvar variants from this gene

### Results

#### Does training w/ clinvar do better than MPC>=2?
For a globally trained MPC cutoff, Clinvar is better for cardiomyopathy, geneDx, and noonan. Clinvar is worse for UC epilsepsy. For a single gene model, clinvar is better for GeneDx, Noonan, and limited UC.

#### Does training with the panel do better than MPC>=2?
For the global model, the panel is always better, except for total UC epilepsy.

#### Does training with panel genes from clinvar do better than all of clinvar?
Yes, except for Noonan.

#### Does training with the panel do better than training with limited clinvar (only genes on panel)?
It is better for all but total GeneDx and Noonan.

#### Do gene specific clinvar models do better than global panel model?
No, except for Noonan and total GeneDx.

#### Do gene specific clinvar models do better than global clinvar?
Yes, except for cardiomyopathy.

#### Does increasing ClinVar annotation quality yield better results?
Up to a the single evidence limit. Multiple evidence and expert seems to restricted.

#### Is Denovo-db training better than MPC>2?
Mostly no.

#### Conclusion
* The baseline MPC>2 can be beaten with training
* Training with the disease panel is generally better than using clinvar
* Gene specific models based on clinvar are not generally better than using the total panel - will investigate single gene models from epi data
