## Supplementary material for "Genetic variant pathogenicity prediction trained using disease-specific clinical sequencing datasets": Source Code: todo.md

## next
* format indels (frameshifts)
* tally pfam eff types
* find most path/benign domains
* see if domains help w/ mpc predictions
* add mtr

### fix fg indels

### include frameshift burden
* lof
* missese + inframe

### are enriched doamains more likely to have pathogenic vars?    

### use burden domain feature + other features to predict benign/likely benign vs pathogenic

### incorporate mtr
* http://mtr-viewer.mdhs.unimelb.edu.au

### find lof fields that give best results. current fields no diff from exac
