## Supplementary material for "Genetic variant pathogenicity prediction trained using disease-specific clinical sequencing datasets": Source Code: requirements.yaml

```
name: mahdi_epi
channels:
- defaults
dependencies:
- appnope=0.1.0=py36_0
- bleach=1.5.0=py36_0
- decorator=4.1.2=py36_0
- entrypoints=0.2.3=py36_0
- html5lib=0.999=py36_0
- icu=54.1=0
- ipykernel=4.6.1=py36_0
- ipython=6.1.0=py36_0
- ipython_genutils=0.2.0=py36_0
- ipywidgets=6.0.0=py36_0
- jedi=0.10.2=py36_2
- jinja2=2.9.6=py36_0
- jsonschema=2.6.0=py36_0
- jupyter=1.0.0=py36_3
- jupyter_client=5.1.0=py36_0
- jupyter_console=5.1.0=py36_0
- jupyter_core=4.3.0=py36_0
- markupsafe=0.23=py36_2
- mistune=0.7.4=py36_0
- mkl=2017.0.3=0
- nbconvert=5.2.1=py36_0
- nbformat=4.3.0=py36_0
- notebook=5.0.0=py36_0
- numpy=1.13.1=py36_0
- openssl=1.0.2l=0
- pandas=0.20.3=py36_0
- pandocfilters=1.4.1=py36_0
- path.py=10.3.1=py36_0
- pexpect=4.2.1=py36_0
- pickleshare=0.7.4=py36_0
- pip=9.0.1=py36_1
- prompt_toolkit=1.0.14=py36_0
- ptyprocess=0.5.2=py36_0
- pygments=2.2.0=py36_0
- pyqt=5.6.0=py36_2
- python=3.6.2=0
- python-dateutil=2.6.1=py36_0
- pytz=2017.2=py36_0
- pyzmq=16.0.2=py36_0
- qt=5.6.2=2
- qtconsole=4.3.0=py36_0
- readline=6.2=2
- setuptools=27.2.0=py36_0
- simplegeneric=0.8.1=py36_1
- sip=4.18=py36_0
- six=1.10.0=py36_0
- sqlite=3.13.0=0
- terminado=0.6=py36_0
- testpath=0.3.1=py36_0
- tk=8.5.18=0
- tornado=4.5.1=py36_0
- traitlets=4.3.2=py36_0
- wcwidth=0.1.7=py36_0
- wheel=0.29.0=py36_0
- widgetsnbextension=2.0.0=py36_0
- xlrd=1.0.0=py36_0
- xz=5.2.2=1
- zlib=1.2.8=3
- pip:
  - ipython-genutils==0.2.0
  - jupyter-client==5.1.0
  - jupyter-console==5.1.0
  - jupyter-core==4.3.0
  - prompt-toolkit==1.0.14
prefix: /Users/perry/miniconda3/envs/mahdi_epi
```
