## Supplementary figures and images for "Genetic variant pathogenicity prediction trained using disease-specific clinical sequencing datasets"

### Cardiomyopathy_cv_mpc_eval.png

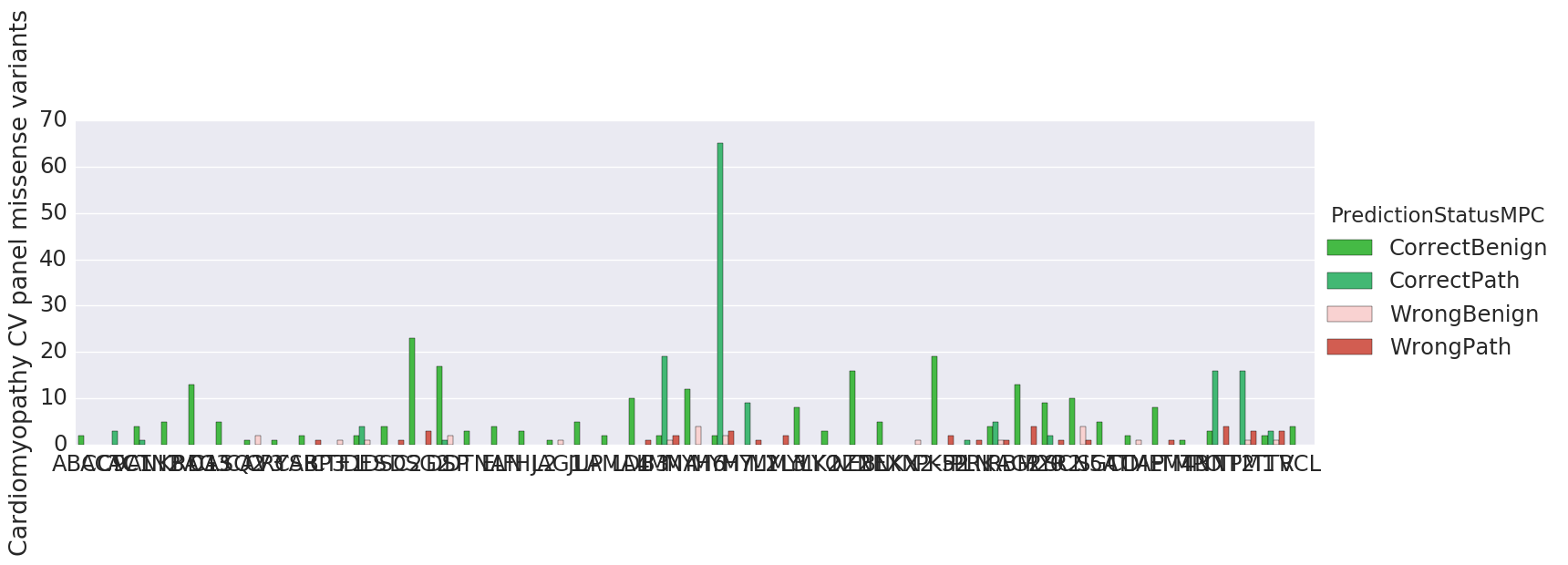

### clinvar_burden_gene_eval.png

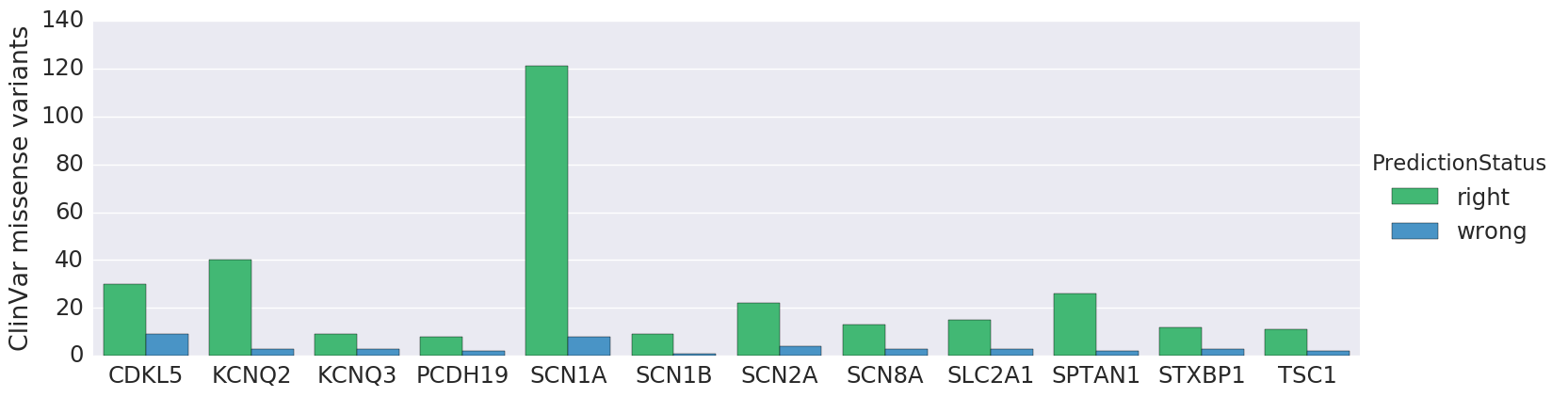

### clinvar_CDKL5_roc.png

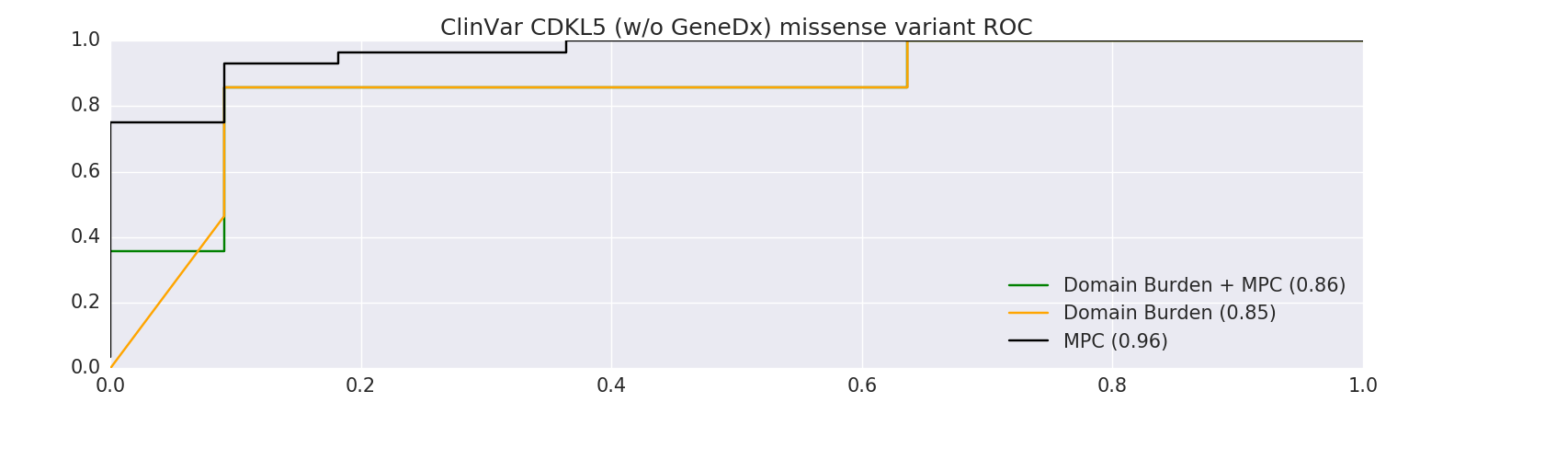

### clinvar_KCNQ2_roc.png

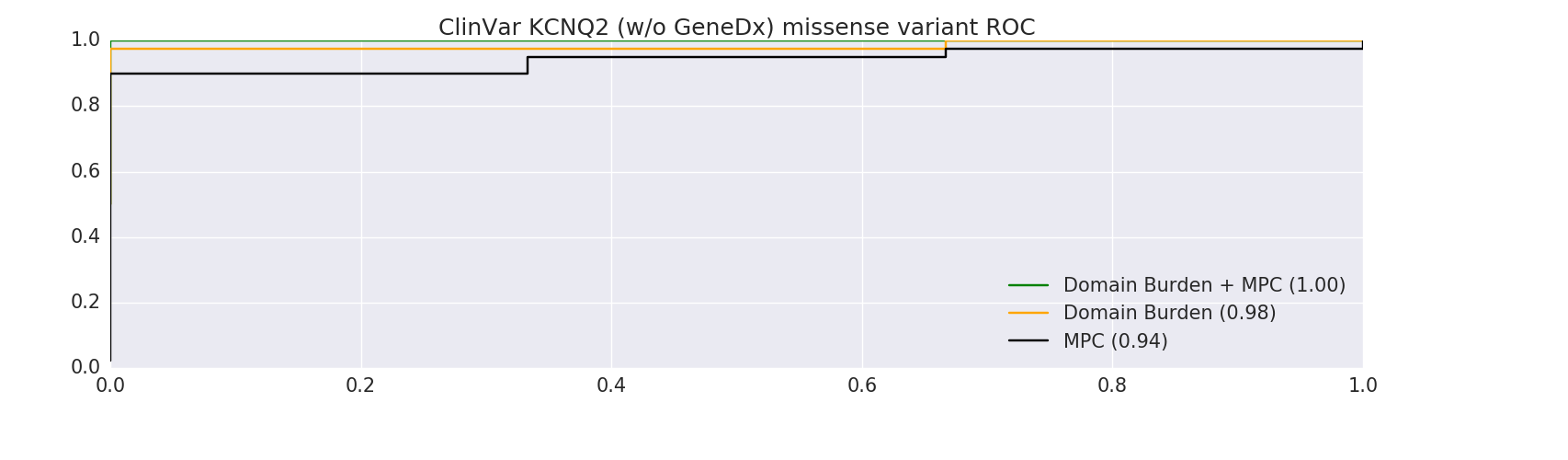

### clinvar_KCNQ3_roc.png

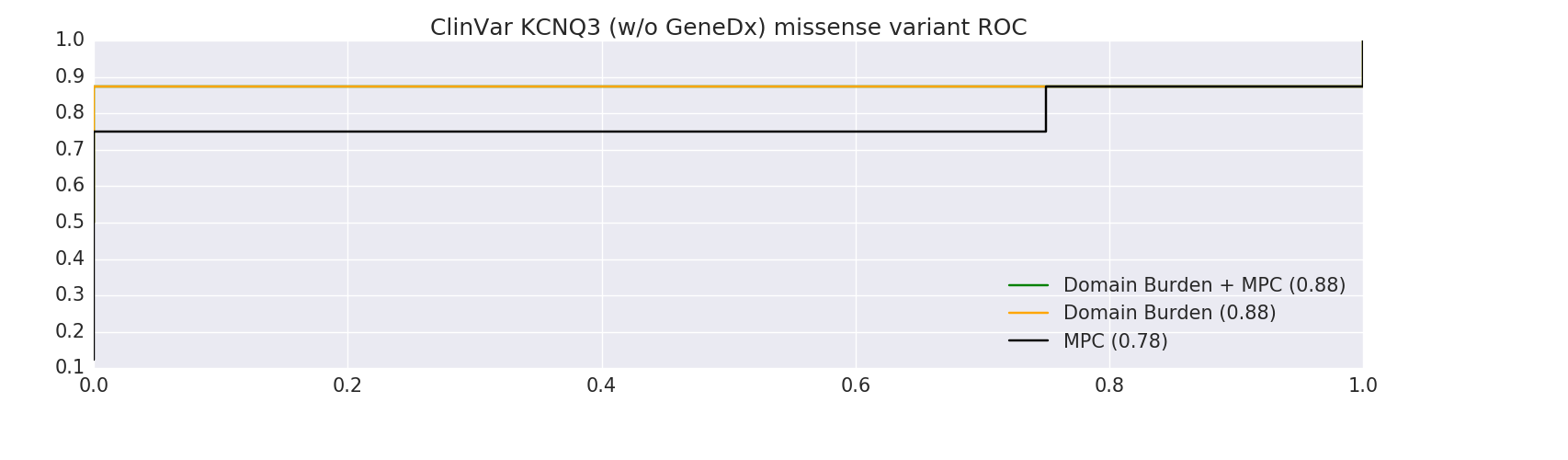

### clinvar_mis_mpc_roc.png

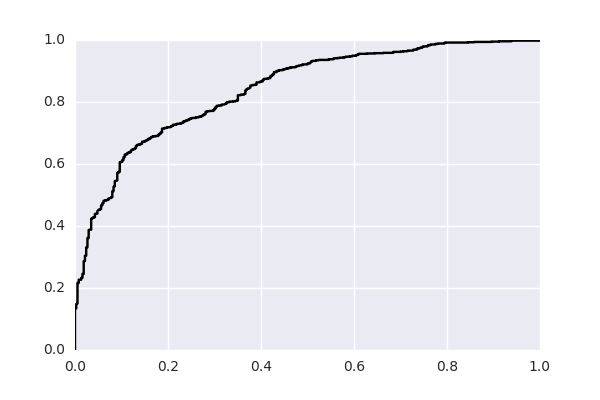

### clinvar_mis_mtr_roc.png

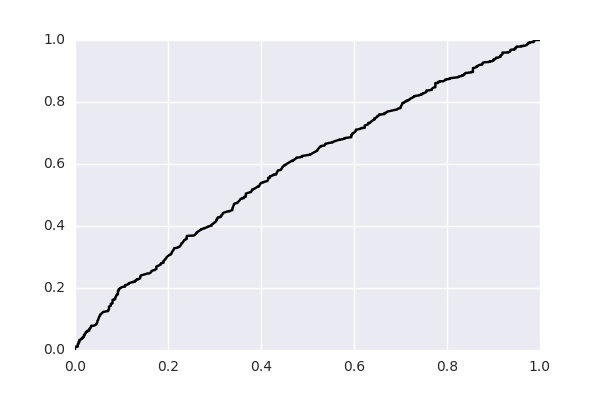

### clinvar_mpc_eval.png

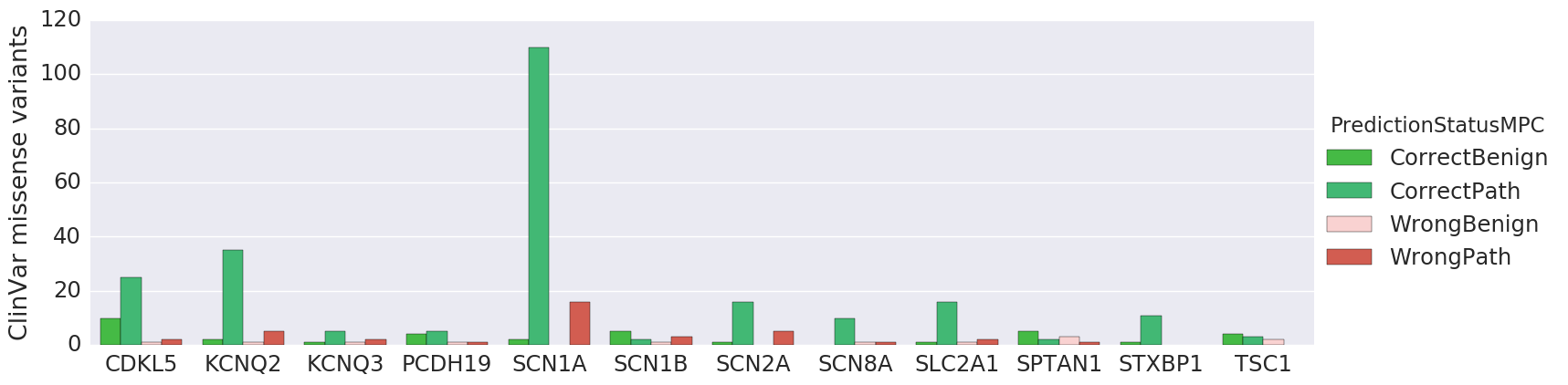

### clinvar_PCDH19_roc.png

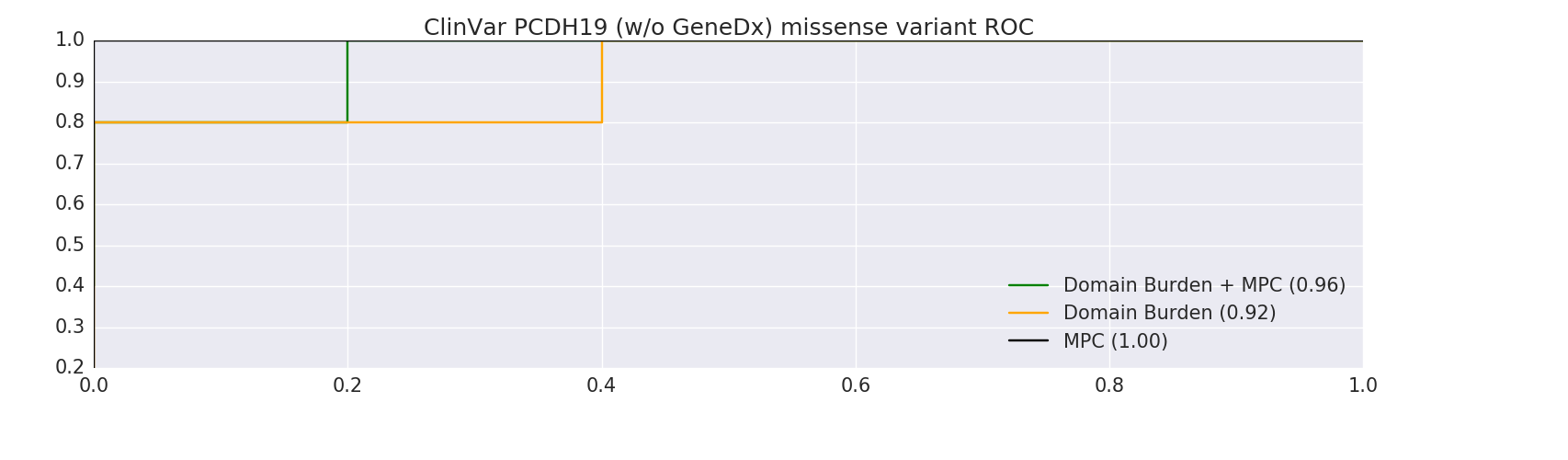

### clinvar_SCN1B_eval.png

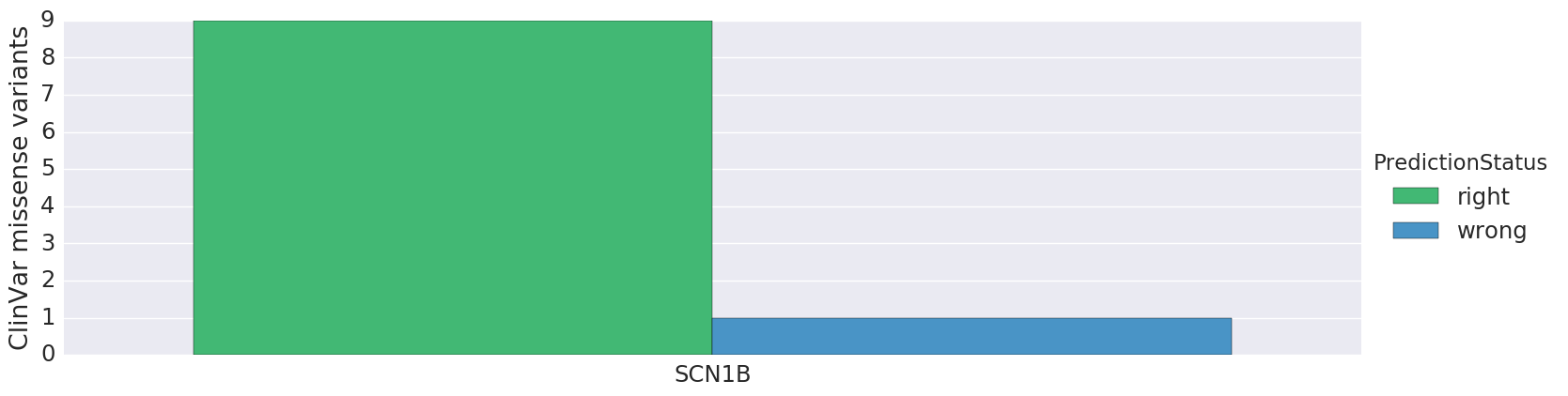

### clinvar_SCN1B_roc.png

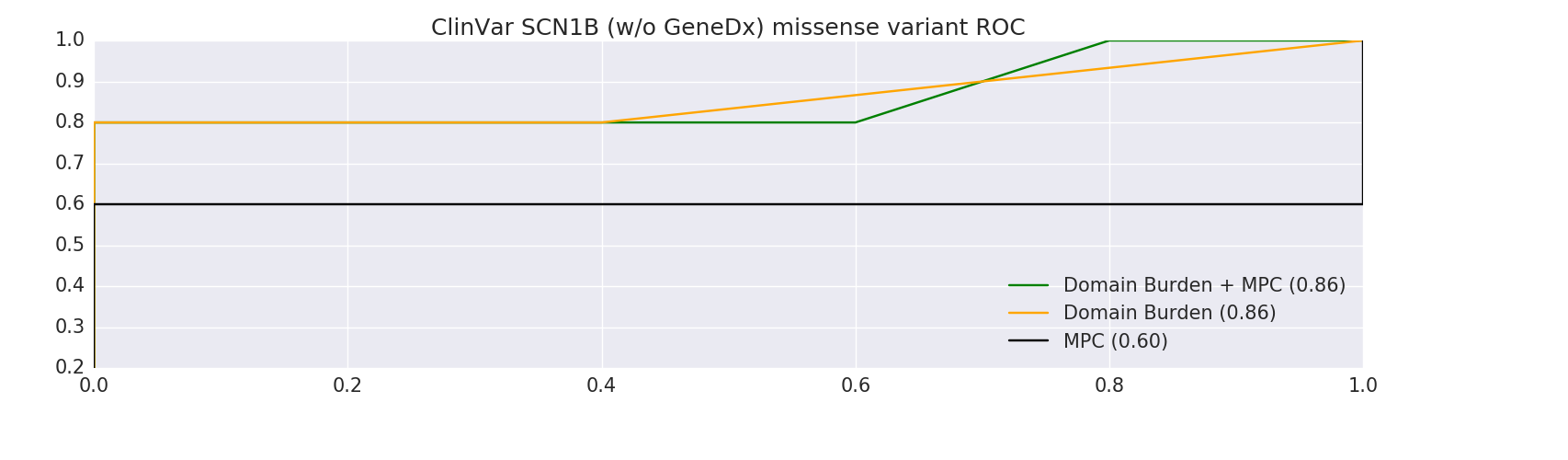

### clinvar_SCN8A_eval.png

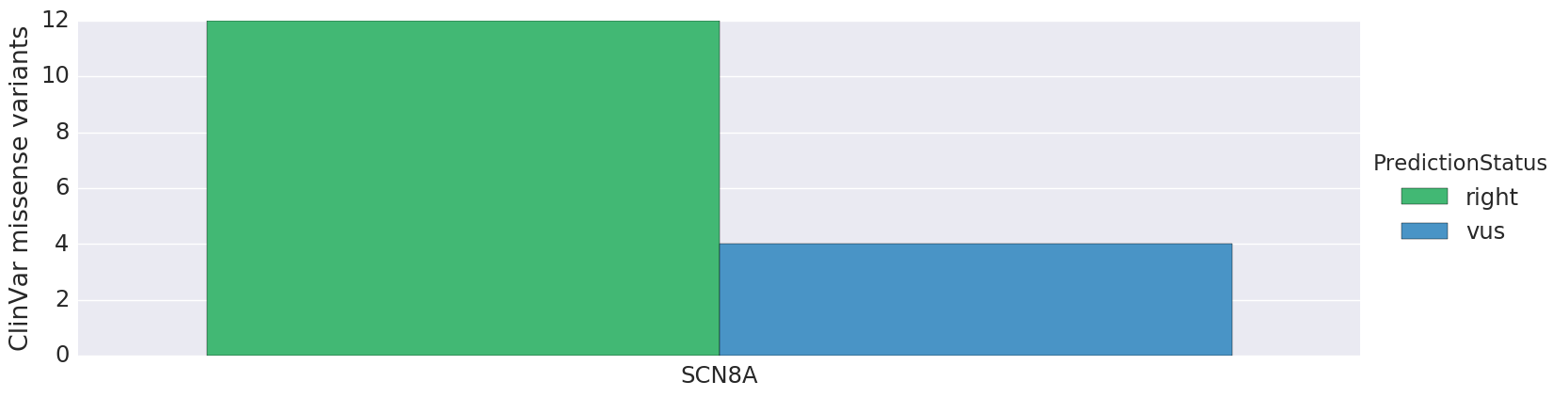

### clinvar_SCN8A_roc.png

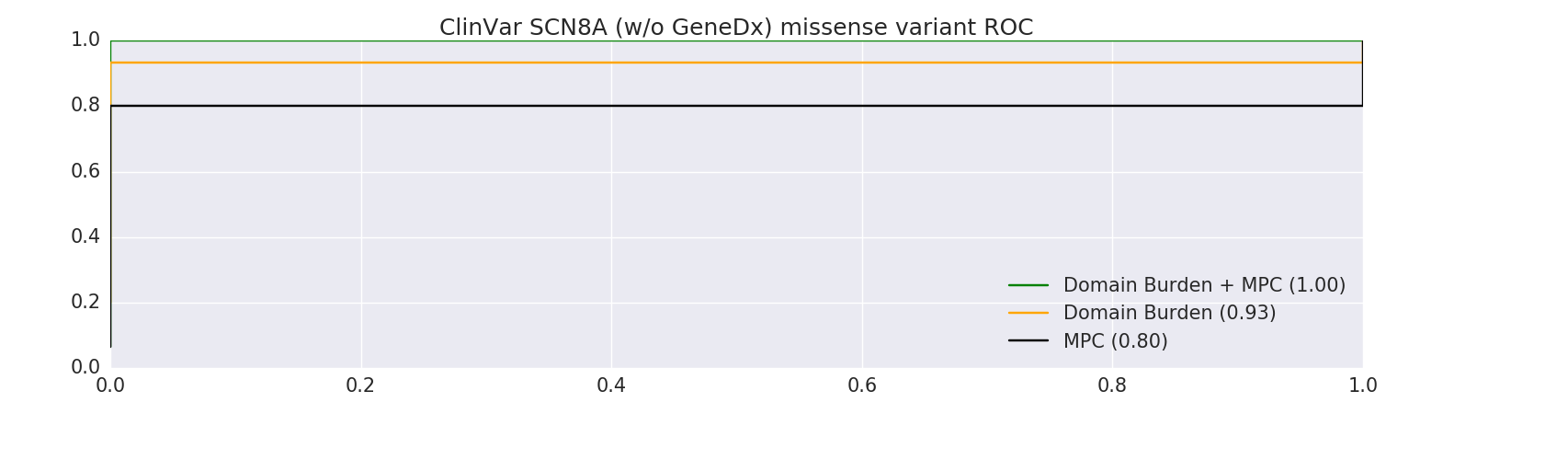

### clinvar_SLC2A1_roc.png

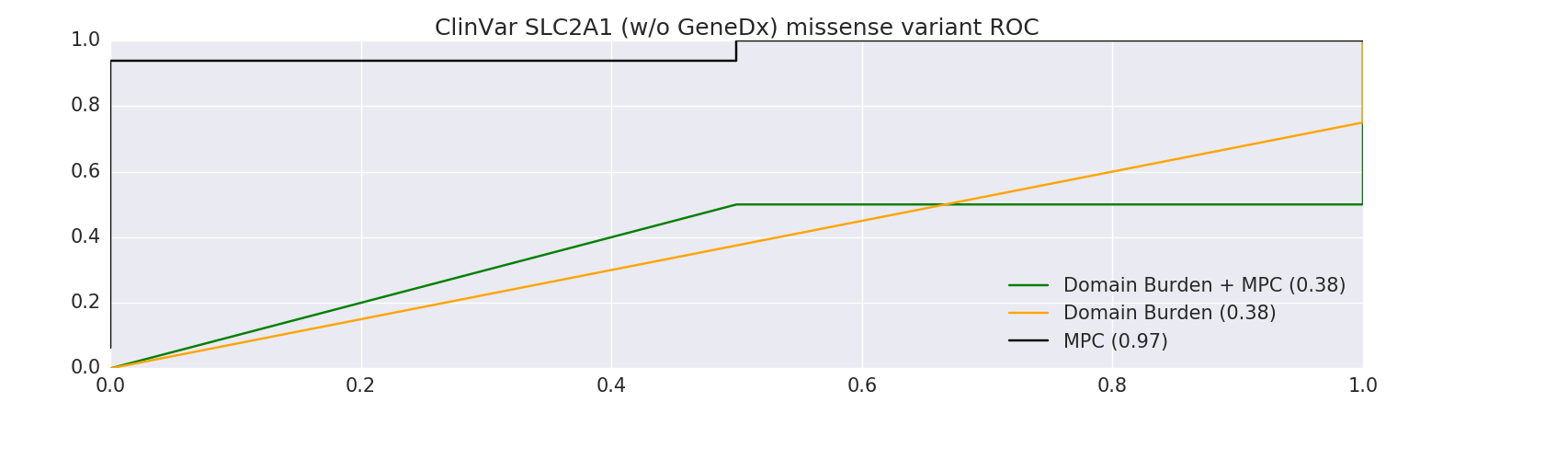

### denovo_mpc_eval.png

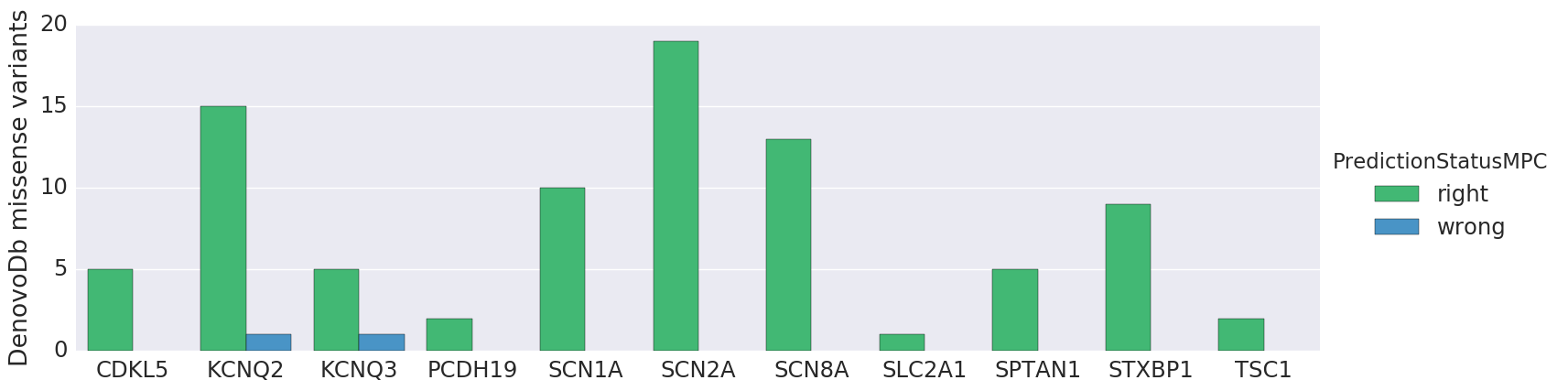

### eval_dat.var_count.png

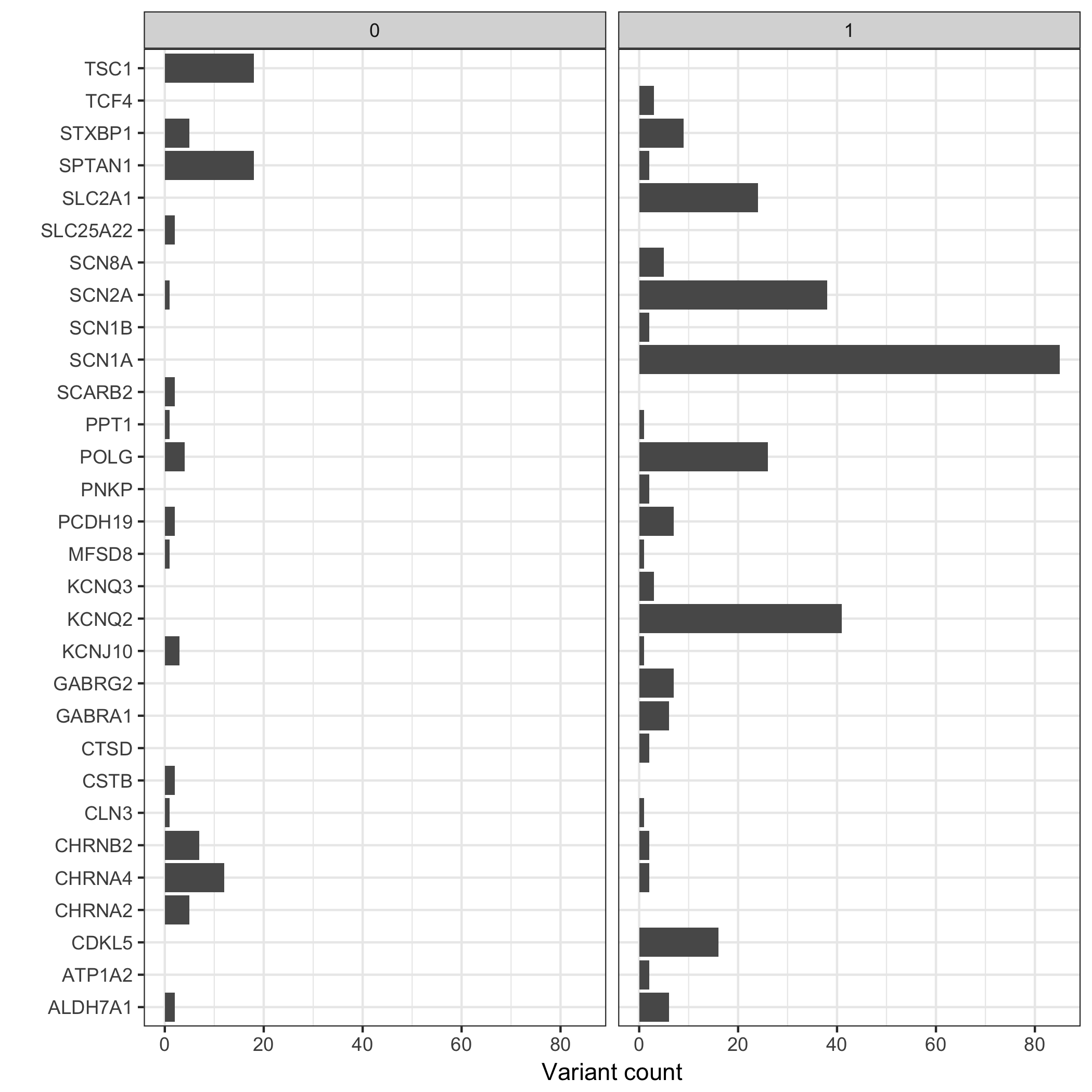

### Hearing Loss_cv_mpc_eval.png

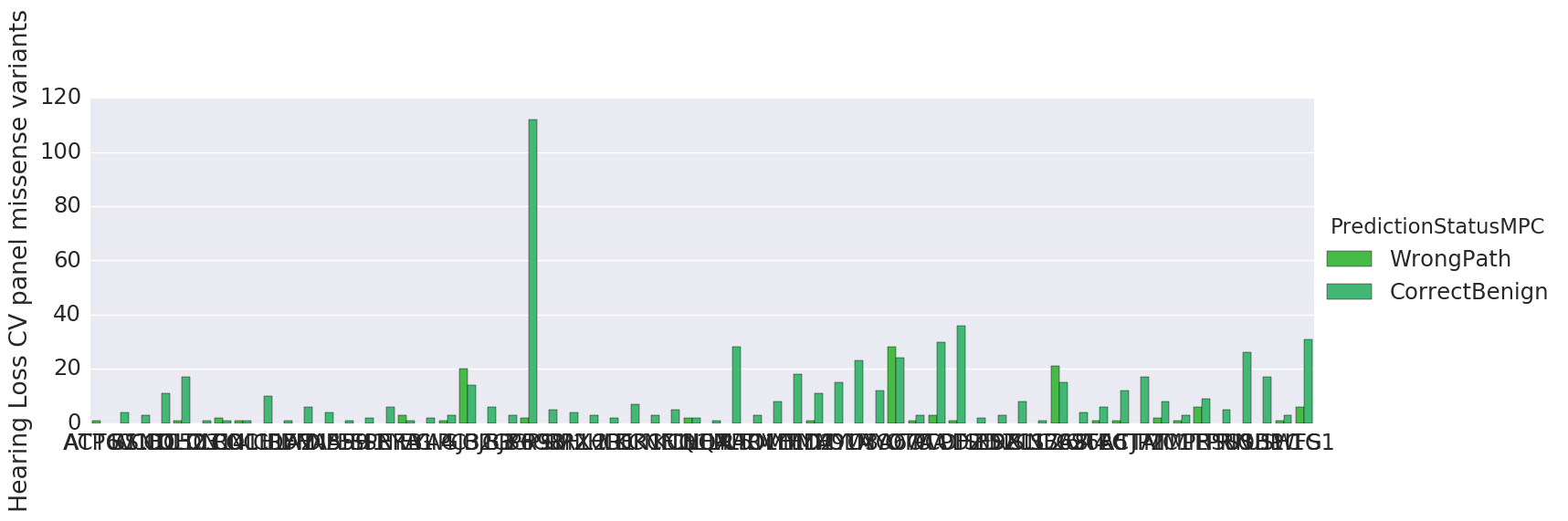

### mis_mpc_roc.png

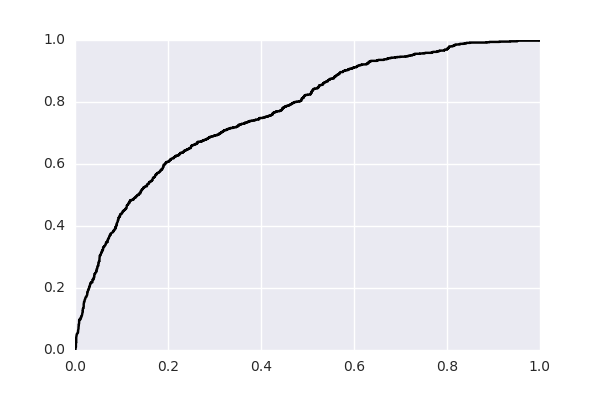

### missense_clinvar_roc_feature_union.PCDH19.png

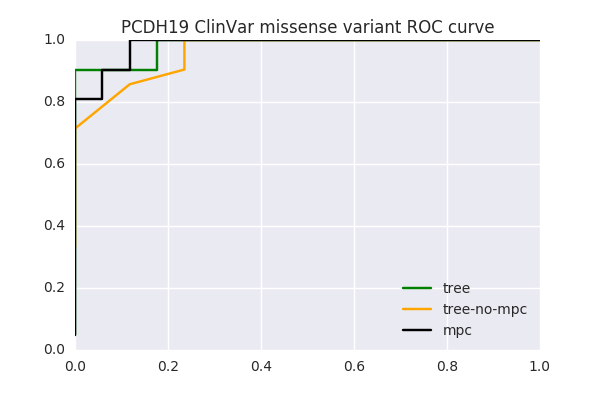

### missense_clinvar_roc_feature_union.png

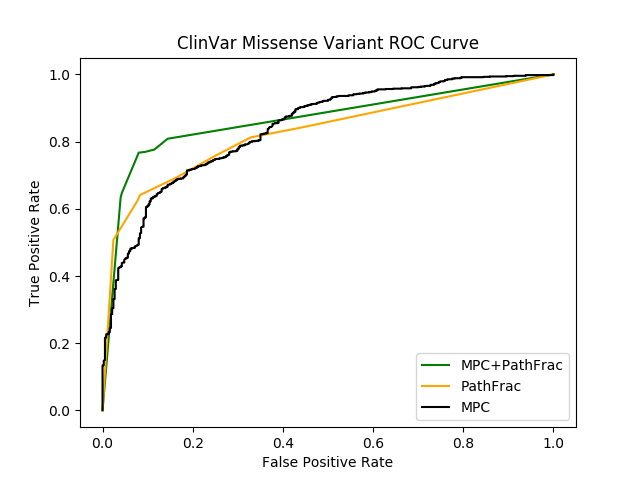

### missense_clinvar_roc_feature_union.SCN1A,SCN2A,KCNQ2,KCNQ3.png

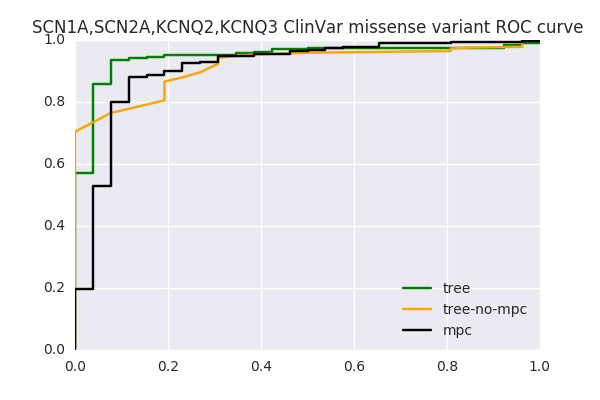

### missense_clinvar_roc_feature_union.SCN8A.png

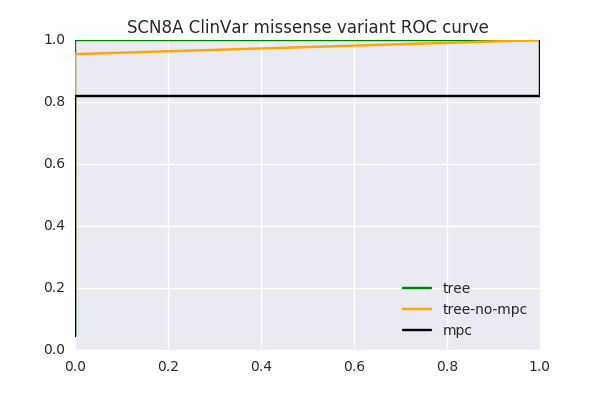

### missense_clinvar_roc_feature_union.SLC2A1.png

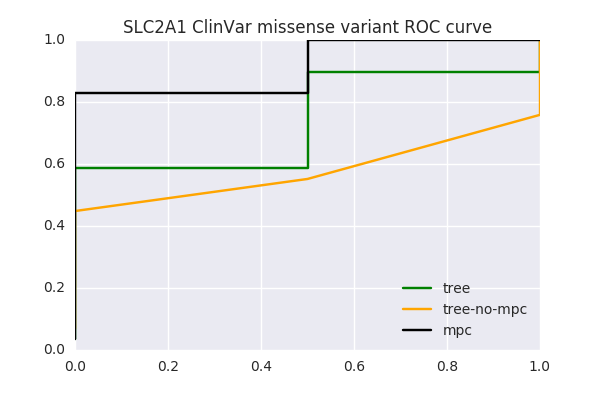

### rare.benign_frac_w_vus.pfam.mpcLow_0.mpcHigh_100.png

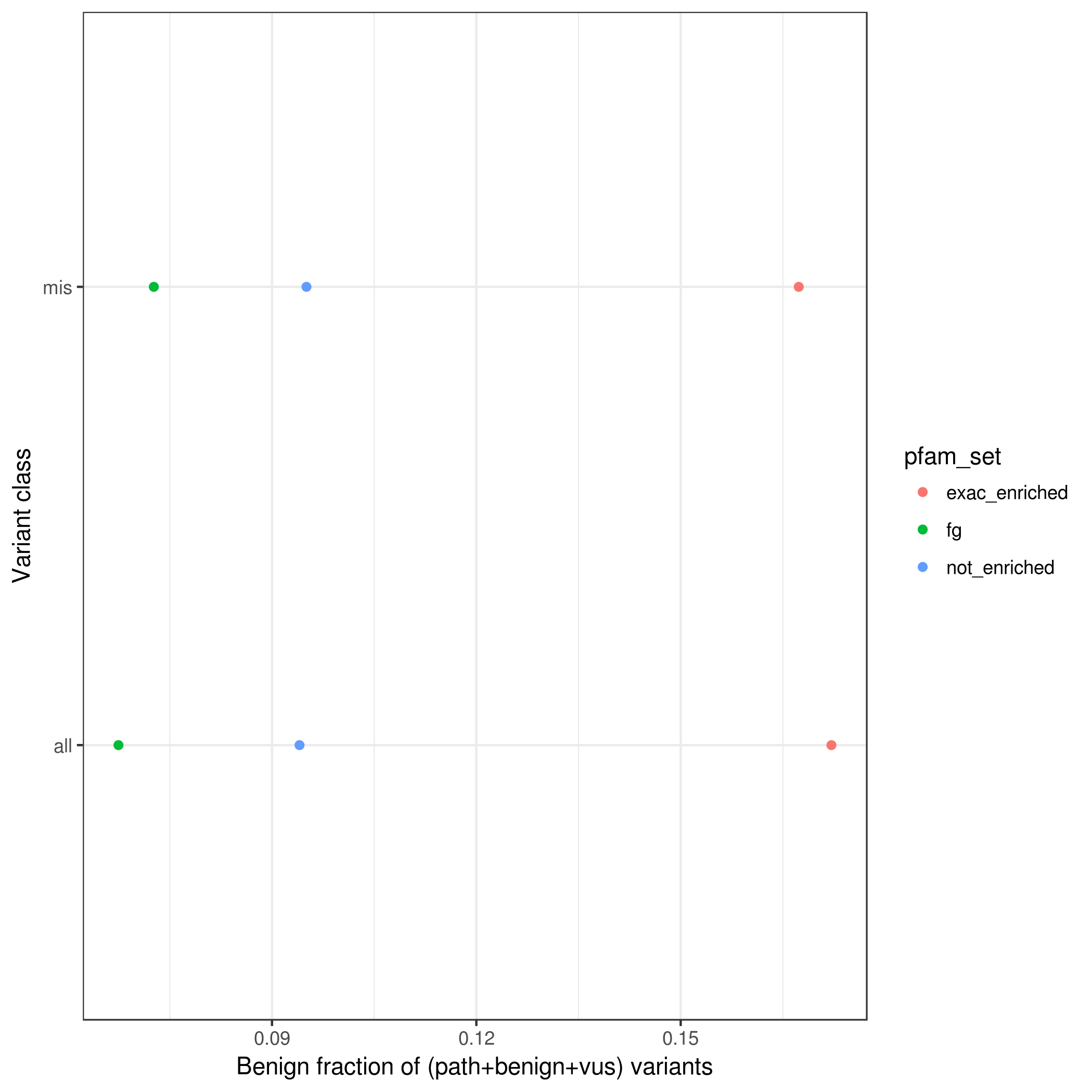

### rare.benign_frac_w_vus.pfam.mpcLow_1.4.mpcHigh_100.png

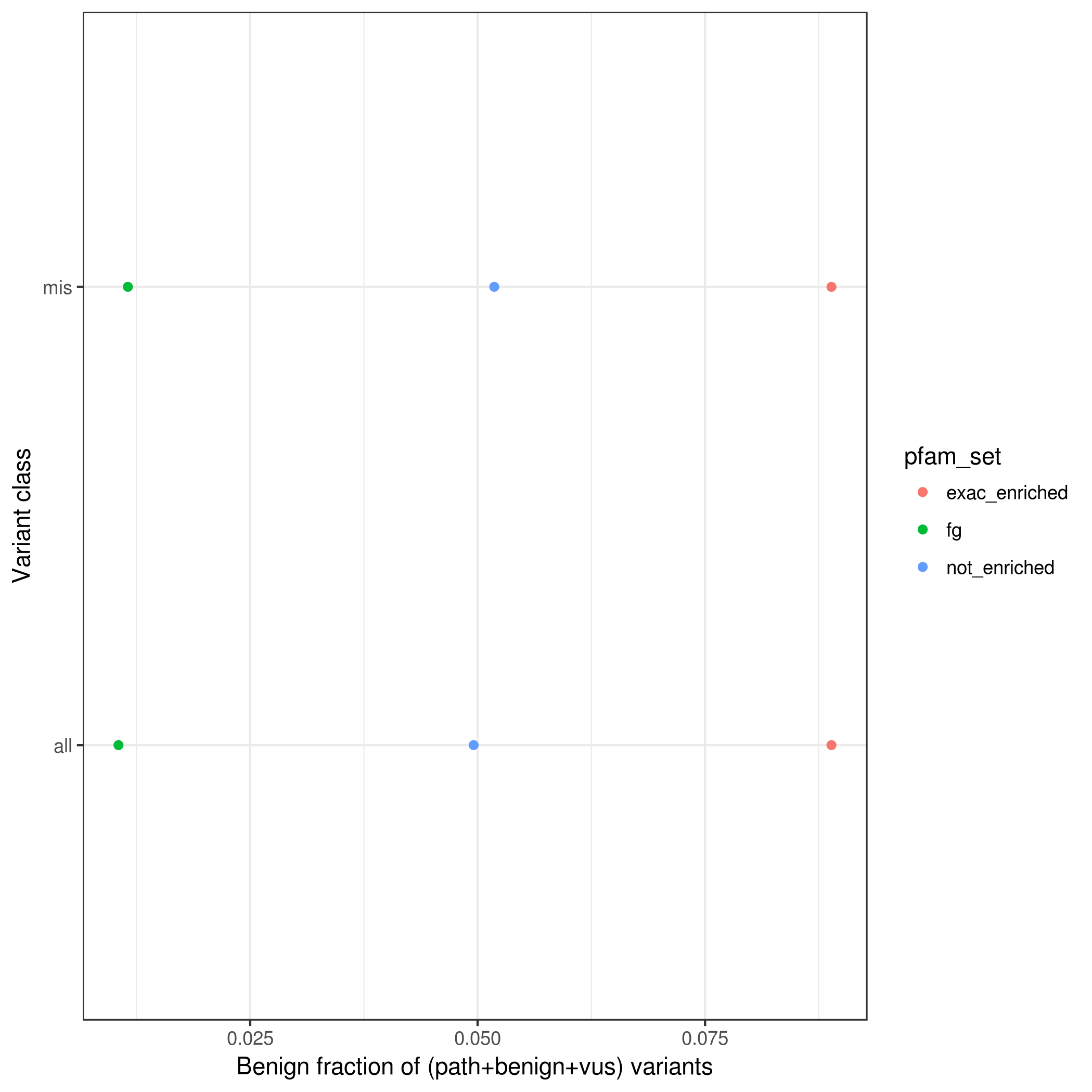

### rare.benign_frac_w_vus.pfamMerge.mpc_100.png

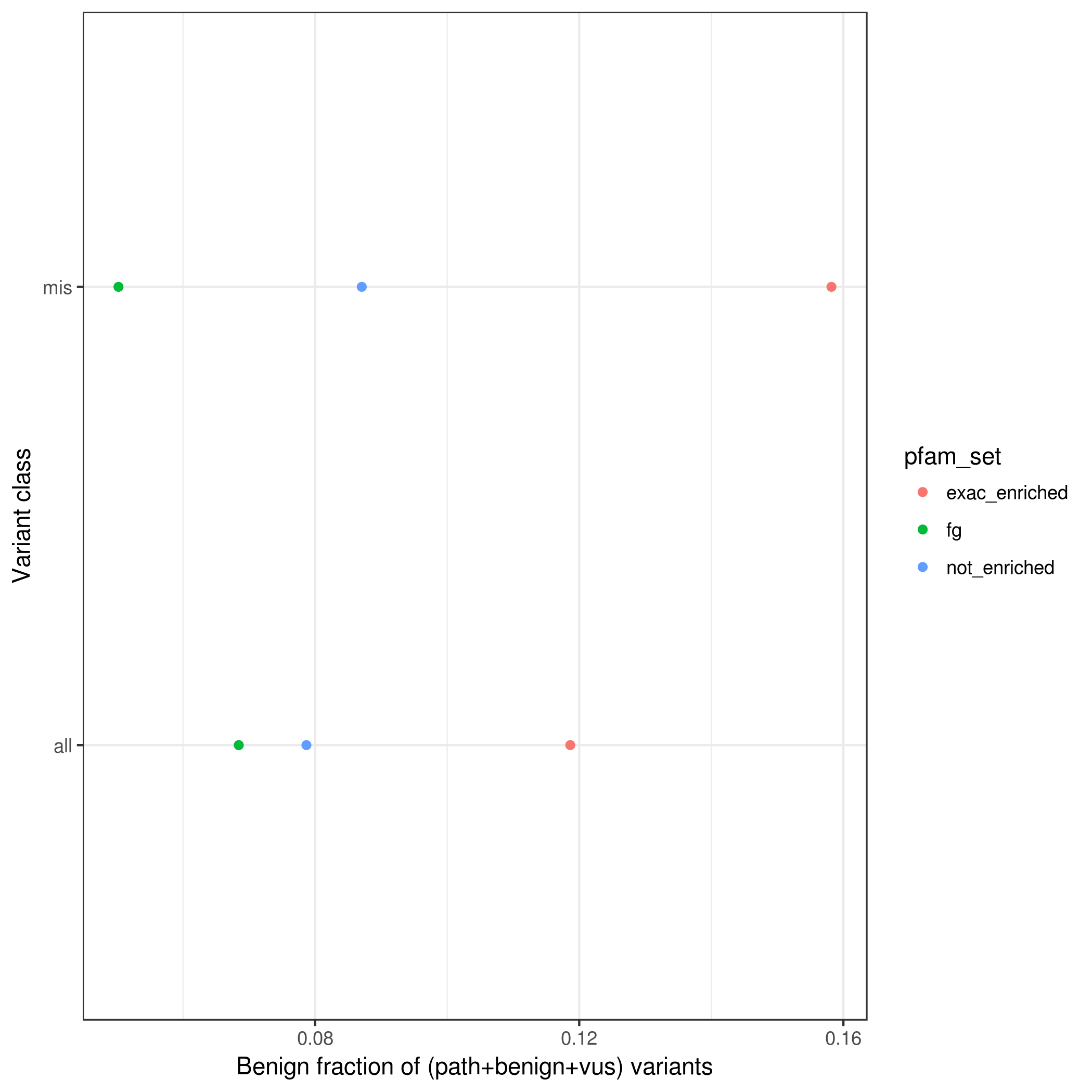

### rare.benign_frac_w_vus.pfamMerge.mpcLow_1.4.mpcHigh_100.png

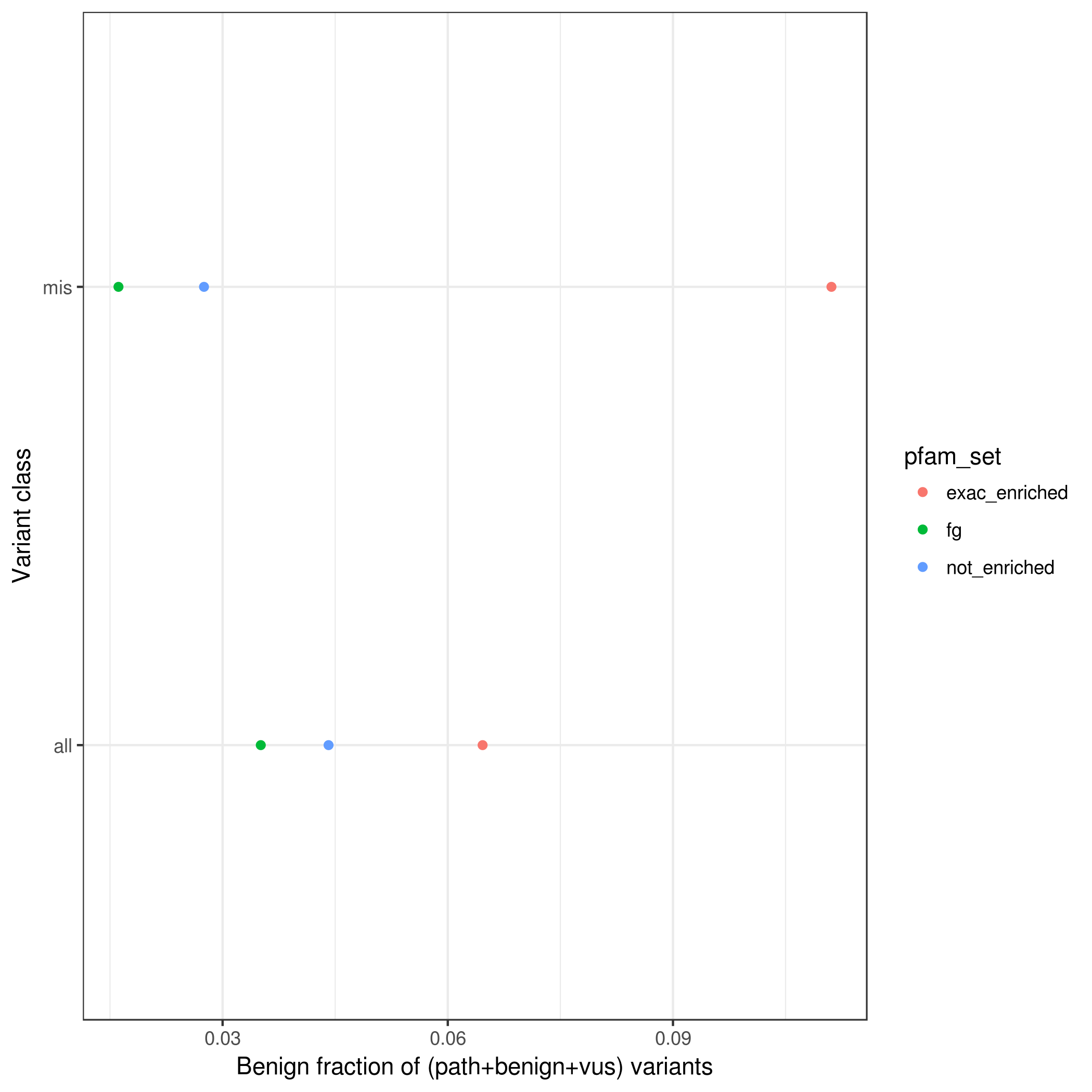

### rare.path_frac_w_vus.pfam.mpcLow_0.mpcHigh_100.png

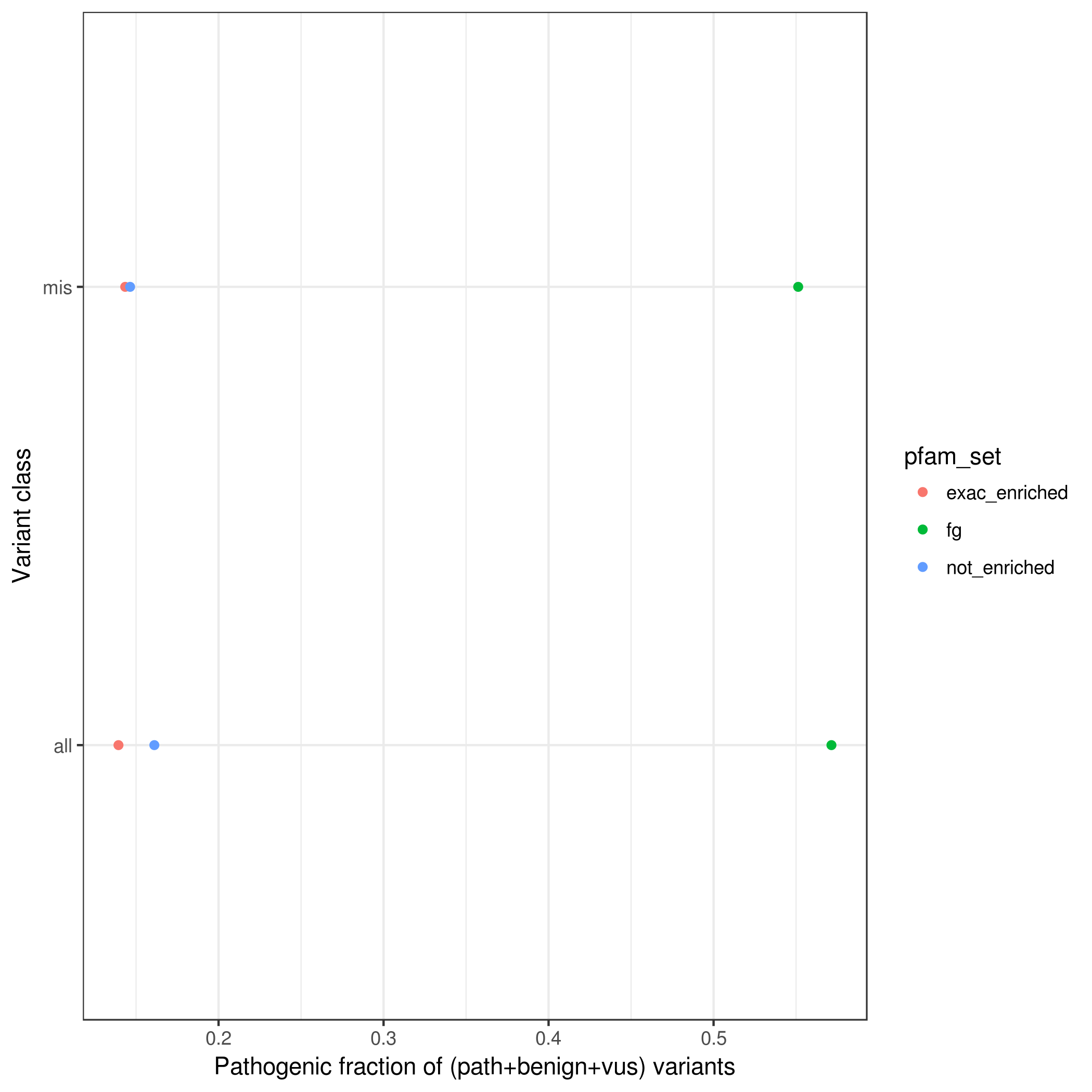

### rare.path_frac_w_vus.pfamMerge.mpcLow_1.4.mpcHigh_100.png

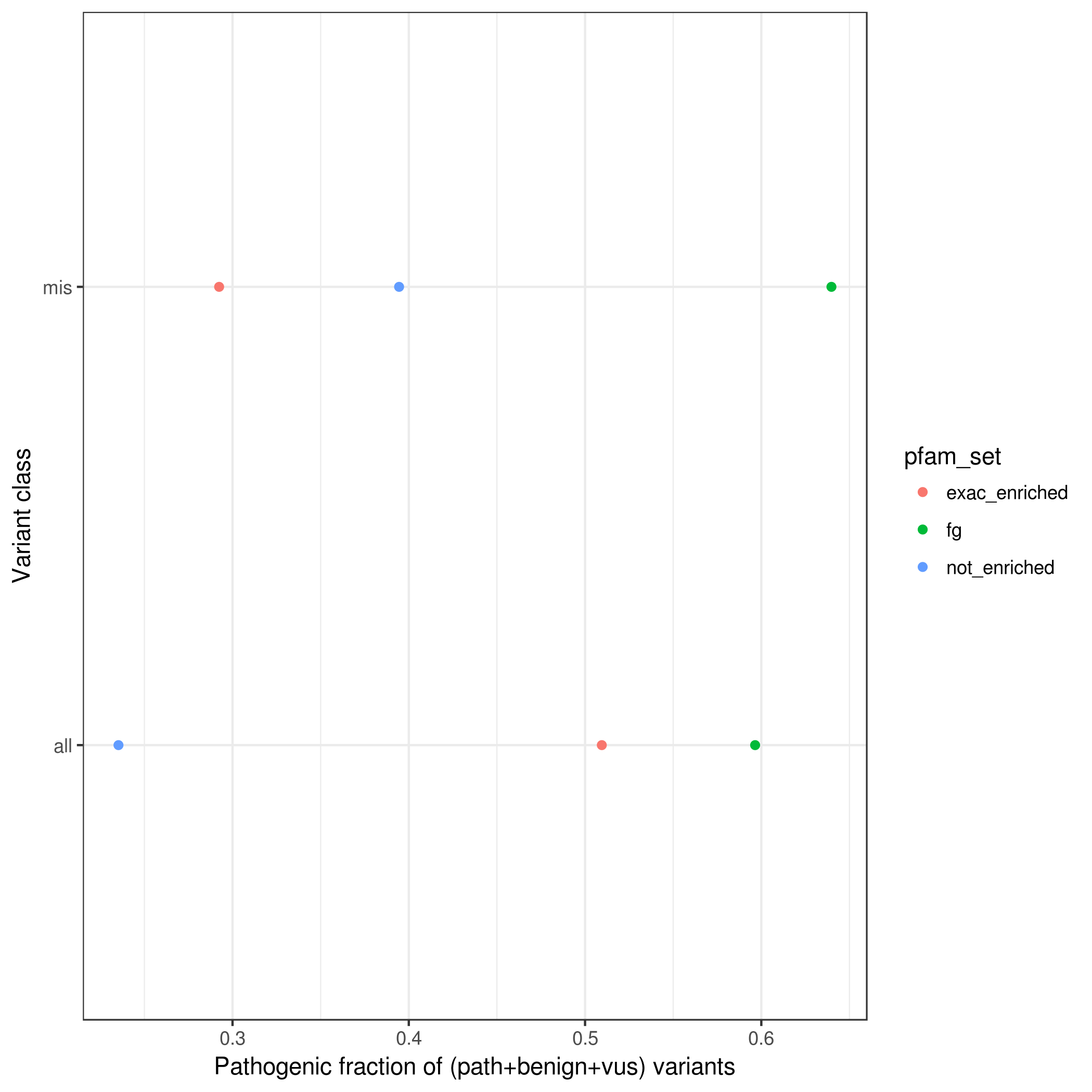

### rare.path_frac_wo_vus.pfam.mpcLow_0.mpcHigh_100.png

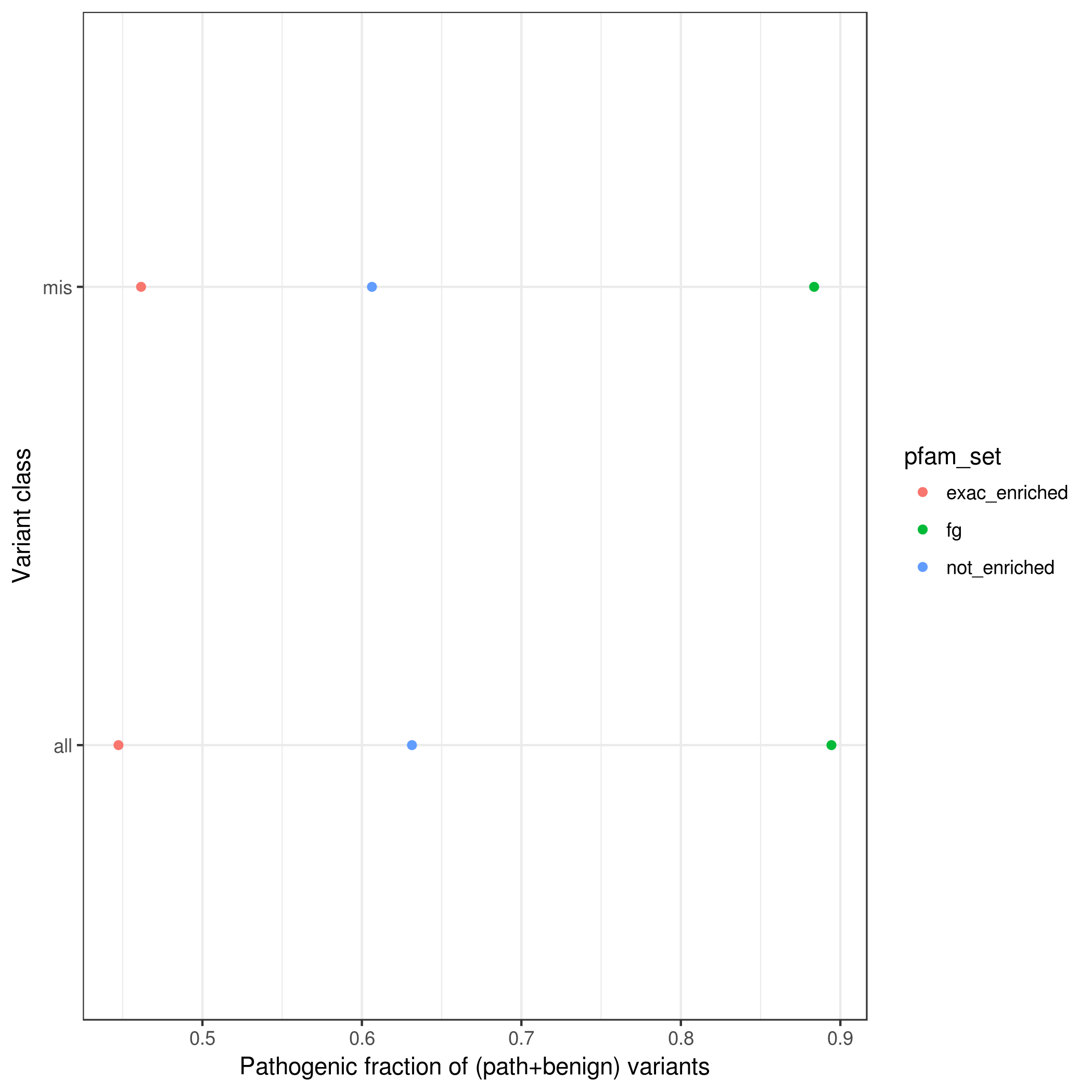

### rare.path_frac_wo_vus.pfamMerge.mpcLow_0.mpcHigh_100.png

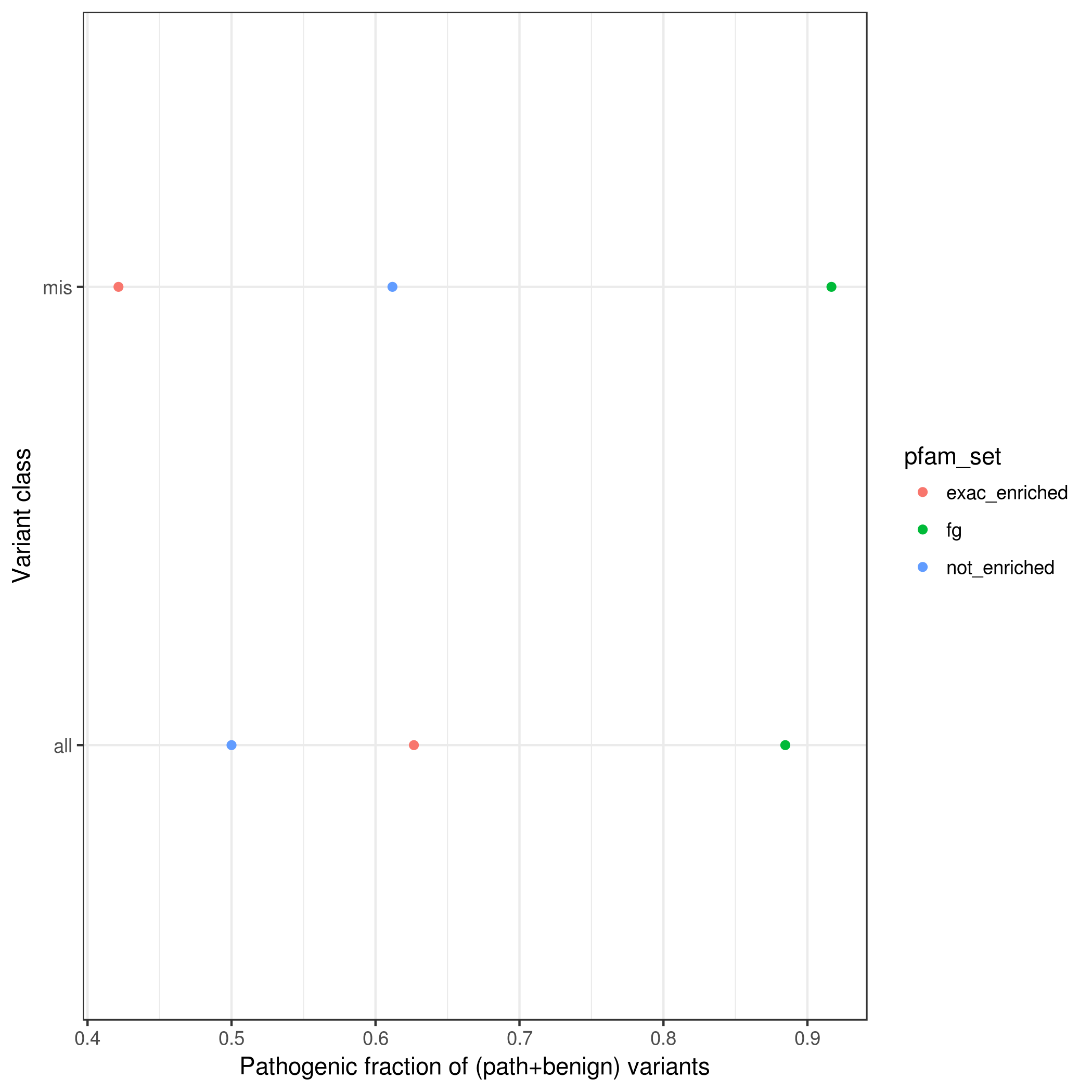

### rare.var_counts_by_enrichment.pfam.mpc_1.4.png

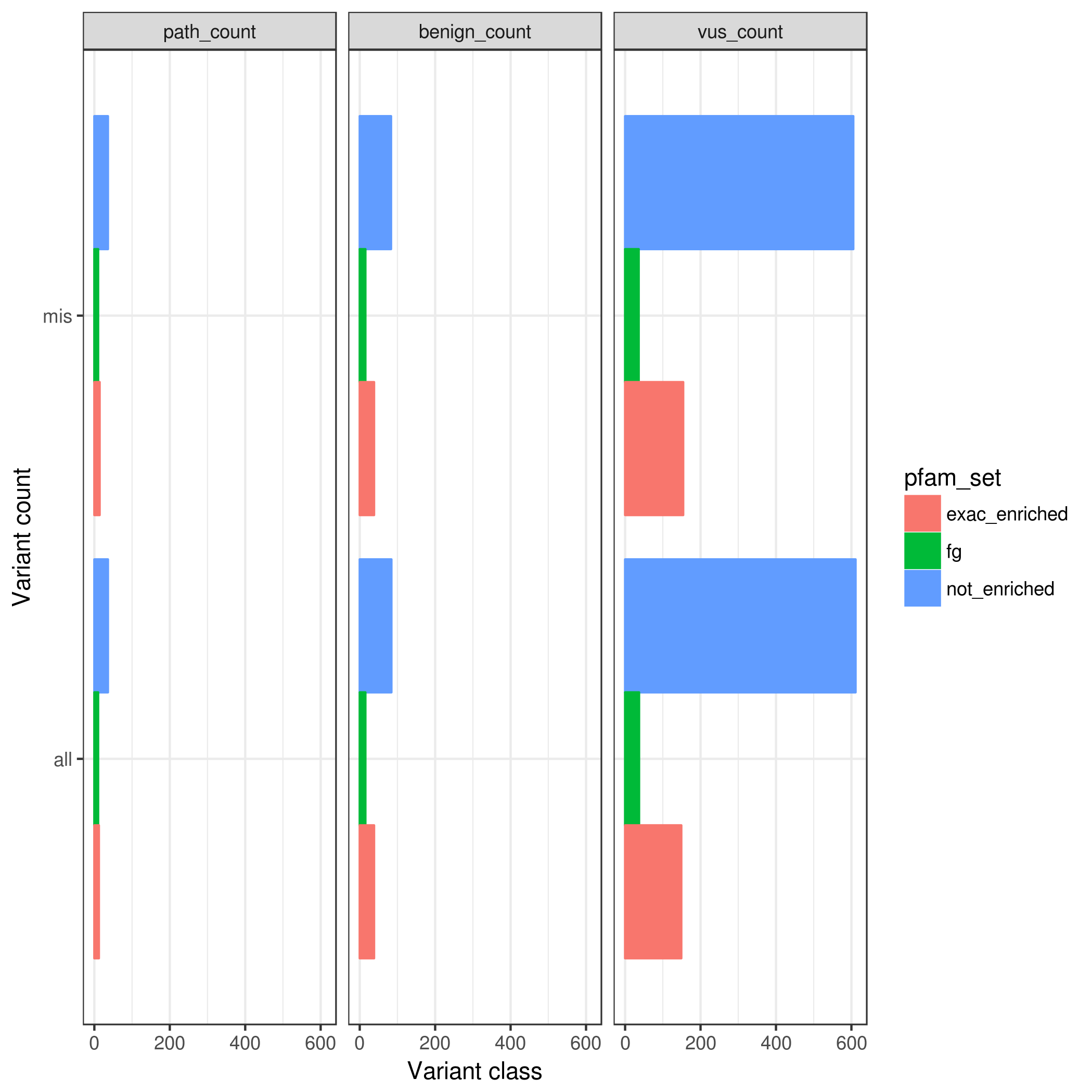

### rare.var_counts_by_enrichment.pfamMerge.mpcLow_1.4.mpcHigh_100.png

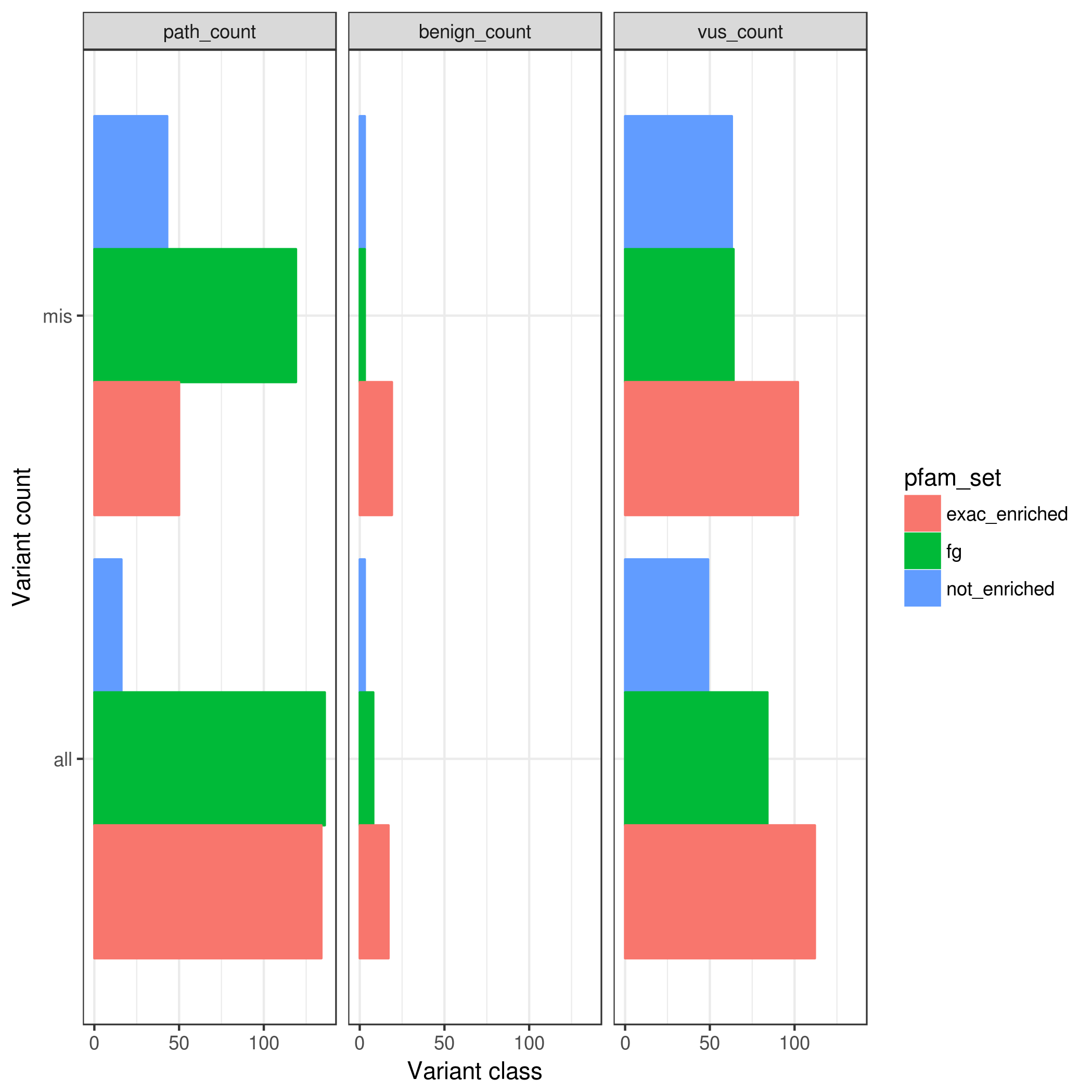

### rarem.benign_frac_w_vus.pfam.mpc_1.4.png

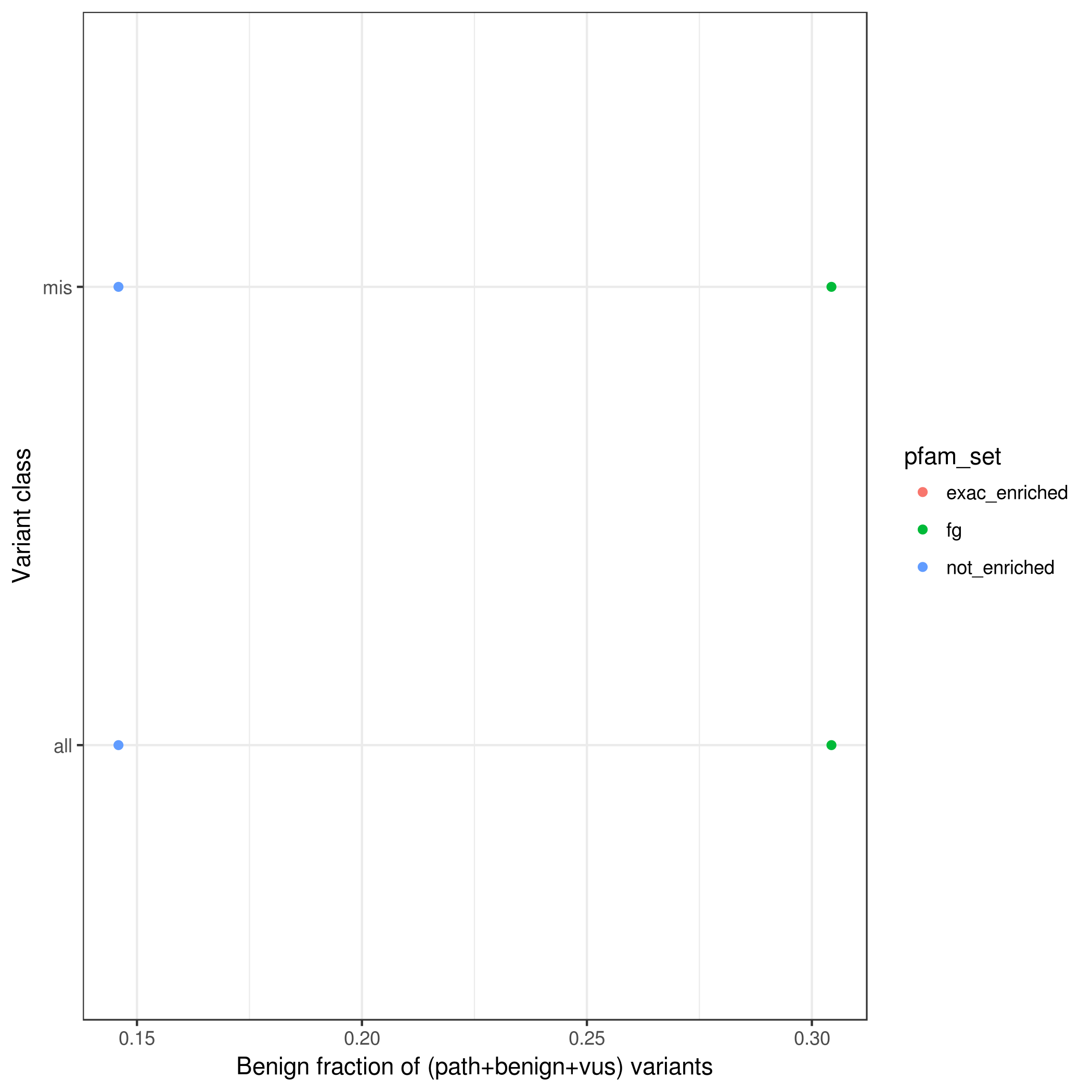

### rarem.path_frac_w_vus.pfam.mpc_1.4.png

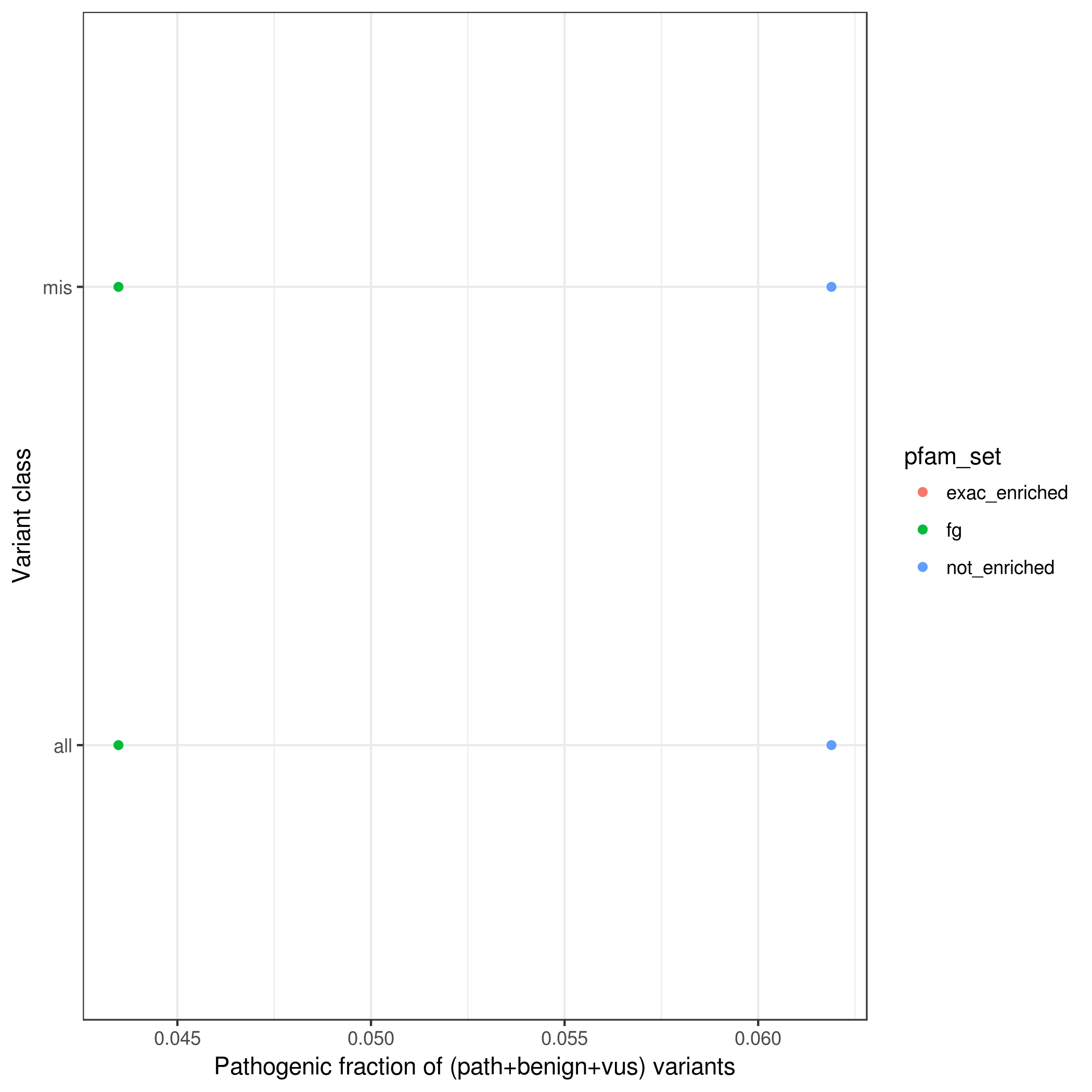

### rarem.path_frac_w_vus.pfam.mpcLow_0.mpcHigh_100.png

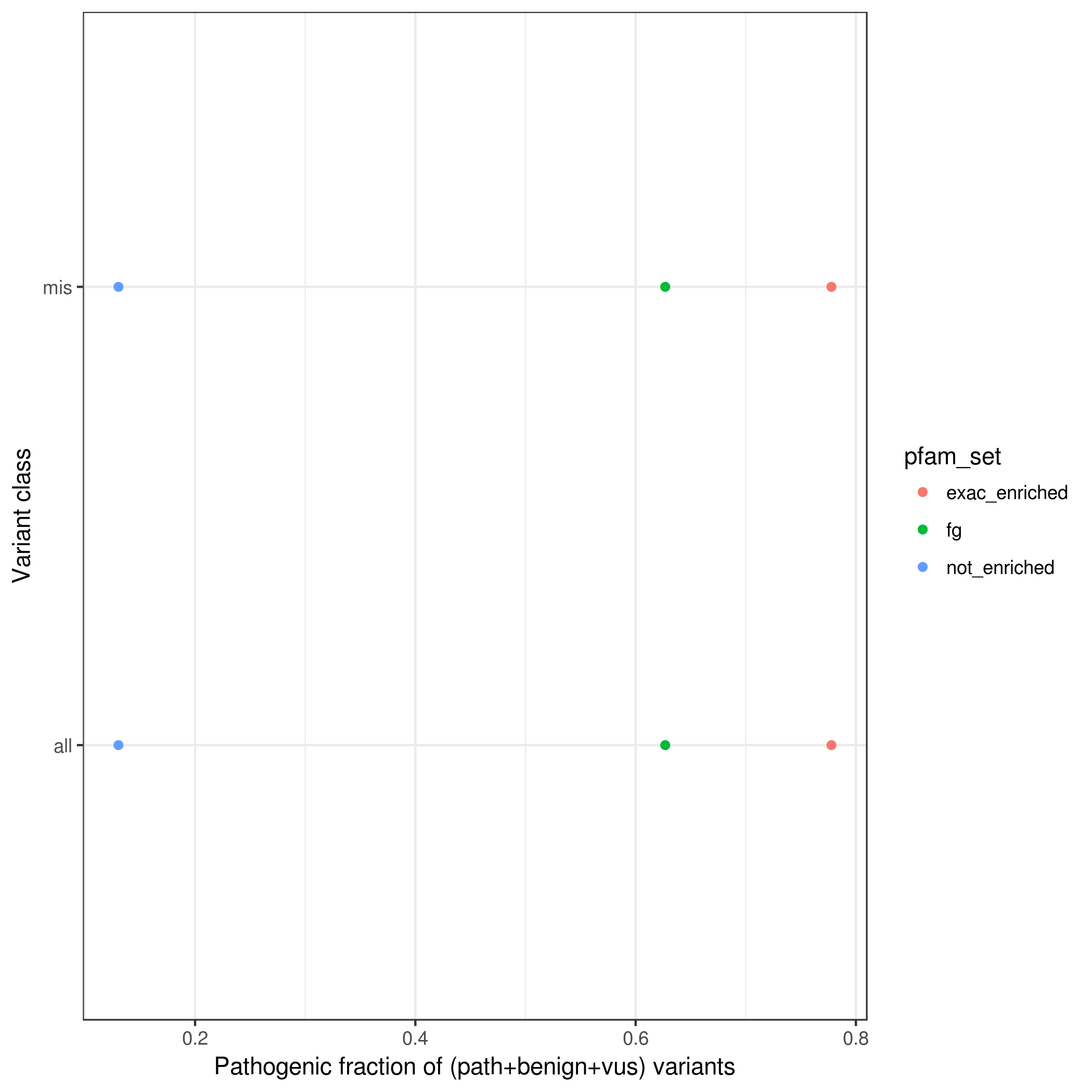

### rarem.path_frac_w_vus.pfamMerge.mpcLow_0.mpcHigh_100.png

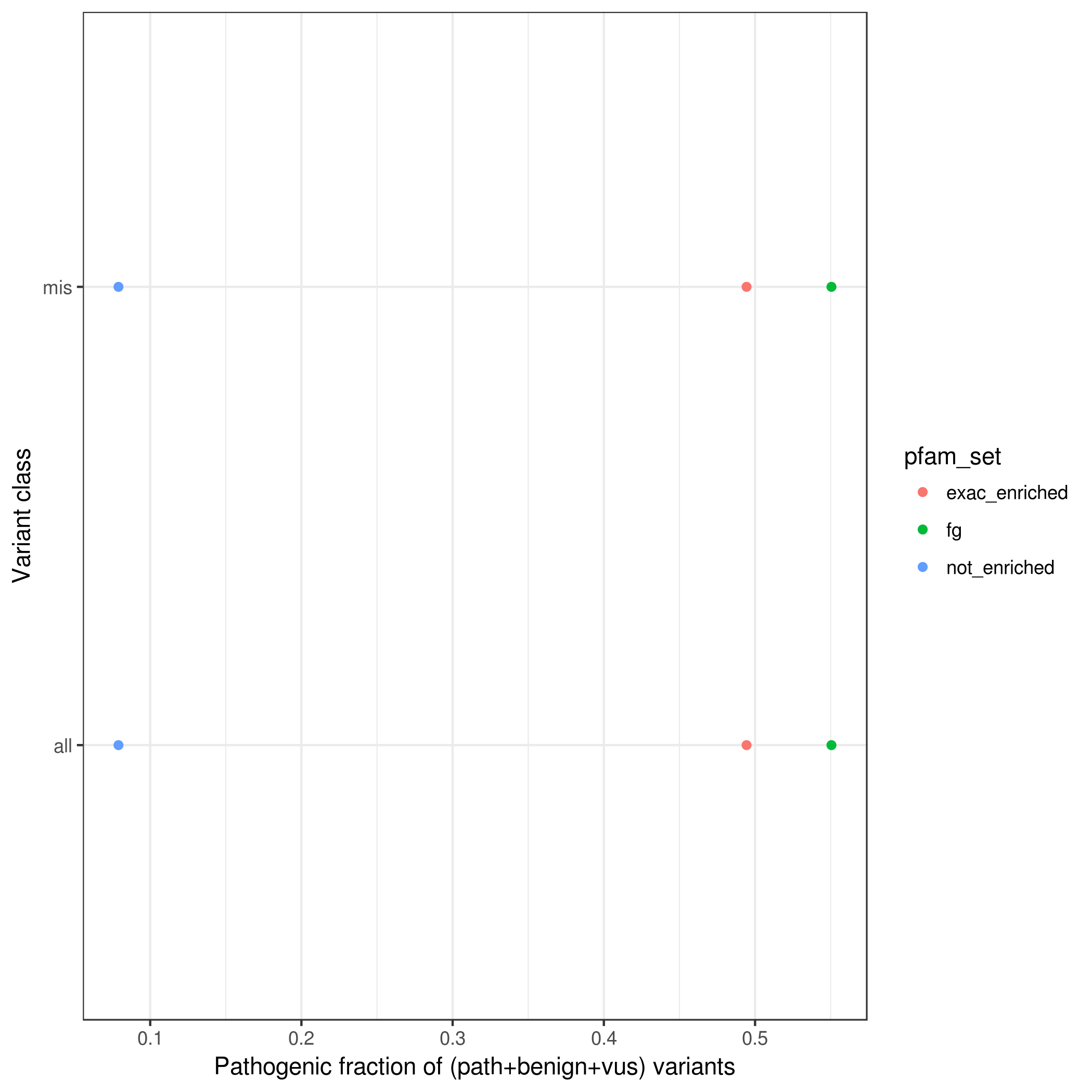

### rarem.path_frac_w_vus.pfamMerge.mpcLow_1.4.mpcHigh_100.png

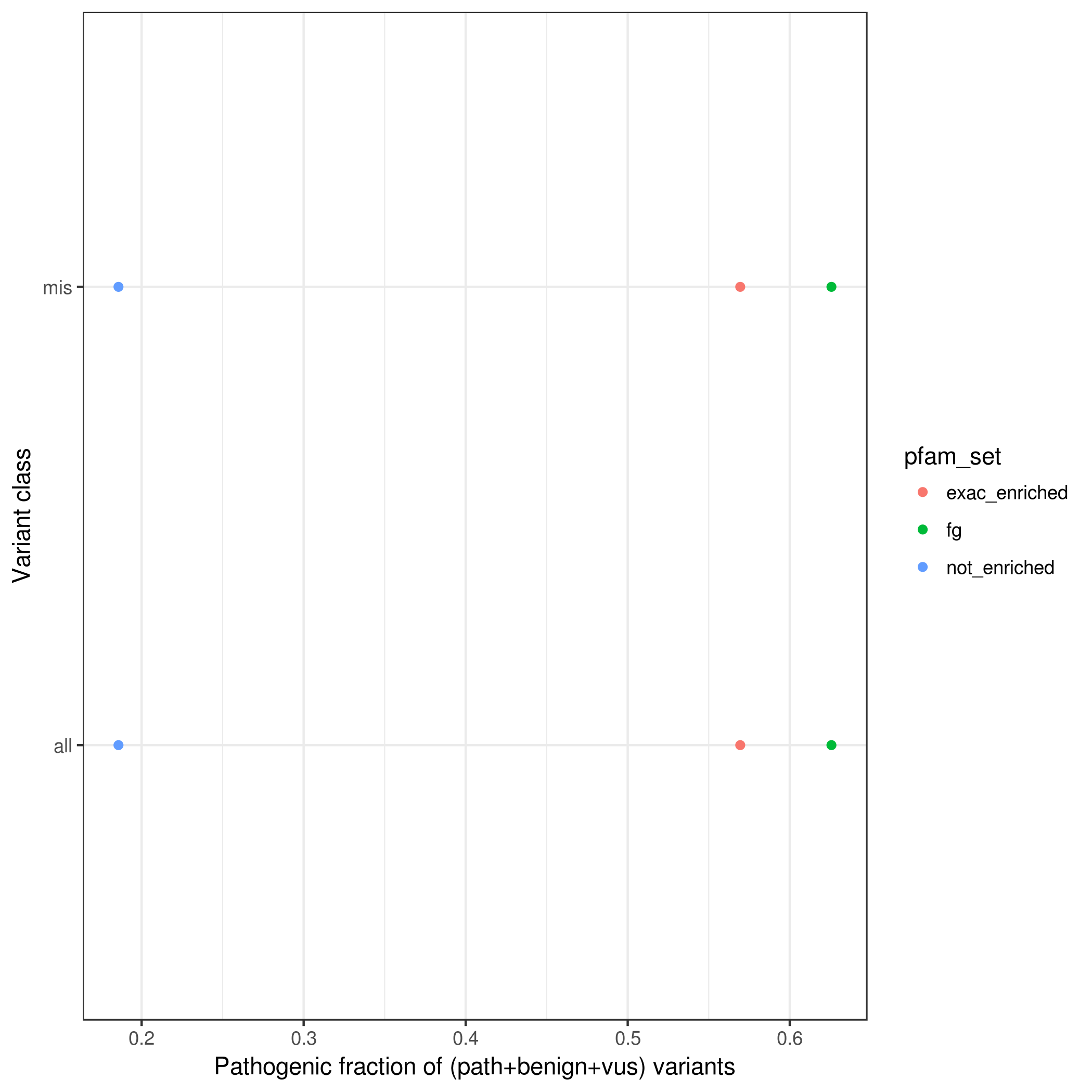

### rarem.path_frac_wo_vus.pfam.mpcLow_0.mpcHigh_100.png

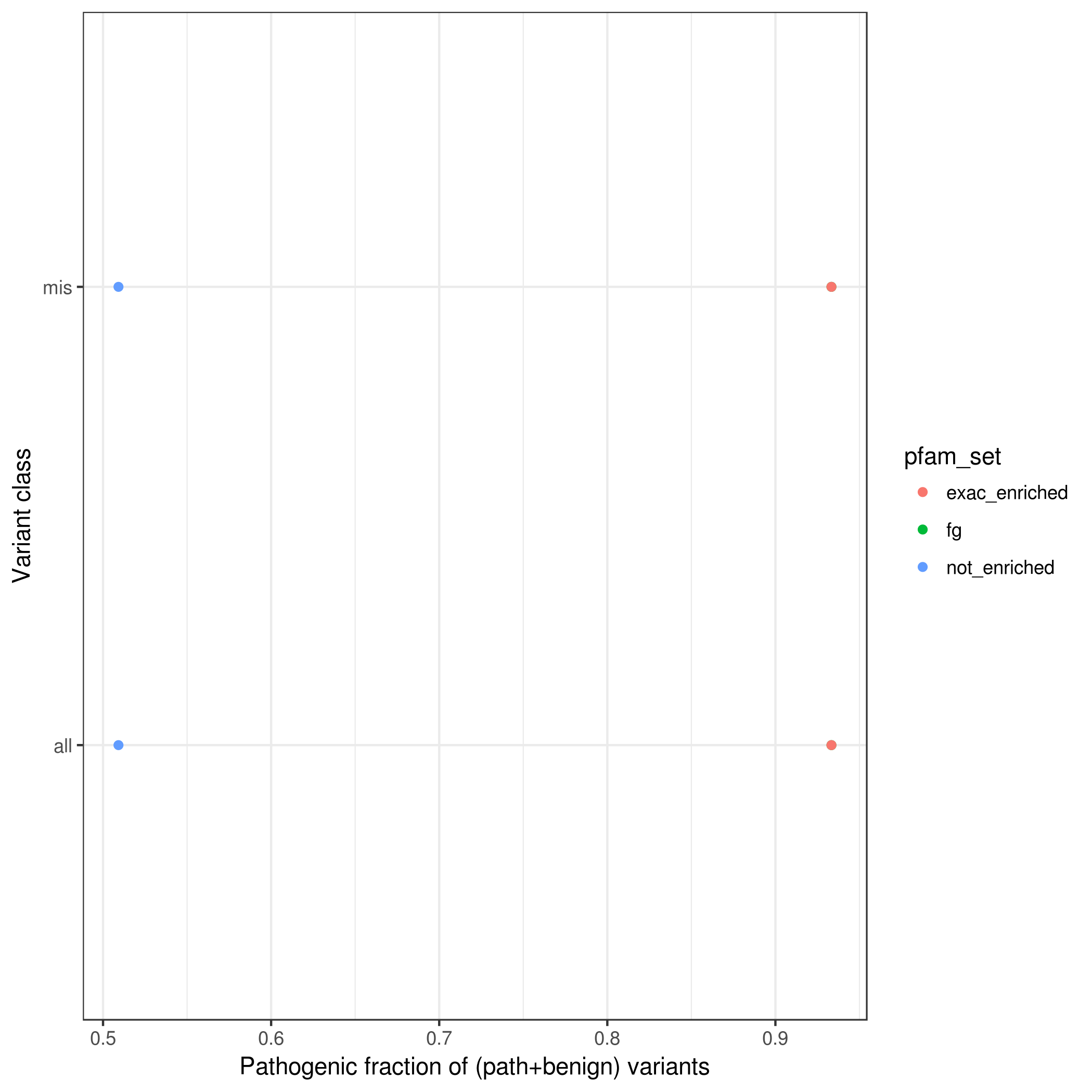

### rarem.path_frac_wo_vus.pfamMerge.mpcLow_0.mpcHigh_1.4.png

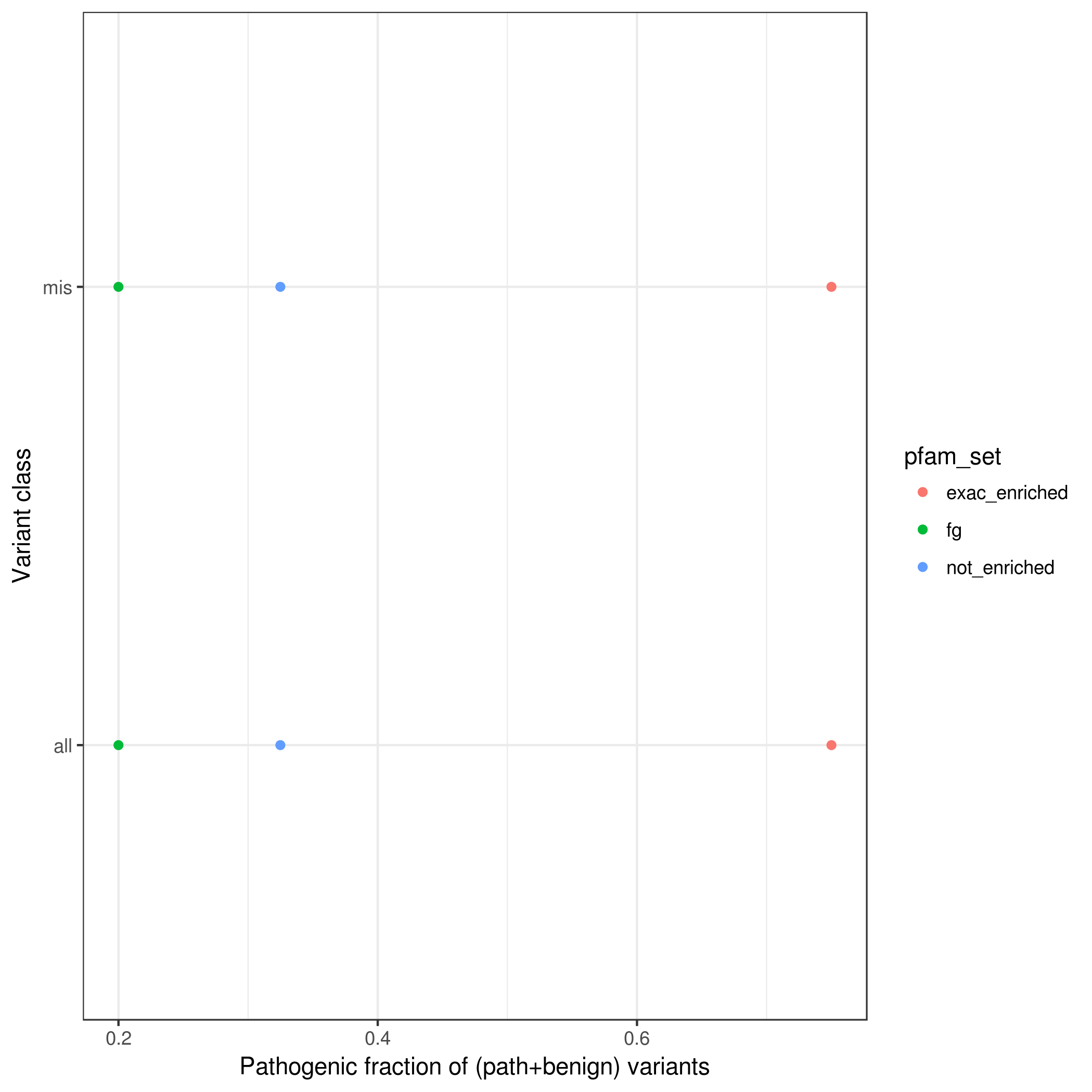

### rarem.var_counts_by_enrichment.pfam.mpcLow_0.mpcHigh_1.4.png

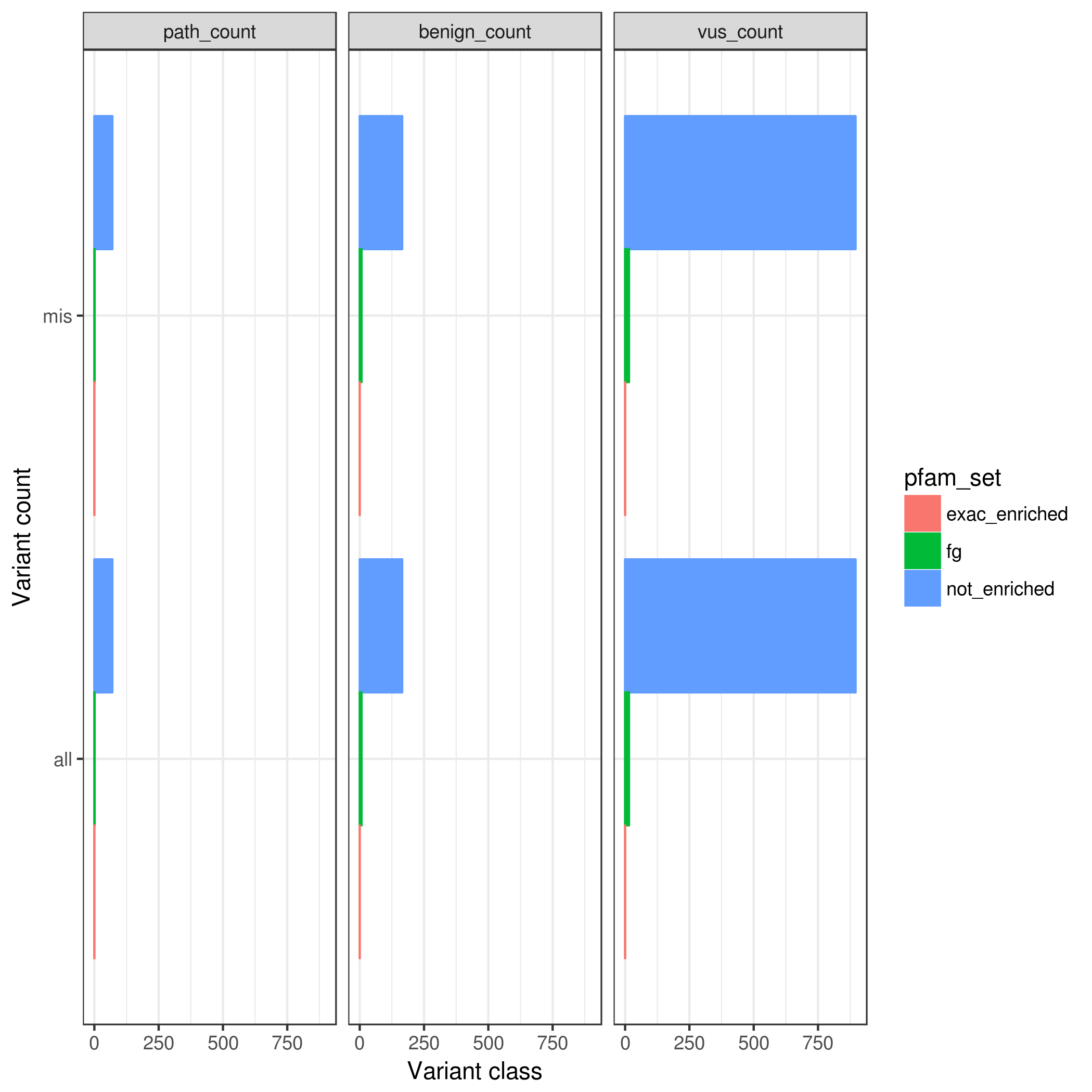

### rarem.var_counts_by_enrichment.pfamMerge.mpc_100.png

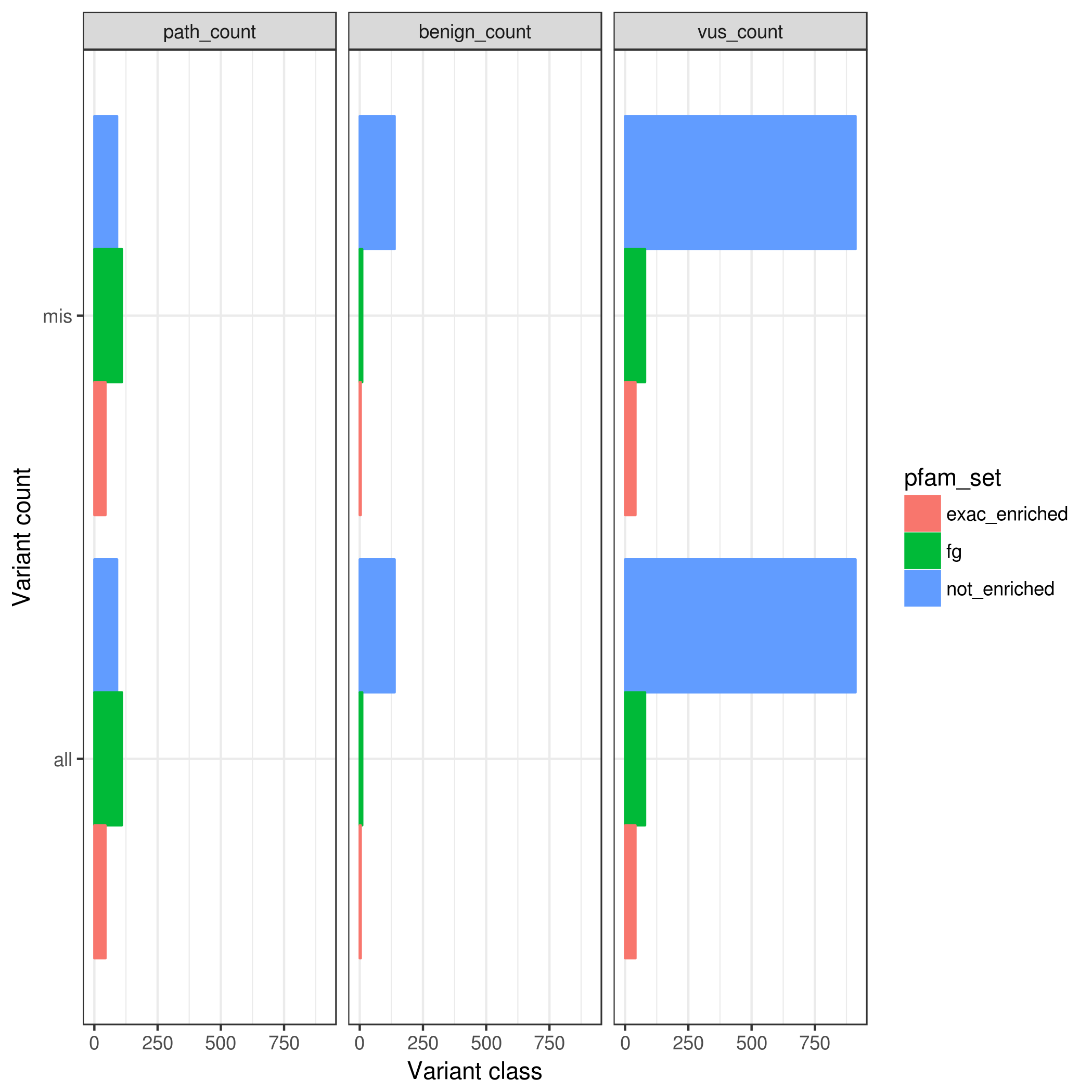

### rarem.var_counts_by_enrichment.pfamMerge.mpcLow_1.4.mpcHigh_100.png

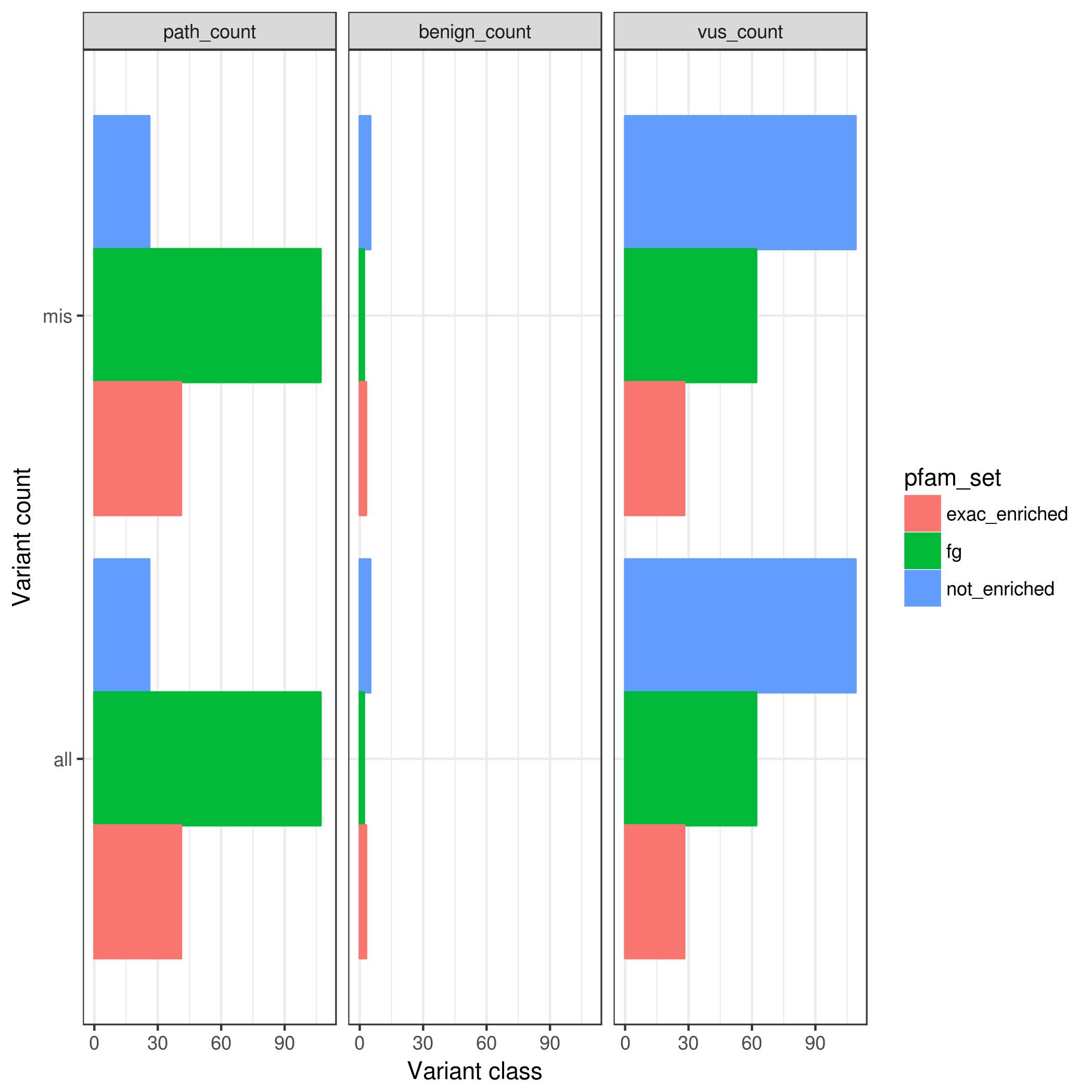

### roc.png

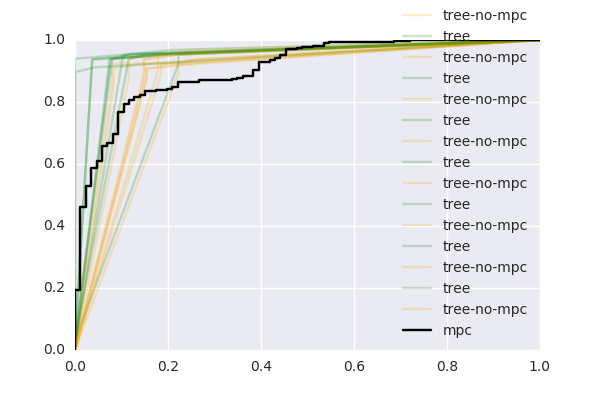
