## Supplementary figures and images for "Genetic variant pathogenicity prediction trained using disease-specific clinical sequencing datasets"

### CDKL5.clinvar.lolly.png

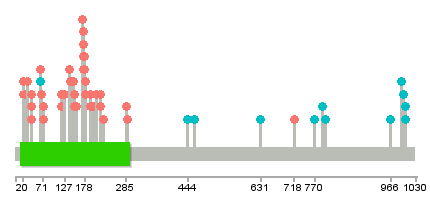

### CDKL5.denovo_db.lolly.png

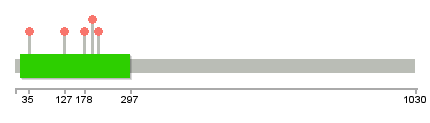

### clinvar_KCNQ2_eval.png

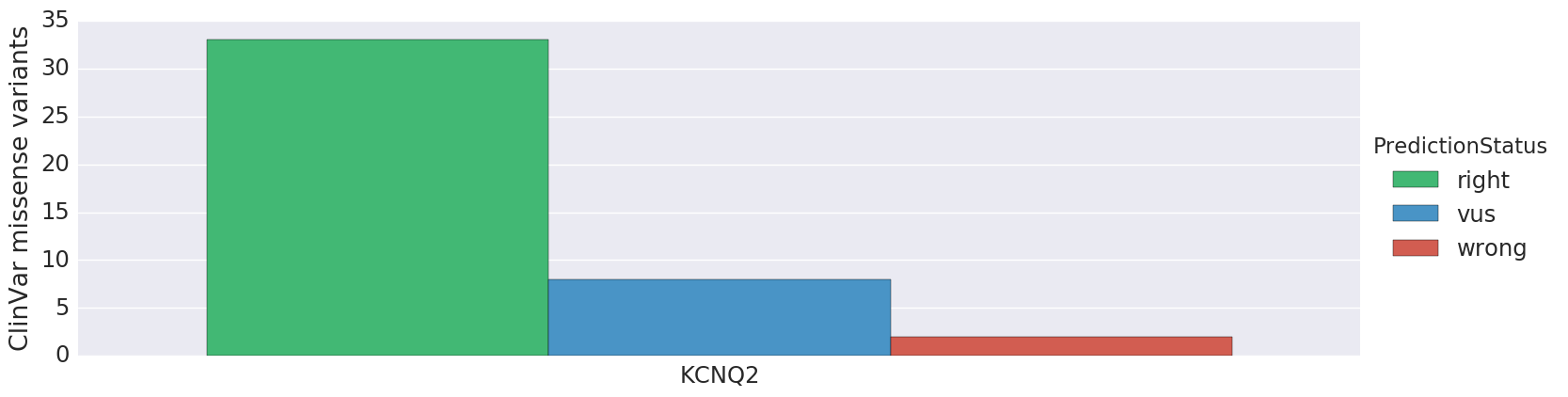

### clinvar_KCNQ3_eval.png

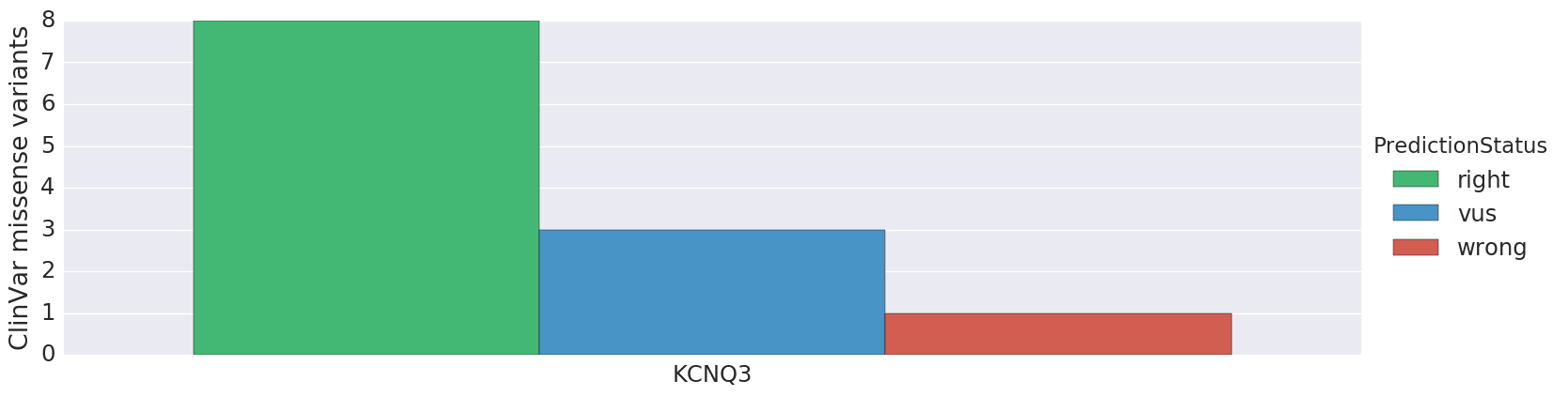

### clinvar_roc.png

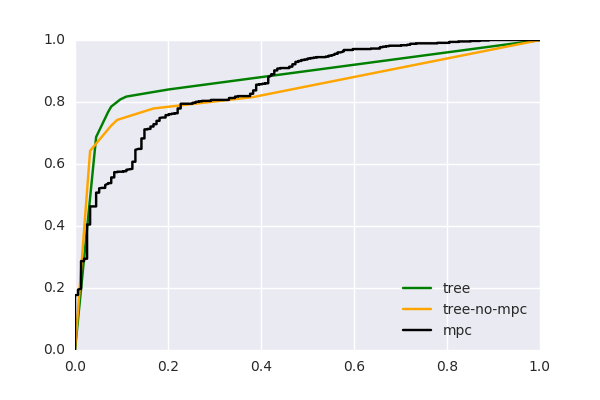

### clinvar_SCN1A_roc.png

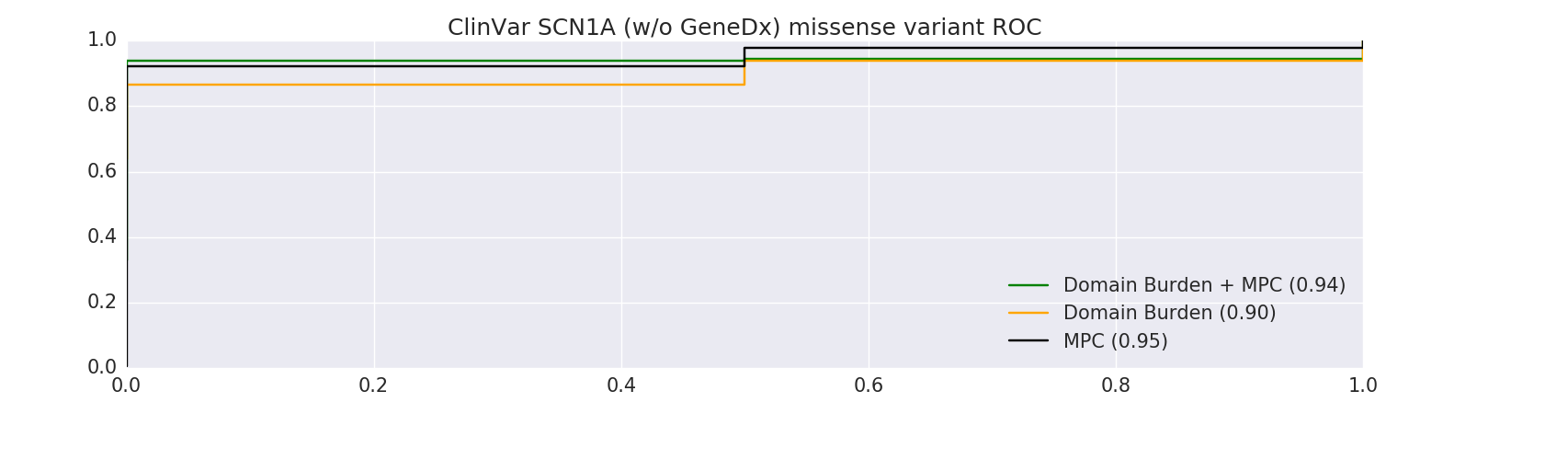

### clinvar_SCN2A_eval.png

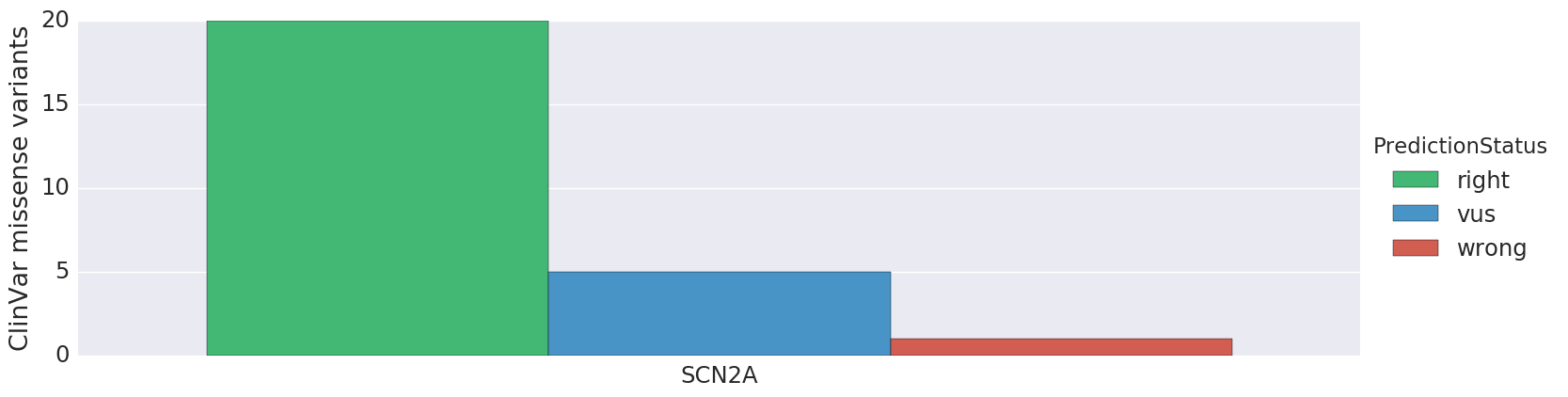

### clinvar_SPTAN1_eval.png

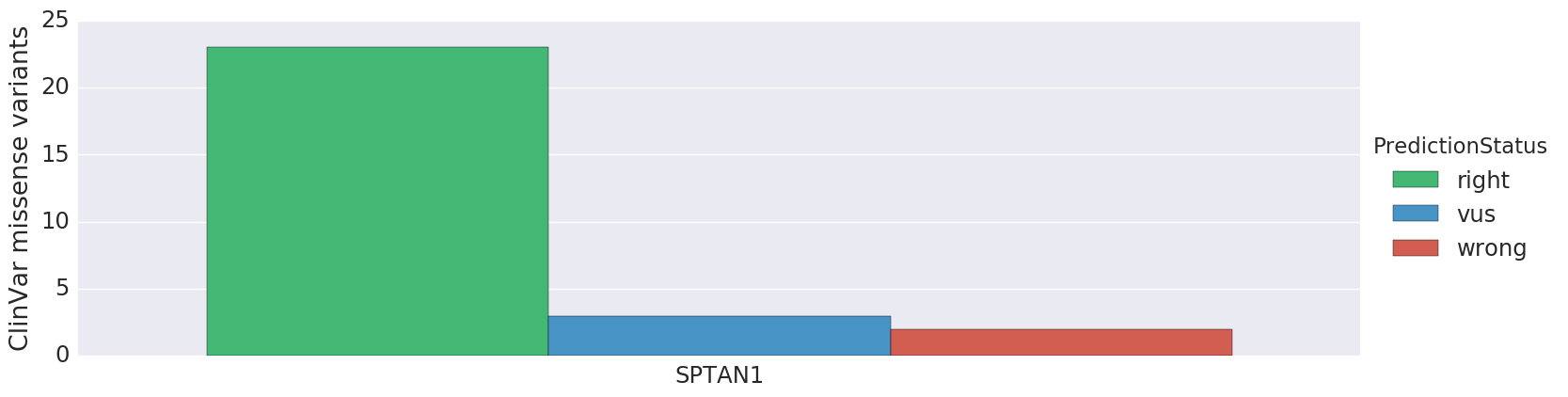

### clinvar_SPTAN1_roc.png

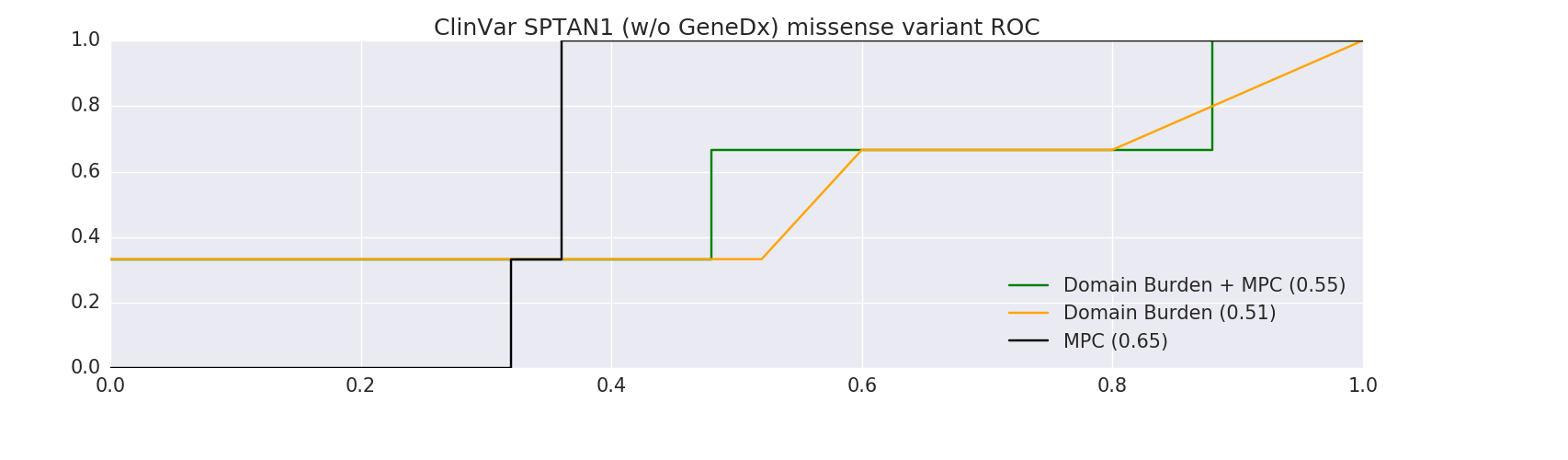

### clinvar_STXBP1_eval.png

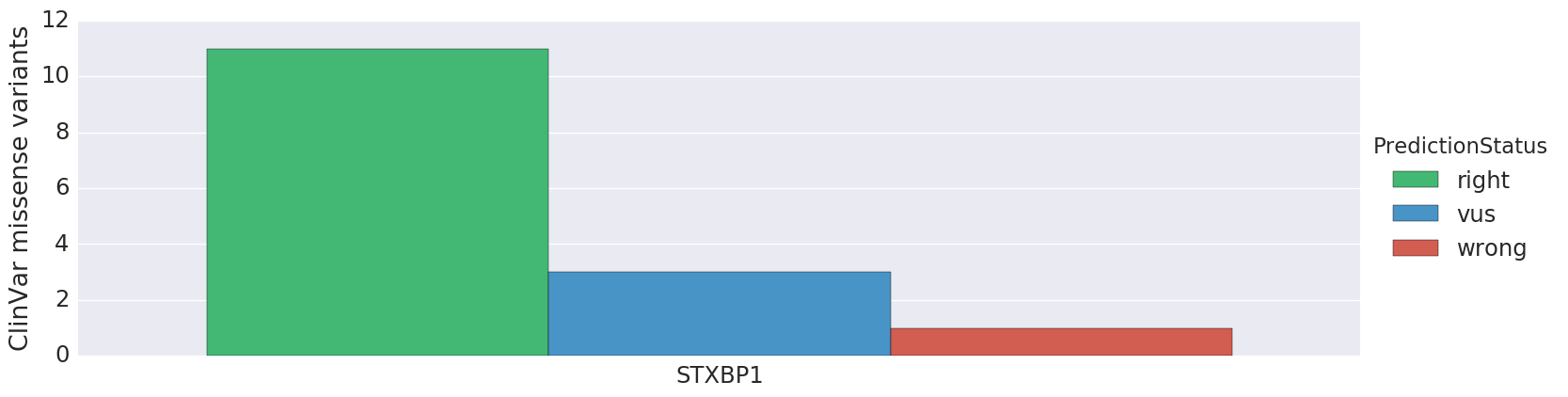

### clinvar_TSC1_roc.png

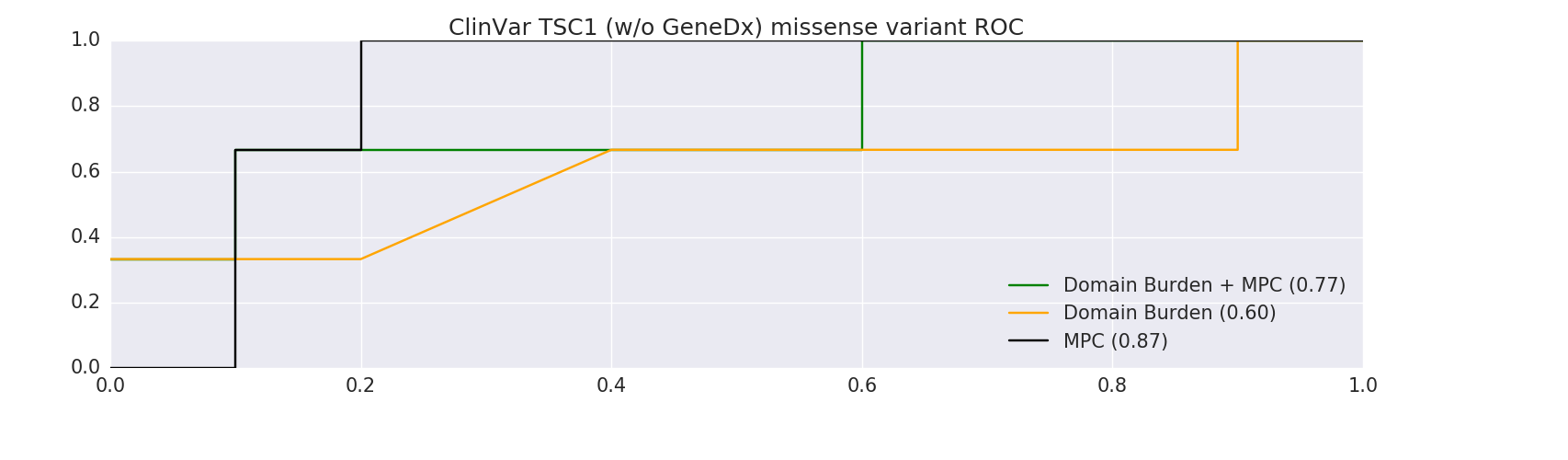

### denovo_burden_gene_eval.png

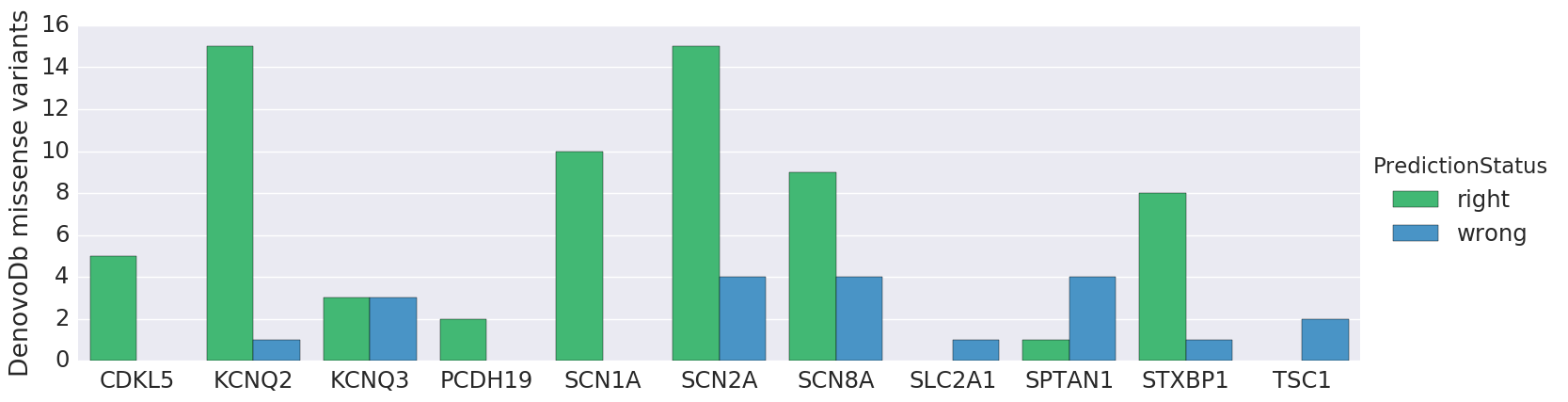

### fig2a_featureImportance.pdf

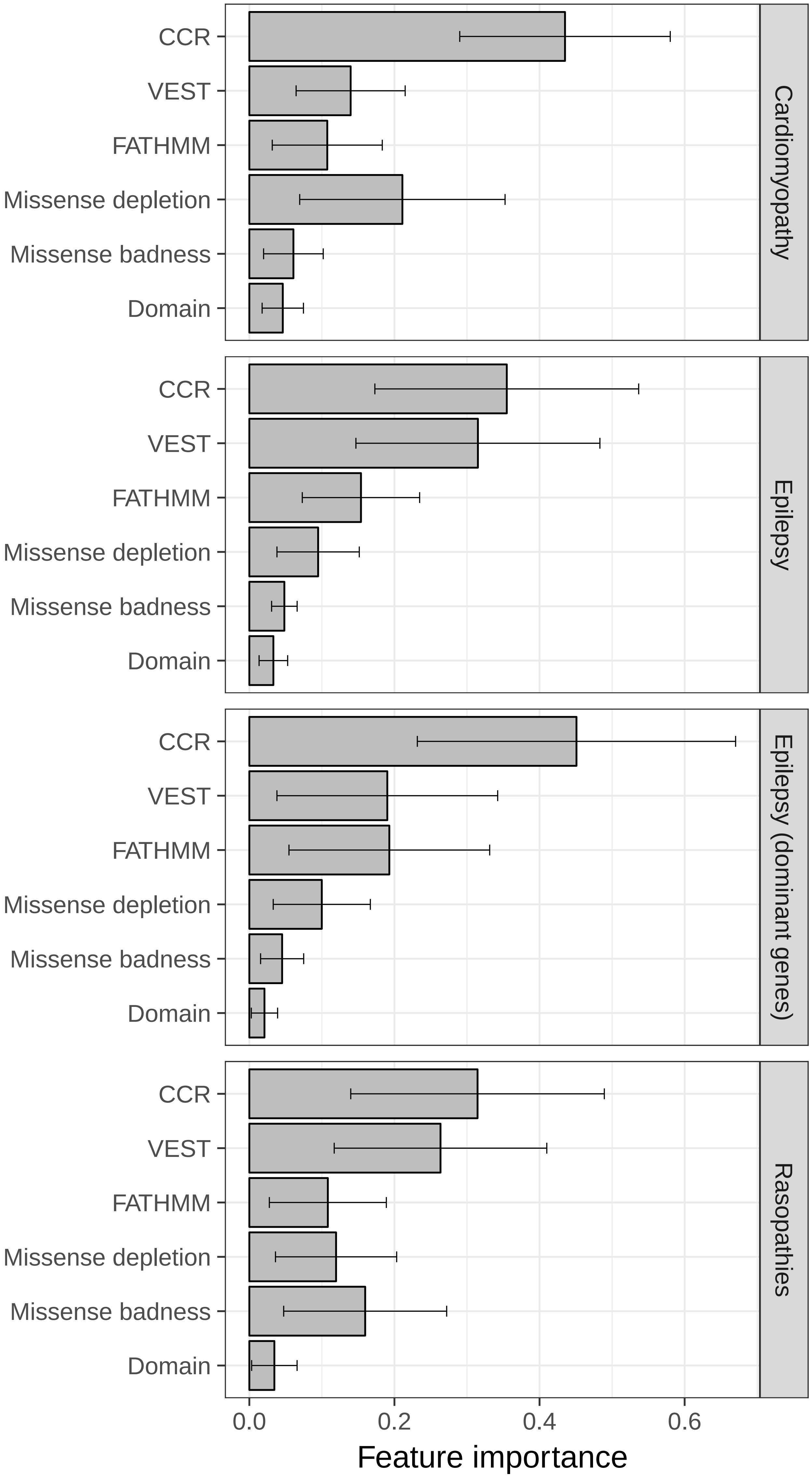

### fig2b_featureCor.pdf

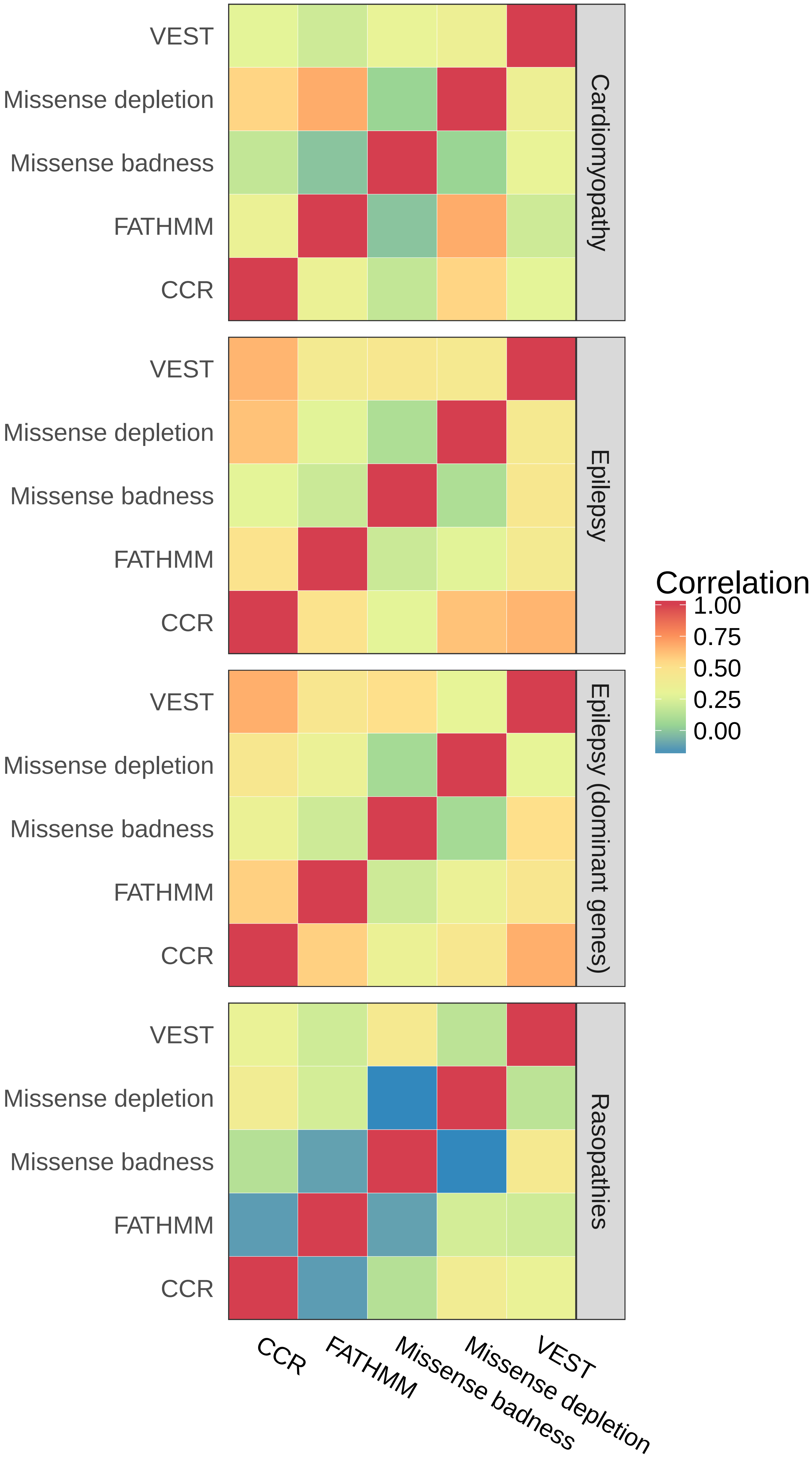

### fig6_byGene.pdf

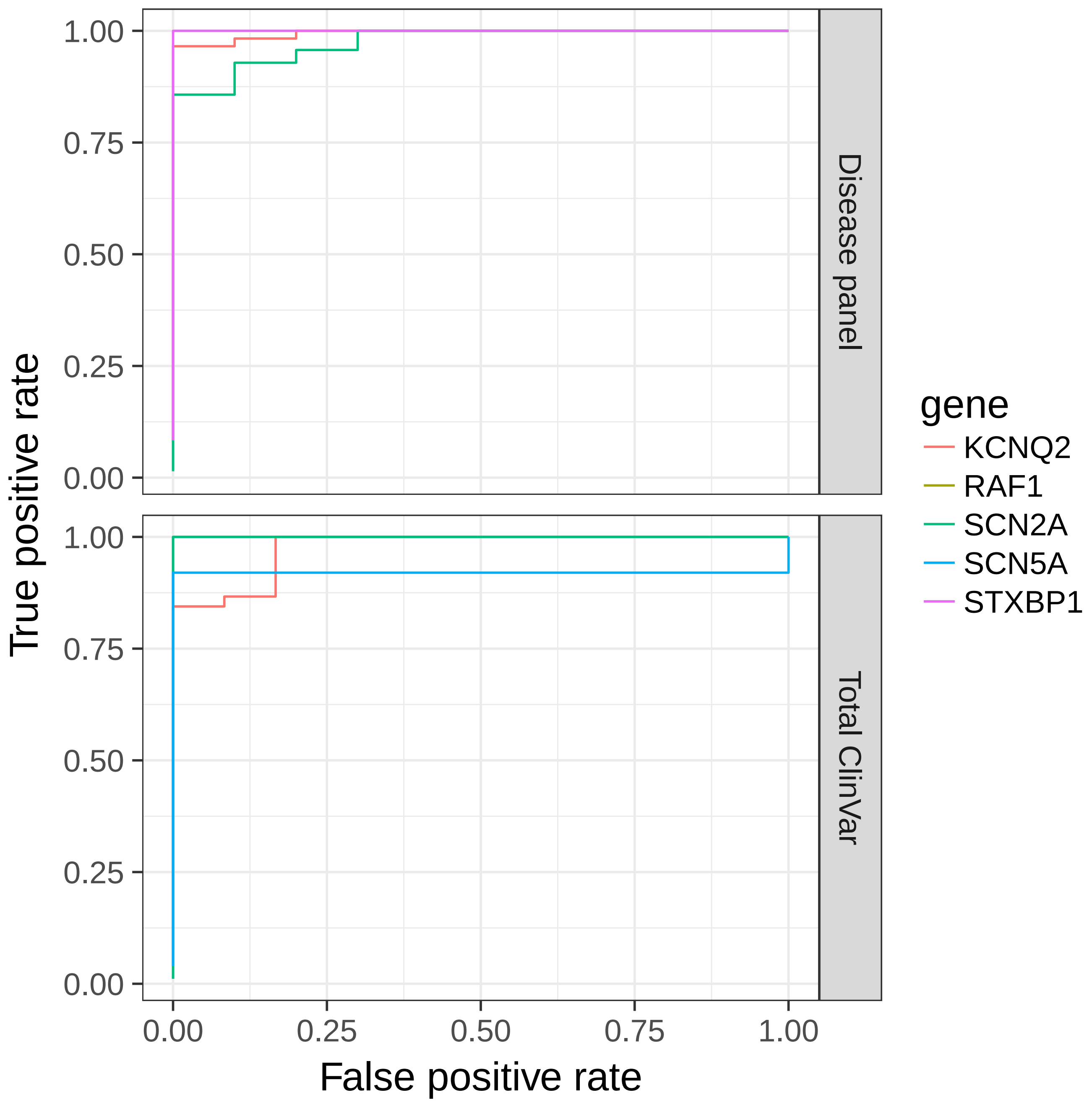

### global.byDisease.png

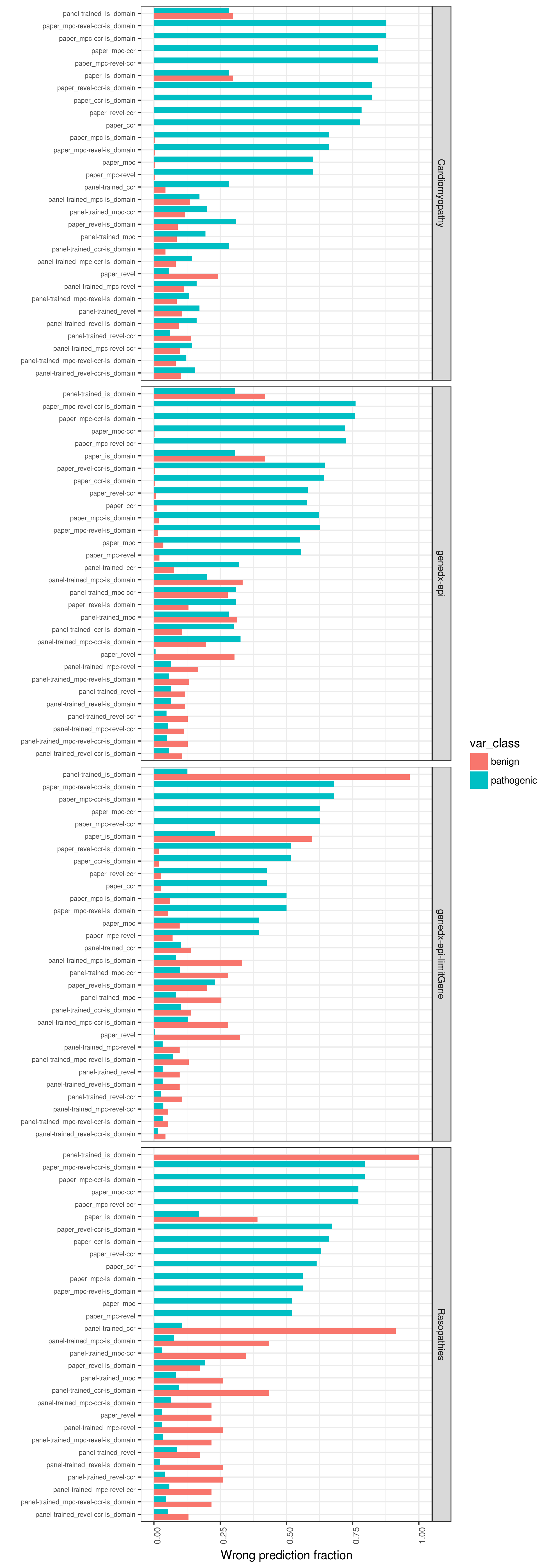

### global.eval_clinvar.totWrong.png

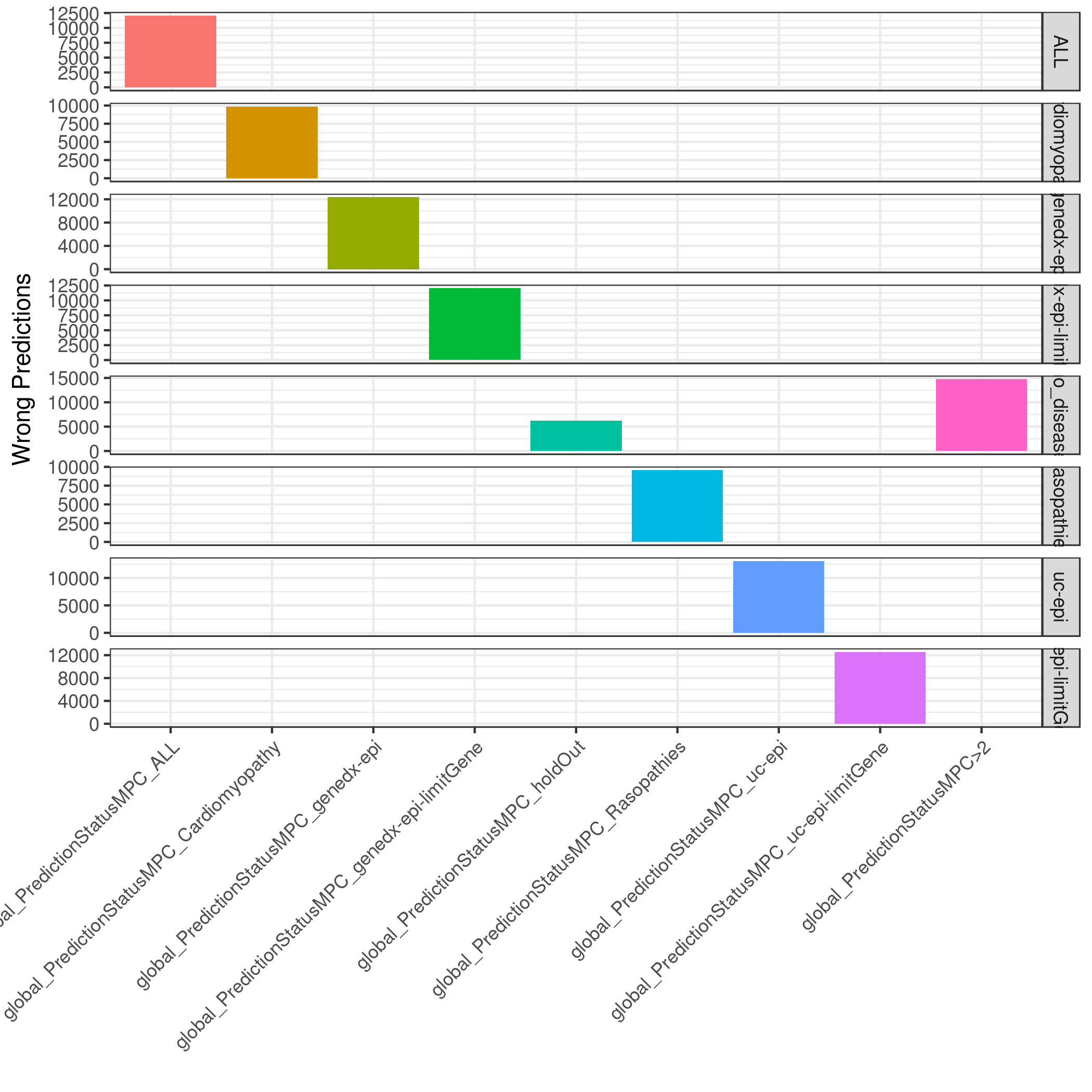

### global.eval_clinvar_exp.totWrong.png

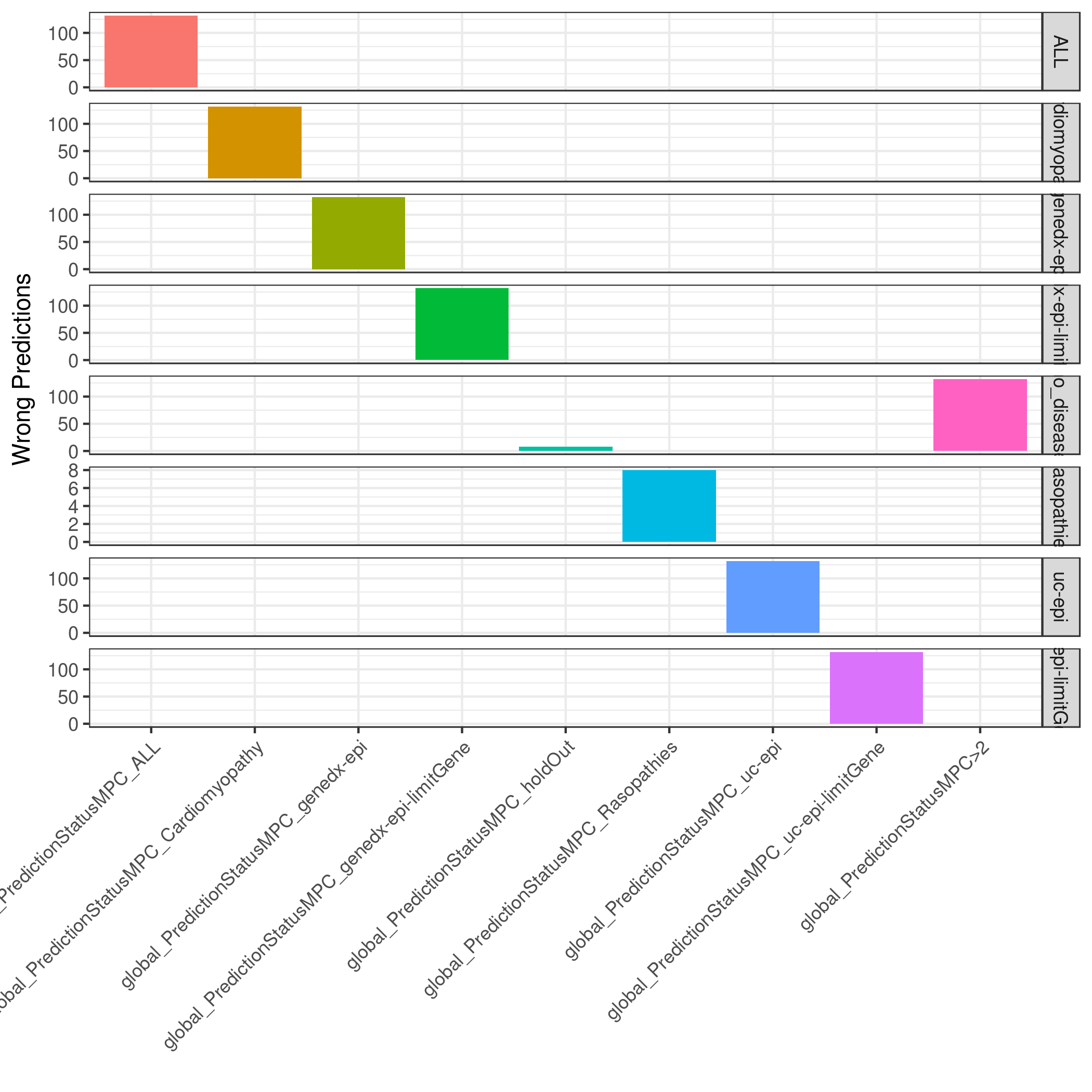

### global.eval_clinvar_mult.totWrong.png

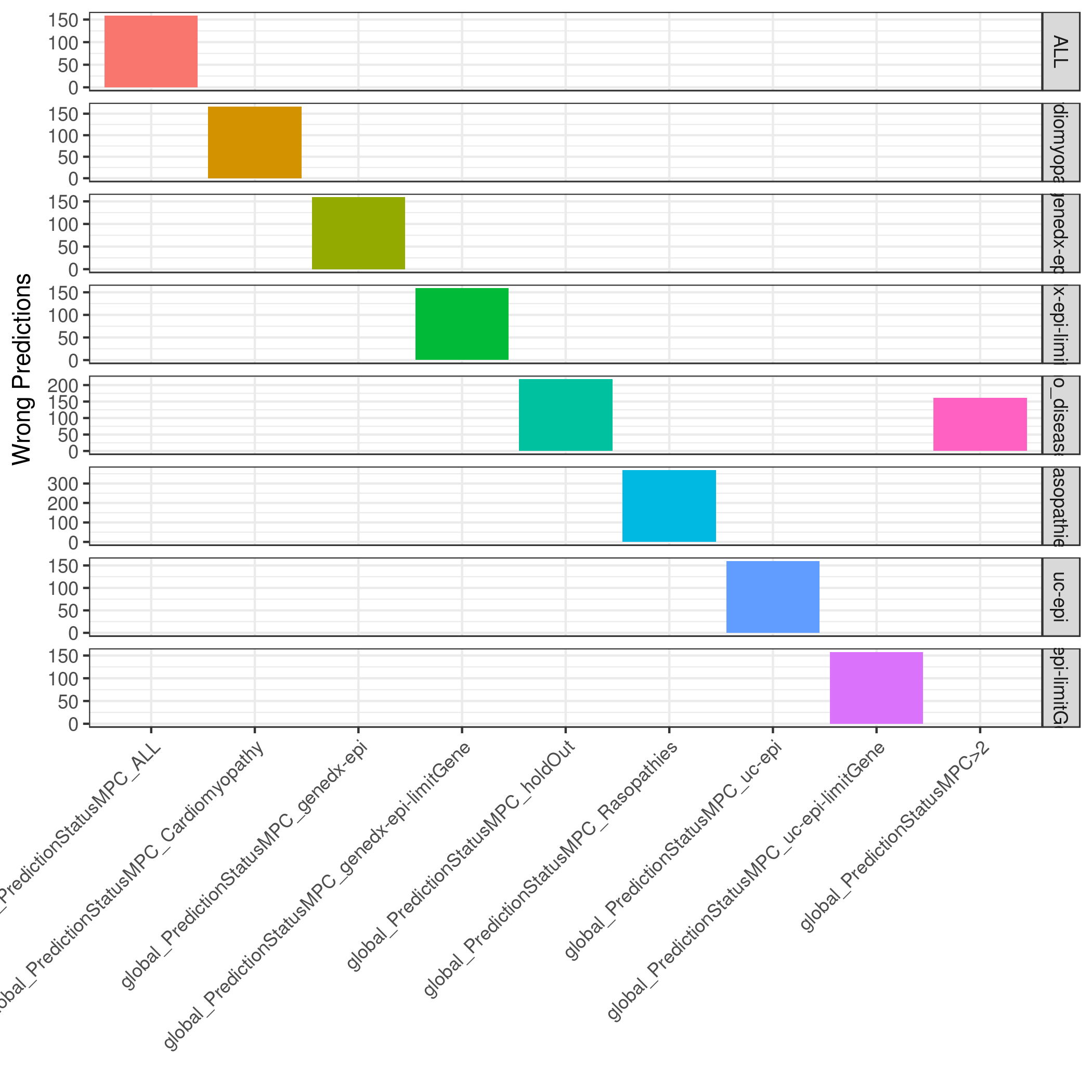

### global.eval_clinvar_single.totWrong.png

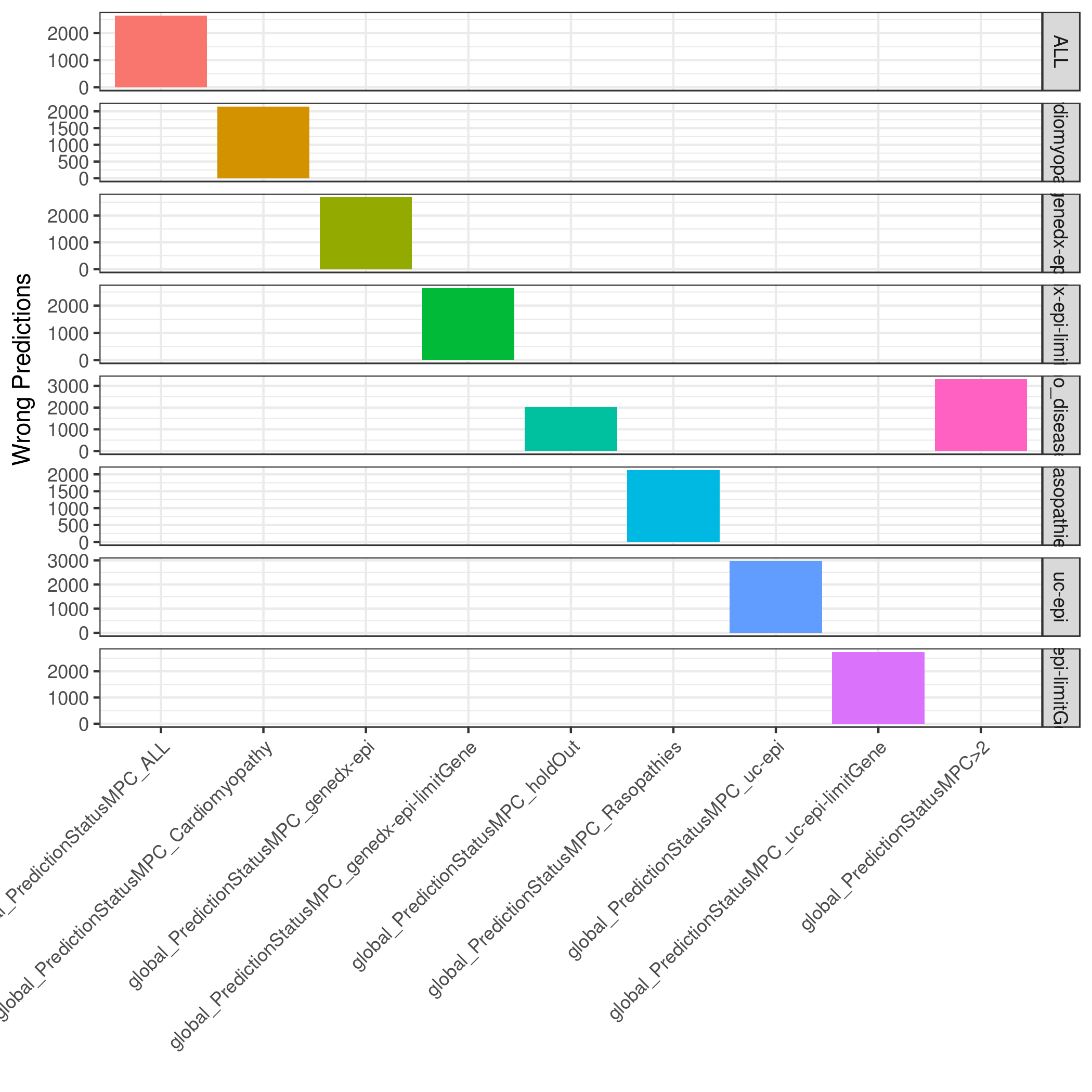

### global.eval_denovo.totWrong.png

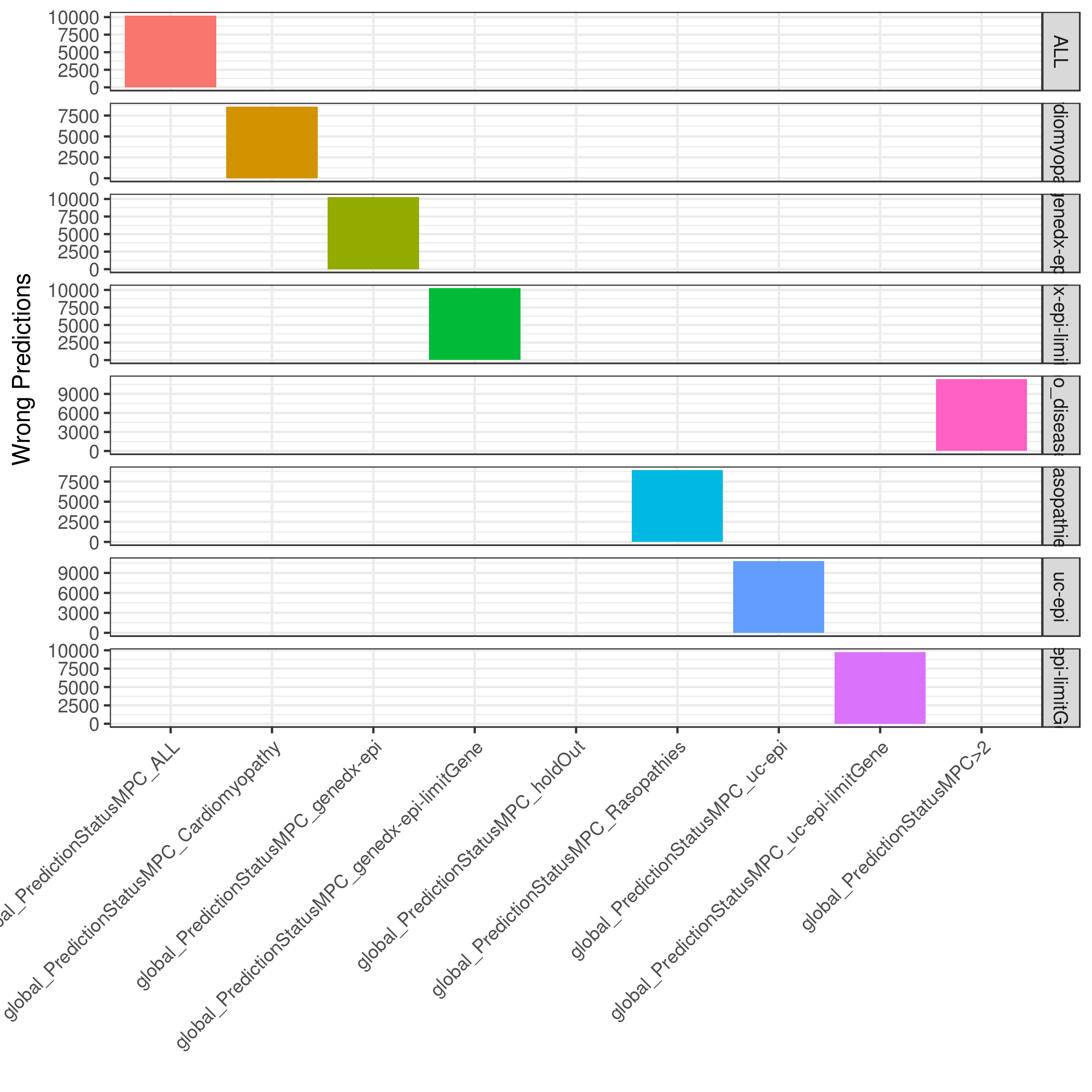

### global.eval_panel.ccr.totWrong.png

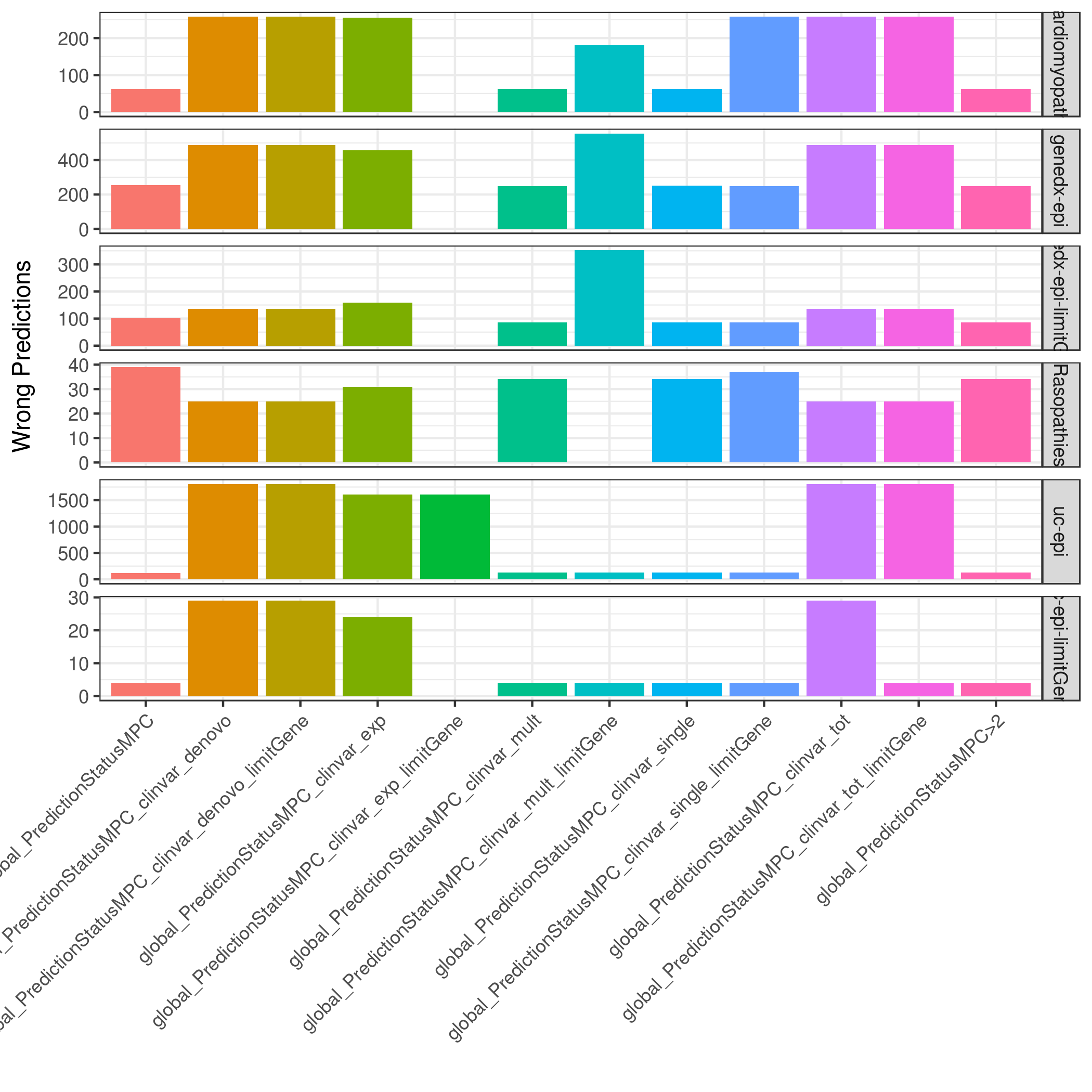

### global.eval_panel.mpc-ccr.totWrong.png

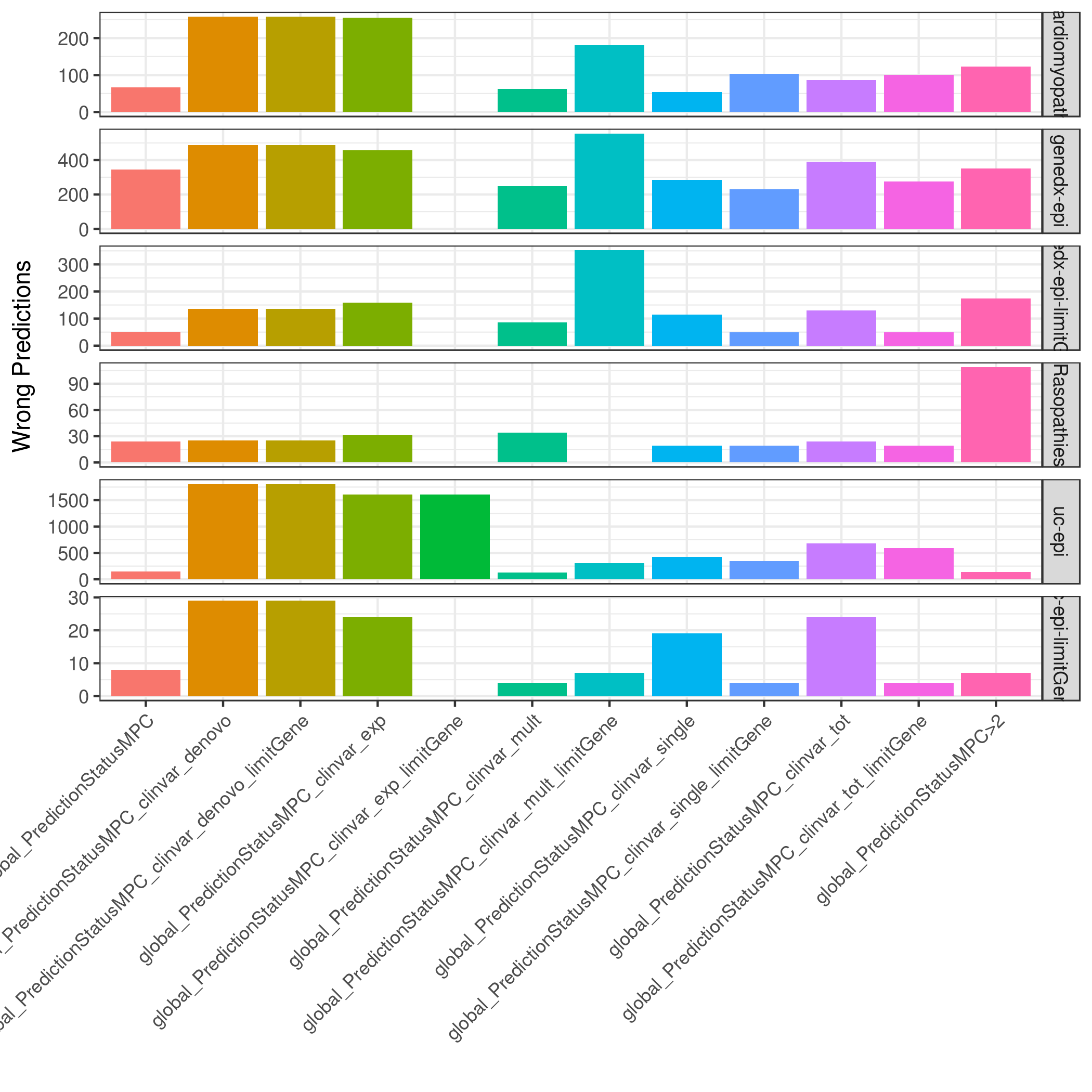

### global.eval_panel.mpc-revel-ccr.totWrong.png

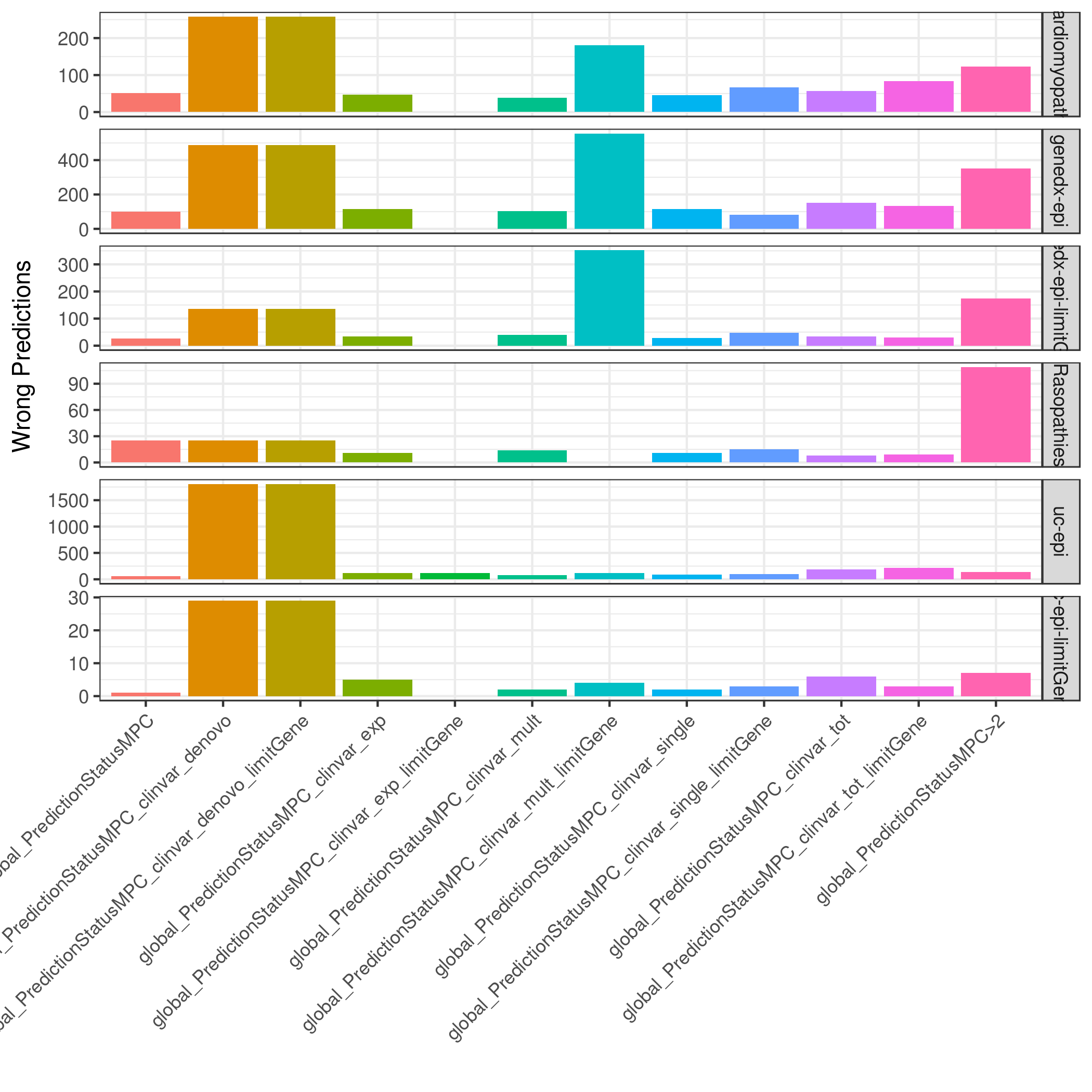

### global.eval_panel.mpc-revel.totWrong.png

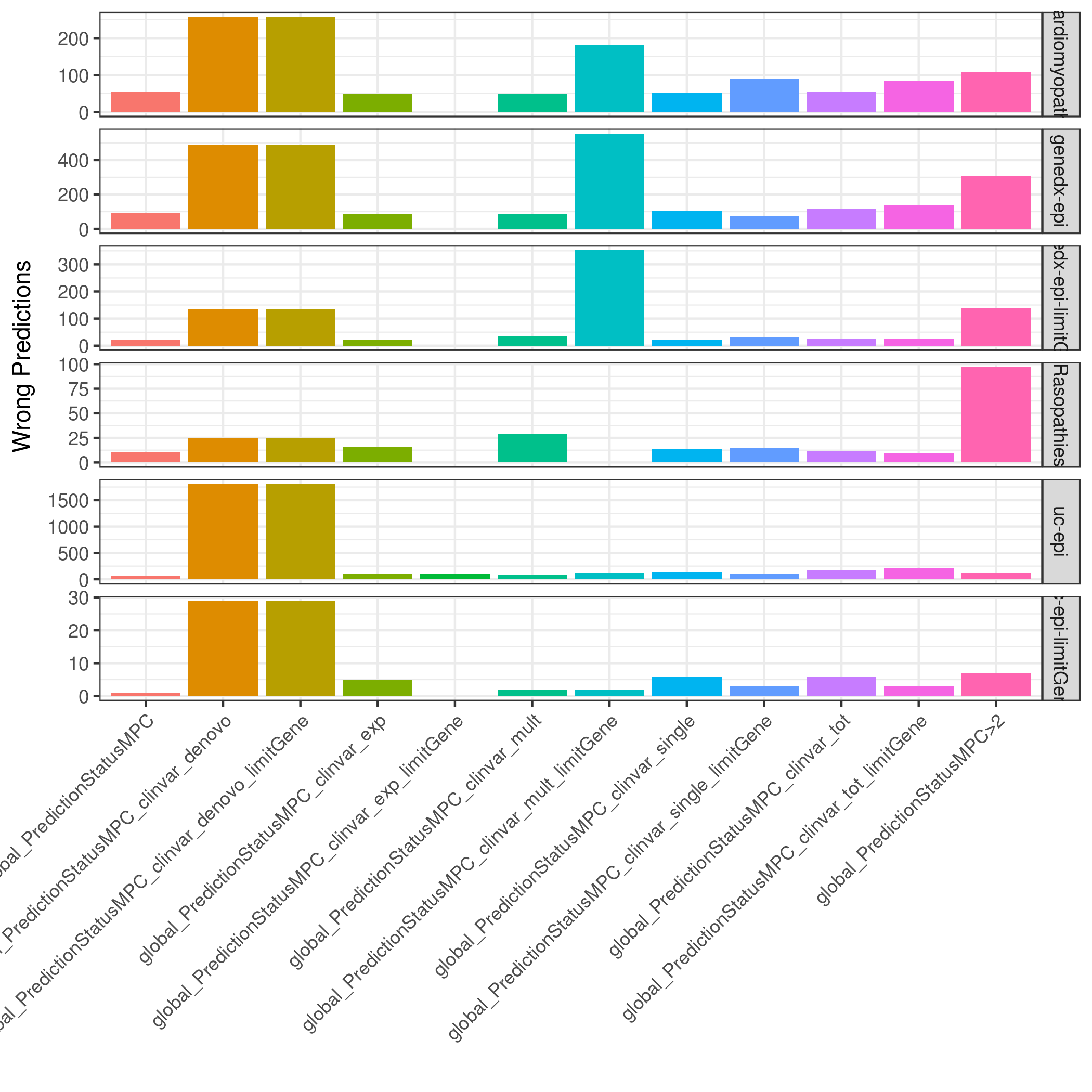

### global.eval_panel.mpc.totWrong.png

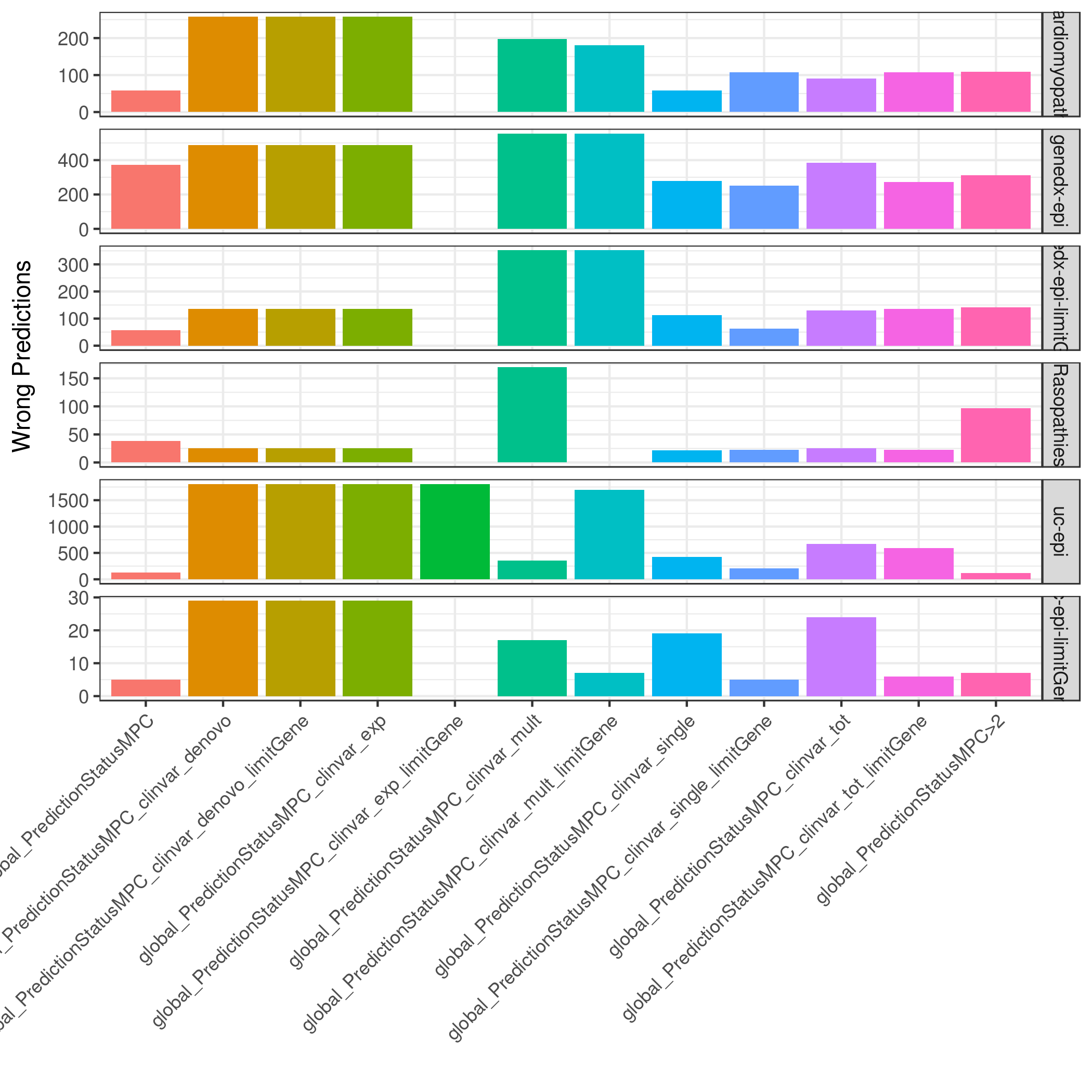

### global.eval_panel.revel-ccr.totWrong.png

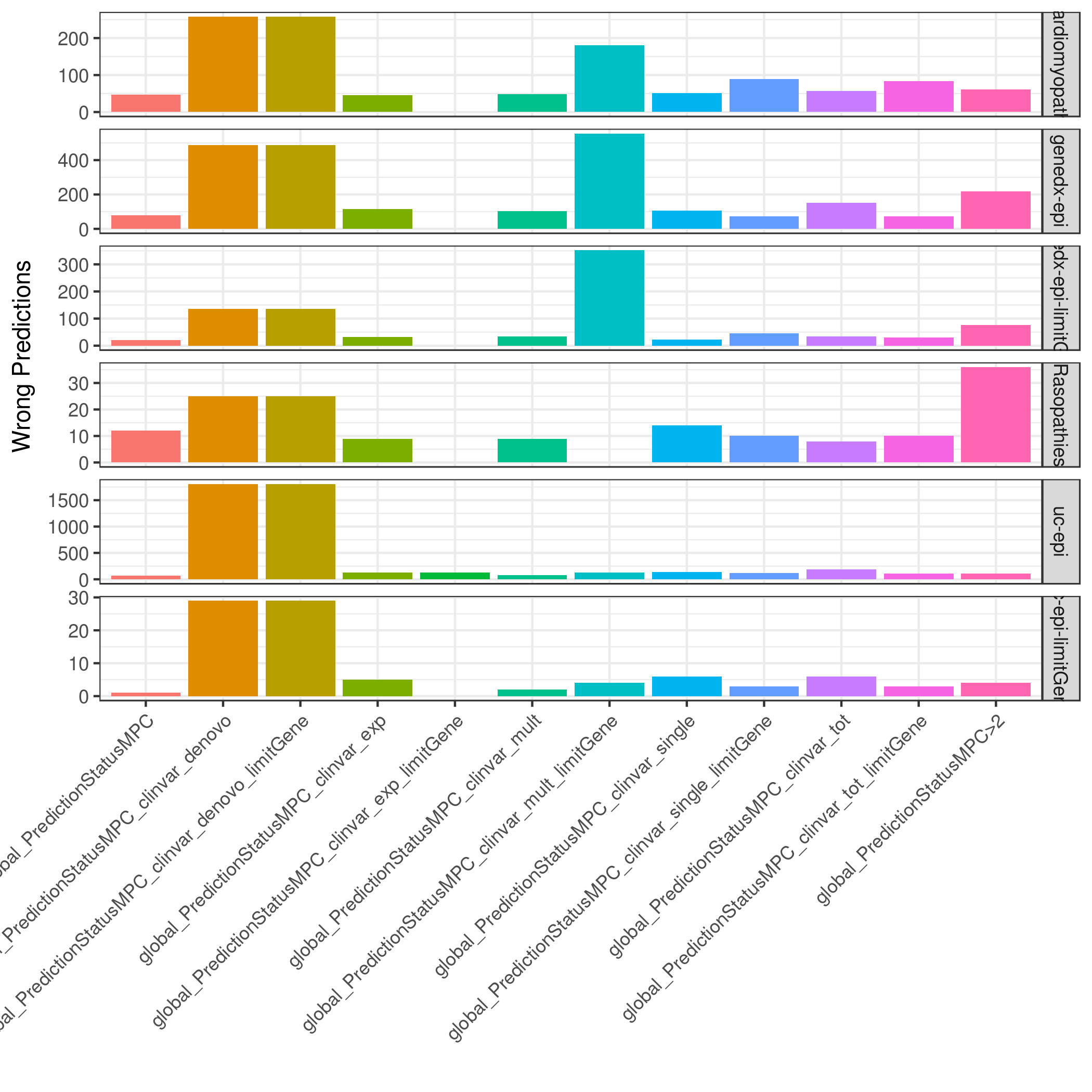

### global.eval_panel.revel.totWrong.png

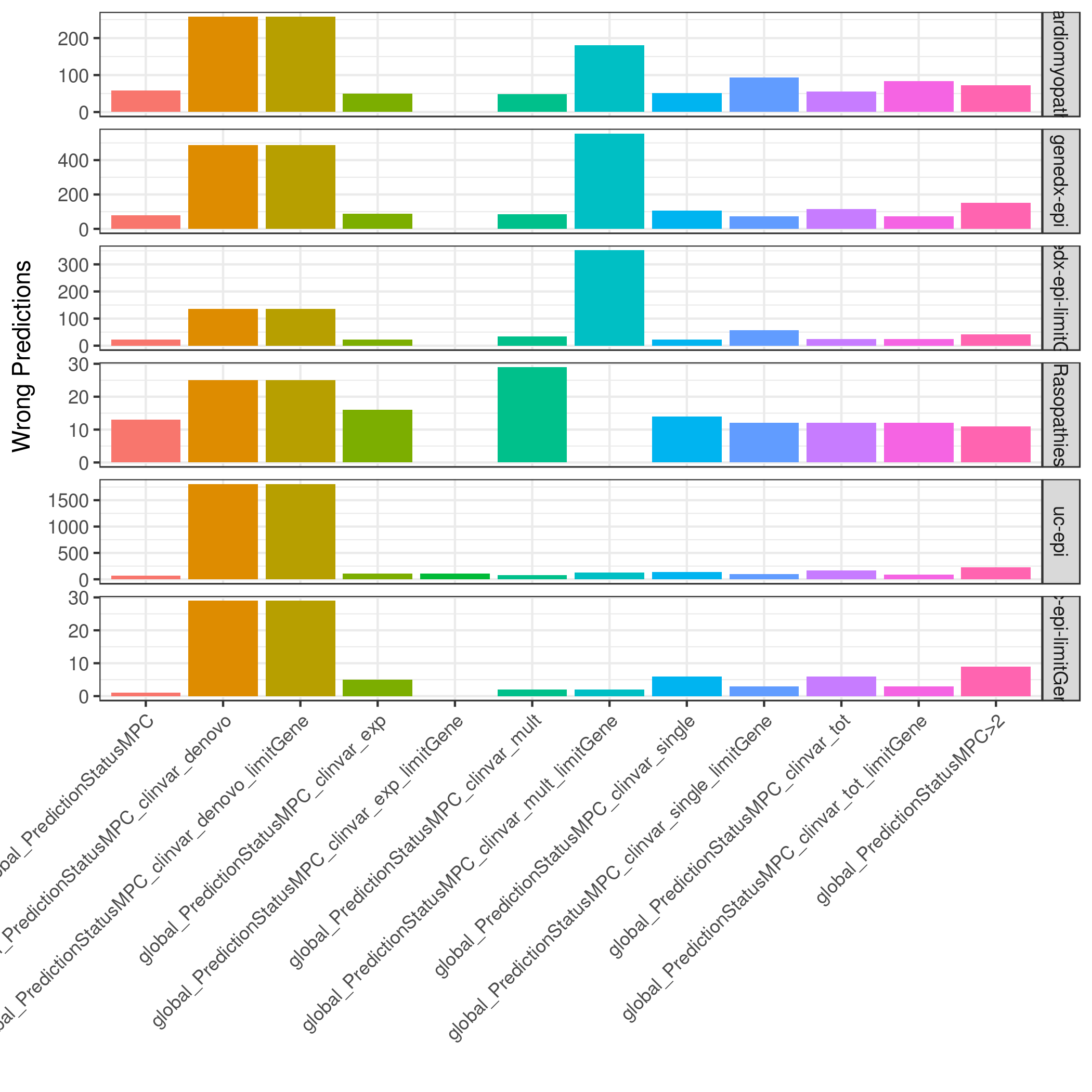

### global.eval_panel.totWrong.png

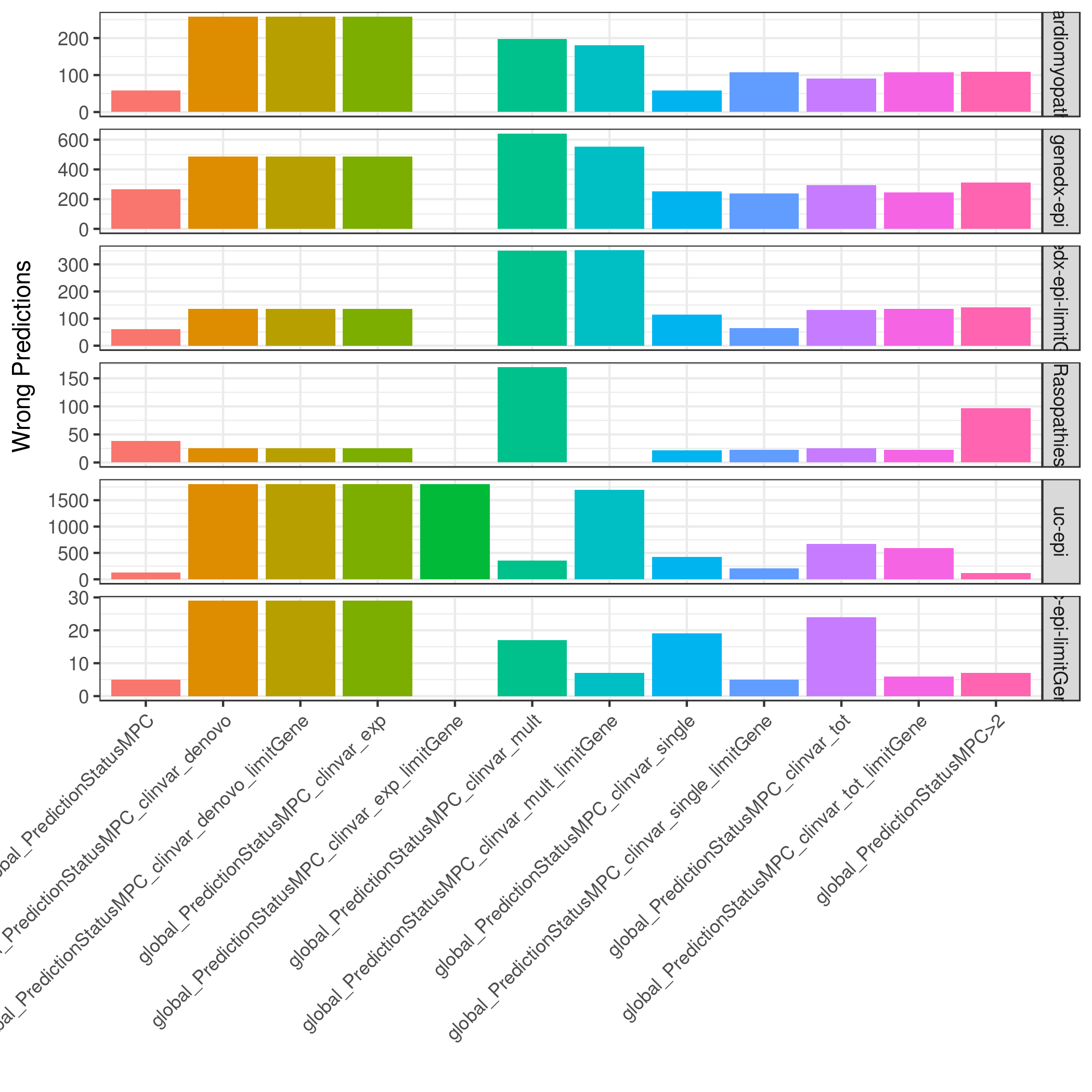

### global.mpc-revel.byDisease.png

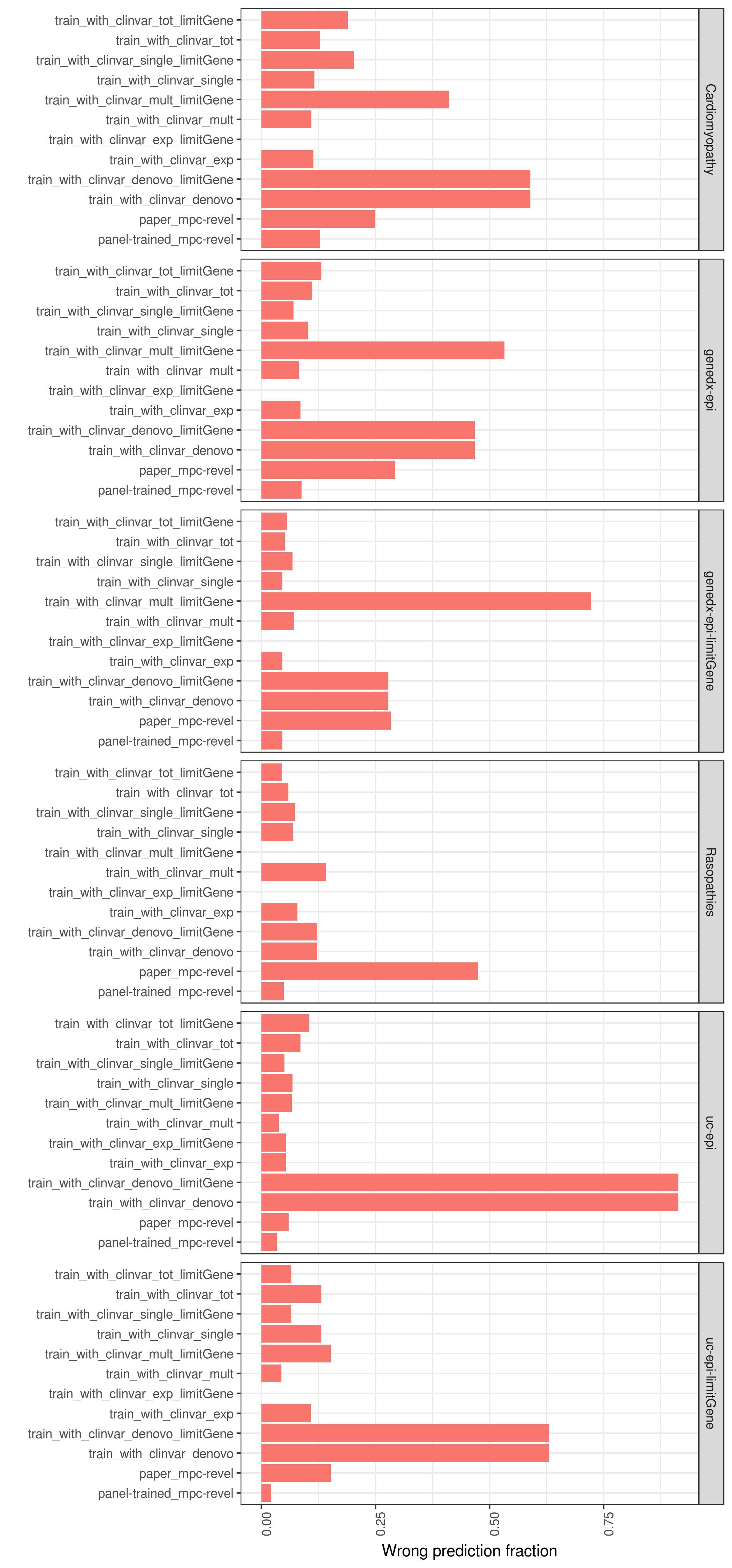

### global.mpc.byDisease.png

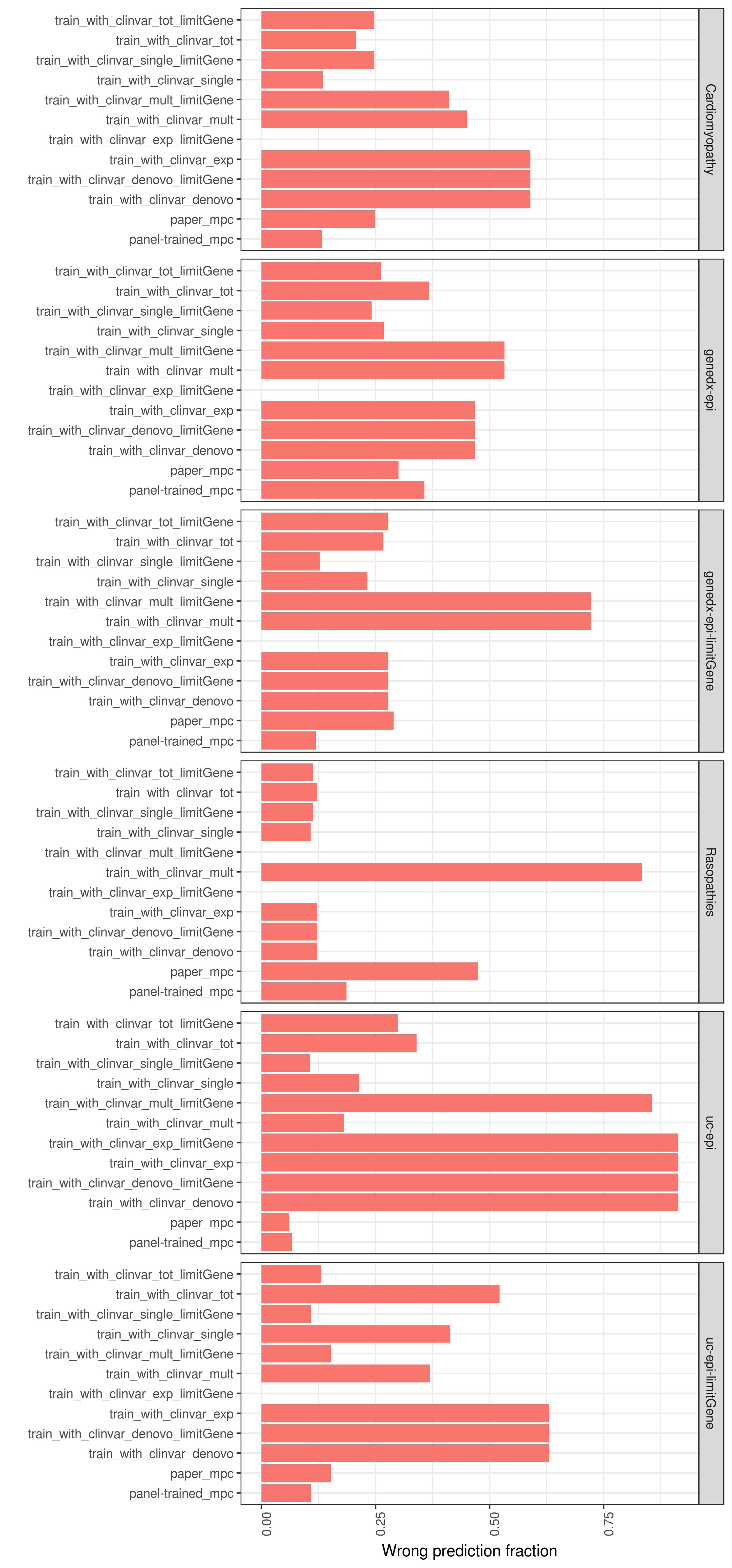

### global.other.ccr.totWrong.png

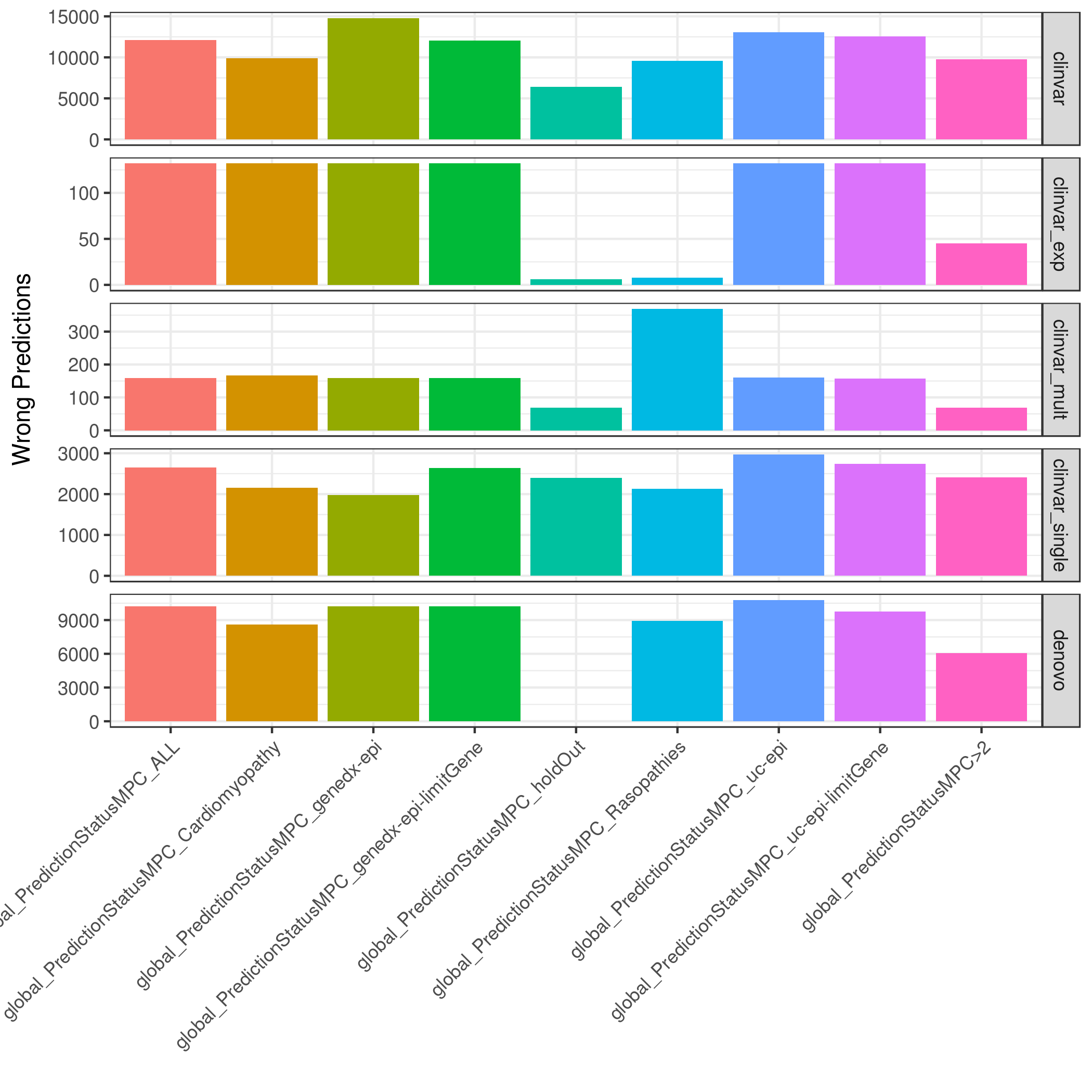

### global.other.mpc-ccr.totWrong.png

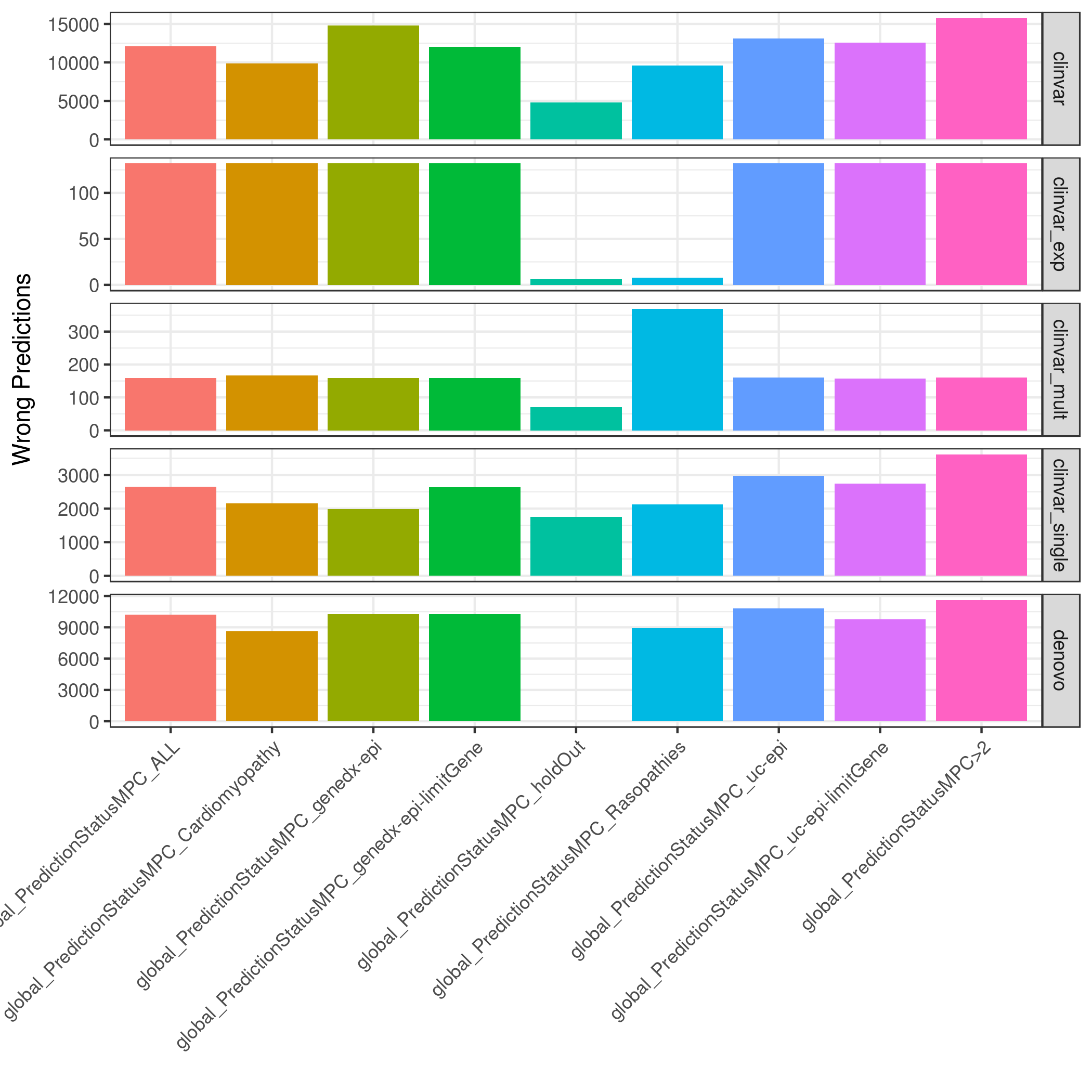

### global.other.mpc-revel-ccr.totWrong.png

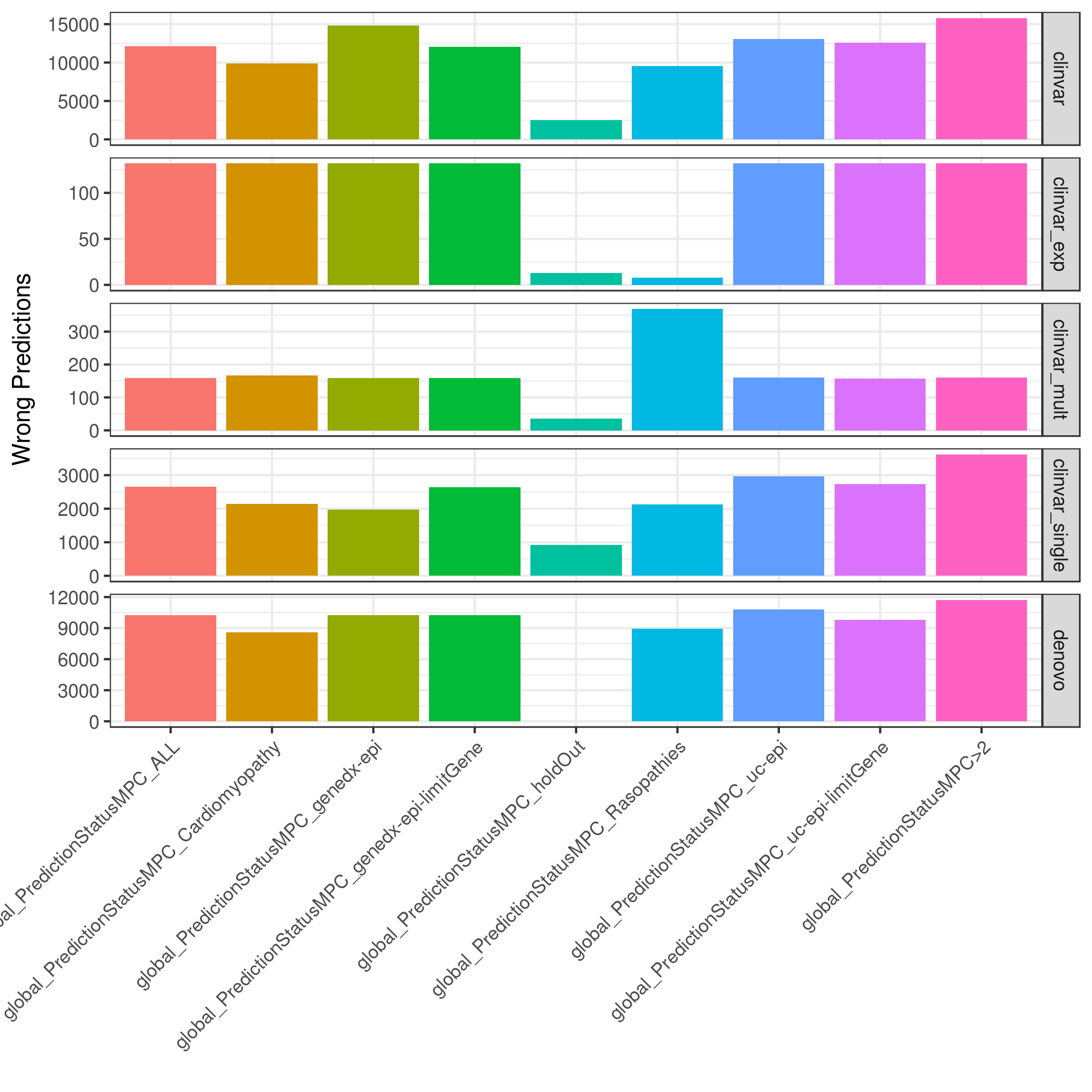

### global.other.mpc.totWrong.png

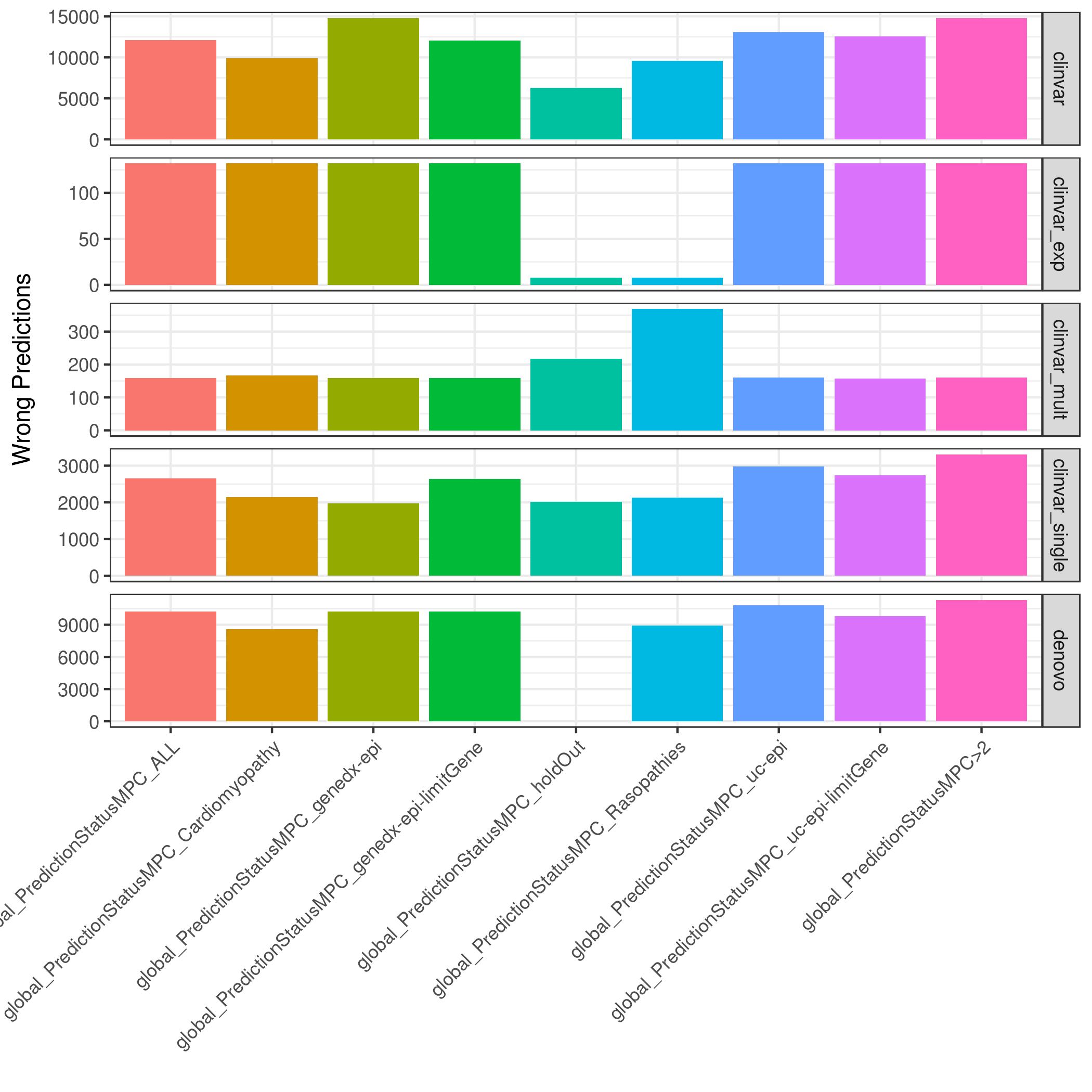

### global.other.revel-ccr.totWrong.png

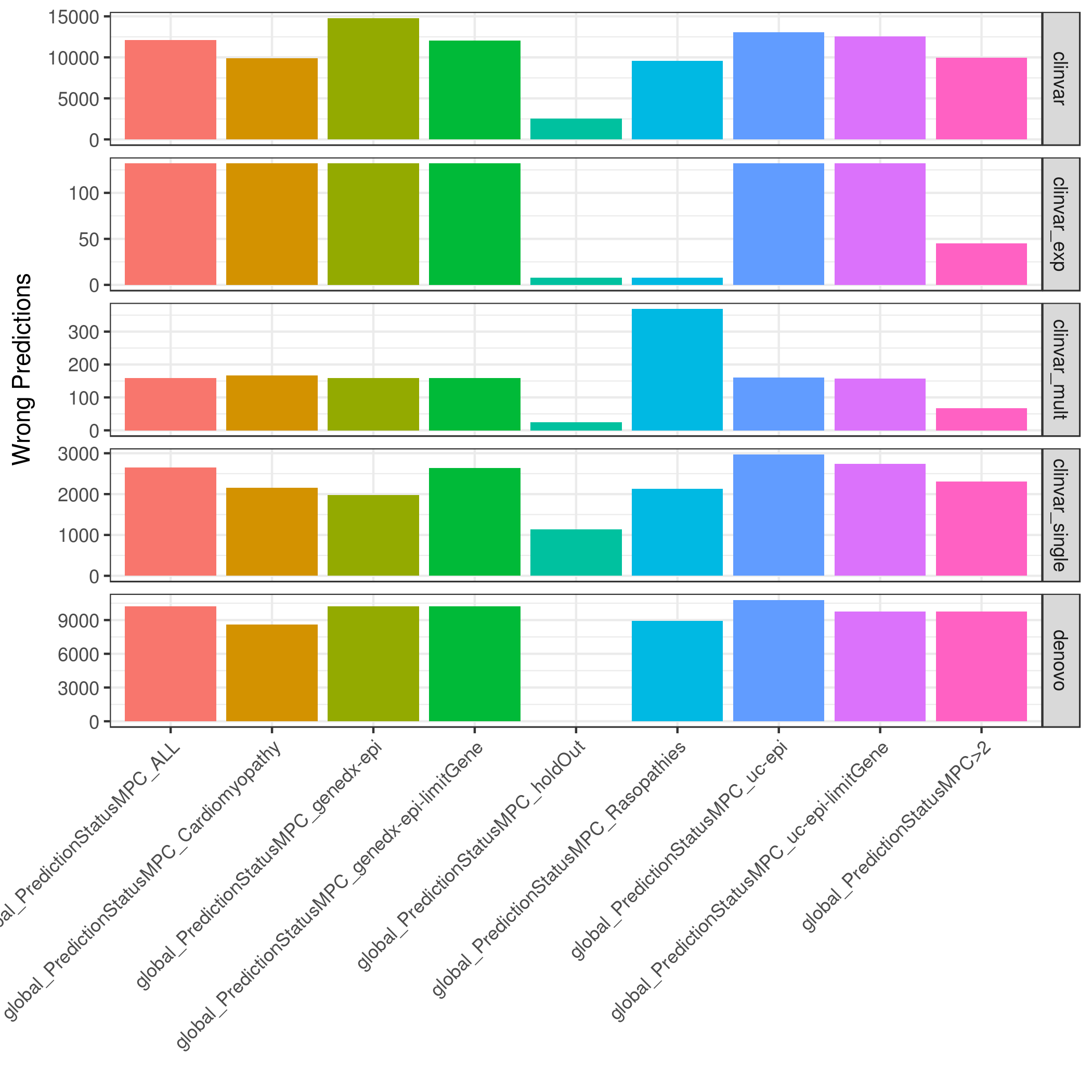

### global.other.revel-mpc.totWrong.png

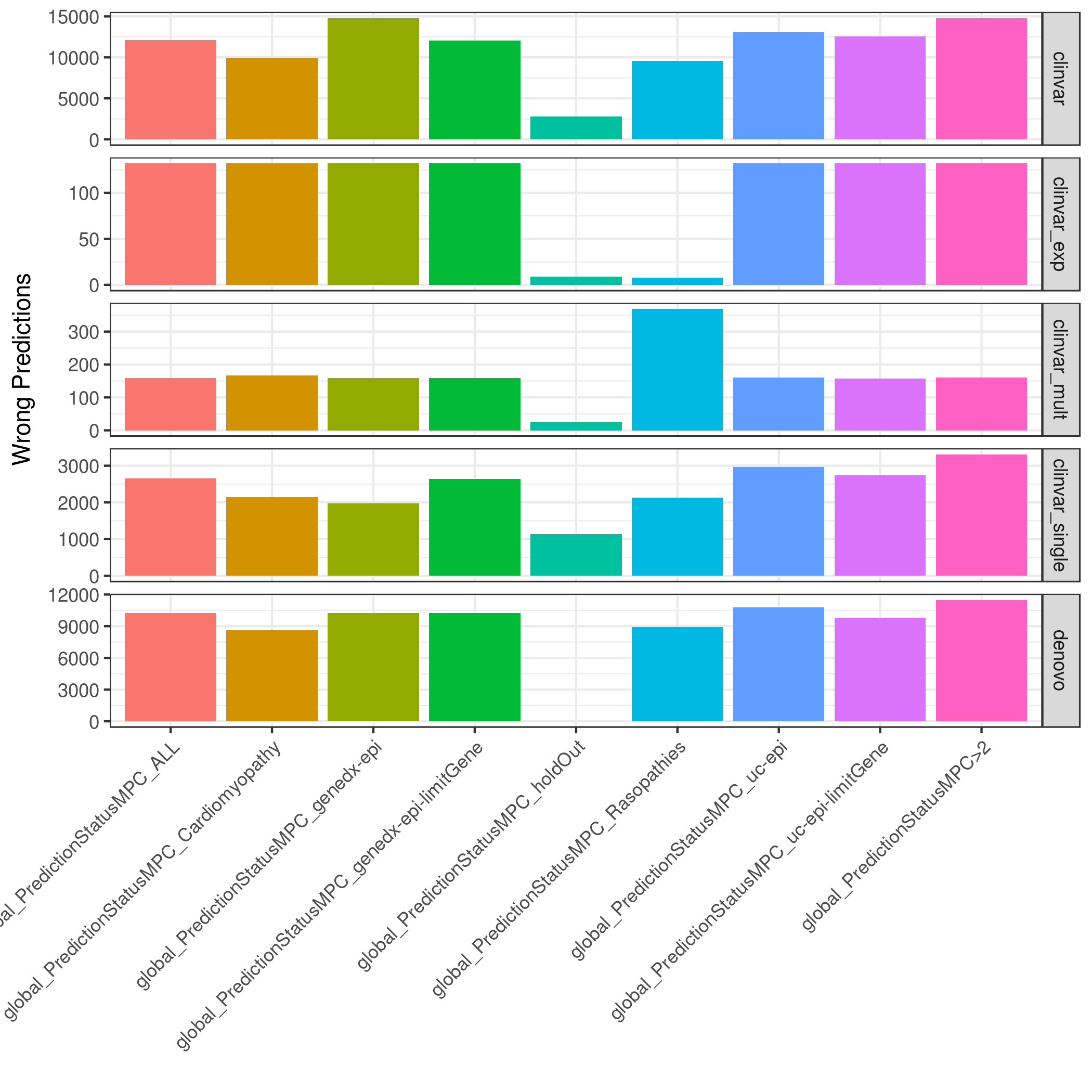

### global.other.revel.totWrong.png

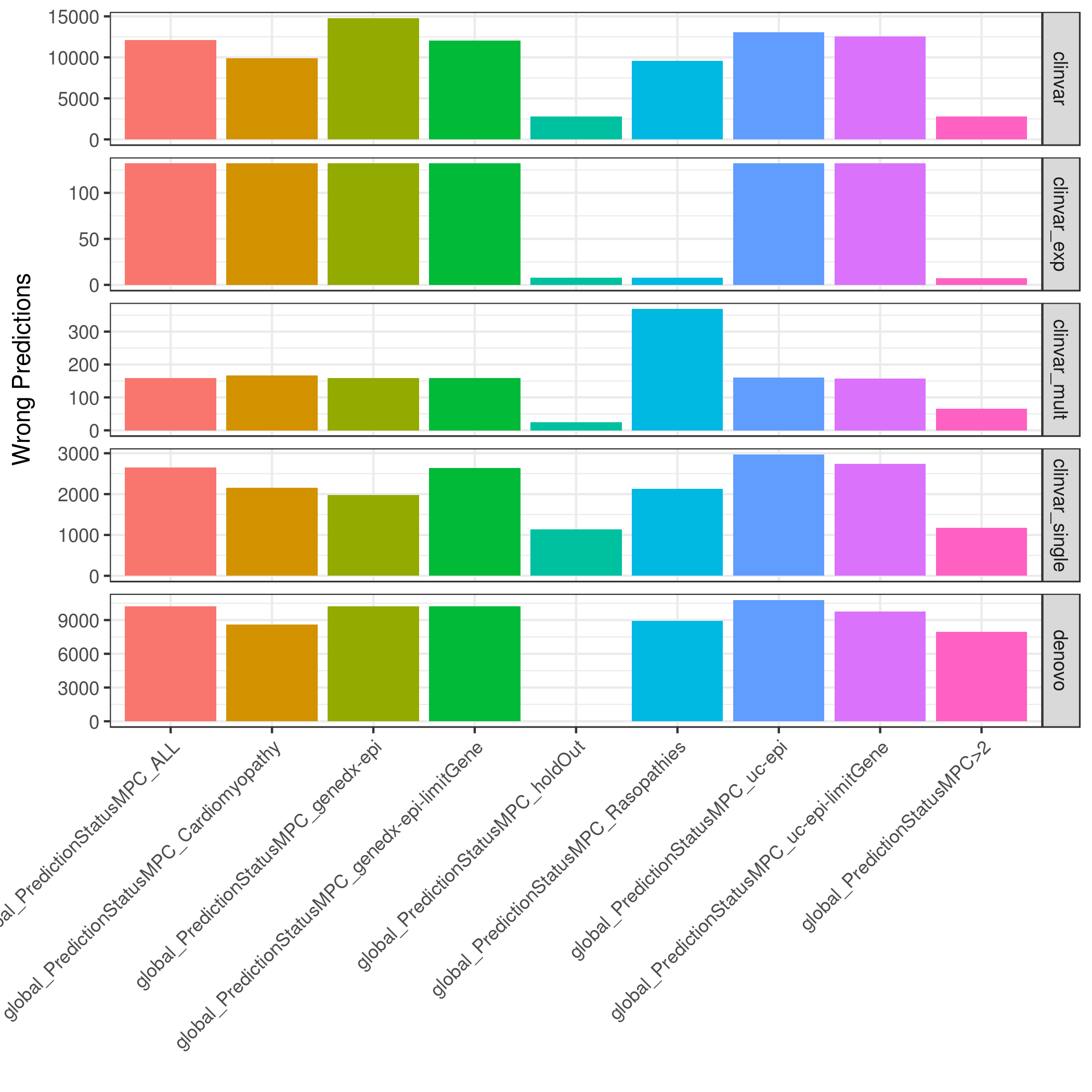

### global.other.totWrong.png

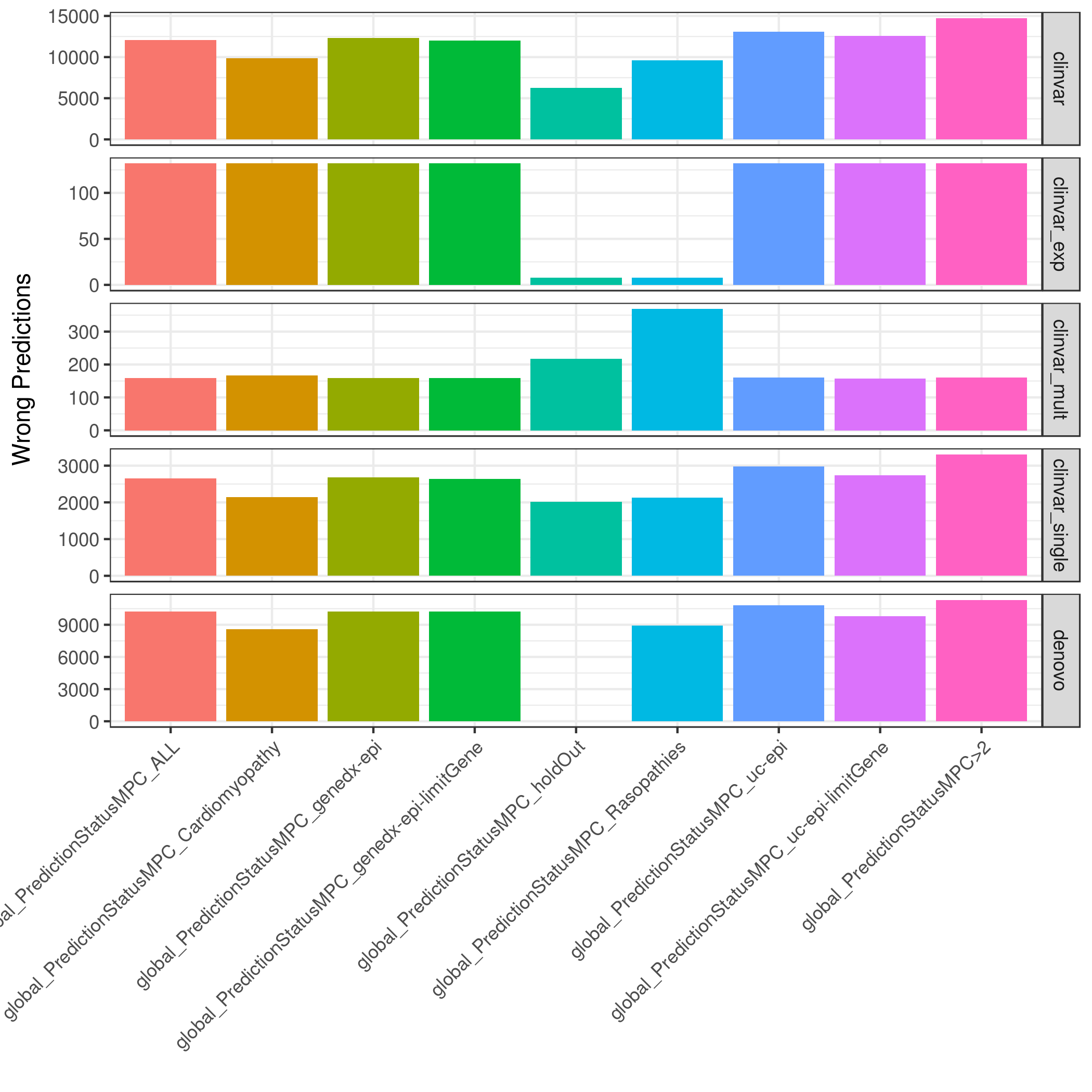

### global.revel.byDisease.png

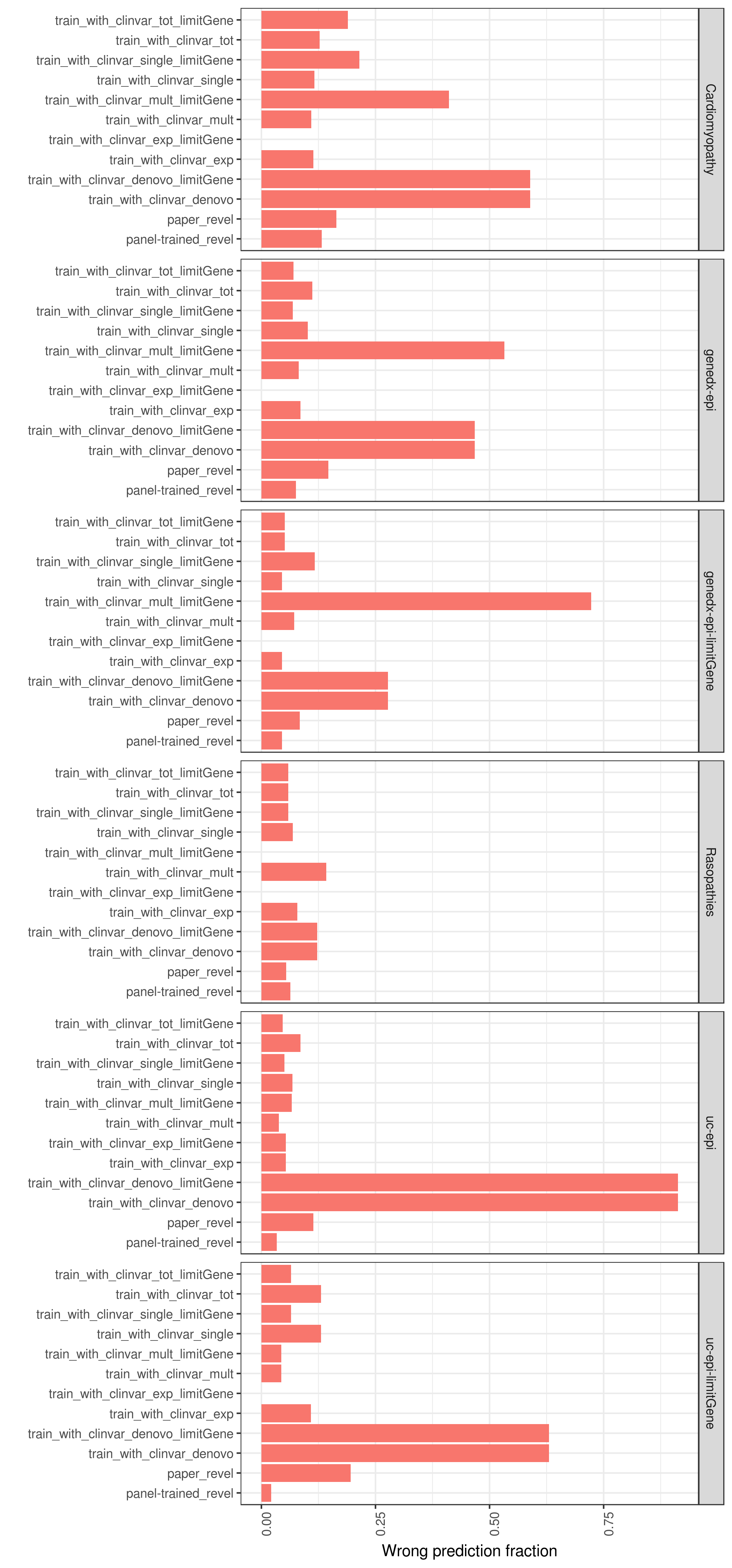

### GRIN2A.clinvar.lolly.png

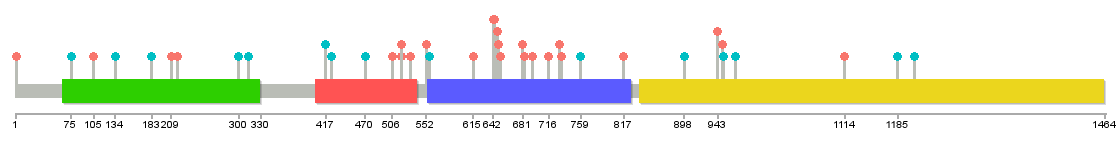

### KCNQ2.clinvar.lolly.png

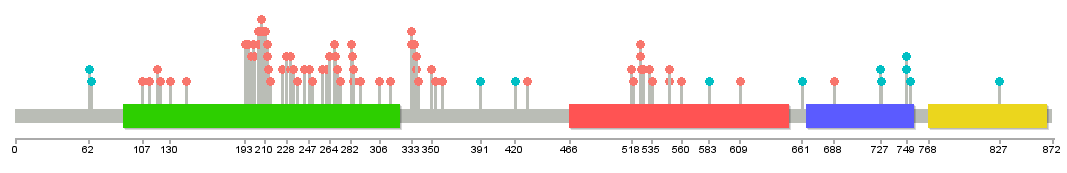

### KCNQ2.denovo_db.lolly.png

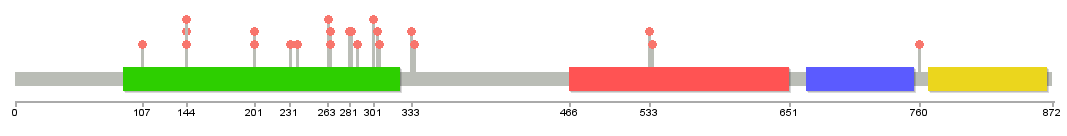

### KCNQ3.clinvar.lolly.png

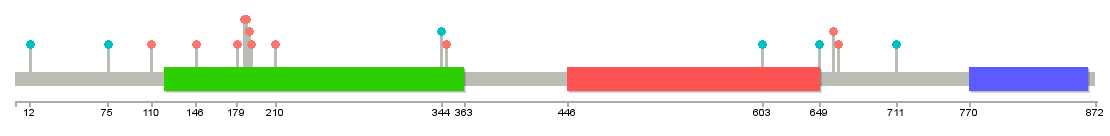

### KCNQ3.denovo_db.lolly.png

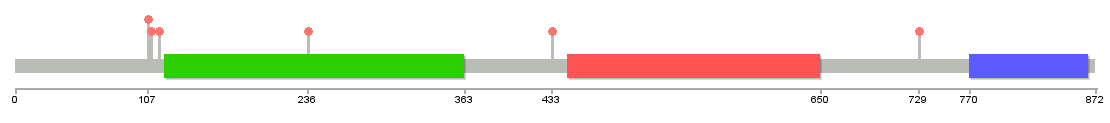

### lab2_gene_eval.png

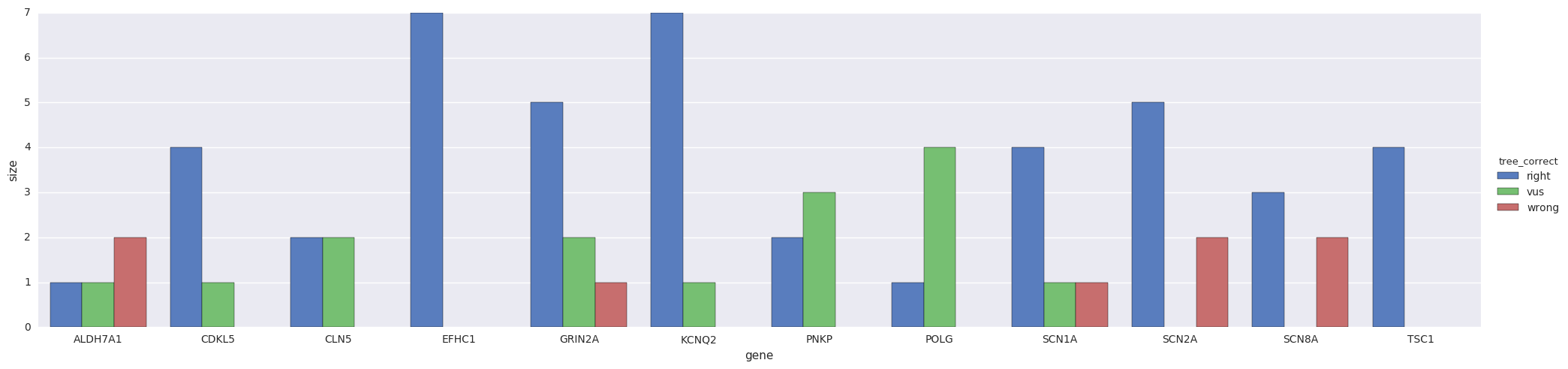

### missense_clinvar_roc_feature_union.CDKL5.png

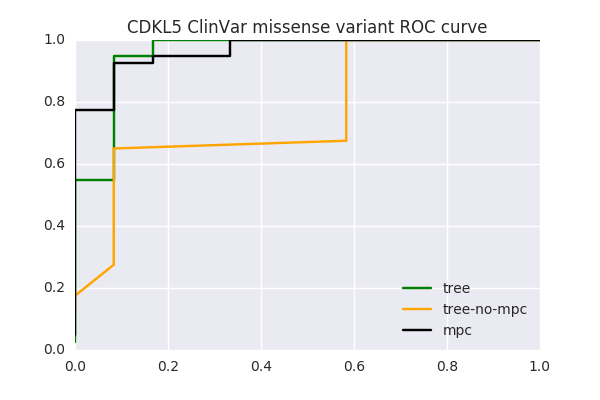

### missense_clinvar_roc_feature_union.KCNQ2.png

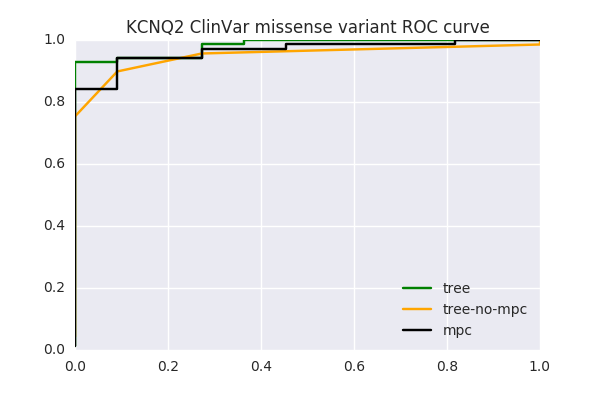

### missense_clinvar_roc_feature_union.KCNQ3.png

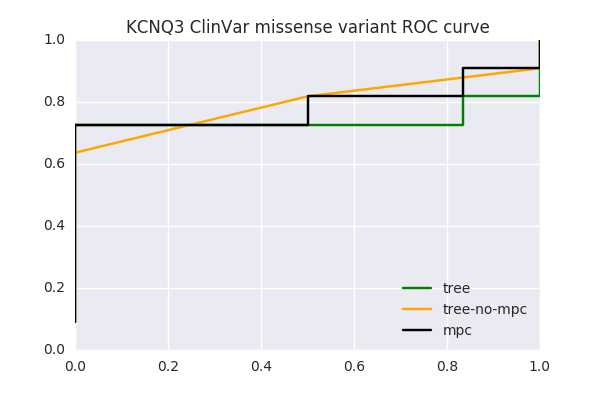

### missense_clinvar_roc_feature_union.SCN1A,SCN2A,KCNQ2,KCNQ3,CDKL5,PCDH19,SCN1B,SCN8A,SLC2A1,SPTAN1,STXBP1,TSC1.png

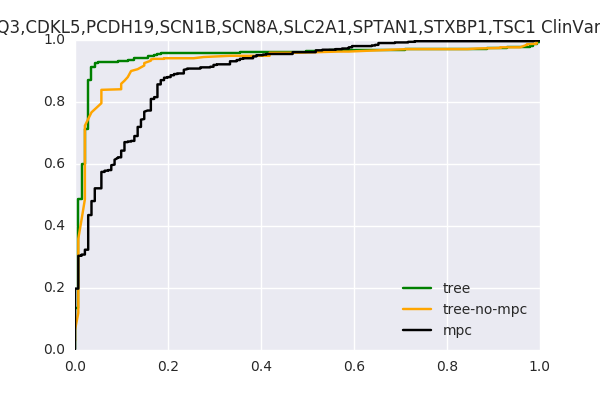

### missense_clinvar_roc_feature_union.SCN1A,SCN2A,KCNQ2,KCNQ3,CDKL5.png

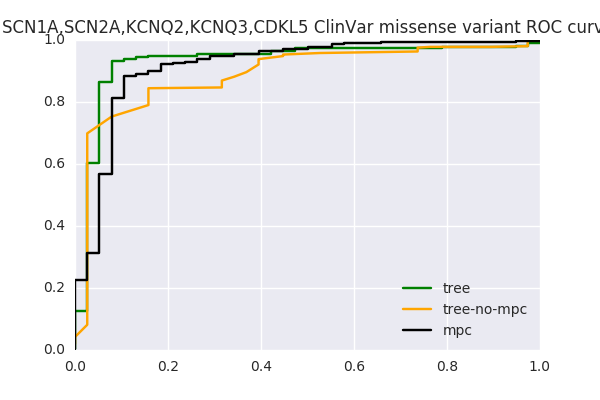

### missense_clinvar_roc_feature_union.SCN1A.png

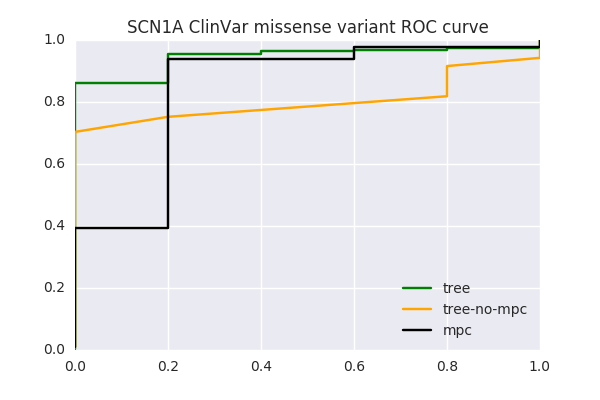

### missense_clinvar_roc_feature_union.SCN1B.png

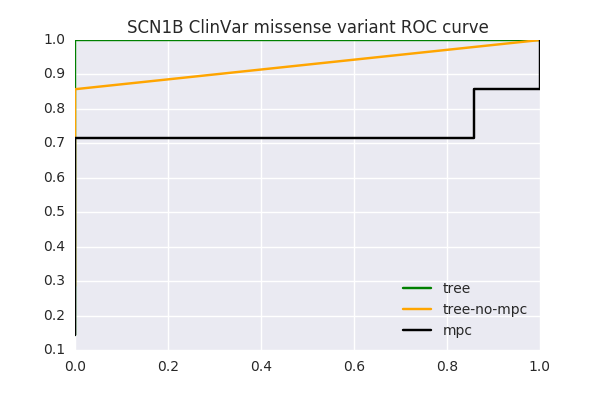

### missense_clinvar_roc_feature_union.STXBP1.png

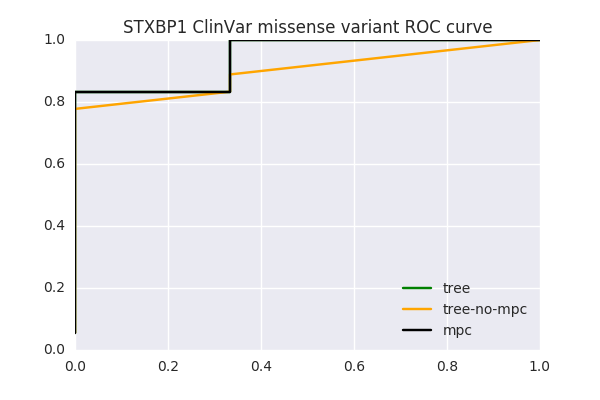

### missense_clinvar_roc_feature_union.TSC1.png

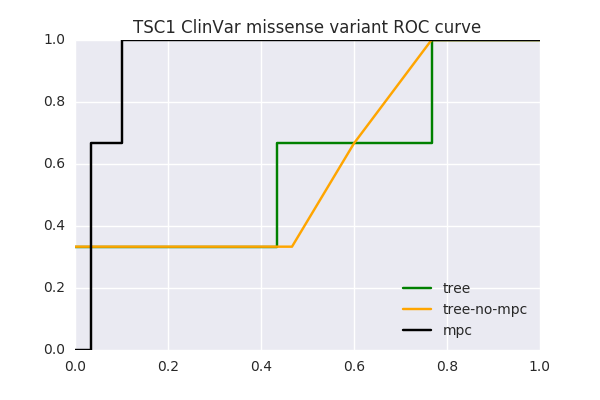

### missense_fg_roc.png

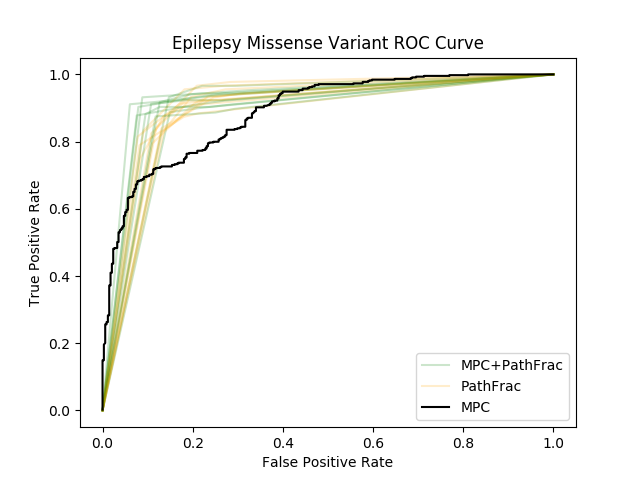

### mpc.by_pfam_enrichment.png

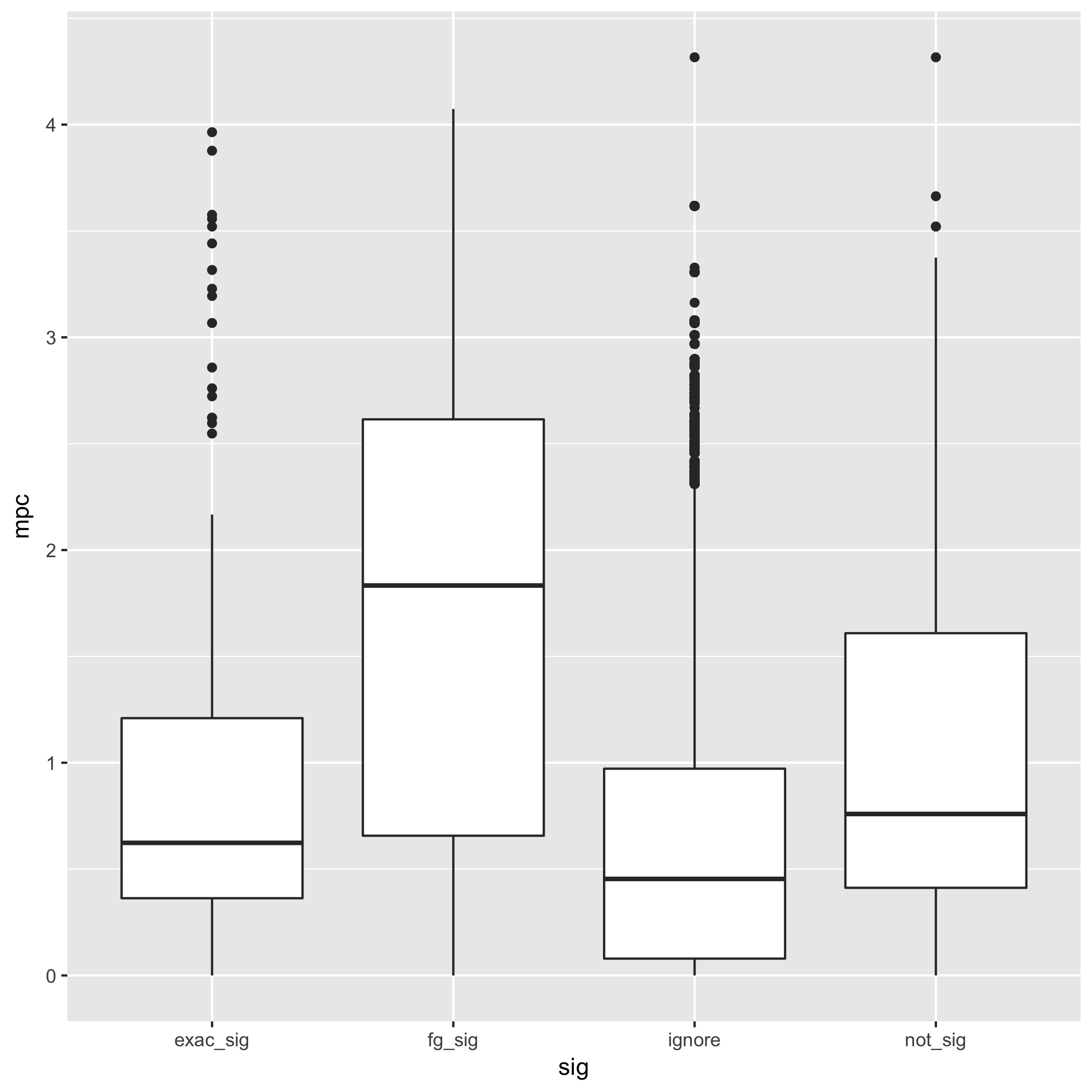

### PCDH19.clinvar.lolly.png

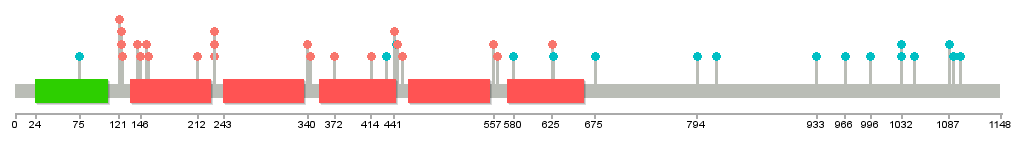

### PCDH19.denovodb.lolly.png

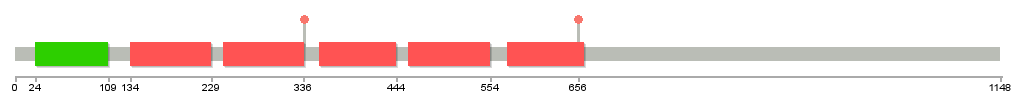

### rare.benign_frac_w_vus.pfam.mpcLow_0.mpcHigh_1.4.png

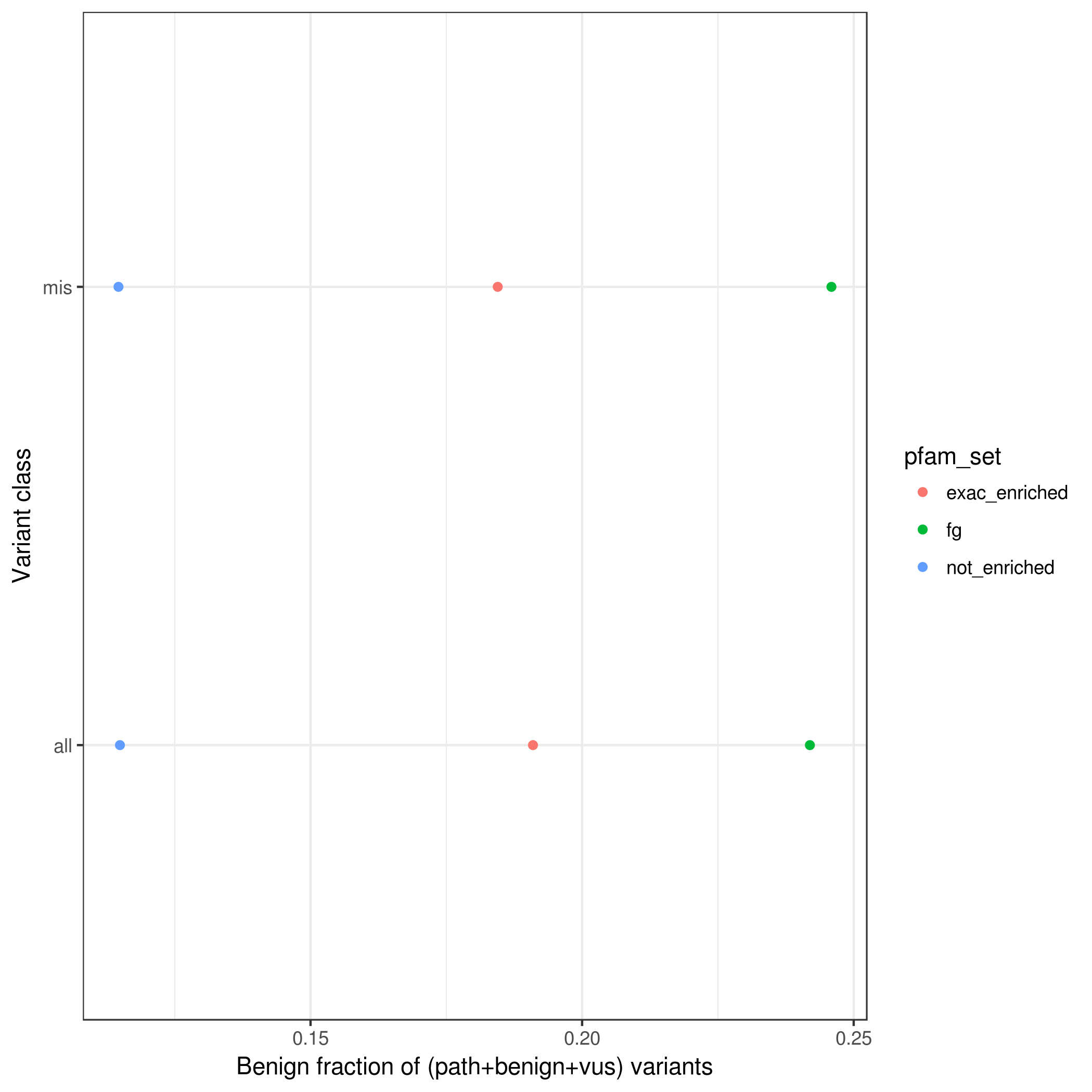

### rare.path_frac_w_vus.pfam.mpcLow_1.4.mpcHigh_100.png

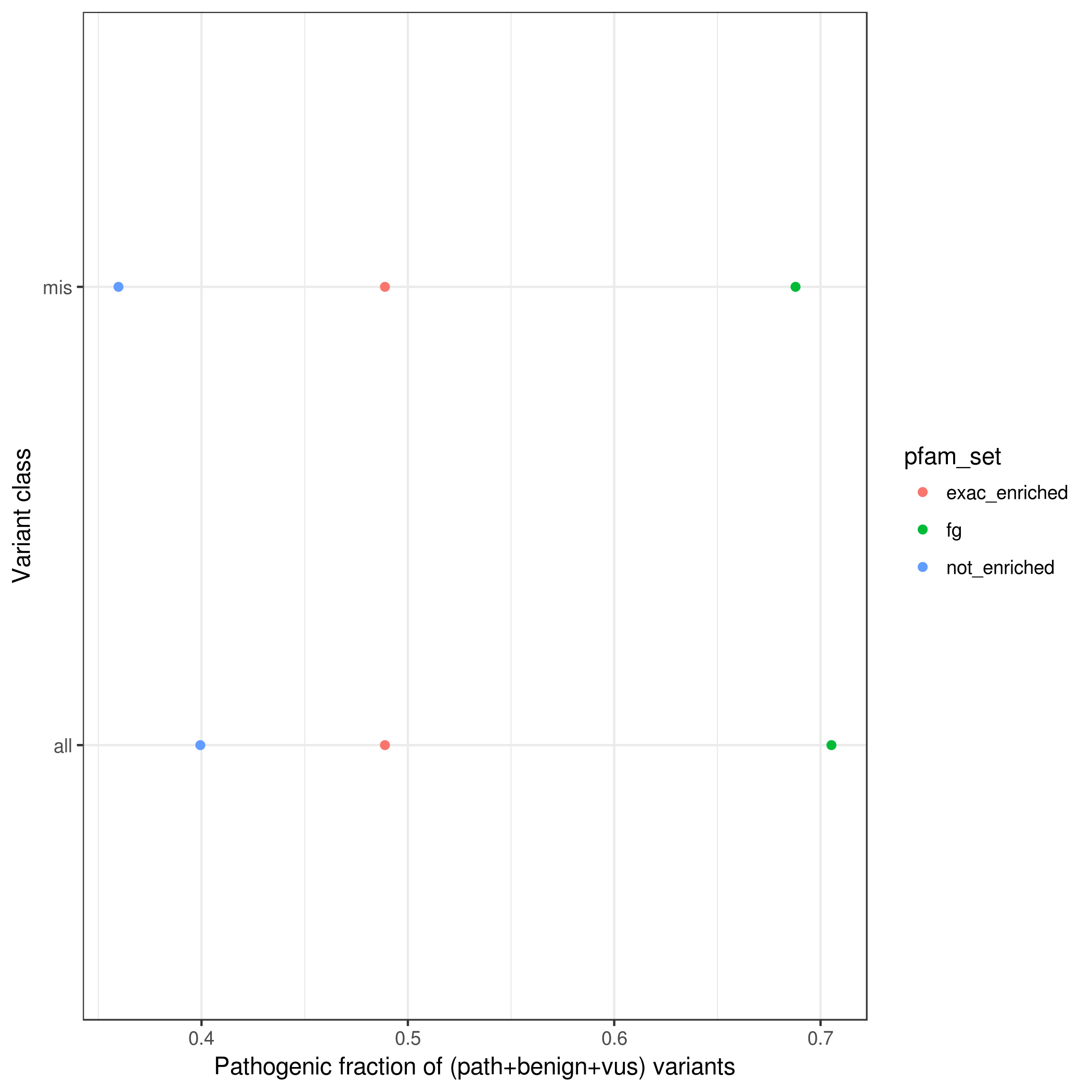

### rare.path_frac_w_vus.pfamMerge.mpcLow_0.mpcHigh_1.4.png

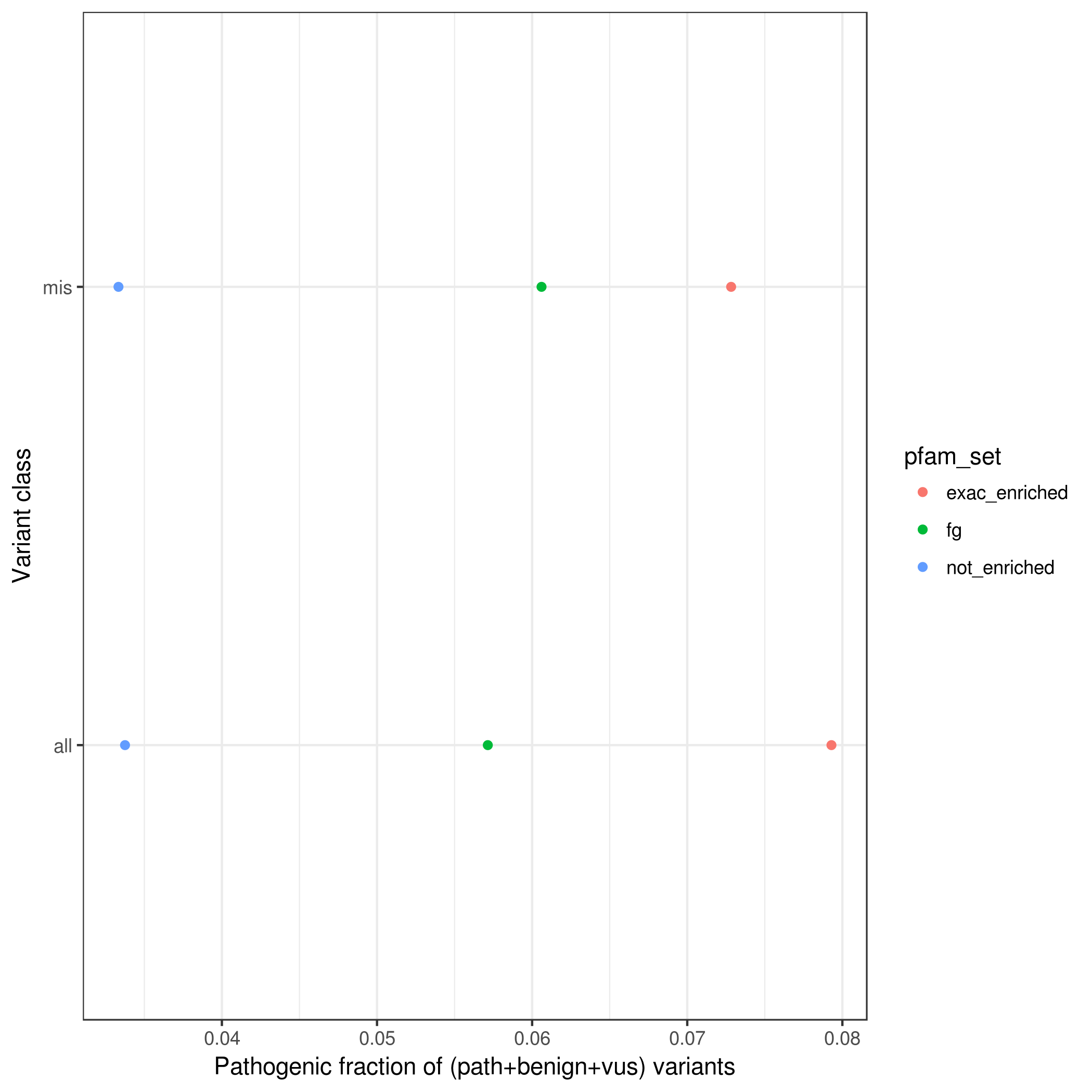

### rare.path_frac_w_vus.pfamMerge.mpcLow_0.mpcHigh_100.png

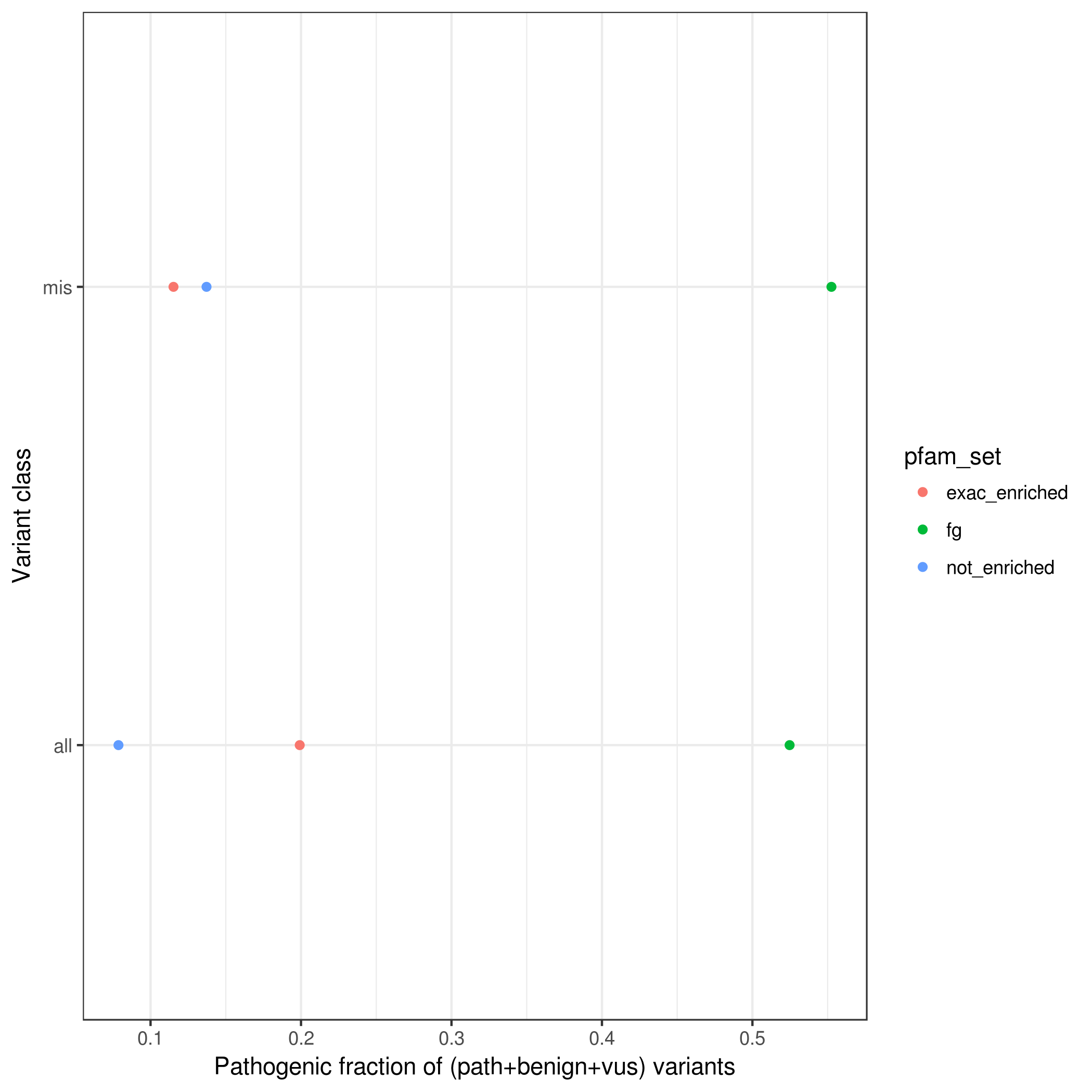

### rare.path_frac_wo_vus.pfam.mpcLow_0.mpcHigh_1.4.png

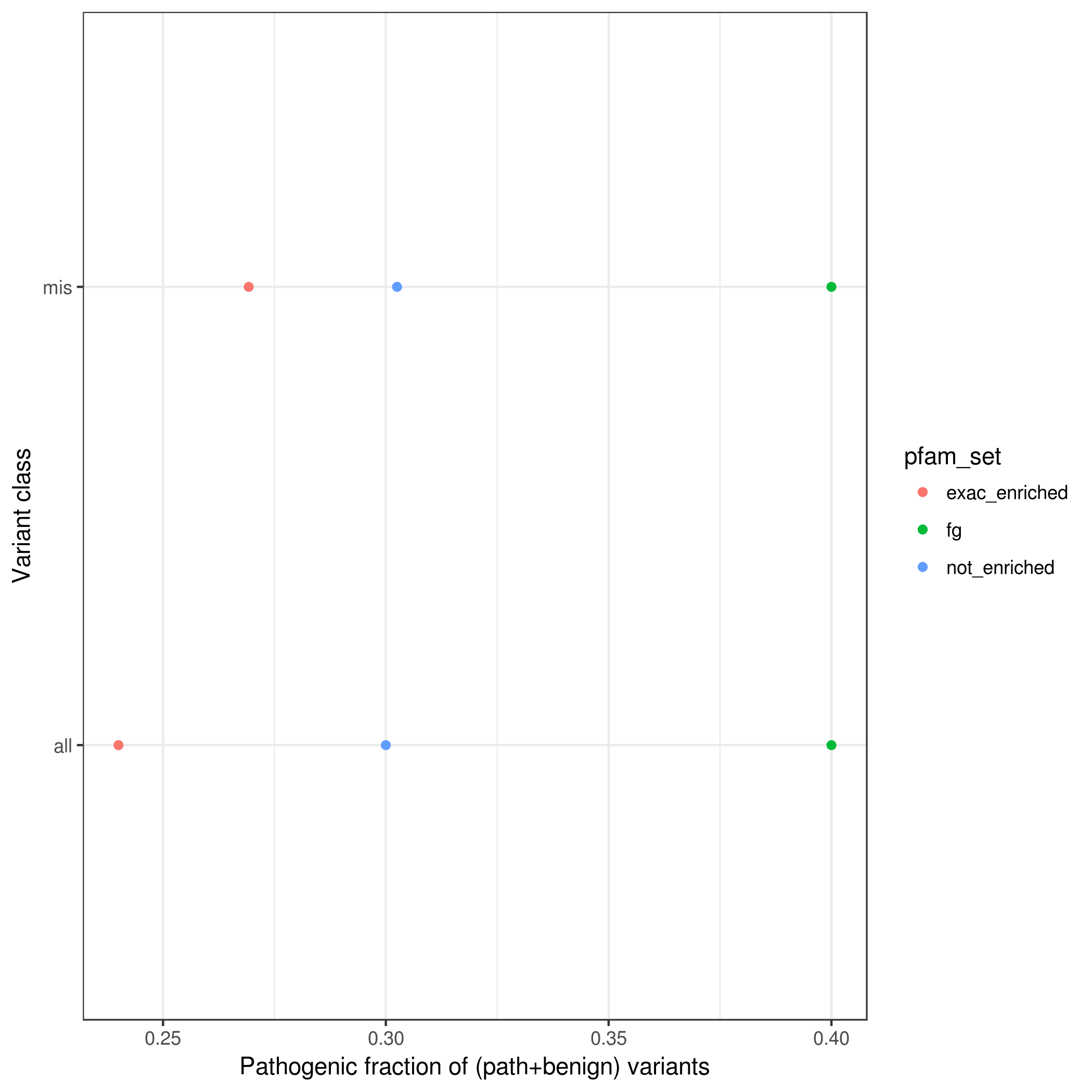

### rare.path_frac_wo_vus.pfam.mpcLow_1.4.mpcHigh_100.png

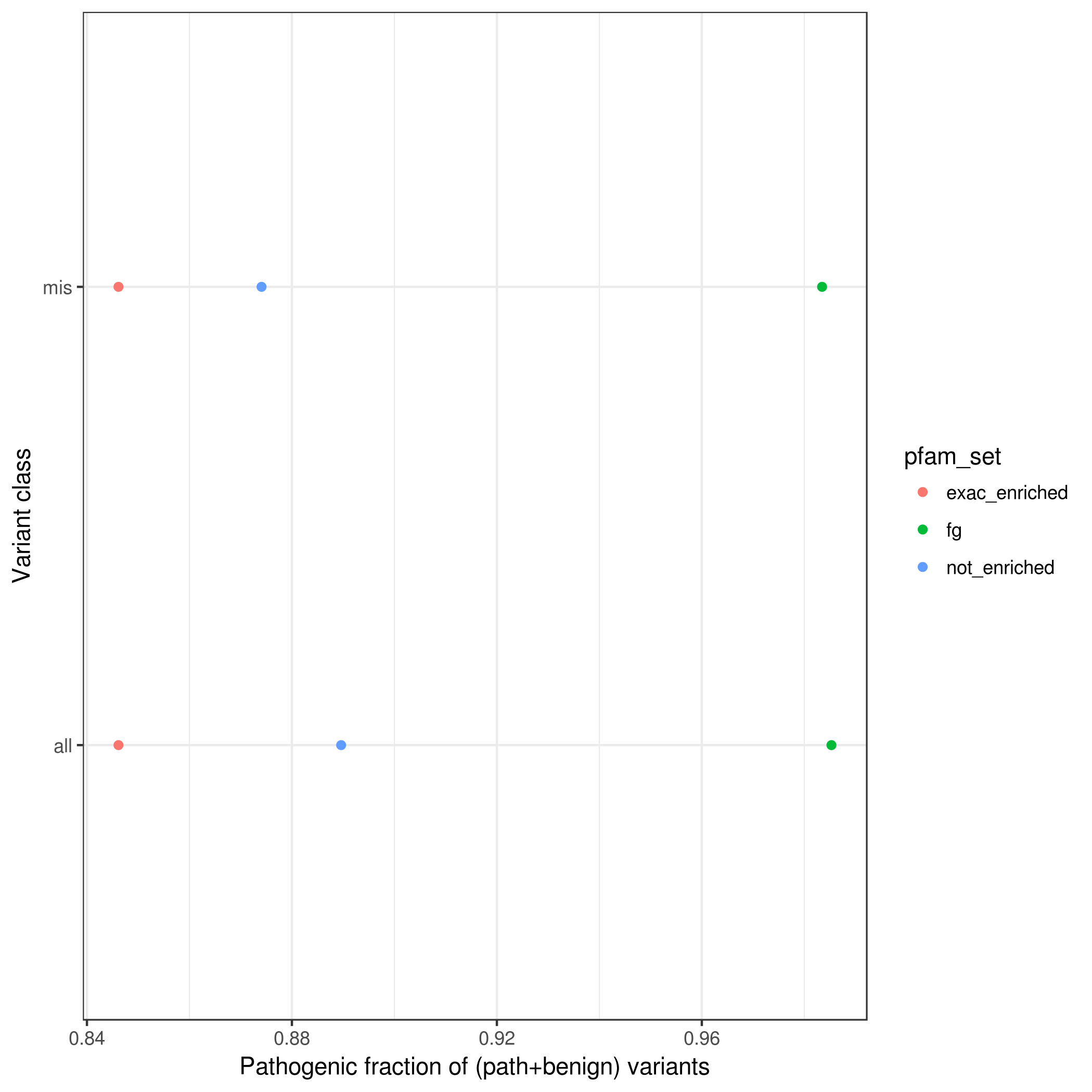

### rare.path_frac_wo_vus.pfamMerge.mpcLow_0.mpcHigh_1.4.png

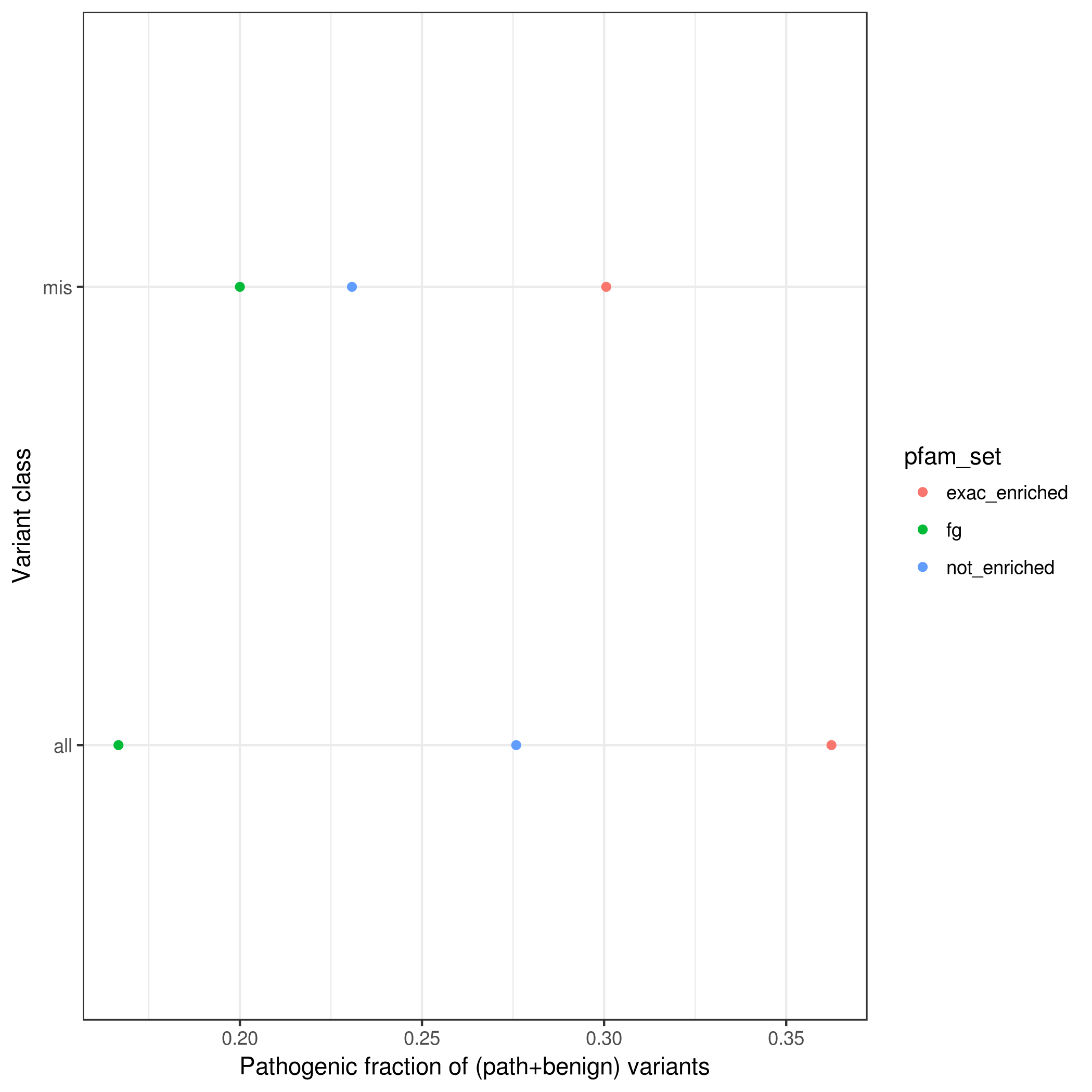

### rare.var_counts_by_enrichment.pfam.mpcLow_0.mpcHigh_100.png

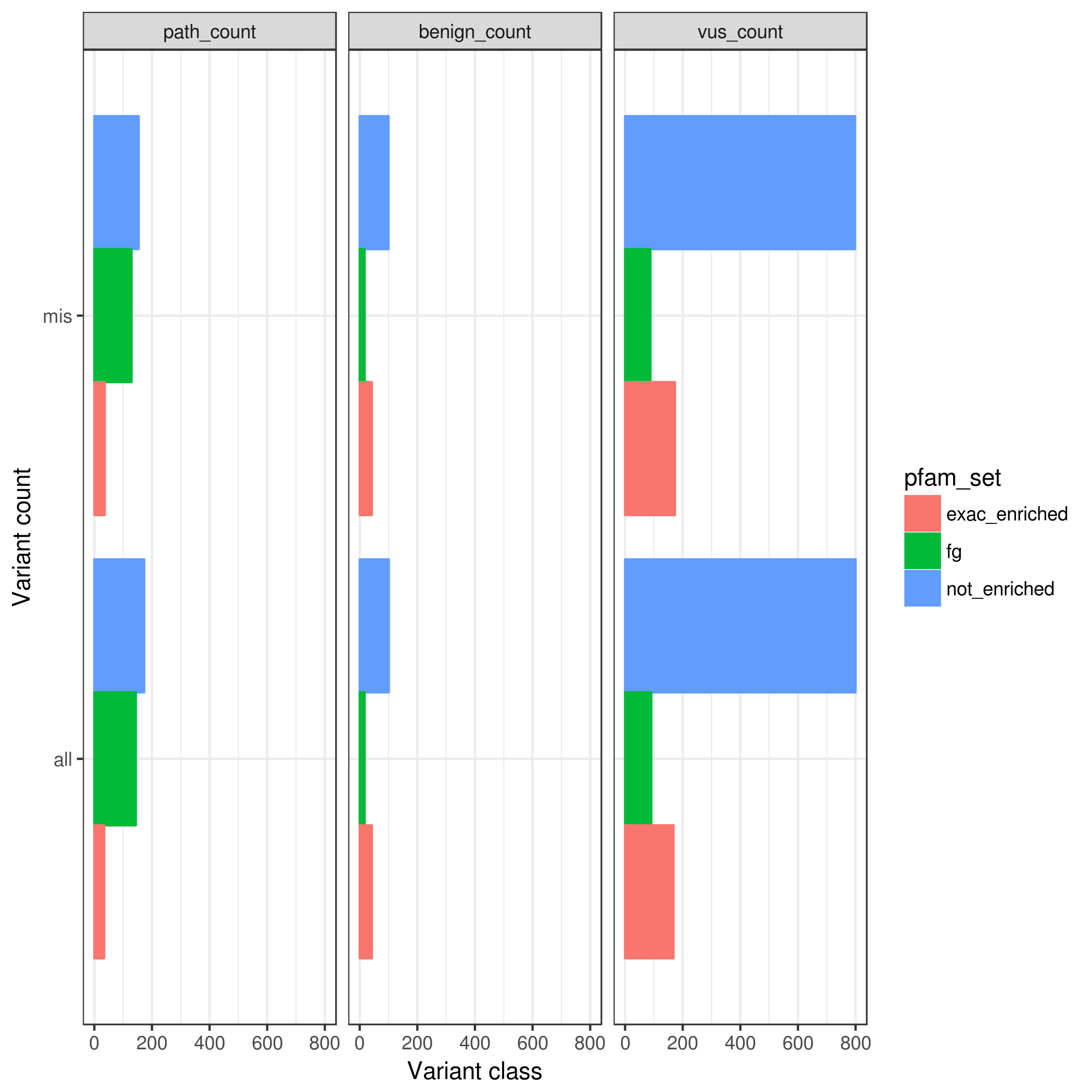

### rare.var_counts_by_enrichment.pfam.mpcLow_1.4.mpcHigh_100.png

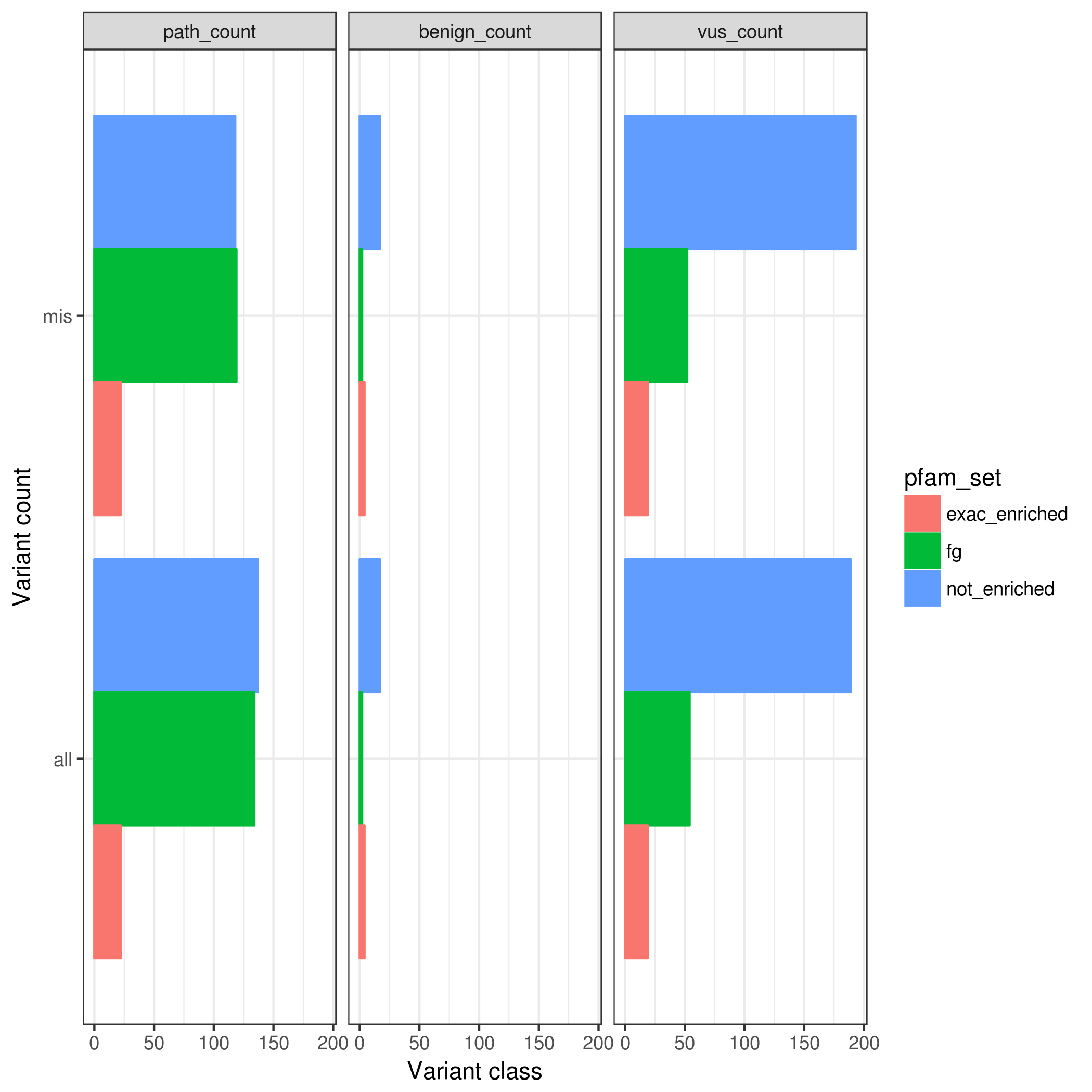

### rare.var_counts_by_enrichment.pfamMerge.mpc_1.4.png

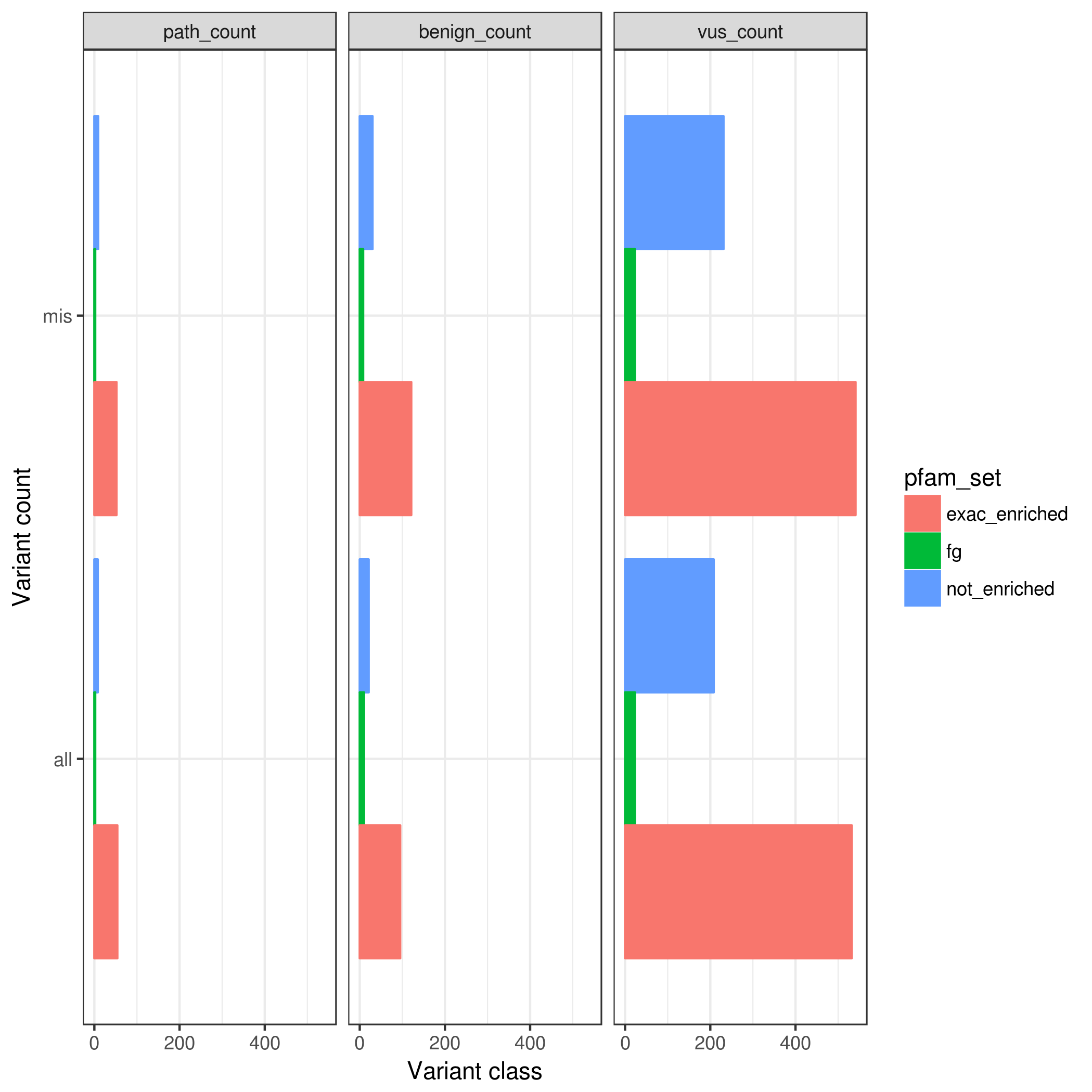

### rare.var_counts_by_enrichment.pfamMerge.mpcLow_0.mpcHigh_100.png

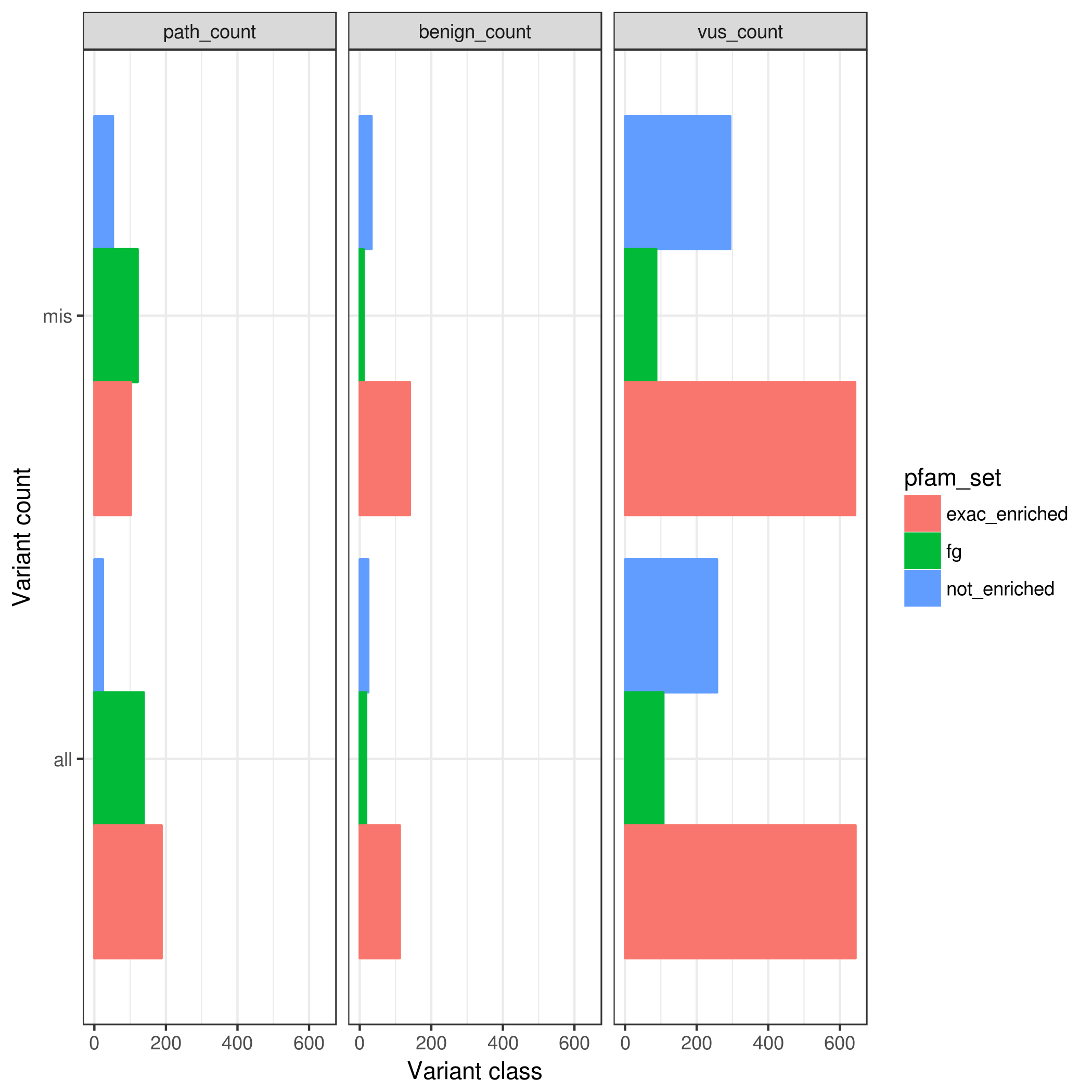

### rarem.benign_frac_w_vus.pfam.mpc_100.png

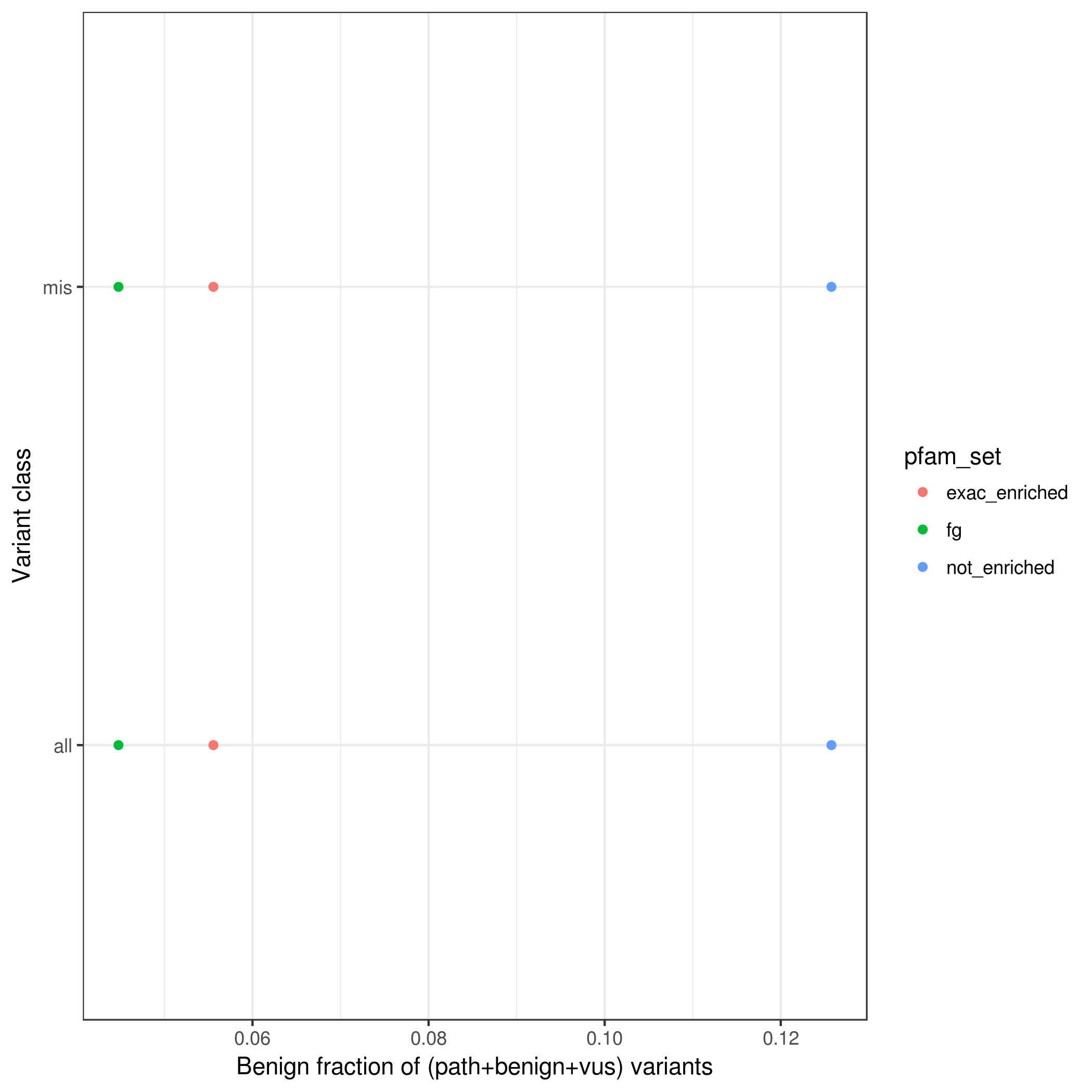

### rarem.benign_frac_w_vus.pfamMerge.mpc_1.4.png

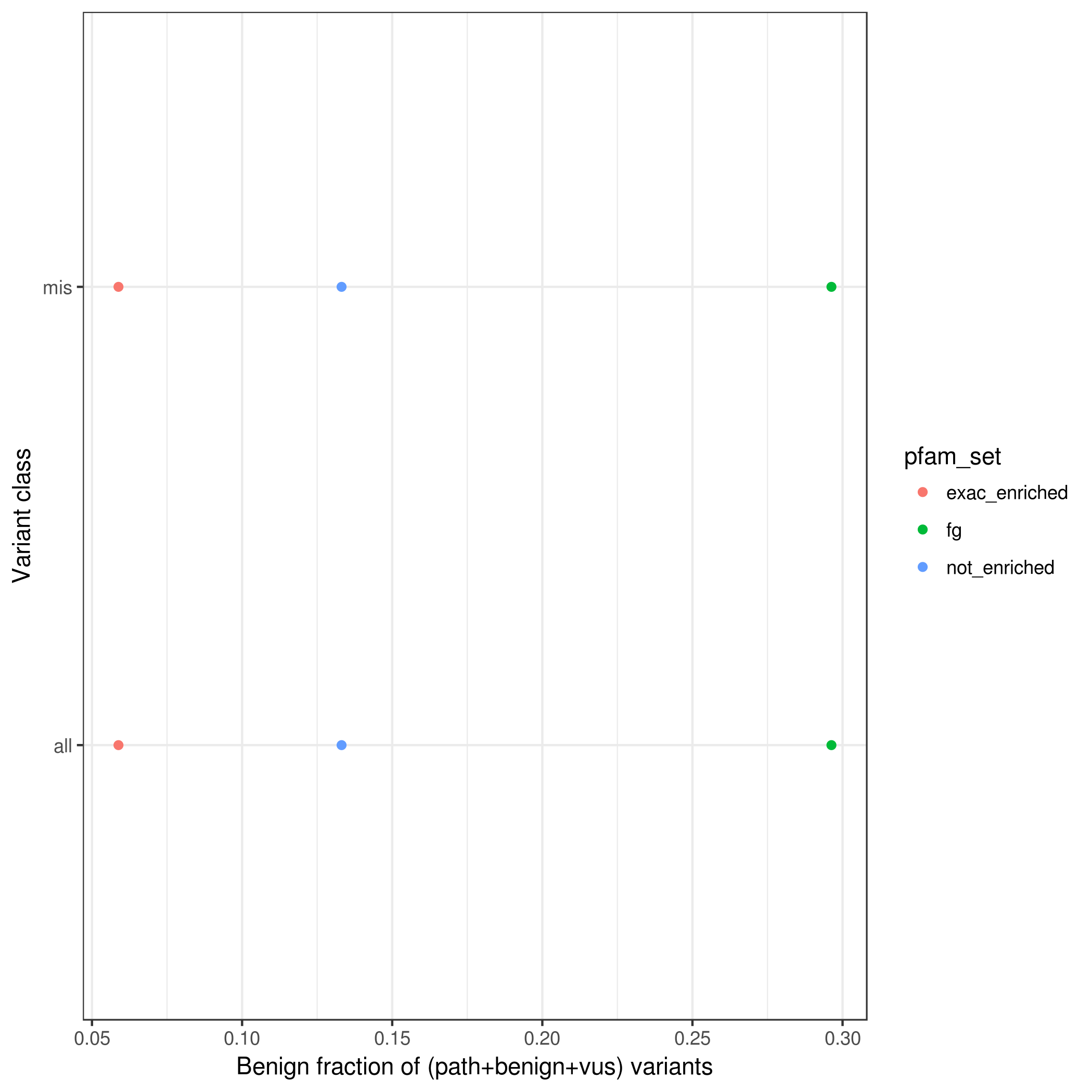

### rarem.benign_frac_w_vus.pfamMerge.mpcLow_0.mpcHigh_100.png

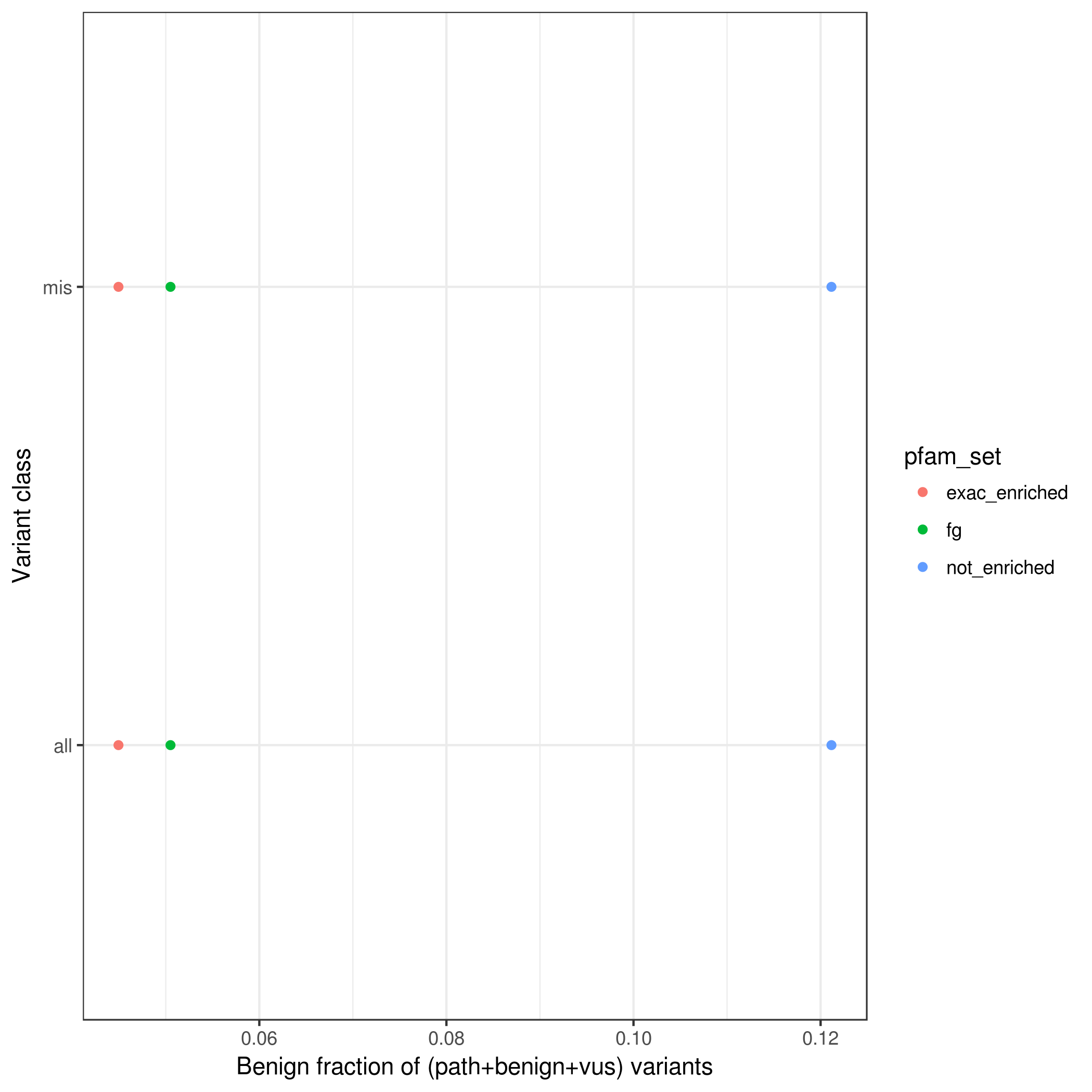

### rarem.path_frac_wo_vus.pfam.mpcLow_0.mpcHigh_1.4.png

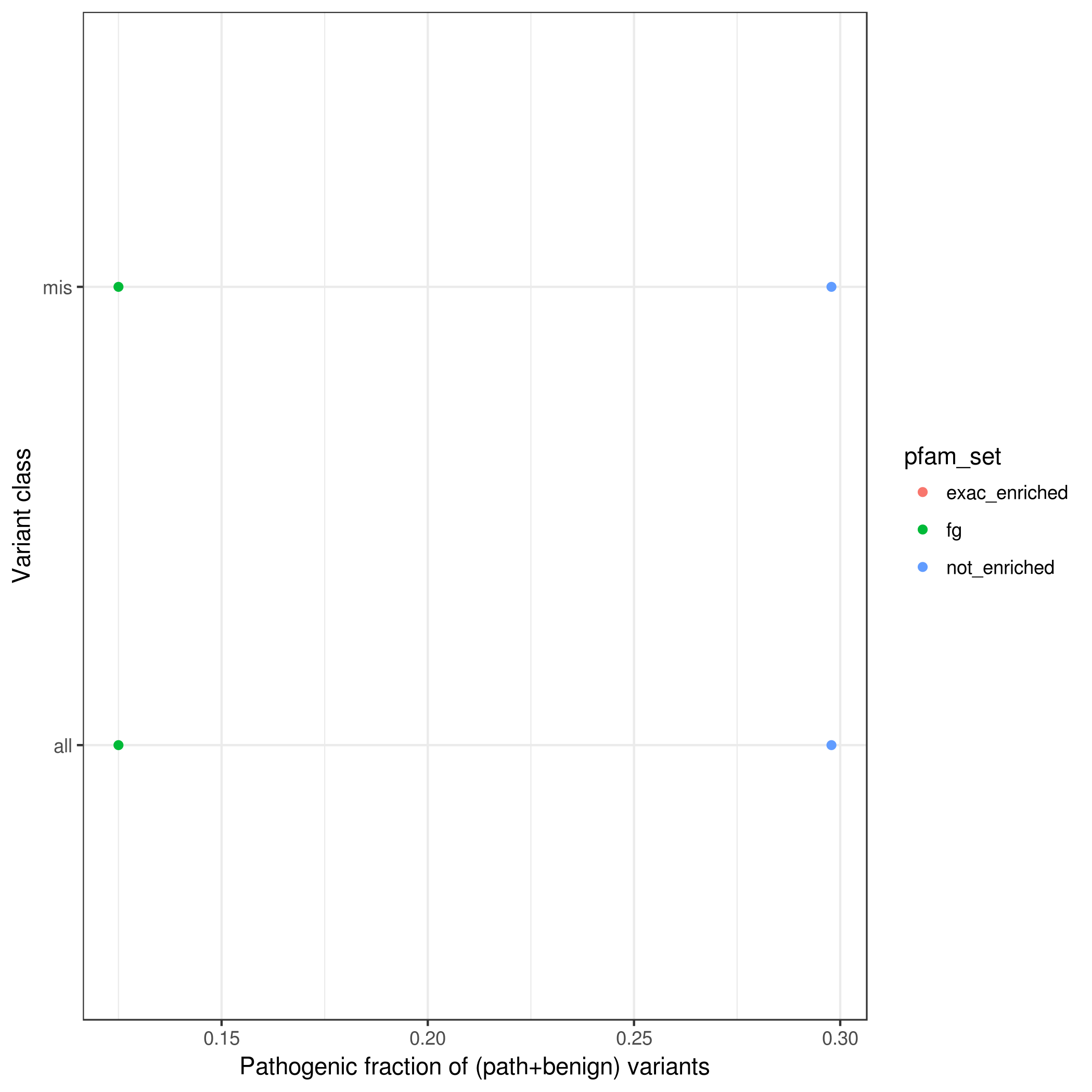

### rarem.path_frac_wo_vus.pfam.mpcLow_1.4.mpcHigh_100.png

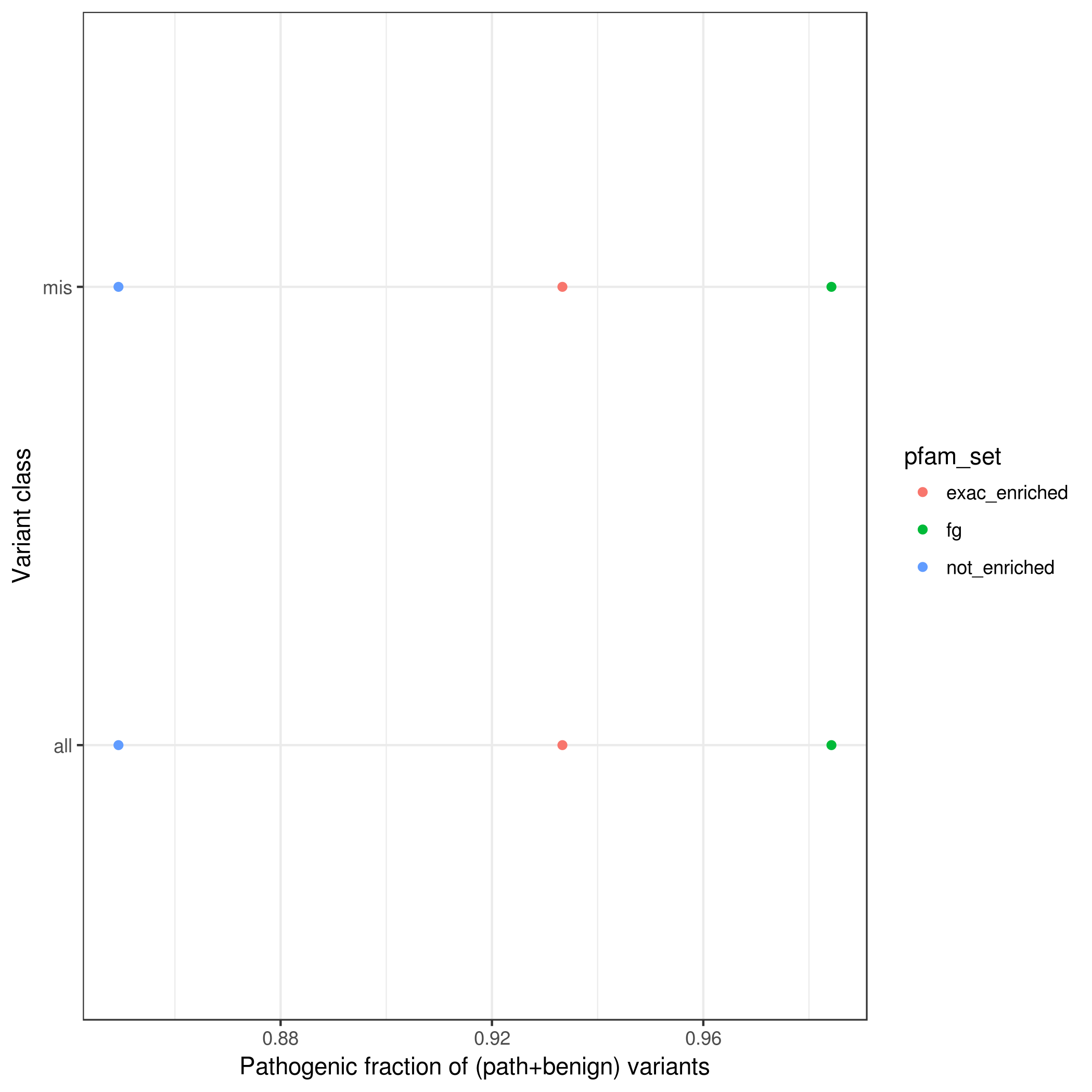

### rarem.path_frac_wo_vus.pfamMerge.mpc_100.png

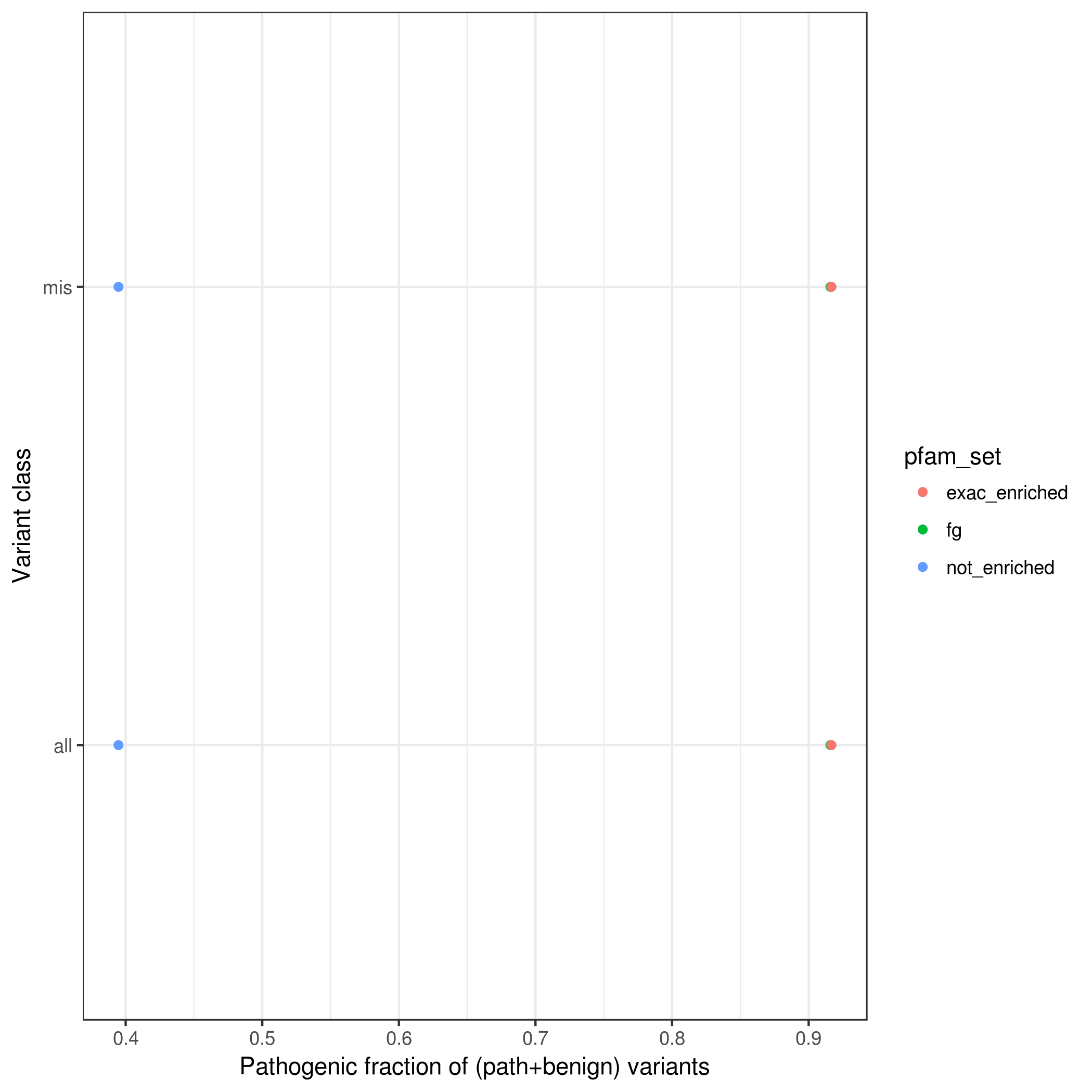

### rarem.path_frac_wo_vus.pfamMerge.mpcLow_1.4.mpcHigh_100.png

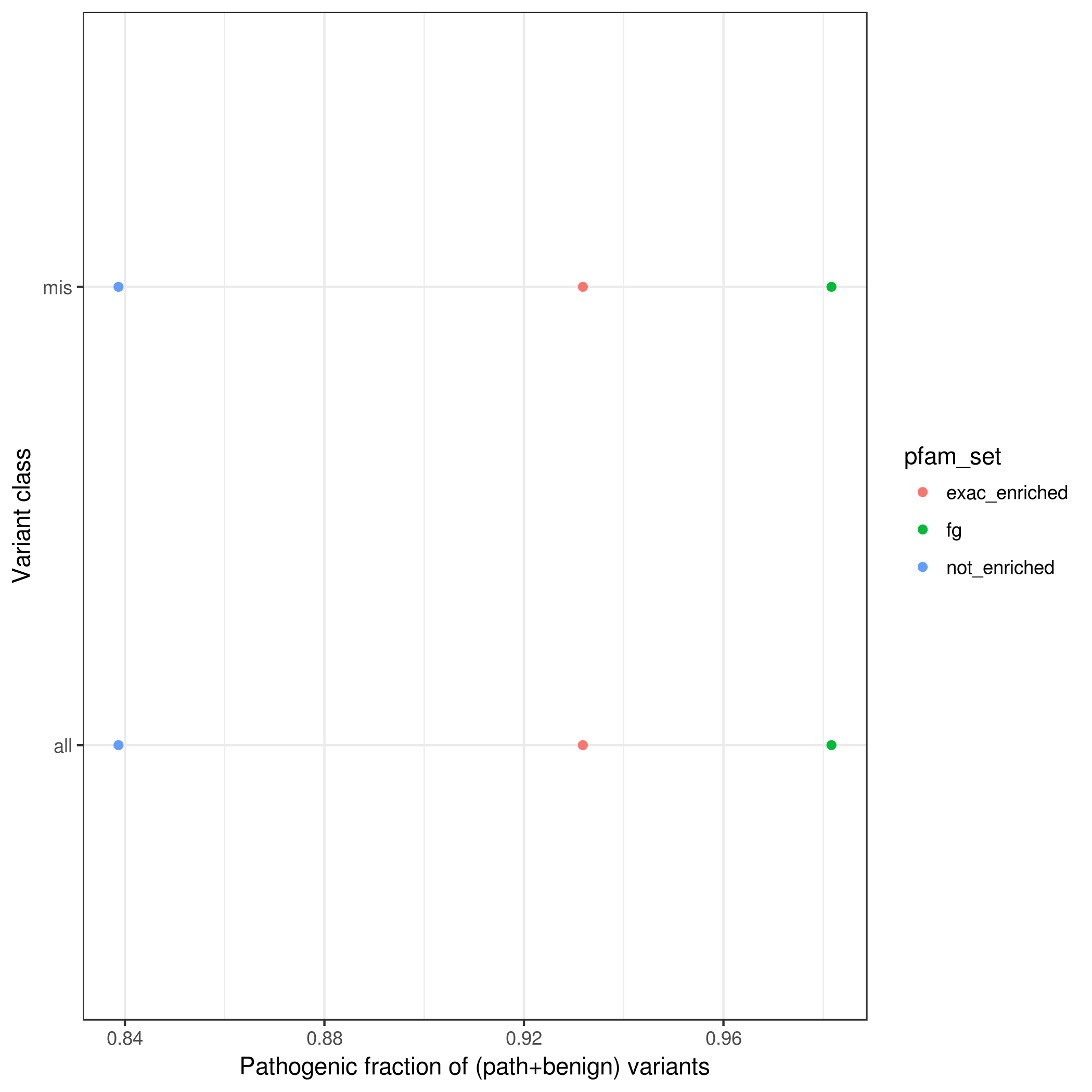

### rarem.var_counts_by_enrichment.pfam.mpcLow_0.mpcHigh_100.png

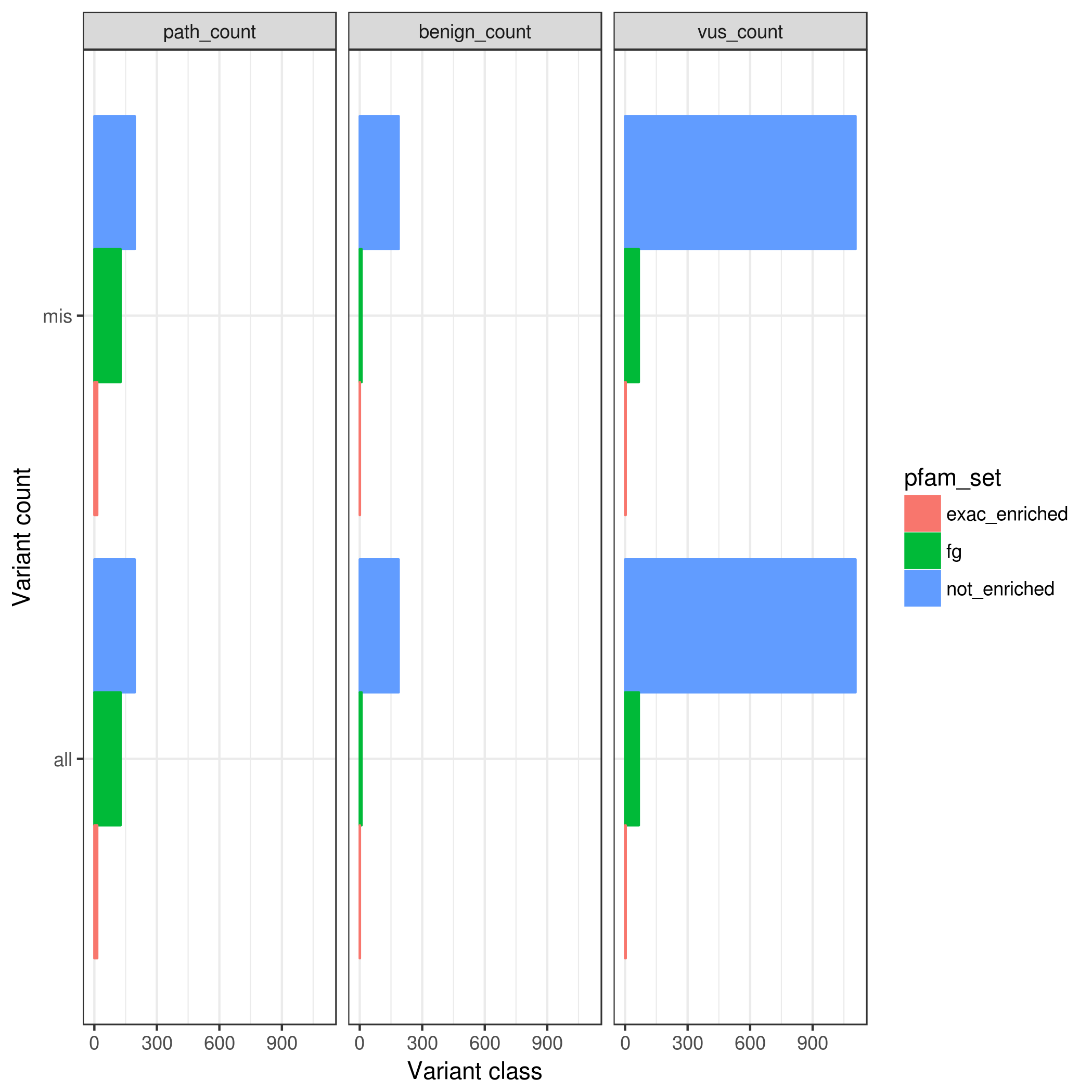

### rarem.var_counts_by_enrichment.pfamMerge.mpc_1.4.png

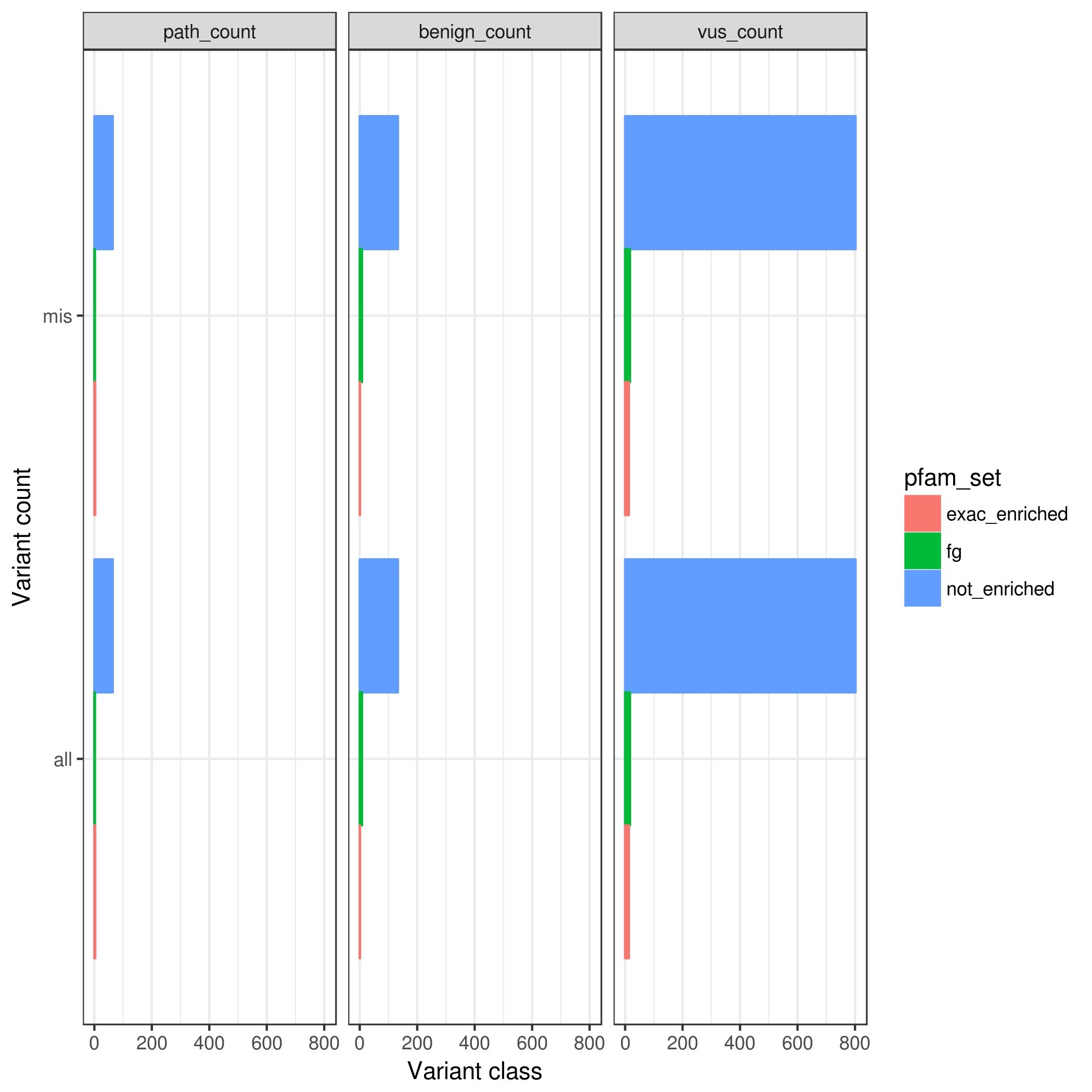

### SCN1A.clinvar.lolly.png

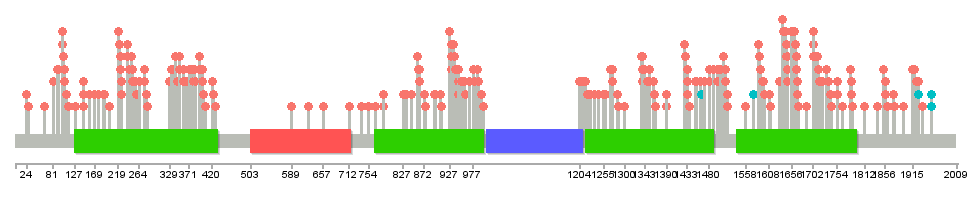

### SCN1A.denovo_db.lolly.png

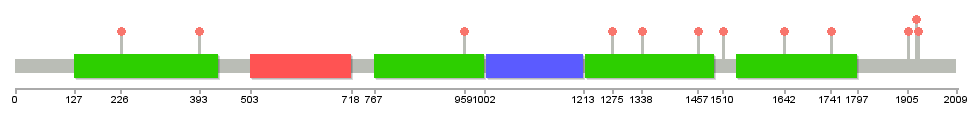

### SCN1B.clinvar.lolly.png

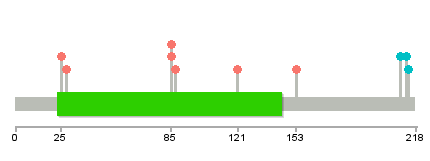

### SCN1B.denovo_db.lolly.png

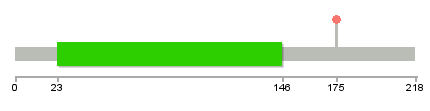

### SCN2A.clinvar.lolly.png

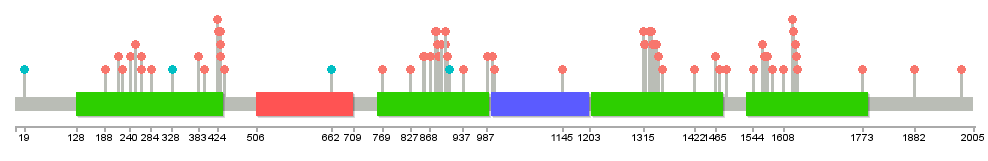

### SCN2A.denovodb.lolly.png

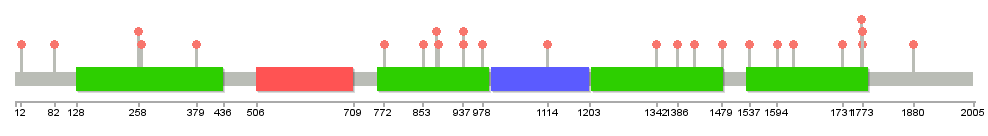

### SCN8A.clinvar.lolly.png

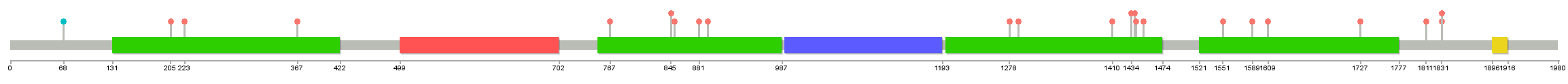

### SCN8A.denovodb.lolly.png

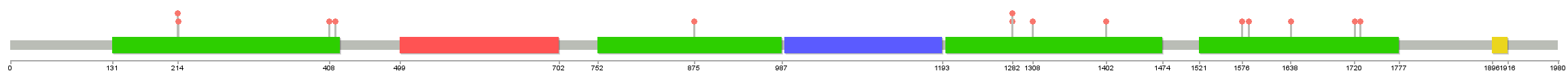

### single_gene_.003_1.pdf

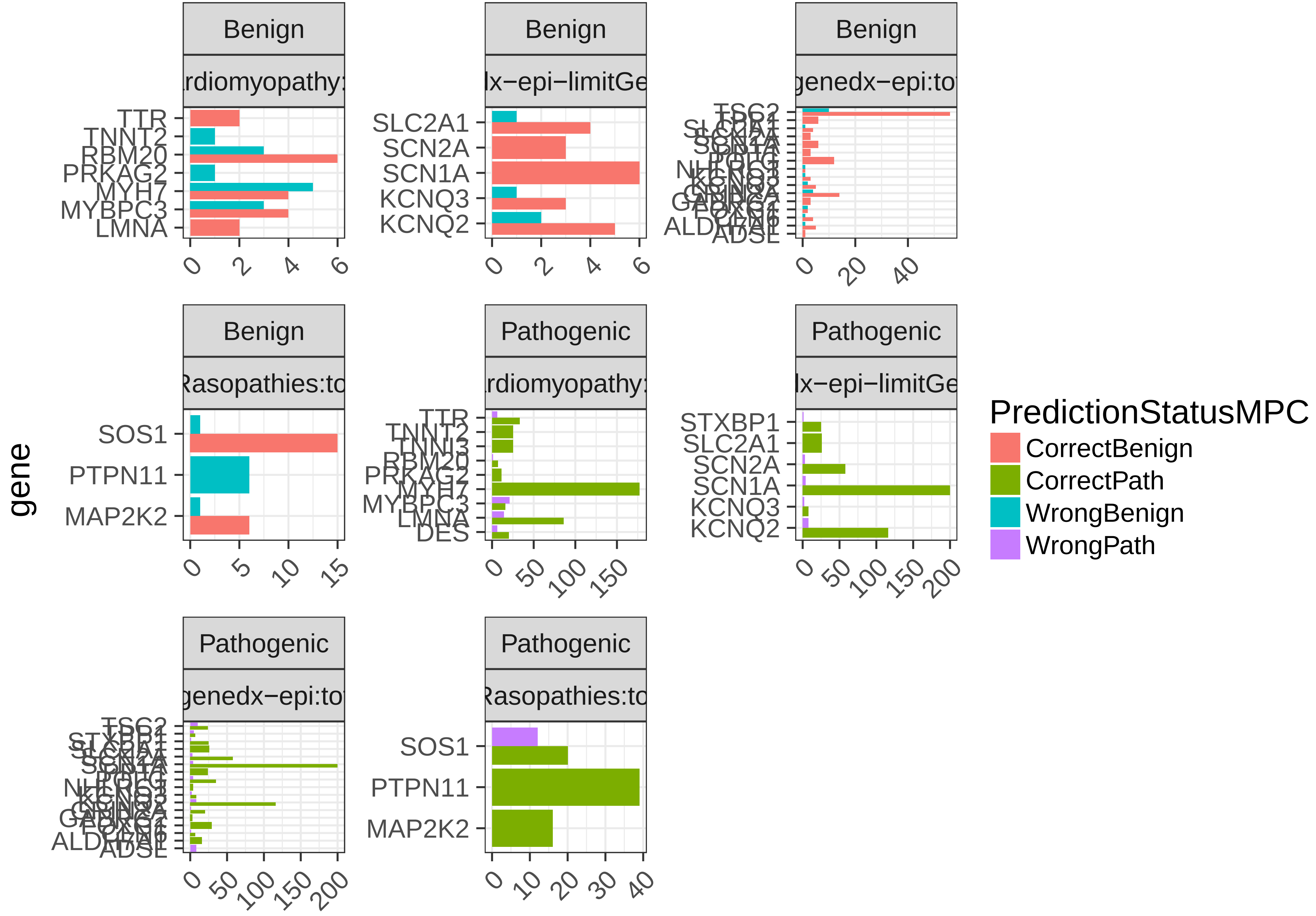

### single_gene_.003_.1.pdf

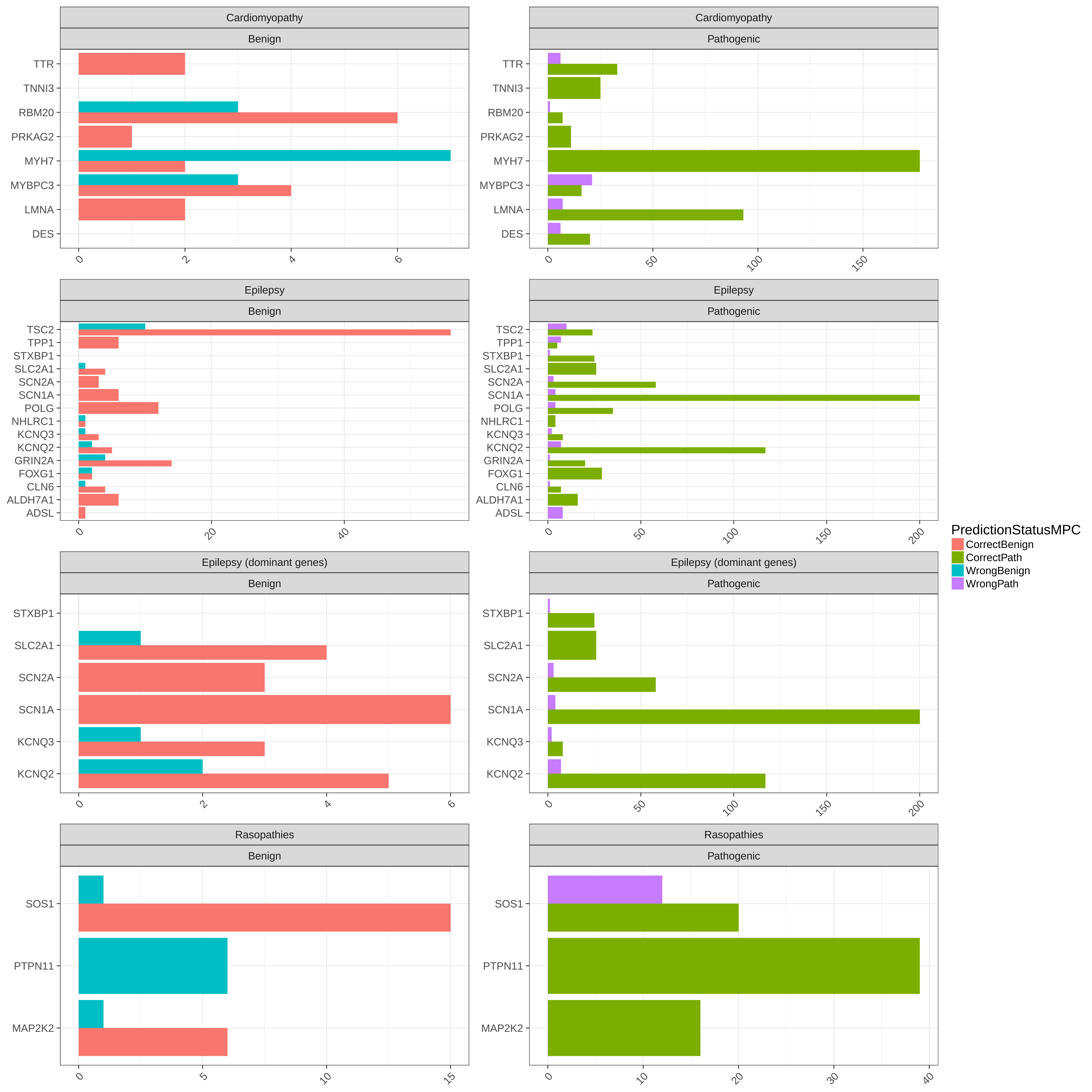

### single_gene_collapse_.003_1.pdf

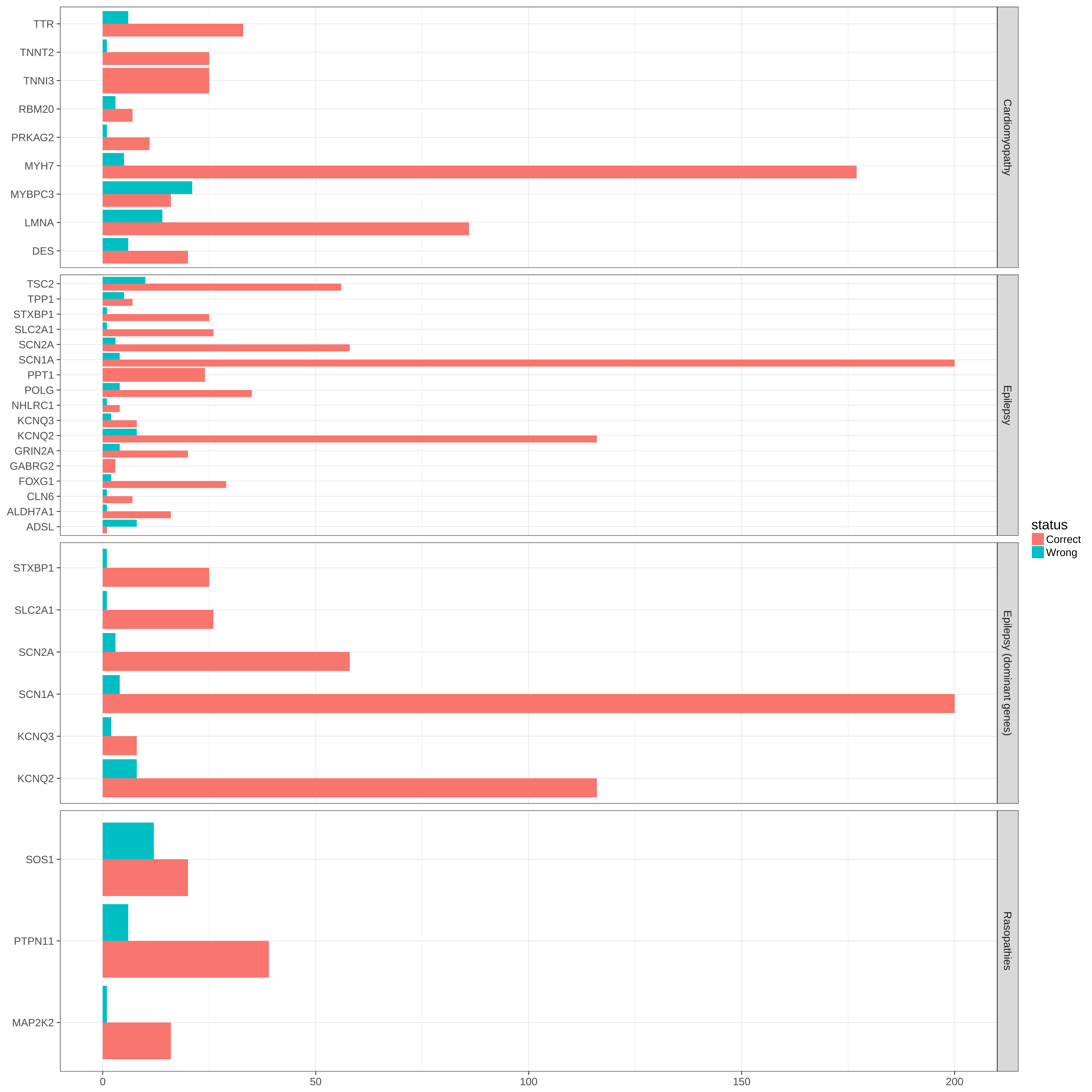

### single_gene_collapse_.003_.1.pdf

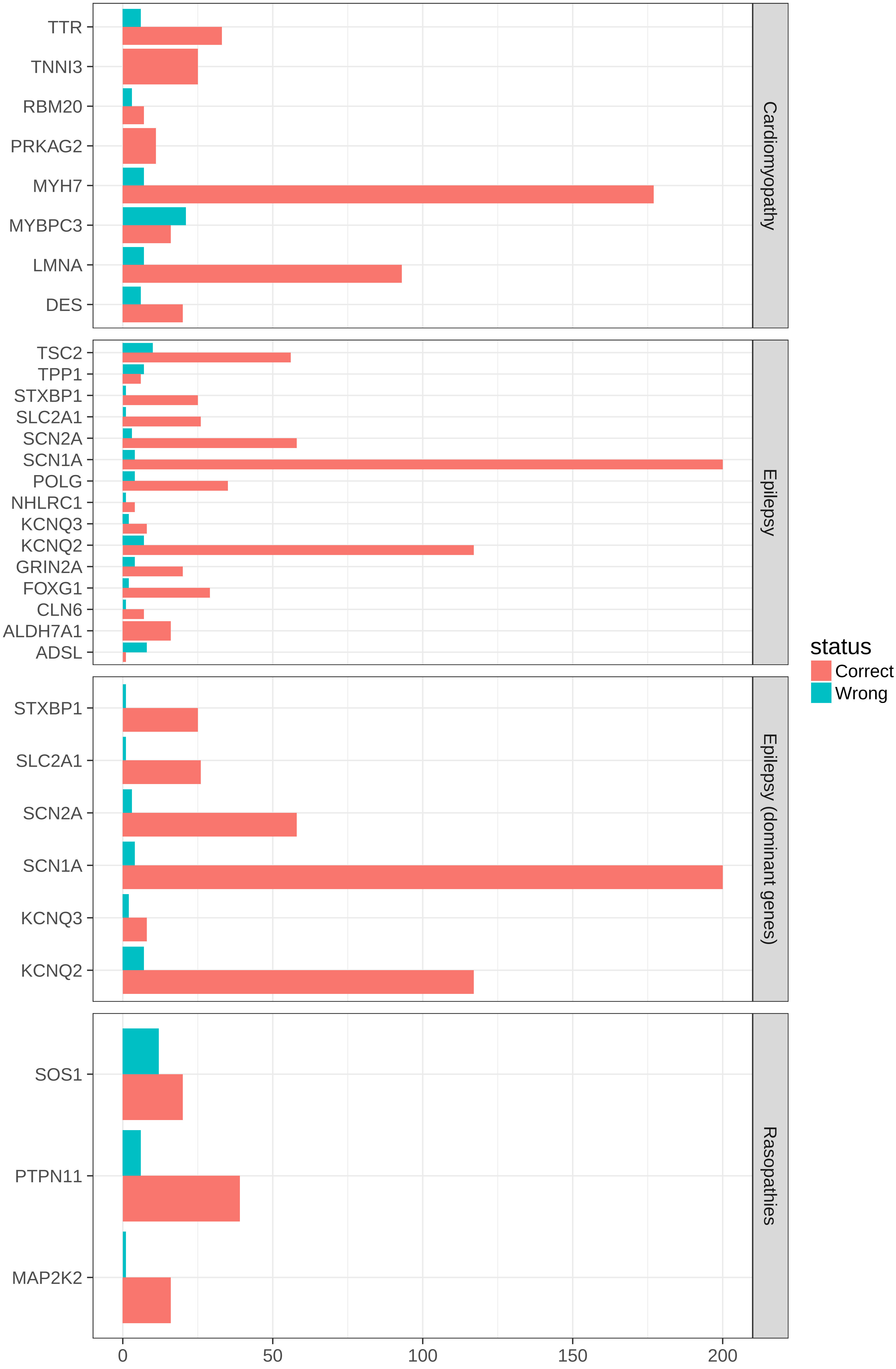

### SLC2A1.clinvar.lolly.png

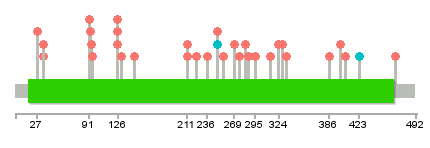

### SLC2A1.denovodb.lolly.png

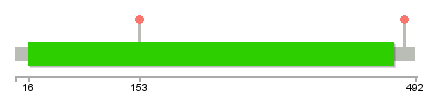

### SPTAN1.clinvar.lolly.png

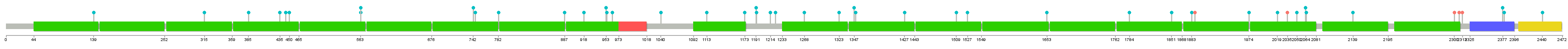

### SPTAN1.denovodb.lolly.png

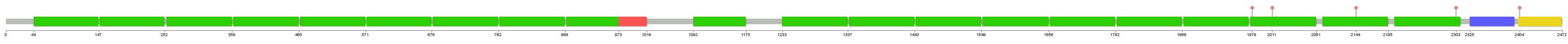

### STXBP1.clinvar.lolly.png

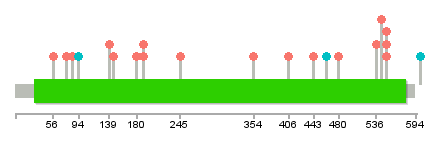

### STXBP1.denovo_db.lolly.png

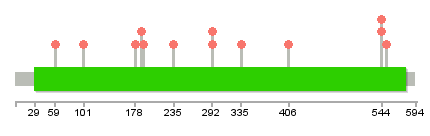

### TSC1.clinvar.lolly.png

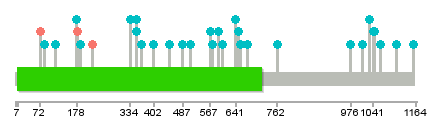

### TSC1.denovodb.lolly.png

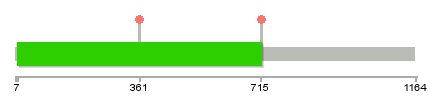

### uc_mpc_eval.png

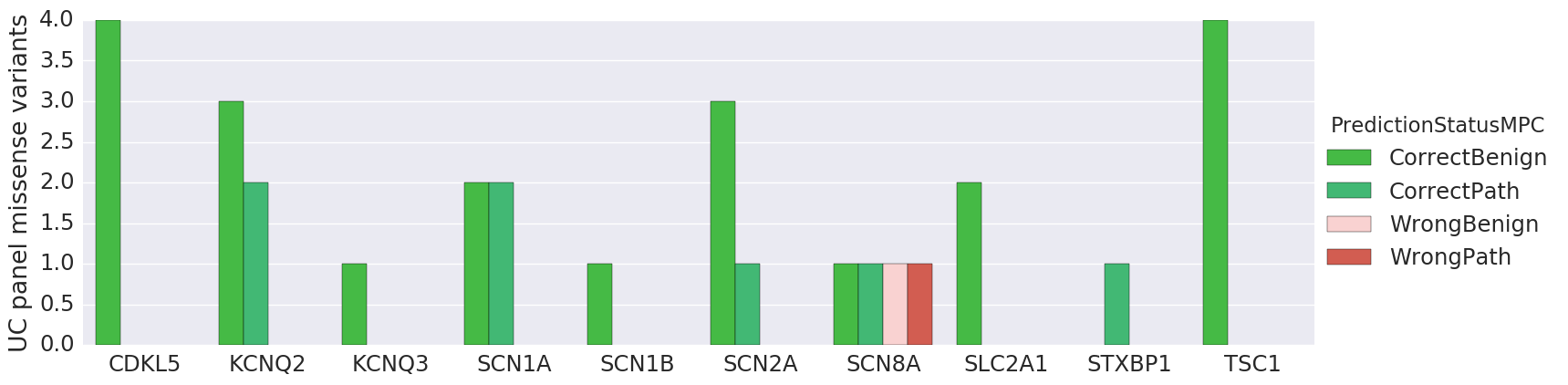
